# PKN2 deficiency leads both to prenatal ‘congenital’ cardiomyopathy and defective angiotensin II stress responses

**DOI:** 10.1101/2022.05.24.493130

**Authors:** Jacqueline J T Marshall, Joshua J Cull, Hajed O Alharbi, May Zaw Thin, Susanna TE Cooper, Christopher Barrington, Hannah Vanyai, Thomas Snoeks, Bernard Siow, Alejandro Suáarez-Bonnet, Eleanor Herbert, Daniel J Stuckey, Angus Cameron, Fabrice Prin, Andrew C. Cook, Simon L Priestnall, Sonia Chotani, Owen J L Rackham, Daniel N Meijles, Tim Mohun, Angela Clerk, Peter J Parker

## Abstract

**Background:** The protein kinase PKN2 is required for embryonic development, and PKN2 knockout mice die as a result of failure in expansion of mesoderm tissues, cardiac development and neural tube closure. In the adult, cardiomyocyte PKN2 and PKN1 (in combination) are required for cardiac adaptation to pressure-overload. The role of PKN2 in contractile cardiomyocytes during development and its role in the adult heart remain to be fully established.

**Methods:** We used mice with cardiomyocyte-directed knockout of PKN2 or global PKN2 haploinsufficiency. Cardiac function and dimensions were assessed with high resolution episcopic microscopy, MRI, micro-CT and echocardiography. Biochemical and histological changes were assessed.

**Results:** Cardiomyocyte-directed PKN2 knockout embryos displayed striking abnormalities in the compact myocardium, with frequent myocardial clefts and diverticula, ventricular septal defects and abnormal heart shape. The sub-Mendelian homozygous knockout survivors developed cardiac failure. RNASeq data showed upregulation of PKN2 in patients with dilated cardiomyopathy, suggesting an involvement in adult heart disease. Given the rarity of homozygous survivors with cardiomyocyte-specific deletion of PKN2, this was explored using mice with constitutive heterozygous PKN2 knockout. Cardiac hypertrophy resulting from hypertension induced by angiotensin II was reduced in haploinsufficient PKN2 mice relative to wild-type littermates, with suppression of cardiomyocyte hypertrophy and cardiac fibrosis.

**Conclusions:** Cardiomyocyte PKN2 is essential for heart development and formation of compact myocardium, and is also required for cardiac hypertrophy in hypertension. Thus, PKN signalling may offer therapeutic options for managing congenital and adult heart diseases.

## INTRODUCTION

Heart disease, a major cause of death and disability worldwide, develops from numerous causes including genetic/environmental interactions resulting in congenital cardiac defects,^1^ in addition to diseases in later life resulting from various pathophysiological stressors.^2-4^ The heart contains three main cell types (cardiomyocytes, endothelial cells, fibroblasts) with cardiomyocytes providing the contractile force. In the embryo/foetus, cardiomyocytes proliferate whilst the heart develops, but withdraw from the cell cycle in the perinatal/postnatal period, becoming terminally-differentiated.^5^ Further growth of the heart to the adult size requires an increase in size and sarcomeric/myofibrillar apparatus of individual cardiomyocytes (maturational growth). The adult heart experiences pathophysiological stresses (e.g. hypertension) requiring an increase in contractile function. This is accommodated by cardiomyocyte hypertrophy (sarcomeric replication in parallel or series) with associated cardiac hypertrophy (enlargement of the heart).^6^ This adaptation is initially beneficial, but pathological hypertrophy develops with cardiomyocyte dysfunction and death, loss of capillaries and deposition of inelastic fibrotic scar tissue.^6^ These processes are all regulated by a complex interplay of intracellular signalling pathways, driven by numerous protein kinases that play central roles both in mammalian development and in adult tissue homeostasis.^7^ These regulatory proteins offer themselves as potential targets for intervention both in management of congenital cardiomyopathies and in pathological states. Insight into the key regulatory players at each and every stage is central for prevention, in addition to the development and delivery of improved treatments and outcomes.

The Protein Kinase N (PKN) family of kinases are emerging as potential therapeutic targets for heart disease.^8^ Whilst PKN1 and PKN2 are ubiquitously expressed in tissues throughout the body, PKN3 is expressed in a smaller subset of tissues, especially endothelial cell types.^9, 10^ Only *Pkn2* knockout is embryonic lethal. This is due to failure in the expansion of mesoderm tissues, failure of cardiac development and compromised neural tube closure.^11, 12^ Further studies with conditional knockouts of *Pkn2* employed cell-targeted Cre under the control of a smooth muscle protein 22α (SM22α) promoter.^12^ SM22α (and therefore SM22α-Cre) is expressed in the heart tube from embryonic day E7.5/8,^13^ with expression declining from E10.5 becoming restricted to the right ventricle by E12.5. By E13.5, SM22α is undetectable in the heart^14^ and expression is subsequently confined to smooth muscle cells and myofibroblasts.^13, 15-17^ Conditional gene deletion of *Pkn2* results in sub-mendelian survival of *SM22α-Cre*^*+/-*^ *Pkn2*^*fl/fl*^ offspring, with approximately 1/3 of mice surviving to 4 weeks postnatally.^12^ These data indicate that PKN2 is not only important in the heart during embryonic development, but (whilst the phenotype is not fully penetrant in the *SM22α-Cre* model) is also required for maturational growth of the heart. This raises the question of what is compromised and how loss of PKN2 manifests in the adult.

There are few studies of PKNs in the adult heart. *In vivo* studies in mice suggest there may be redundancy between PKN1 and PKN2 in cardiomyocytes, and double knockout of both kinases simultaneously in cardiomyocytes inhibits cardiac hypertrophy in pressure-overload conditions induced by transverse aortic constriction or angiotensin II (AngII).^20^ Fundamental questions remain concerning the functional redundancy of PKN1 and 2 in cardiomyocytes. Here, we demonstrate that the loss of PKN2 in cardiomyocytes has a catastrophic effect on ventricular myocardial development, suggesting that alterations in PKN2 signalling may contribute to congenital cardiac problems/cardiomyopathy. We also demonstrate that PKN2 haploinsufficiency compromises cardiac adaptation to hypertension in adult mouse hearts.

## METHODS

### Data availability

Details of experimental procedures are in the Data Supplement. Data, methods and study materials are available on request from the corresponding authors.

## RESULTS

### Cardiac-specific knockout of *Pkn2*

Evidence from the *SM22α-Cre* conditional *Pkn2* knockout indicates reduced survival (Supplementary Table S1 and ^12^), which might reflect in part an impact on heart function but may also be determined by loss of *Pkn2* expression in smooth muscle. To dissect the functional contributions more selectively, we sought to refine the pattern of knockout by using the *XMLC-Cre* line, where *Cre* expression is cardiac restricted.^22^ Genotyping pups from the *Pkn2*^*fl/fl*^ mice crossed with XMLC2-Cre^+/-^ *Pkn2*^*fl/+*^ animals at 2-4 weeks of age identified only one *XMLC2-Cre*^*+/-*^ *Pkn2*^*fl/fl*^ mouse (of 126), consistent with a severe phenotype with this Cre line (Supplementary Table S1). This potentially results from higher efficiency of *XMLC2-Cre* activity in cardiomyocytes (95%^22^) compared with *SM22α-Cre* (75-80%^15^) and more specifically indicates that there are cardiac-associated phenotypes of PKN2 loss. The single surviving male was small (11.2 g relative to 22.4g for a *XMLC2-Cre*^*+/-*^ *Pkn2*^*fl/+*^ littermate) and was culled at 5 weeks due to poor condition. Histological analysis of the heart from this animal showed that it was highly abnormal. Both ventricles were dilated, with a hypertrophic right ventricle and thin-walled (hypotrophic) and disorganised left ventricular myocardium and interventricular septum. There were partial discontinuities in the cardiac muscle of the compact layer, and sections with highly disorganised cardiomyocytes and fibrosis (Figure 1a). It was noted also that the lungs from the *XMLC2-Cre*^*+/-*^ *Pkn2*^*fl/fl*^ mouse displayed grossly enlarged alveolar spaces (Figure 1b).

**Figure 1.**
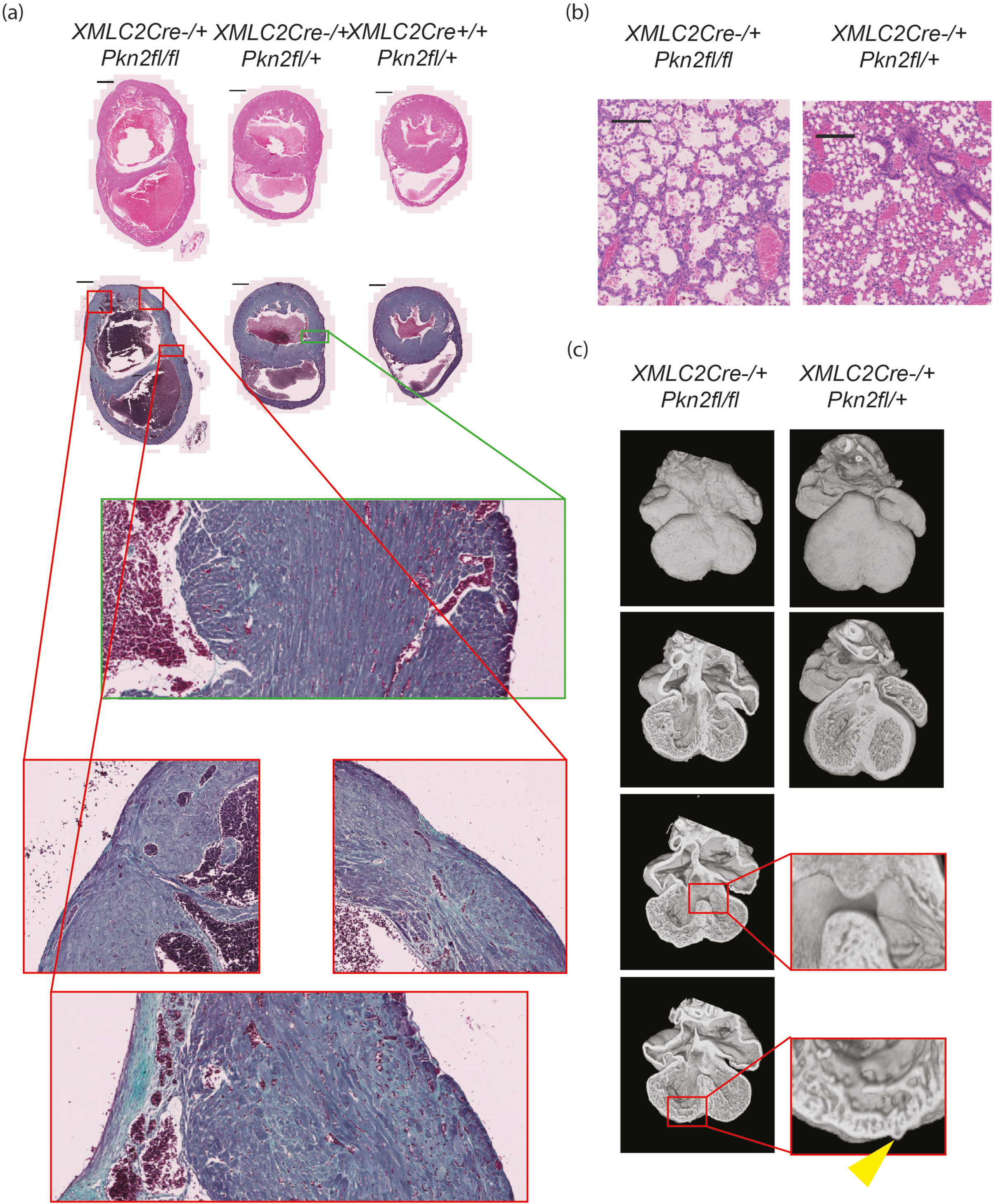
PKN2 knockout in cardiomyocytes causes defective embryonic heart development leading to failure prior to adulthood. **(a)** H&E and Gomori’s Trichrome stained histological cardiac short-axis sections of the longest surviving *XMLC2Cre*^*+/-*^ *Pkn2*^*fl/fl*^ genotype mouse (left; male), and two littermates (middle & right; male *XMLC2Cre*^*+/-*^ *Pkn2*^*fl/+*^ *&* female *XMLC2Cre*^*+/+*^ *Pkn2*^*fl/fl*^, respectively) all culled aged 5 weeks. Scale bars are 1 mm. In lower boxed regions, various sections are also shown at 20x zoom relative to whole-heart cross-sections. **(b)** H&E sections of lungs of the same *XMLC2Cre*^*+/-*^ *Pkn2*^*fl/fl*^ genotype mouse (left), and its *XMLC2Cre*^*+/-*^ *Pkn2*^*fl/+*^ littermate (right; both male). Scale bars are 200 µm. **(c)** Images of High-Resolution Episcopic Microscopy (HREM) reconstructions of E14.5 embryo hearts for the genotypes indicated, showing surface and four chamber views. Boxed regions of sections are shown at 4x zoom; a diverticulum is indicated by the yellow arrowhead.

To explore this impact of PKN2 further, we genotyped gestational day 14.5 embryos from *Pkn2*^*fl/fl*^ mice crossed with *XMLC2-Cre*^*+/-*^ *Pkn2*^*fl/+*^ and found that 6 of 24 were *XMLC2-Cre*^*+/-*^ *Pkn2*^*fl/fl*^ consistent with a Mendelian distribution at this developmental stage. However, analysis of the hearts using high resolution episcopic microscopy (HREM) showed various abnormalities (Figure 1c; Video 1; Figure S1). Notably, hearts from these mouse embryos displayed multiple surface nodules (diverticula; see below) distributed across both ventricles, unusually large perimembranous ventricular septal defects and thin compact myocardium indicative of significant cardiac developmental complications.

### Characterising the cardiac requirement of PKN2

Whilst the *XMLC*-driven knockout of *Pkn2* clearly demonstrated a cardiac phenotype associated with PKN2 loss, the fatal consequences of the XMLC-driven knockout of Pkn*2* led us instead to investigate in more detail the more frequent survivors derived in the *SM22α-Cre* model. On re-derivation of the *SM22α* model into a new facility and crossing the *Pkn2*^*/fl*^ with *SM22αCre*^*+/-*^ *Pkn22*^*fl/+*^ animals, we found there remained a sub-Mendelian distribution of the *SM22α*^*+/-*^ *Pkn2*^*fl/fl*^ genotype (28% of expected numbers with no male/female bias; Supplementary Table S1). Crossing 10 of these mice with P*kn2*^*fl/fl*^ animals produced 62 weaned pups but again a sub-Mendelian representation of the *SM22α*^*+/-*^ P*kn2*^*fl/fl*^ genotype (32% of expected).

Typically, surviving *SM22α*^*+/-*^ *Pkn2*^*fl/fl*^ mice became overtly unwell as they aged, displaying a range of adverse phenotypes including body weight differences compared to littermates, loss of condition, hunched appearance or reduced activity. Amongst a cohort of 23 animals, there were 50% asymptomatic *SM22α-Cre*^*+/-*^ *Pkn2*^*fl/fl*^ mice at 24 weeks of age, with the oldest two mice reaching 72 weeks and only then started to display phenotypes (echocardiography showed aortic valve stenosis and echo-dropout across the long-axis view of the left ventricle in one, suggestive of mitral annular calcification; see below). A typical example of an ageing *SM22α-Cre*^*+/-*^ *Pkn2*^*fl/fl*^ animal was a 38-week-old fertile female which upon culling due to loss of condition displayed a heart weight: body weight ratio = 0.99% (compared to 0.37-0.48% for female littermates).

Similar to the single weaned *XMLC2-Cre*^*+/-*^ *Pkn2*^*fl/fl*^ mouse, histology of the *SM22α-Cre*^*+/-*^ *Pkn2*^*fl/fl*^ mice showed hypertrophic ventricular walls with disorganised cardiomyocytes and extensive fibrosis, consistent with a form of cardiomyopathy and heart failure (Figure 2a,b; Figure S2). Histological assessment of the lungs showed *SM22α-Cre*^*+/-*^ *Pkn2*^*fl/fl*^ mice culled with breathlessness had grossly enlarged alveolar spaces (Figure 2c), corresponding to the anecdotal finding in the one *XMLC-Cre*^*+/-*^ *Pkn2*^*fl/fl*^ survivor. The timing of symptom onset is illustrated in Figure 2d. Functional cardiac MRI analysis of a further set of five surviving *SM22α-Cre*^*+/-*^ *Pkn2*^*fl/fl*^ mice with initial symptoms of heart failure, indicated that they had reduced left ventricular ejection fraction (Figure 2e,f). Imaging did not identify a single unifying cause, with examples of reduced right ventricular mass, potentially associated with pulmonary hypertension (n=2 of 5), or completely abnormal architecture with hearts exhibiting a bulbous shape (n=3 of 5) (see histology and HREM in Figures S2 and S3).

**Figure 2.**
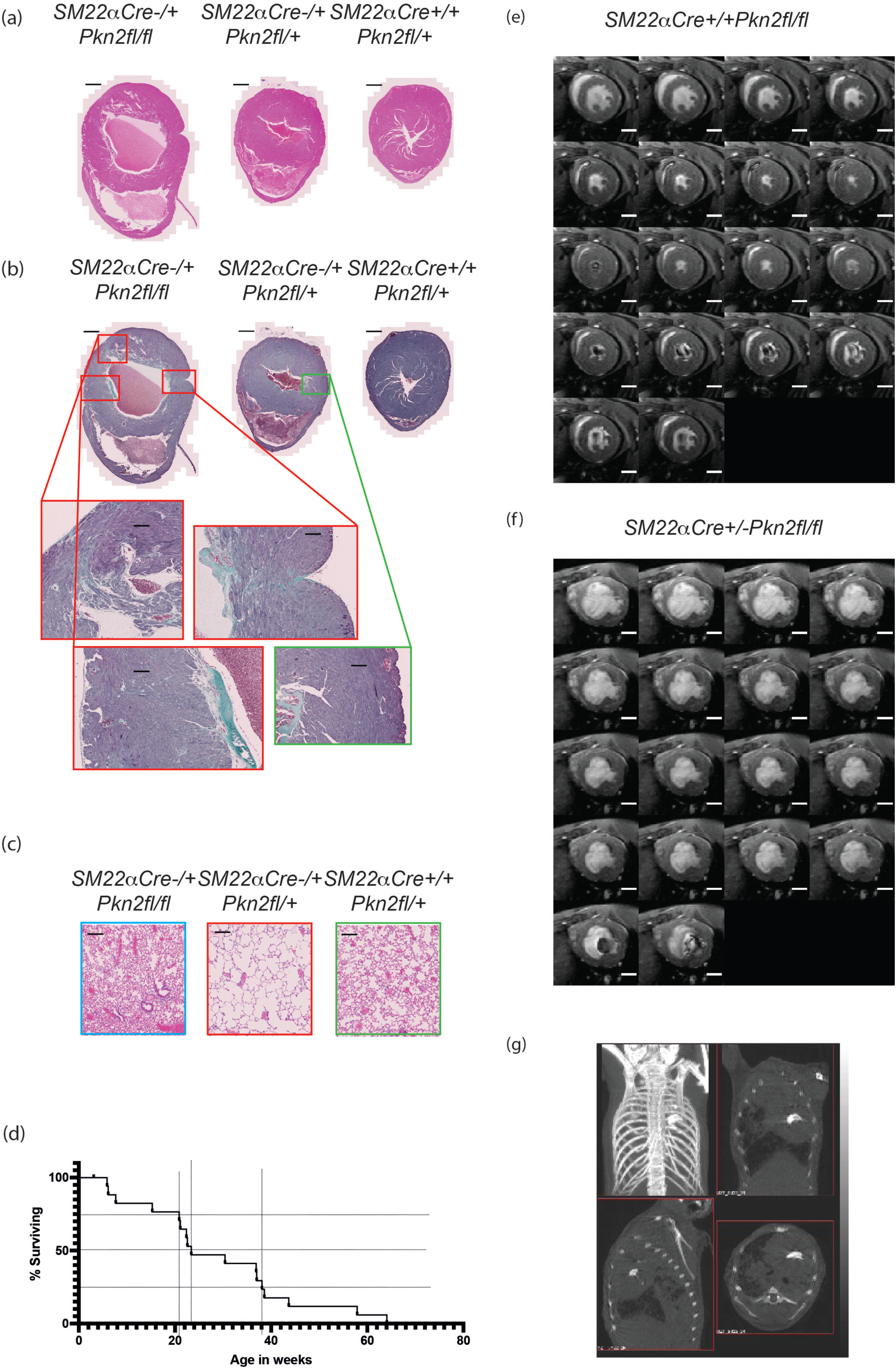
*SM22aCre*^*+/-*^ *Pkn2*^*fl/+*^ mice suffer cardiac failure on aging. **(a**,**b)** Short-axis mid-ventricle sections of hearts of 39 week male littermates of the genotypes indicated, stained with H&E (a), or Gomori’s Trichrome (b). **(c)** H&E stained sections of lungs, littermates as in (a,b). **(d)** Time course of symptom onset for mice of genotype *SM22αCre*^*+/-*^ *Pkn2*^*fl/fl*^ (n=23). **(e**,**f)** Montage of time-series through the cardiac cycle recorded by cine-cardiac-MRI through the short-axis plane at mid-ventricle from *SM22αCre*^*+/+*^ *Pkn2*^*fl/fl*^ (e) and *SM22αCre*^*+/-*^ *Pkn2*^*fl/fl*^ (f) 32 week old female littermates. **(g)** micro-CT of the thorax of a 24 week *SM22αCre*^*+/-*^ *Pkn2*^*fl/fl*^ female shown as a projected image and slice images in 3 planes. Scale-bars in histopathology images are 1 mm in whole heart sections (panels a and b) and 200 µm in zoomed sections (panels b and c). Scale-bars in MRI montages are 2.6 mm (panels e and f).

Micro-CT imaging was used to assess disease in a further subset of survivors. This revealed variously: pulmonary oedema and pleural effusion, and marked, unusual calcification in the centre of the rib cage (Figure 2g). Subsequent micro-CT of fixed hearts showed that calcification appeared to be associated with both the aortic and mitral valves (Video 2). The evidence suggests that surviving *SM22α-Cre*^*+/-*^ *Pkn2*^*fl/fl*^ mice had congenital defects in the heart and/or lungs resulting in heart failure and as with human congenital heart disease, there was heterogeneity with respect to age of disease onset, severity of disease and specific cardiac phenotype.

### Cardiomyocyte PKN2 is essential for normal cardiac development

Although previous studies reported that PKN2 is essential for embryonic development, E13.5 embryos were produced at a normal Mendelian ratio.^12^ Data from surviving *SM22α-Cre*^*+/-*^ *Pkn2*^*fl/fl*^ mice (above) suggest the cardiac defect is most likely the result of developmental cardiac abnormalities, however defects in lung development resulting from loss of Pkn2 in smooth muscle cells would impact cardiac function and also lead to heart failure. We therefore collected embryos at E14.5-E18.5 generated by crossing *Pkn2*^*fl/fl*^ and *SM22α-Cre*^*+/-*^ *Pkn2*^*fl/+*^ mice for further analysis to examine the Mendelian ratios at these later embryonic stages, and to enable pathological examination of the developing hearts and lungs. Genotype distribution was Mendelian at both E14.5 and E18.5 (Supplementary Table S1). Notably, histological analysis failed to identify any difference between lungs from *SM22α-Cre*^*+/-*^ *Pkn2*^*fl/fl*^ embryos and those from littermates that were either *Cre* negative or heterozygous for the floxed *Pkn2* allele or both (Figure 3a), suggesting normal lung development to E18.5 and implying that the enlarged alveolar spaces in some of the rare surviving adults may be a secondary consequence of cardiac abnormalities.

**Figure 3.**
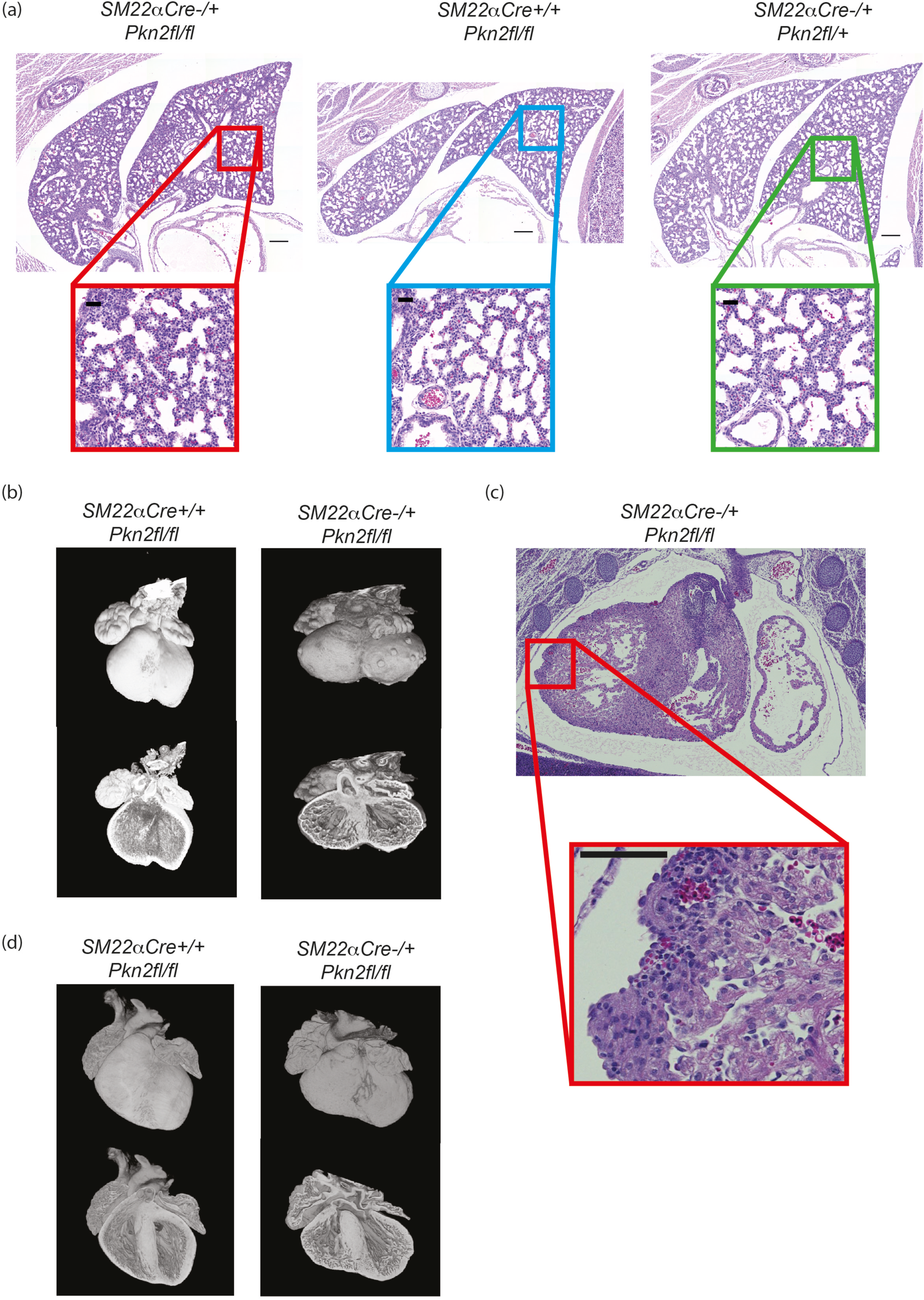
The heart-specific defects in *SM22αCre*^*+/-*^*Pkn2*^*fl/fl*^ mice. **(a)** H&E stained sections of (E18.5) lungs of littermates with the indicated genotypes. Lower boxed panels are 5x zoomed; scale bars indicate 200 µm in upper panels and 40 µm in lower zoomed panels. (**b**) Reconstructions of E14.5 hearts imaged by HREM are shown in surface (top images) and slice (lower images) views. The genotypes are as indicated. (**c**) H&E stained E14.5 heart sections featuring an outer ventricular surface nodule, identified as a diverticulum. Scale bars are 100 µm. **(d)** Reconstructions of E18.5 hearts imaged by HREM shown as per panel (b). Genotypes are as indicated.

Given that the E18.5 genotype distribution was Mendelian but only ∼30% of *SM22α-Cre*^*+/-*^ *Pkn2*^*fl/fl*^ mice survived to weaning, this suggested that lethality occurred in the perinatal period. Analysis of deceased pups including part-cannibalised carcasses demonstrated that this genotype was selectively lost in this very early neonatal stage (Supplementary Table S1). Veterinary pathologist characterisation of P1.5-P5.5 carcasses showed that all had normal palates, lungs that floated and milk spots. There were no observations of pericardial bleeding, which might have been observed if one of the cardiac diverticula had ruptured causing tamponade (see below).

Analysis of embryo hearts using HREM showed that of 16 *SM22α-Cre*^*+/-*^ *Pkn2*^*fl/fl*^ E14.5 embryos, 15 had overt defects in cardiac development compared with littermates (exemplified in Figure 3b; see also Supplementary Figure S3). Defects included perimembranous ventricular septal defects (VSDs), small muscular VSDs (VSDs), thin compact myocardium (right and left ventricle) and overt nodules on the external surface of either or both ventricles. The degree of phenotype varied, with the most abnormal hearts showing perimembraneous VSD with overriding of the aorta, and many showing an overall abnormally squat shape with an indistinct apex. The nodules on the surface of the ventricles were examined histologically and identified as diverticula with internal lumens connected to the ventricular cavities (Figure 3c).

Cardiac septation would expect to be completed by E15, however of 60 embryos analysed in crosses of *Pkn2*^*fl/fl*^ and *SM22α-Cre*^*+/-*^ *Pkn2*^*fl/+*^ mice, we obtained 18 *SM22α-Cre*^*+/-*^ *Pkn2*^*fl/fl*^ embryos of which seven had persistent VSDs long with thin compact myocardium/ventricular walls. The nodules on the ventricle surfaces apparent at E14.5 also persisted. There was additional abnormal development of the trabecular layer in the ventricular walls – analogous to hypertrabeculation (Figure 3d).

The congruence of developmental defects in the *SM22α-Cre* and the *XMLC-Cre* strains indicates that the dominant effect of tissue-specific *Pkn2* loss relates to the shared aspect between these models, namely an impact on cardiac myocytes rather than, for example any later embryonic stage stromal loss of *Pkn2*. Although there is the potential for additional influences of the *SM22α-Cre Pkn2* knockout model via vascular smooth muscle, this is unlikely to have any profound impact given the phenotype of the XMLC-Cre strain is more penetrant not less so. It is surmised that this reflects the higher efficiency of this latter promoter in cardiac myocytes ^22^. These observations of developmental abnormalities in the heart are consistent with the conclusion that the failure of these *Cre*^*+/-*^ *Pkn2*^*fl/fl*^ mice to thrive, and in particular the cardiac abnormalities in the rare survivors, reflect congenital problems. The rarity of these survivors and the legacy of the developmental defects compromise the assessment of the role of PKN2 in adults in these models and alternative strategies are required.

### Expression of PKN2 in the adult heart

Pkn2 is expressed in adult cardiomyocytes.^7^ Expression relative to total protein declines during postnatal development due to the substantial increase in cardiomyocyte size, but the relative amount of PKN2 per cell increases in the adult (Supplementary Figure S4). To determine if variations in *PKN2* expression are associated with human heart failure, we mined an RNASeq database of patients with dilated cardiomyopathy *vs* normal controls.^21^ All PKN isoforms were detected in human hearts, but *PKN2* expression increased in dilated cardiomyopathy hearts relative to controls, whilst *PKN1* and *PKN3* declined (Figure 4a). To determine if PKN2 may be involved in disease aetiology, we assessed expression in a mouse model of hypertension induced by AngII (0.8 mg/kg/d, 7 d). Expression of PKN2 (not PKN1) increased with AngII treatment relative to vehicle-treated controls (Figure 4b). Thus, altered expression may contribute to the adaptive response to hypertension and/or may be associated with progression towards a pathological state. To assess whether altered expression might impact this response and in view of the impact of *Pkn2* knockout as described above, we focused on the effects of haploinsufficiency (global loss of a single allele, i.e. *Pkn2*Het mice). PKN2 protein expression was reduced in hearts from male *Pkn2*Het mice relative to wild-type (WT) littermates (Figure 4c). There was no compensatory increase in PKN1 expression although the relative level of phosphorylation of PKN2 was increased. *Pkn2*Het mice thus provide a model for assessment of altered expression in the adult heart.

**Figure 4.**
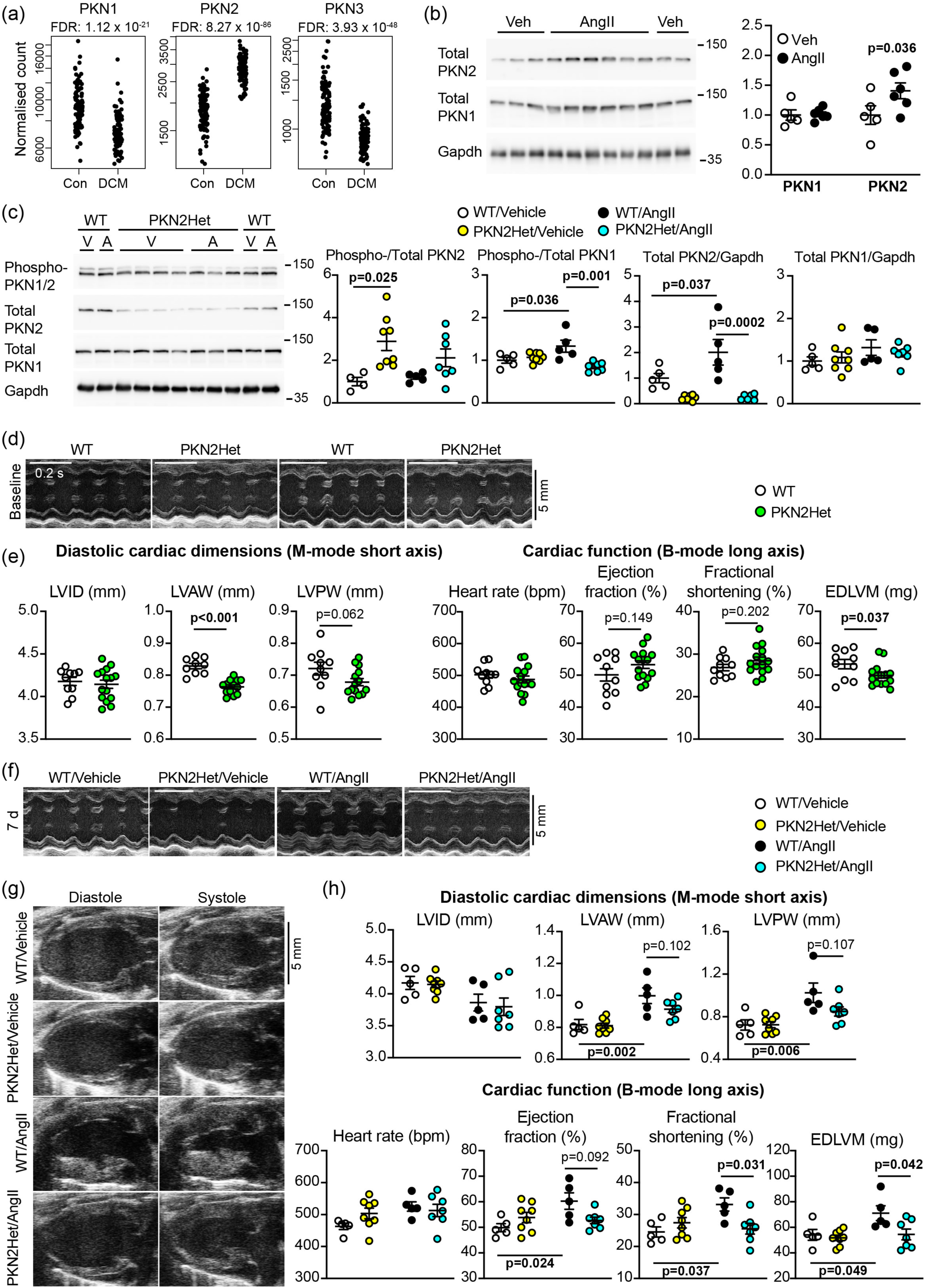
PKN2 is associated with human heart failure and required for cardiac adaptation to hypertension in mice. (**a**), Expression of *PKN1, PKN2* and *PKN3* in human hearts. Data were from an RNASeq database of patients with dilated cardiomyopathy (DCM; n=97) and normal controls (Con; n=108). (**b**,**c**), Immunoblotting of phospho-PKN1/2, total PKN1, total PKN2 and GAPDH in hearts from wild-type (WT) or *Pkn2*Het mice treated for 7d with vehicle (Veh, V) or 0.8 mg/kg/d angiotensin II (AngII, A). Immunoblots are on the left with densitometric analysis on the right (normalised to the mean of vehicle-treated controls). Analysis used 2-way ANOVA with Holm-Sidak’s post-test. (**d-h**), Echocardiography of hearts from WT and *Pkn2*Het mice at baseline (**d**,**e**) or treated with vehicle or AngII (7d) (**f-h**). Representative short-axis M-mode images used for assessment of cardiac dimensions are shown at baseline (**d**) and after treatment (**f**) (the same animals are shown). (**g**), Representative long-axis B-mode images used for speckle-tracking and strain analysis to assess cardiac function are shown (7d). (**e**,**h**), Echocardiograms were analysed. Additional data are in Supplementary Tables S2 and S3. Analysis used unpaired, two-tailed t tests (**e**) or 2-way ANOVA with Holm-Sidak’s post-test (**g**). LV, Left ventricle; ID, internal diameter; AW, anterior wall; PW, posterior wall; EDLVM, end diastolic LV mass. (**b**,**c**,**e**,**h**) Individual data points are provided with means ± SEM.

### The role of PKN2 in the adult heart

We assessed baseline dimensions and function of the hearts from *Pkn2*Het mice and WT littermates using echocardiography (Figure 4d and 4e; Supplementary Table S2). M-mode imaging of the short-axis view revealed that *Pkn2*Het mice had a small but significant reduction in left ventricle (LV) wall thickness compared to WT littermates. Assessment of cardiac function using speckle-tracking strain analysis confirmed that overall LV mass was significantly decreased in *Pkn2*Het mice and there was a small, albeit non-significant, increase in ejection fraction (Figure 4e; Supplementary Table S2). We conclude that there are abnormalities in surviving *Pkn2*Het mice and, although these changes appear relatively minor, they may compromise cardiac adaptation to pathophysiological stresses such as hypertension.

To determine if the hearts from *Pkn2*Het mice can adapt to hypertension, adult male *Pkn2*Het mice or WT littermates (11-14 weeks) were treated with AngII or vehicle for 7d and cardiac dimensions/function were assessed by echocardiography (Figure 4f-h; Supplementary Table S3). As in previous studies,^23, 24^ AngII promoted cardiac hypertrophy in WT mice, with decreased LV internal diameter and significantly increased LV wall thickness (Figure 4f,h). AngII promoted similar changes in *Pkn2*Het mice, although ventricular wall thickening appeared reduced with no significant difference relative to vehicle-treated mice (Figure 4h). Strain analysis of B-mode images confirmed that the increase in LV mass induced by AngII was attenuated in *Pkn2*Het mice (Figure 4g,h). In addition, AngII significantly increased ejection fraction and fractional shortening in WT mice, but not *Pkn2*Het mice, indicating that cardiac adaptation to hypertension in response to AngII was attenuated. Histological staining showed that AngII increased cardiomyocyte cross-sectional area in WT mice, but not in *Pkn2*Het mice (Figure 5a). AngII increased cardiac fibrosis in interstitial areas of the myocardium, particularly at the junctions between the outer LV wall and the interventricular septum, but this was similar in both WT and *Pkn2*Het mice (Figure 5b). AngII also increased fibrosis in the perivascular regions of arteries/arterioles, and this was reduced in *Pkn2*Het hearts compared with hearts from WT mice (Figure 5c). Overall, cardiac adaptation to hypertension was reduced in *Pkn2*Het mice compared with WT littermates, with both reduced cardiomyocyte hypertrophy and perivascular fibrosis.

**Figure 5.**
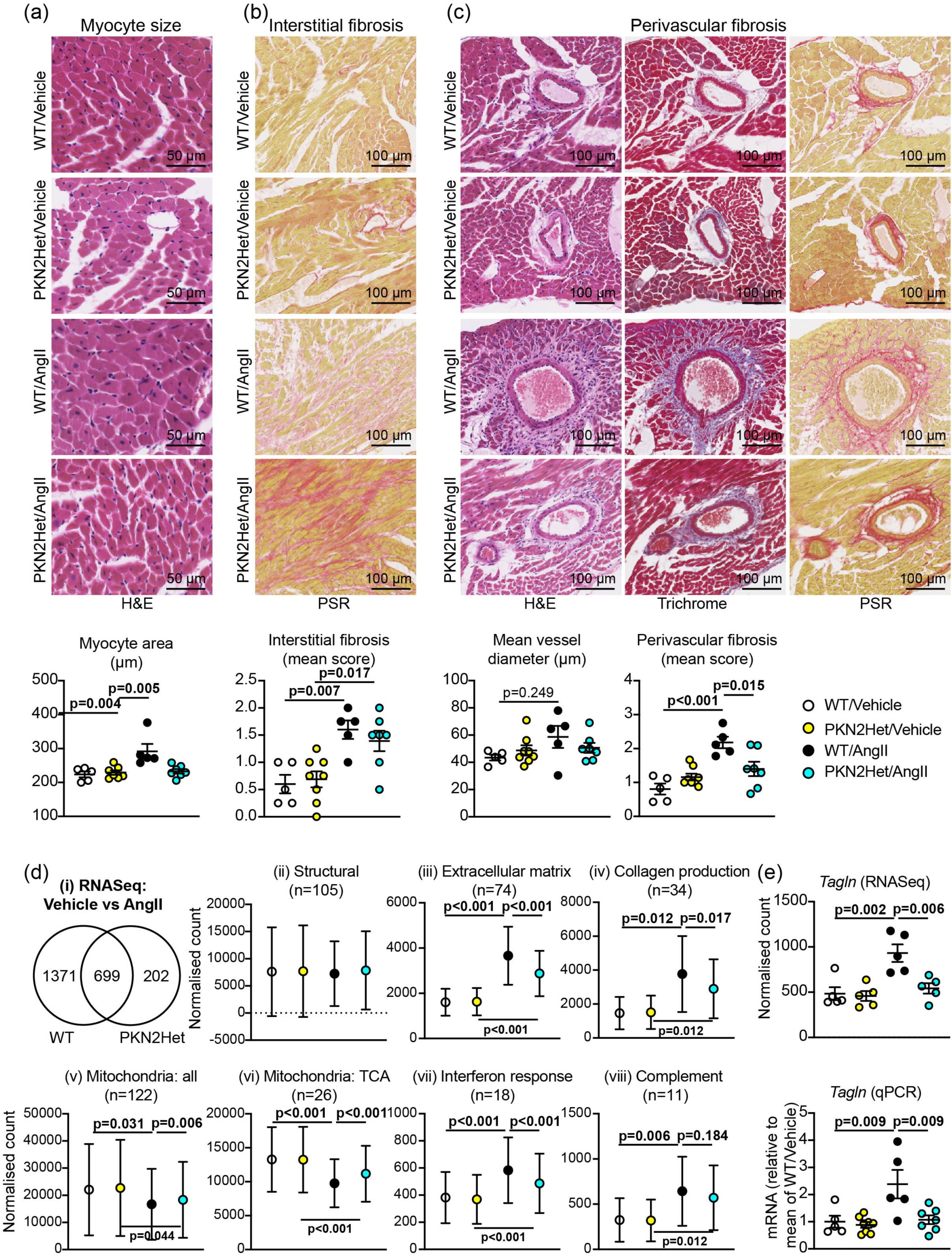
*Pkn2* haploinsufficiency reduces cardiac adaptation to hypertension, affecting cardiomyocyte size, perivascular fibrosis and gene expression. WT or *Pkn2*Het mice were treated with vehicle or AngII (0.8 mg/kg/d, 7d). (**a-c**), Sections of mouse hearts were stained with haematoxylin and eosin (H&E), picrosirius red (PSR), or Masson’s Trichrome. Representative images are shown with quantification below. Analysis used 2-way ANOVA with Holm-Sidak’s post-test. (**d**), RNASeq of hearts from WT and *Pkn2*Het mice. (i) Summary of differentially-expressed genes (DEGs; p<0.01) resulting from AngII treatment. (ii)-(viii) Clusters of DEGs according to function. Results are the mean normalised count values with 95% CI for the n values indicated. Analysis used 1-way ANOVA with Holm-Sidak’s post-test. (**e**), Comparison of *Tagln* (*SM22α*) mRNA expression assessed by RNASeq and qPCR. (**a-c**,**e**) Individual data points are shown with means ± SEM. Analysis used 2-way ANOVA with Holm-Sidak’s post-test.

To gain mechanistic insight into the effects of PKN2 in the cardiac response to hypertension, we used RNASeq to assess the transcriptional differences in the AngII response of hearts from *Pkn2*Het mice compared with WT littermates (Figure 5d). No differentially-expressed genes (DEGs) were identified when comparing *Pkn2*Het and WT hearts from mice treated with either vehicle or AngII. AngII-treatment resulted in 2272 DEGs in hearts from WT or *Pkn2*Het mice (p<0.01): 699 were identified in both genotypes, 1371 were only detected in WT hearts and 202 were only detected in *Pkn2*Het hearts (Figure 5d(i), Supplementary Tables S4-S9). Clustering the DEGs according to function highlighted significant changes in a subset of genes for the myofibrillar apparatus and cytoskeletal structures, particularly the actin cytoskeleton, but there were no overall differences between the genotypes/treatment in these gene classes as a whole (Figure 5d(ii)). In contrast, genes associated with fibrosis were significantly upregulated by AngII (Figure 5d(iii)), particularly those associated with collagen production (Figure 5d(iv)). The response in hearts from *Pkn2*Het mice was reduced relative to WT littermates, consistent with a reduction in perivascular fibrosis seen by histology (Figure 5c). Another feature of the AngII response was the reduction in expression of genes for mitochondrial proteins (Figure 5d(v)). The overall response was reduced in *Pkn2*Het mice, but the effect was more pronounced for some genes, particularly those of the tricarboxylic acid (TCA) cycle (Figure 5d(vi)). AngII cardiac hypertrophy is associated with inflammation and we detected a clear interferon response (Figure 5d(vii)) with upregulation of the complement pathway (Figure 5d(viii)). Both of these responses were reduced in hearts from *Pkn2*Het mice relative to WT littermates. Data for individual genes in each of the clusters are in Supplementary Table S10. We mined the data for classical markers of cardiac hypertrophy and detected increased expression of *Myh7, Nppa* and *Nppb* with AngII as expected^25^, but expression was similar in *Pkn2*Het and WT mice (Supplementary Figure S5). We also detected upregulation of *Tagln* (SM22α) by AngII in WT, but not *Pkn2*Het, mouse hearts (Figure 5e). *SM22α* is a marker of smooth muscle cells in the adult, and this is potentially a reflection of the perivascular fibrosis induced by AngII around arteries/arterioles.

The importance of PKN2 in cardiac development and disease led us to investigate what the consequences might be with aging, comparing echocardiography data for male mice with an average age of 12 weeks with those obtained from mice with an average age of 42 weeks. As expected, hearts from the older mice had a significantly greater LV mass than the 12 week animals, but there was no difference between WT and *Pkn2*Het mice (Figure 6a). The largest diameter of the aortae (measured during cardiac systole) increased with age to some extent in both genotypes, but the distensibility of the aorta was compromised in the older mice as shown by a reduction in the ratio between the diameter measured at cardiac systole and after the aortic valve has closed when the diameter is at its narrowest (Figure 6b). This appeared to be mitigated to some degree in the *Pkn2*Het mice. Pulsed-wave Doppler was used to assess blood flow from the heart into the aorta and the pulmonary artery. We detected no differences in blood flow in either vessel in the young mice (Figure 6c-f; Supplementary Table S2). Older WT and *Pkn2*Het mice had reduced pulmonary velocity time interval, with a reduction in velocity and gradient, and the degree of change was similar (Figure 6c,d; Supplementary Table S2). In WT mice, the aortic velocity time interval, along with the mean/peak velocity and gradient were all significantly reduced in older mice compared with the young mice, but aortic VTI, velocity and gradients were relatively preserved in the older *Pkn2*Het mice (Figure 6e,f; Supplementary Table S2). This may be a consequence of greater flexibility/elasticity of the aorta in these mice compared with the WT animals (Figure 6b). This raises the possibility that some of the long survival of a minor subset of *SM22aCre*^*+/-*^ *Pkn2*^*fl/*fl^ mice may have been supported by loss of PKN2 in the aorta.

**Figure 6.**
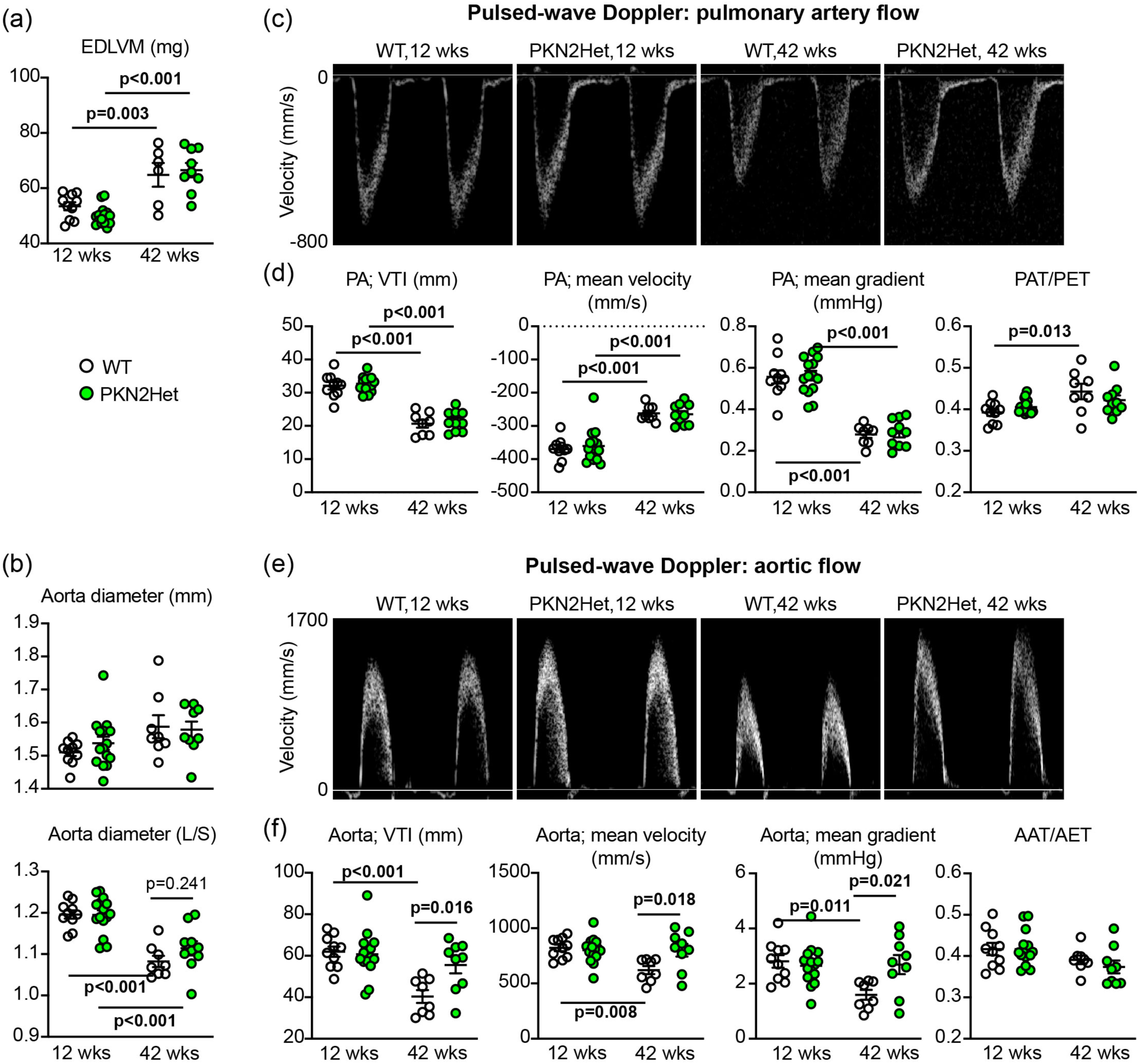
Effects of aging on aortic and pulmonary blood flow in WT and *Pkn2*Het mice. Echocardiograms were from *Pkn2*Het and WT littermates with an average age of 12 or 42 weeks. (**a**), End diastolic left ventricular mass (EDLVM) was estimated from long-axis B-mode images using speckle-tracking strain analysis. (**b**), The width of the aorta was measured from B-mode images from the widest diameter taken at cardiac systole (upper panel) and assessing the ratio of this to the narrowest diameter taken after the aortic valve closed (L/S). (**c-f**), Pulsed wave Döppler was used to assess blood flow as it leaves the heart into the pulmonary artery (**c**,**d**) and the aorta (**e**,**f**). Additional data are in Supplementary Table S2. Representative images are shown (**c**,**e**) and with the analysis (**d**,**f**). Individual data points are shown with means ± SEM. Analysis used 2-way ANOVA with Holm-Sidak’s post-test.

Overall, our studies of PKN2 in the adult heart indicate that it plays a significant role in cardiovascular adaptation to pathophysiological stresses, both in disease and as animals age, affecting both the contractile cardiomyocytes themselves and the major vessels.

## DISCUSSION

This study addresses the role of PKN2 throughout development, from its importance in cardiomyocytes in the embryo, through influence on cardiac remodelling in pathological cardiac hypertrophy in the adult heart and to a potential role in aging. It shows that there can be a continuous spectrum between heart disease defined as being ‘congenital’ and what might be considered as ‘acquired’ heart disease in adults. As previously reported,^11, 12^ PKN2 is critical for embryonic cardiac development but our data demonstrate that PKN2 plays a crucial role in cardiomyocytes, having a particular effect on development of the compact myocardium. PKN2 also supports cardiac remodelling in response to hypertension, but has apparently little impact on the aging heart, potentially having a greater effect on the vasculature.

Previous studies using the SM22α promoter in mice suggested that PKN2 is required in cardiomyocytes for embryonic cardiac development.^12^ Our data, using the XMLC promoter for specific cardiomyocyte deletion of *Pkn2* reinforce this conclusion and *XMLC*-driven knockout of *Pkn2* was of even greater severity, with only one homozygote survivor from the *XMLC*-driven knockout. In both models, embryos were produced at a normal Mendelian ratio, but the ventricular walls of the hearts in E14.5-E18.5 embryos were very thin, and the integrity and contiguousness of these walls was compromised, impacting on the overall architecture of the heart as it developed in utero. The heart is the earliest functioning organ to form in the embryo and, in mice, cardiac looping is generally completed by E9.0 with chamber development apparent by E9.5.^26-28^ Although looping may be delayed, cardiac development appears relatively normal through to E11.5 with *SM22α*-driven *Pkn2* knockout, with no evidence of any significant effect on cardiomyocyte proliferation or global cardiac structure.^12^ Here, we showed that by E14.5, there were significant effects of *Pkn2* deletion in cardiomyocytes (Figures 1 and 3): cardiac trabeculae had developed between E9.5 and E14.5, projecting into the ventricular chambers as expected, but there was a failure in formation of the compact myocardium, resulting in thin walls. Consistent with this, amongst other changes, there is a decrease in expression of the compact layer marker, *Hey2*^29^ in these mutant embryos (unpublished observations). This phase of cardiac development is still poorly understood, but cardiomyocytes required for compaction develop largely from a different pool of cells from the base of the trabeculae that are less differentiated and have higher proliferative potential.^26, 27^ The compaction process requires proliferation of these cells and, although the trabecular cells contribute to the compact myocardium, they also form part of the vasculature within the myocardium and the Purkinje fibre network. PKN2 can control migration and influence intercellular adhesion in other cells (e.g. fibroblasts, epithelial cells, skeletal muscle myoblasts^9, 12, 30, 31^), factors that potentially influence cardiac compaction in embryonic development.

Although mice with cardiomyocyte *Pkn2* knockout had normal Mendelian ratios throughout embryonic development, even with the milder *SM22α*-driven knockout model, only ∼30% survived the perinatal phase. This is probably because of myocardial developmental malformations in particular the lack of an adequate compact myocardium compromising neonatal cardiac adaptation to an increased workload and oxidative metabolism, along with cell cycle withdrawal of cardiomyocyte and a switch from hyperplasia to maturational (hypertrophic) growth^28^. The data have clear implications for congenital heart disease both with respect to PKN2 itself and its downstream signalling, particularly in relation to a potential role in formation of the compact myocardium and left ventricular non-compaction. Although left ventricular hypertrabeculation or non-compaction is a recognised genetic disease, it varies in presentation and severity. Cardiomyocyte *Pkn2* knockout appears to have the greatest similarity with the most severe form associated with fetal/neonatal disease that is often lethal by or within the first year of life and which (as for the *Pkn2* knockout mice) is often associated with structural cardiac abnormalities.^32, 33^

Mice with heterozygous PKN2 deletion have no obvious abnormalities and cardiac development appears normal.^12^ Moreover, we did not detect any profound differences in cardiac function/dimensions between WT and *Pkn2*Het mice into middle/old age (42 weeks). This means that PKN2 is largely dispensable once the heart has formed and is only required for cardiac remodelling in response to a severe stress such as that induced by sudden imposition of pressure-overload in the AngII experiments. We did detect a difference in aortic flow and the reduction in flow in the WT mice was largely prevented in the *Pkn2*Het mice. The meaning of this is not currently clear, but there may be some adaptation of the aorta that preserved LV function. The difference was not apparent in pulmonary flow suggesting that the effect was specific for the highly muscularised aortic wall.

Even though it may have little role in a non-stressed heart, PKN2 is important in the adult cardiac response to pathophysiological stressors since expression is increased in patients with DCM and in mouse hearts subjected to pressure-overload resulting from AngII treatment. In our hands, PKN2 haploinsufficiency compromised cardiac adaptation to AngII, with reduced LV hypertrophy and an overall reduction in the increase in the LV mass, resulting from inhibition of cardiomyocyte enlargement and fibrotic extracellular matrix.

This contrasts with a recent study reporting that tamoxifen-inducible, cardiomyocyte-specific knockout of both PKN1 or PKN2 (not either gene alone) in adult mice, reduced cardiac hypertrophy in models of pressure-overload, namely thoracic aortic constriction or AngII.^20^ The most obvious difference is our use of global heterozygotes rather than cardiomyocyte-specific knockout, and inhibition of fibrosis induced by AngII in our studies may result from PKN2 haploinsufficiency in cardiac non-myocytes, causing a reduction in fibrosis that reduces cardiomyocyte workload. Alternatively, the effect may be due to PKN2 haploinsufficiency in cardiomyocytes, and the differences reflect the higher degree of stress imposed on the heart in our studies (0.8 mg/kg/d AngII) compared with Sakaguchi et al., who used 0.1 mg/kg/d, a sub- or slow pressor dose.^34^ With global knockout, there are also potential systemic effects on the heart that influence cardiac adaptation to AngII. Although PKN2 does not appear to play a significant role in endothelial cells during development,^12^ endothelial cell specific knockout in adult mice increases blood pressure *in vivo*, potentially due to loss of phosphorylation of endothelial nitric oxide synthase (eNOS) with consequent reduction in NO production in the peripheral vasculature. There was no evidence for this in our studies of mice with global PKN2 haploinsuffiency, with no increase in fibrosis or cardiomyocyte hypertrophy at baseline. Furthermore, we detected no difference in phosphorylation of eNOS (data not shown). A final consideration relating to all of these studies is the background strain of the mice. Our studies used mice with a C57Bl/6J background, rather than C57Bl/6N and the cardiac responses of these two strains can differ substantially (see, for example, ^35^).

The PKNs family of enzymes remain poorly understood. PKN1 is the most well-investigated and is implicated in protection against ischaemia/reperfusion injury in *ex vivo* models but, even for this family member, there is little information on mechanism of action. For PKN2, there is less. It may influence gene expression directly via interaction and phosphorylation of HDAC5,^36^ preventing HDAC5 import into the nucleus and thus increasing chromatin remodelling or it may act via the transcription factor MRTF to regulate hypertrophy-associated gene expression.^20^ Our RNASeq data are not consistent with this because, although AngII induced changes in gene expression as expected with increases in classic hypertrophy-associated gene expression (e.g. *Nppa, Myh7, Nppb*), there was no apparent effect of *Pkn2* haploinsufficiency. Instead, *Pkn2* haploinsufficiency had a more general effect to moderate the changes induced by AngII on collagen production, mitochondrial TCA genes, interferon response genes and genes in the complement system (it is noted that mitochondrial TCA gene expression is also down-regulated in embryonic hearts from *SM22α-Cre+/-Pkn2*^*fl/fl*^ mice; data not shown). The net effect would be to maintain cardiac energetics, reduce inflammation and reduce fibrosis. The marked inhibition of the induction of Transgelin (*Tagln*, or *SM22α*) upon AngII treatment of *Pkn2*Het mice compared to WT littermates, suggests a potential mechanism. *SM22α* is an actin binding protein although its function is still obscure. In vascular smooth muscle cells, SM22α facilitates stress fibre formation and contractility^37^ so, with reduced SM22α, the response to AngII would be dampened in *Pkn2*Het mice, as observed. The mechanism for *SM22α* upregulation may be due to hypoxia and activation of HIF2α.^38^

Irrespective of the mechanism of action, this study adds to an increasing body of work indicating that the PKNs play an important role in cardiac remodelling in the adult; for PKN2, haploinsufficiency impacting the reponse to AngII and on early knockout (driven by XMLC2 or SM22α) formation of the heart during development. With further understanding of regulation, specific targets and functions, PKN signalling may offer therapeutic options for managing congenital and adult heart diseases. This undoubtedly requires further research but, since the PKNs are not redundant and have specific roles in different cells/tissues, it will be essential to develop specific tools for inhibition and manipulation of the different family members.

The Rho kinase (ROCK) inhibitor Fasudil is a known broad specificity PKN inhibitor,^39^ already in clinical use in Japan and China as a vasodilator.^40, 41^ More discriminating inhibitors are necessary and, although it may prove challenging to target individual PKNs by small molecule approaches,^39^ other approaches may be useful. For example, an siRNA against *PKN3* (Atu027) has been in clinical trials as a novel chemotherapeutic agent for solid cancers and pancreatic cancer (in combination with Gemcitabine),^42-44^ and exogenous application of an auto-inhibitory peptide from PKN1 has been explored also.^45^ Similar approaches might work for PKN2^46^ and, as with other protein kinases, developing these inhibitory systems will not only form the basis for novel therapeutics for the future, but their use as biochemical tools to elucidate mechanisms of action can facilitate the identification of other targets in the pathway.

## NON-STANDARD ABBREVIATIONS AND ACRONYMS

AngII: angiotensin II
DCM: dilated cardiomyopathy
DEG: differentially-expressed gene
HREM: high resolution episcopic microscopy
LV: left ventricle
PKN: Protein kinase N
PKN2het: mice heterozygous for PKN2 gene deletion
TCA: tricarboxylic acid
VSD: Ventricular septal defect
WT: wild-type

