## Supplementary Figures and Tables for "PKN2 deficiency leads both to prenatal ‘congenital’ cardiomyopathy and defective angiotensin II stress responses"

### SUPPLEMENTARY MATERIALS

**Expanded Materials and Methods**

**Supplementary Figure S1.** HREM XMLC2

**Supplementary Figure S2.** SM22a PKN2 mouse adult heart

**Supplementary Figure S3.** Embryonic defects SM22a Pkn2 mice

**Supplementary Figure S4.** Expression of PKN2 in rat ventricular myocytes

**Supplementary Figure S5.** Expression of hypertrophy associated genes in hearts from WT or Pkn2Het mice.

**Supplementary Table S1.** Surviving genotypes of conditional PKN2 knockouts.

**Supplementary Table S2.** Echocardiography: baseline data.

**Supplementary Table S3.** Echocardiography: mice treated for 7 d with angiotensin II or vehicle.

**Supplementary Table S4.** RNASeq analysis: mRNAs significantly upregulated by AngII in PKN2Het or WT hearts.

**Supplementary Table S5.** RNASeq analysis: mRNAs significantly downregulated by AngII in PKN2Het or WT hearts.

**Supplementary Table S6.** RNASeq analysis: mRNAs significantly upregulated by AngII in WT hearts.

**Supplementary Table S7.** RNASeq analysis: mRNAs significantly downregulated by AngII in WT hearts.

**Supplementary Table S8.** RNASeq analysis: mRNAs significantly upregulated by AngII in PKN2Het hearts.

**Supplementary Table S9.** RNASeq analysis: mRNAs significantly downregulated by AngII in PKN2Het hearts.

**Supplementary Table S10.** RNASeq data: gene clusters.

**Supplementary Table S11.** Body weights of adult mice.

**Supplementary Table S12.** qPCR primers.

**Supplementary References.**

**Supplementary legends for Video files.**

**Expanded Materials and Methods:**

**Ethics statement for mouse experiments.**

Animals were housed in the Biological Resource Unit (BRU) at Cancer Research UK's London Research Institute (LRI to 2015), University College London (UCL)'s Centre for Advanced Biomedical Imaging (for pilot imaging experiments) and the Biological Research Facility (BRF) at the Francis Crick Institute (from 2016 onward), or the BioResource Facility at St. George's University of London. Each site is UK registered with a Home Office certificate of designation. Studies were performed in accordance with European Parliament Directive 2010/63/EU on the protection of animals used for scientific purposes, institutional animal care committee procedures (CRUK's London Research Institute, University College London, The Francis Crick Institute, University of Reading and St. George's University of London) and the UK Animals (Scientific Procedures) Act 1986 (under Procedure Project Licences 77/8066, P166DEA98, 70/7474, 70/8248, 70/8249, 70/8709 and P8BAB0744).

**Mouse strains, *in vivo* mouse imaging and experiments, and *ex vivo* imaging.**

*Pkn2*Het, floxed *Pkn2* and *SM22α-Cre* mouse strains were maintained on a C57Black6J background, and each sourced as described in Quetier *et al.*.[1] The *XMLC2-Cre* mouse strain[2] was provided from within the Francis Crick Institute and was on a mixed background, subsequently in this study back-crossed onto the C57Black6J background. Most Cre mice in this study also carried the mTmG reporter allele at the Rosa26 locus, sourced as described previously.[1]

*In vivo* imaging of mice from *SM22α-Cre* and *XMLC2-Cre* crosses was carried out at UCL-CABI (up to 2017; SM22α-Cre only) or the Francis Crick Institute (post 2017). MRI, micro-CT and ultrasound imaging technologies were utilised. All *in vivo* mouse imaging was carried out under continuous inhalation anaesthesia using isoflurane (1.5-5%) supplied with oxygen at 1-2 L/min, and with appropriate restraint. We note 5% isoflurane was used to induce anaesthesia initially, and maintenance was typically at 1.5-2%, but some strongly phenotypic *SM22α-Cre*$^{+/-}$ *Pkn2*$^{fl/fl}$ mice required higher levels of isoflurane for successful maintenance. Procedures typically lasted only 20 minutes, with exception for cine-MRI, which necessitated longer non-recovery procedures. At CABI, the micro-CT was performed on mice in the supine position in a nanoScanPET/CT scanner (Mediso, Hungray) with a 50 kVP X-ray source, with 300 ms exposure time in 720 projections with an acquisition time of 8 minutes. Ultrasound of young mice was performed using a VEVO2100 (VisualSonics Inc., Toronta, ON, Canada) with MS400 18-38 MHz transducer mouse probe whilst middle-aged mice were imaged using a VEVO3100 with a MS-550D 25-55 MHz transducer. Mice were restrained on a heated VEVO Imaging Station, and cardiac scans of the parasternal long-axis and short-axis were recorded in B mode and M mode, and flow velocity waveforms of the aorta near the aortic valve were obtained with colour Doppler and then placing the pulsed wave Doppler sample gate over the colour Doppler signals. At the Francis Crick Institute, cardiac cine-MRI was performed using a 9.4T MRI (Bruker GmbH) equipped with a 4-channel receive only mouse cardiac coil and 86 mm volume transmit coil, and Paravision 6.0.1 software. Mice were set up lying prone head-first with a heat pad and breathing movement sensor pad. A series of fast low-angle shot (FLASH) scans used for localization of the heart and to determine the short-axis. Short-axis retrospectively-gated cine-MRI (intragate-FLASH sequence) was performed using the follow parameters: 0.8 mm slices covering the entire left and right ventricles; 128 x 128 pixels matrix and field of view of 25 x 25 mm, giving a resolution of ~195 μm; TR=5.5ms, TE=2.233ms, 10° flip angle; 300x oversampling with 24 cine frames reconstruction. The ultrasound used at the Francis Crick Institute was a VEVO3100, but otherwise scans were performed as described above.

MRI data was converted using in-house Matlab scripts to obtain tiffs, followed by ImageJ for image analysis, following published procedures,[3] and entailed determining the area of lumen in each the LV and RV in each slice at each systole and diastole, and calculating an approximate volume for the lumens of LV and RV at each systole and diastole based on the slice thickness of 0.8mm. Stroke volumes and the ejection fractions were determined by comparing diastolic and systolic chamber volumes. The relative volumes of the LV and RV, and number of slices containing LV and RV enabled assessment of abnormalities of shape of the hearts, as reported in Results.

*Ex vivo* imaging of 10% buffered formalin-fixed, and subsequently PBS-soaked (24h) whole carcasses or extracted plucks (heart and lungs) from a cohort of mice from the SM22α-Cre crosses was carried out at the Francis Crick Institute by micro-CT using a SkyScan1176 CT scanner (Bruker MicroCT, Kontich, Belgium). 394 projections were acquired over a 180º trajectory with an exposure time of 65ms, frame averaging of 3, X-ray source voltage and current of 50kV and 500µA, and a 0.5mm Al filter. Scans were reconstructed at a 34.2µm isotropic resolution using nRecon software (version 1.6.10.1, Bruker MicroCT). Video 2 was generated by segmentation of heart, lungs and cardiac calcification using Analyze (version 12.0, AnalyzeDirect, Overland Park, KS USA) using threshold based segmentation.

For *in vivo* basal and AngII-challenge studies with young mice, male *Pkn2*Het and wild-type (WT) littermates (average age 11 weeks) on a C57Bl/6J background were imported into the BioResource Facility at St. George's University of London and allowed to acclimatise for 7 d. Mice were randomly allocated to each treatment group; body weights are provided in Supplementary Table S11. Drug delivery used 1007D Alzet osmotic pumps, filled according to the manufacturer's instructions. Mice received minipumps for delivery of

0.8 mg/kg/d AngII (Merck) or vehicle (acidified PBS) Minipumps were incubated overnight in sterile PBS (37°C), then implanted subcutaneously under continuous inhalation anaesthesia using isoflurane (induction at 5%, maintenance at 2-2.5%) mixed with 2 L/min $O_2$. A 1 cm incision was made in the mid-scapular region and mice were given 0.05 mg/kg (s.c.) buprenorphine (Ceva Animal Health Ltd.) to repress post-surgical discomfort. Minipumps were implanted portal first in a pocket created in the left flank region of the mouse. Wound closure used a simple interrupted suture with polypropylene 4-0 thread (Prolene, Ethicon). Mice were allowed to recover singly and returned to their home cage once fully recovered. Echocardiography was performed on anaesthetised mice using the VEVO2100 imaging system equipped with a MS400 18-38 MHz transducer (Visualsonics). Mice were anaesthetised in an induction chamber with isoflurane (5% flow rate) with 1 l/min $O_2$ then transferred to the heated Vevo Imaging Station. Anaesthesia was maintained with 1.5% isoflurane delivered via a nose cone. Baseline scans were taken prior to experimentation (-7 to -3 days). Further scans were taken at intervals following tamoxifen treatment or minipump implantation. Imaging was completed within 20 min. Mice were recovered singly and transferred to the home cage once fully recovered.

For *in vivo* studies with middle-aged mice, male *Pkn2*Het and wild-type (WT) littermates (average age 42 weeks) were housed in the Francis Crick Institute and echocardiograms were taken with the VEVO3100 imaging system (Visualsonics).

Data analysis was performed using VevoLAB software (Visualsonics) by an independent assessor blinded to any AngII intervention. Left ventricular cardiac dimensions were assessed from short axis M-mode images with the axis placed at the mid-level of the left ventricle at the level of the papillary muscles. Data were gathered from two M-mode scans at each time point, taking mean values across 4 cardiac cycles for each echocardiogram. The diameter of the aorta was measured with the calliper function from B-mode images at the end of cardiac systole (with the aorta at its widest) and following aortic contraction, taking an average of measurements across two cardiac cycles. Cardiac function and left ventricular mass were measured B-mode long axis images using Vevo Strain software for speckle tracking. Blood flow was assessed using pulsed-wave Doppler.

Mice were euthanized either by cervical dislocation followed by exsanguination, or by $CO_2$ inhalation followed by cervical dislocation. Hearts were excised quickly, washed in PBS and blotted to remove excess PBS. The apex of the heart was snap-frozen in liquid $N_2$ and the remainder fixed in 10% buffered formalin for histology.

**HREM, histology and assessment of myocyte size and fibrosis.**

Samples for high resolution episcopic microscopy (HREM) were fixed in Bouin's for a minimum of 12h followed by extensive washing in PBS, dehydration in a graded methanol series, incubation in JB-4 (Sigma) /Eosine (Sigma)/Acridine orange (Sigma) mix overnight to ensure proper sample infiltration and then embedded in fresh mix by adding the accelerator (see [4] ). Once polymerised the blocks were imaged as previously described[5, 6.]; details of the process can be found at: https://dmdd.org.uk/hrem/. Samples were sectioned on a Leica sledge microtome at 1 or 2 µm or on a commercial HREM (Indigo Scientific) at 0.85 or 1.7 µm. An image of the surface of the block was then acquiered under GFP excitation wavelenght light using Olympus MVX10 microscope and a high resolution camera (Jenoptik). After acquisition the stacks were adjusted for gray levels using Photoshop CS6 and then processed for isotropic scaling, orthogonal resectioning, 25% downscaling, using a mixture of commercial and homemade software (see Wilson R et al. NAR 2016, Vol. 44 D855-D861). 3D volume rendering of the datasets were typically produced from the 25% downscaled stack using OsirixMD or Horos.

Histological staining and analysis were performed as previously described,[7] by board certified veterinary pathologists, assessing general morphology by haematoxylin and eosin (H&E), general fibrosis by Masson's or Gomori's trichrome (as indicated in figure legends and below) and collagen deposition using picrosirius red (PSR). Images of heart sections were captured and stored digitally using a Hamamatsu slide scanner. For analysis of myocyte cross-sectional area, cells stained by H&E within the LV (excluding epicardial and endocardial regions) were outline traced using NDP.view2 software (Hamamatsu). Only cells with a single nucleus that were clearly in cross-section were included in the analysis, and all cells in a given area meeting these criteria were measured. For assessment of interstitial fibrosis, PSR-stained sections were used and the areas of the myocardium in the middle of the left ventricular free wall, plus the points of intersection between the left ventricle and the interventricular septum were scored (0, no fibrosis; 1, limited fibrosis; 2, significant fibrosis; 3, extensive fibrosis permeating the tissue) and the mean value taken for each mouse. To assess perivascular fibrosis, Masson's Trichrome images were used. All arteries/arterioles with a clearly defined elastic lamina in cross section were measured across the diameter. Vessels > 20μm diameter were scored as for interstitial fibrosis. The mean values for each mouse were taken. Data analysis was performed by an independent assessor blinded to treatment groups.

**RNASeq, qPCR and immunoblotting of adult mouse heart samples.**

The apex of each of the mouse hearts was ground to powder under liquid $N_2$. Samples (10-15 mg) were homogenised with 1 ml RNA Bee (AMS Biotechnology Ltd). RNA was prepared according to the manufacturer's instructions and dissolved in nuclease-free water. The concentration and purity were assessed from the $A_{260}$ and $A_{260}/A_{280}$ values measured using an Implen NanoPhotometer.

The nf-core/rnaseq pipeline version 3.1[8] was used to prepare quantified expression matrices. The pipeline takes FastQ files as input and runs quality control checks on the data, trims reads for low quality nucleotides, aligns reads and quantifies aligned data to gene models. The GRCm38 reference and Ensembl release-95 gene models were provided. The '--aligner star_rsem' option was specified to align the reads with STAR[9] and quantify expression with RSEM[10] via RSEM version 1.3.1. Raw gene-level counts were imported into R using tximport [11] and DESeq2 version 1.32.0[12] used to test for differential expression with a FDR threshold of 1%.

Quantitative PCR (qPCR) was performed as previously described.[13] Total RNA (0.5 µg) was reverse transcribed to cDNA using High Capacity cDNA Reverse Transcription Kits with random primers (Applied Biosystems) according to the manufacturer's instructions. qPCR was performed using an ABI Real-Time PCR 7500 system (Applied Biosystems) using optical 96-well reaction plates and iTaq Universal SYBR Green Supermix (Bio-Rad Laboratories Inc.). *GAPDH* was used as the reference gene for the study. Results were normalized to *GAPDH*, and relative quantification was obtained using the ΔCt (threshold cycle) method; relative expression was calculated as $2^{-\Delta\Delta Ct}$, and normalised to vehicle or time 0. Primers were from Eurofins Genomics; sequences are provided in Supplementary Table S12.

Samples of heart powders (10-15 mg) were extracted in 8 vol (relative to powder weight) Buffer A [20 mmol/L Tris pH 7.5, 1 mmol/L EDTA, 10% (v/v) glycerol, 1% (v/v) Triton X-100, 100 mmol/L KCl, 5 mmol/L NaF, 0.2 mmol/L $Na_3VO_4$, 5 mmol/L $MgCl_2$, 0.05% (v/v) 2-mercaptoethanol, 10 mM benzamidine, 0.2 mM leupeptin, 0.01 mM trans-epoxy succinyl-l-leucylamido-(4-guanidino)butane, 0.3 mM phenylmethylsulphonyl fluoride, 4 µM microcystin]. Samples were vortexed and extracted on ice (10 min). Extracts were centrifuged (10,000 × g, 10 min, 4°C). The supernatants were removed, a sample was taken for protein assay and the remainder boiled with 0.33 vol sample buffer [0.33 mol/L Tris-HCl pH 6.8, 10% (w/v) SDS, 13% (v/v) glycerol, 133 mol/L dithiothreitol, 0.2 mg/mL bromophenol blue]. Protein concentrations were determined by BioRad Bradford assay using bovine serum albumin (BSA) standards.

Proteins were separated by SDS-PAGE using using a BioRad mini-gel system with 8% (w/v) polyacrylamide resolving gels and 6% stacking gels (200 V, 90 min) for PKN1/2 (40 μg total protein), or 12% polyacrylamide resolving gels and 6% stacking gels (200 V, 50 min) for GAPDH (5 μg total protein). Proteins were transferred electrophoretically to nitrocellulose using a BioRad semi-dry transfer cell (10 V, 60 min) and detected as previously described.[13] Bands were visualised by enhanced chemiluminescence using ECL Prime Western Blotting detection reagents and using an ImageQuant LAS4000 system (GE Healthcare). ImageQuant TL 8.1 software (GE Healthcare) was used for densitometric analysis. Raw values for phosphorylated PKN1/2 were normalised to the total kinase. Values for all samples were normalised to the mean of the controls. Primary antibodies for total PKN2 (Cat. No. 2612), phospho-PKN1/2 (Cat. No. 2611) and GAPDH (Cat. No. 2118) were from Cell Signalling Technology. Antibodies for PKN1 were from BD Transduction Laboratories (Cat. No. 610686). Phospho- and total PKN antibodies were used at 1/750 dilution; GAPDH antibodies were used at 1/1000 dilution.

**Bioinformatics analysis for transcript expression in dilated cardiomyopathy.**

mRNA expression of *PKN1* (ENSG00000123143), *PKN2* (ENSG00000065243) and *PKN3* (ENSG00000160447) in control and diseased human hearts was determined using a published RNASeq dataset for left ventricular samples of 97 patients with end-stage dilated cardiomyopathy taken at the time of transplantation or left ventricular assist device implantation, and 108 non-diseased controls.[14] Differential expression analysis was carried out using DESeq2 (V*1.18.1*, Wald test).[12]

### Supplementary Figures

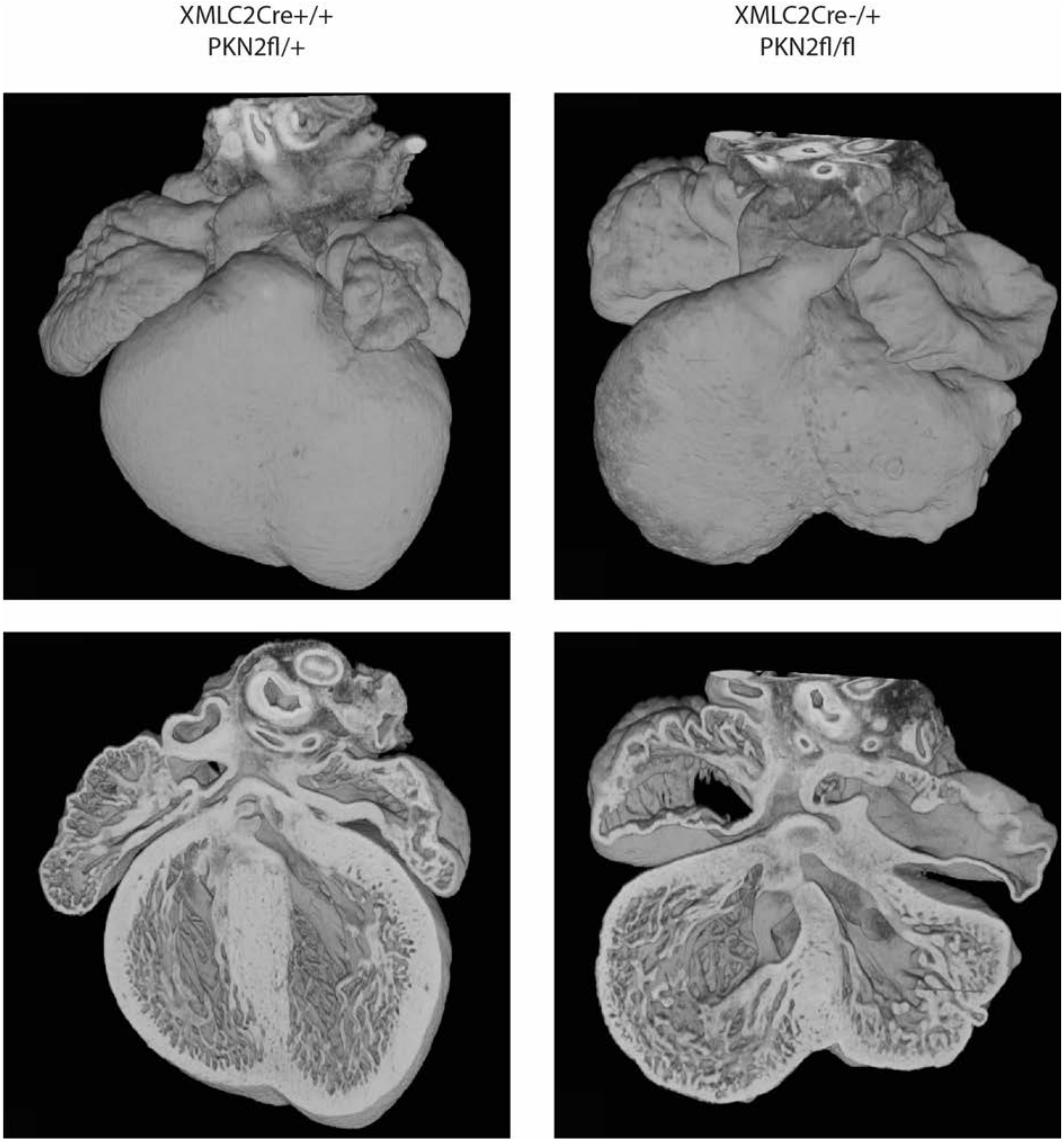

**Supplementary Figure S1. HREM XMLC2**

HREM images are from a reconstruction of a 14.5d XMLC2Cre$^{+/-}$ *Pkn2*$^{fl/+}$ embryo (left) and of a XMLC2Cre$^{+/-}$*Pkn2*$^{fl/fl}$ (right) from the same dam. The upper panels are surface images and the lower panels illustrate sections.

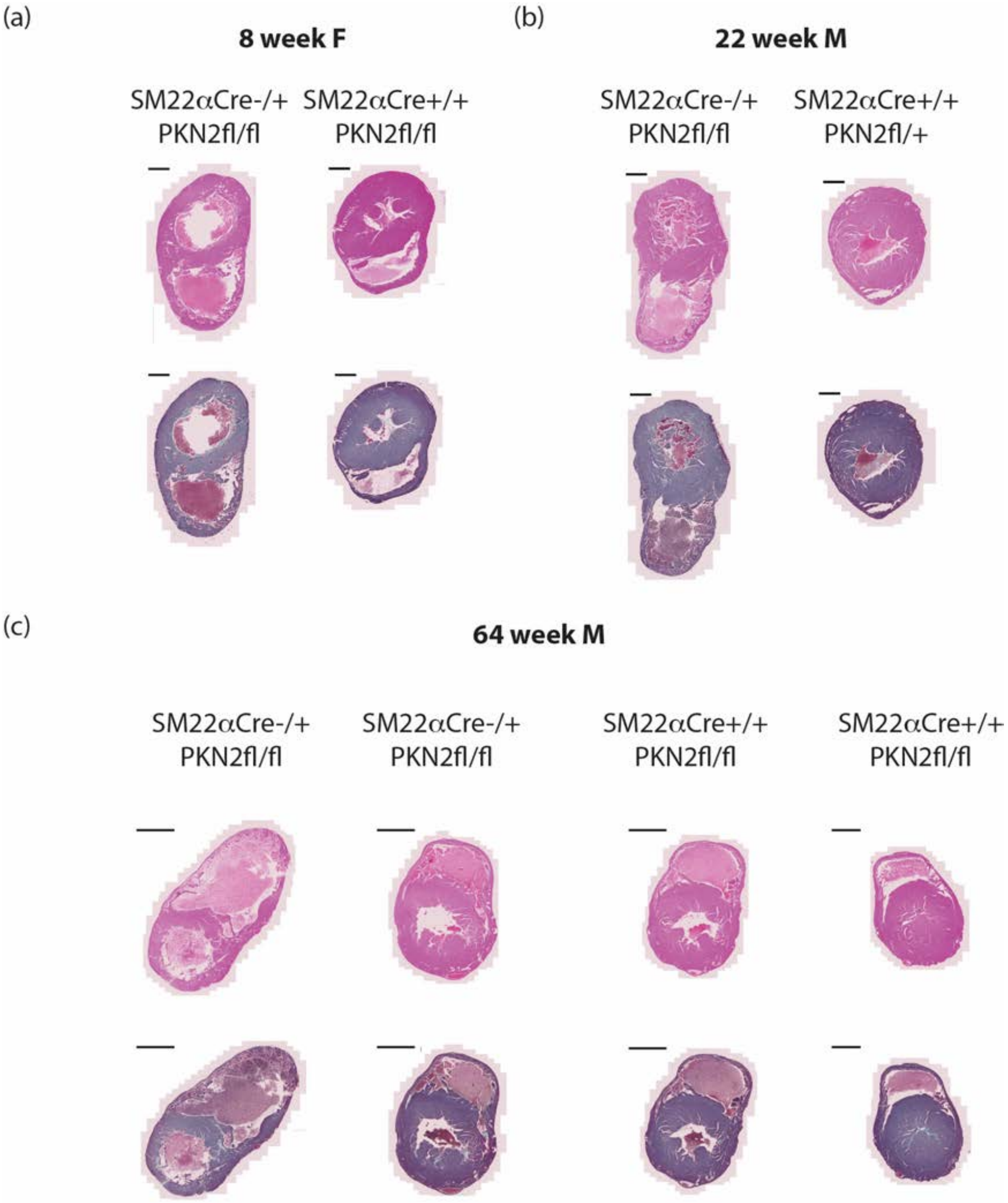

**Supplementary Figure S2. SM22α PKN2 mouse adult heart**

H&E (top rows) and Gomori's Trichrome (lower rows) stained sections through the short-axis of the heart, at 2-3 mm from the apex from littermates of (a) females culled at 8 weeks of age, (b) males at 22 weeks, and (c) males at 64 weeks, with genotypes as labelled. For each group, cull was triggered due to loss of condition of one littermate (left in each group). Scale bars are (a,b) 1 mm or (c) 2 mm.

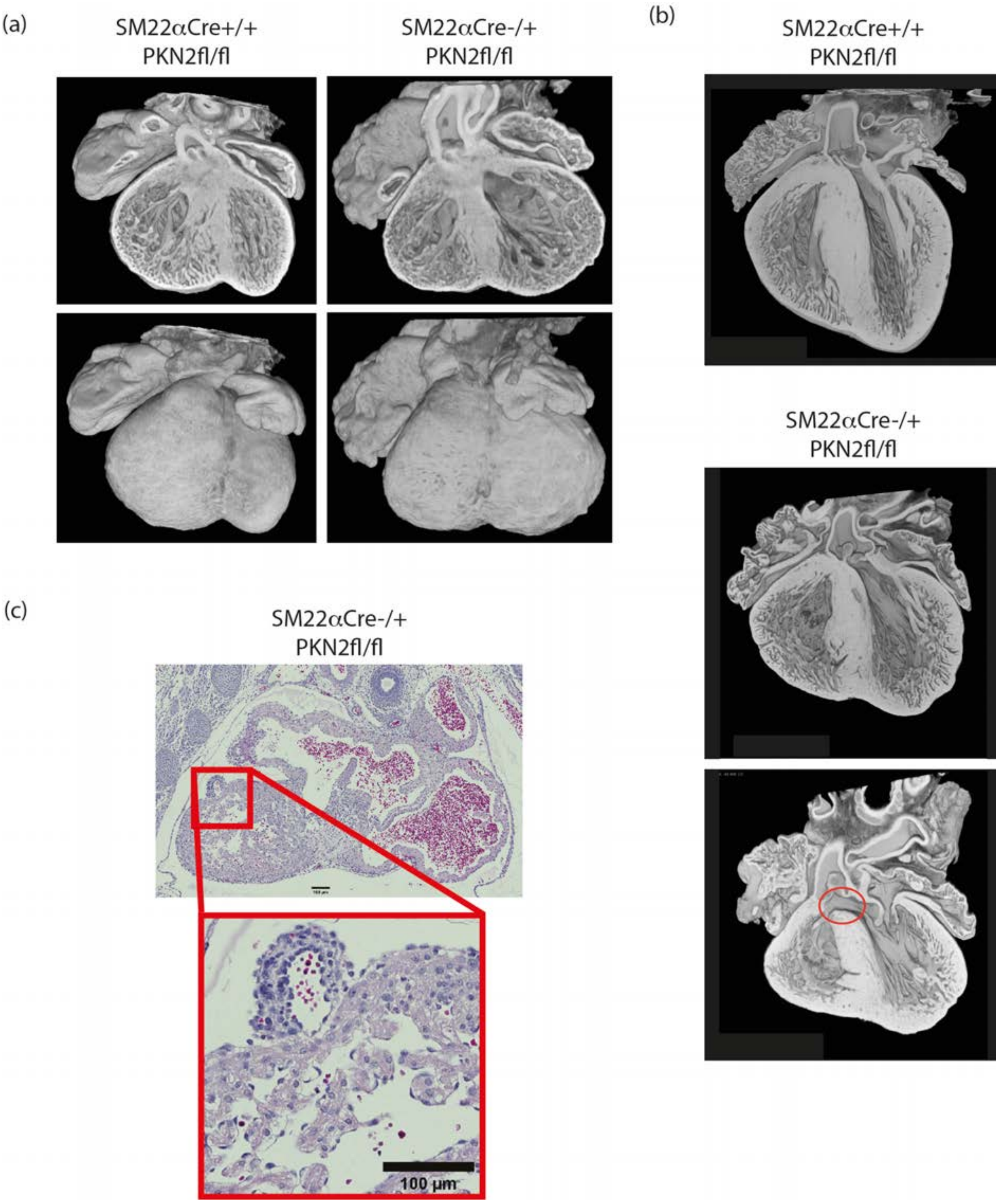

**Supplementary Figure S3 Embryonic defects SM22α Pkn2 mice**

HREM images are from reconstructions of embyros collected at (a) E14.5 and (b) E18.5 days gestation. Genotypes are as indicated. (c) H&E stained section of a SM22αCre$^{+/-}$ *Pkn2*$^{fl/fl}$ E14.5 heart with 100 µm scale, as labelled.

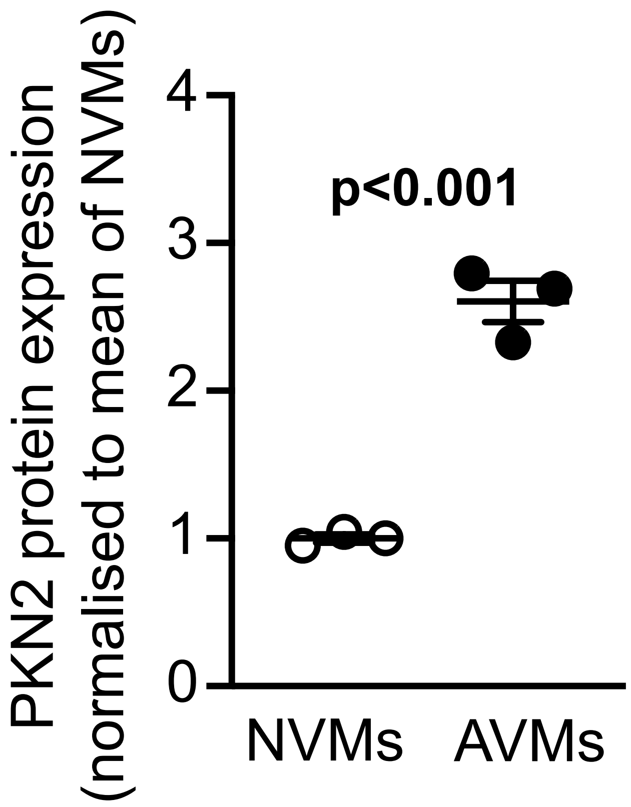

**Supplementary Figure S4. Expression of PKN2 in neonatal rat ventricular myocytes (NVMs) compared with adult rat ventricular myocytes (AVMs) relative to cell size.** Expression data were from Fuller SJ et al. (Cardiovasc Res. 2015; 108: 87-98) adjusted for cell size according to membrane capacitance which increases from 13 pF in 1- to 2-day NVMs to 156 pF in AVMs (Cerbai et al. Cardiovasc Res1999;42:416–423; Banyasz et al. Exp Physiol2008;93:370–382).

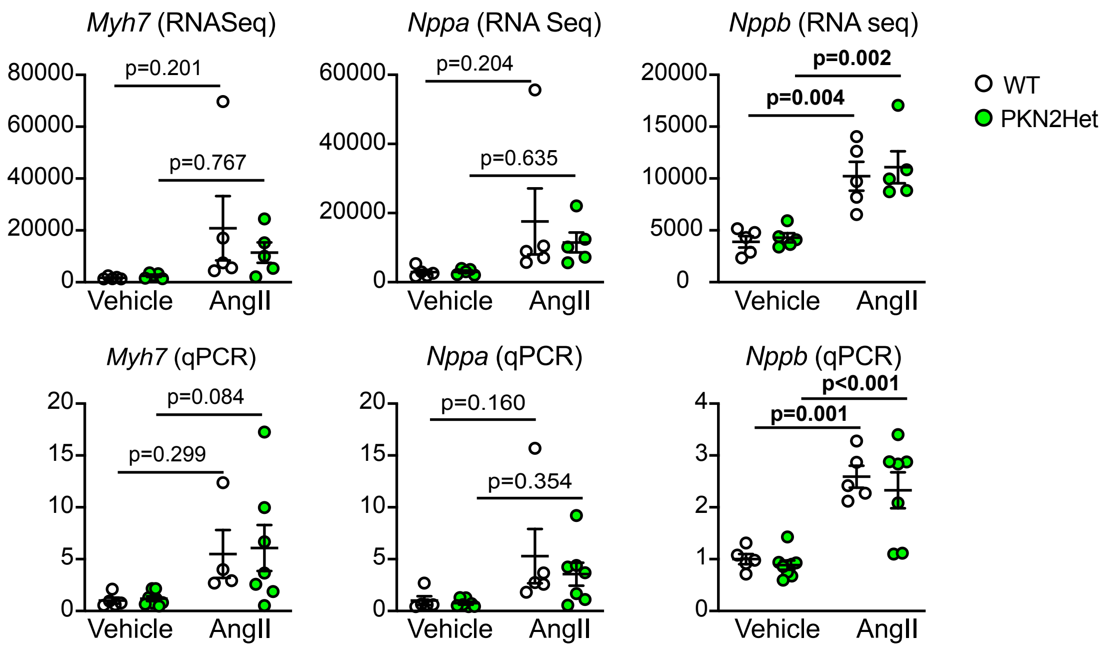

**Supplementary Figure S5. Expression of hypertrophy associated genes in hearts from WT or Pkn2Het mice.** WT (white) or Pkn2Het (green) mice were treated with vehicle (left side of each graph) or AngII (0.8 mg/kg/d; right side of each graph) for 7 d. RNA was prepared and used for RNASeq (upper panels) or qPCR (lower panels). Individual data points are shown with means ± SEM. Analysis used 2-way ANOVA with Holm-Sidak's post-test.

**Supplementary Table S1. Surviving genotypes of conditional PKN2 knockouts.** *SM22αCre* and *XMLC2Cre* mouse strain crosses are indicated in column 1, with numbers of experimentally determined genotypes shown in rows for the age ranges defined. The representation of the PKN2 knockout (ie $PKN2^{fl/fl}$ in the context of Cre expression) was analysed as a function of the all genotypes using Fisher's test.

| Parent Genotypes | Age | Cre negative | | Cre positive | | Fisher's test | Representation |
|---|---|---|---|---|---|---|---|
| | | $PKN2^{fl/+}$ | $PKN2^{fl/fl}$ | $PKN2^{fl/+}$ | $PKN2^{fl/fl}$ | | |
| SM22α-$Cre^{+/-}$$PKN2^{fl/+}$ X $PKN2^{fl/fl}$ | E14.5-E18.5 | 54 | 43 | 33 | 46 | >0.5 | Mendelian |
| | 3 weeks | 166 | 215 | 176 | 44 | 0.0001 | under-represented |
| XMLC2-$Cre^{+/-}$$PKN2^{fl/+}$ X $PKN2^{fl/fl}$ | E14.5-E18.5 | 5 | 3 | 10 | 6 | >0.5 | Mendelian |
| | 3 weeks | 37 | 41 | 47 | 1 | 0.0001 | under-represented |

**Supplementary Table S2. Echocardiography data for 12 and 42 week male mice with heterozygous PKN2 gene deletion (PKN2Het) and wild-type (WT) mice: baseline data.** AAT, aortic acceleration time; AET, Aortic expulsion time; VTI, velocity time interval; PAT, pulmonary acceleration time; PET, pulmonary expulsion time; PA, pulmonary artery; LV, left ventricle; ID, internal diameter; AW, anterior wall; PW, posterior wall.

| | **WT: 12 wk (n=10)** | | **PKN2Het: 12 wk (n=15)** | | **WT: 42 wk (n=8)** | | **PKN2Het: 42 wk (n=10)** | |
|---|---|---|---|---|---|---|---|---|
| | Mean | SEM | Mean | SEM | Mean | SEM | Mean | SEM |
| **Aortic flow (Pulsed Wave Doppler)** | | | | | | | | |
| AAT (ms) | 23.80 | 0.82 | 23.59 | 0.72 | 18.98 | 0.72 | 20.03 | 1.08 |
| AET (ms) | 57.21 | 0.68 | 56.55 | 0.68 | 48.54 | 1.13 | 53.56 | 1.77 |
| AAT/AET | 0.42 | 0.01 | 0.42 | 0.01 | 0.39 | 0.01 | 0.37 | 0.01 |
| Aorta VTI (mm) | 61.95 | 2.47 | 60.71 | 2.86 | 40.31 | 3.09 | 55.47 | 3.77 |
| Aorta Mean Velocity (mm/s) | 821.32 | 29.33 | 800.96 | 29.24 | 620.71 | 36.58 | 800.88 | 54.81 |
| Aorta Mean Gradient (mmHg) | 2.81 | 0.23 | 2.65 | 0.19 | 1.59 | 0.18 | 2.69 | 0.33 |
| Aorta Peak Velocity (mm/s | 1632.92 | 58.53 | 1592.77 | 53.01 | 1271.29 | 82.00 | 1620.63 | 104.77 |
| Aorta Peak Grad (mmHg) | 11.08 | 0.93 | 10.46 | 0.65 | 6.69 | 0.83 | 10.94 | 1.32 |
| Aorta Peak Pressure (mmHg) | 10.85 | 0.86 | 10.31 | 0.65 | 6.64 | 0.82 | 10.81 | 1.31 |
| **Pulmonary flow (Pulsed Wave Doppler)** | | | | | | | | |
| PAT (ms) | 23.71 | 0.53 | 24.90 | 0.37 | 24.97 | 0.98 | 24.71 | 0.72 |
| PET (ms) | 60.47 | 0.92 | 61.43 | 0.65 | 56.74 | 2.15 | 58.94 | 2.42 |
| PAT/PET | 0.39 | 0.01 | 0.41 | 0.00 | 0.44 | 0.02 | 0.42 | 0.01 |
| PA VTI | 32.11 | 1.12 | 32.70 | 0.58 | 20.66 | 1.20 | 21.44 | 0.97 |
| PA Mean Velocity (mm/s) | -369.58 | 10.38 | -360.73 | 12.93 | -262.10 | 8.32 | -264.84 | 9.79 |
| PA Mean Gradient (mmHg) | 0.56 | 0.03 | 0.56 | 0.02 | 0.28 | 0.02 | 0.29 | 0.02 |
| PA Peak Velocity (mm/s | -764.05 | 17.63 | -740.28 | 23.98 | -551.01 | 13.94 | -558.99 | 25.85 |
| PA Peak Grad (mmHg) | 2.37 | 0.11 | 2.36 | 0.08 | 1.22 | 0.06 | 1.28 | 0.12 |
| **Aorta diameter (B-Mode)** | | | | | | | | |
| Widest (mm) | 1.51 | 0.01 | 1.54 | 0.02 | 1.59 | 0.03 | 1.60 | 0.03 |
| Narrowest (mm | 1.28 | 0.02 | 1.29 | 0.02 | 1.47 | 0.04 | 1.44 | 0.04 |
| Wide/narrow | 1.20 | 0.01 | 1.19 | 0.01 | 1.08 | 0.01 | 1.11 | 0.02 |
| **Left ventricle dimensions (short axis M-mode)** | | | | | | | | |
| Heart Rate (bpm | 522.13 | 11.06 | 508.26 | 11.26 | 473.62 | 17.63 | 448.95 | 18.55 |
| LVID;s (mm) | 2.97 | 0.05 | 2.92 | 0.05 | 2.59 | 0.07 | 2.71 | 0.05 |
| LVID;d (mm) | 4.18 | 0.05 | 4.14 | 0.05 | 3.80 | 0.06 | 3.95 | 0.04 |
| LVAW;s (mm) | 1.11 | 0.02 | 1.05 | 0.01 | 1.10 | 0.04 | 1.07 | 0.03 |
| LVAW;d (mm) | 0.84 | 0.02 | 0.77 | 0.01 | 0.84 | 0.02 | 0.81 | 0.02 |
| LVPW;s (mm) | 1.03 | 0.03 | 1.01 | 0.02 | 1.16 | 0.03 | 1.11 | 0.05 |
| LVPW;d (mm) | 0.72 | 0.02 | 0.69 | 0.01 | 0.78 | 0.04 | 0.77 | 0.02 |
| **Cardiac function (speckle-tracking strain analysis; long axis B-mode)** | | | | | | | | |
| Heart rate (bpm | 502.23 | 10.29 | 487.39 | 10.63 | 459.97 | 22.21 | 460.58 | 22.49 |
| Stroke volume (μl) | 25.51 | 1.35 | 26.23 | 0.76 | 24.18 | 2.71 | 24.17 | 2.51 |
| Fractional shortening (%) | 26.85 | 0.76 | 28.45 | 0.85 | 28.04 | 1.43 | 29.33 | 2.73 |
| Ejection fraction (%) | 50.11 | 1.93 | 53.31 | 1.20 | 53.38 | 2.76 | 54.07 | 2.95 |
| Cardiac output (ml/min) | 12.76 | 0.59 | 12.79 | 0.49 | 10.94 | 1.10 | 11.01 | 1.04 |
| End diastolic LV mass | 53.48 | 1.45 | 49.95 | 0.88 | 64.83 | 4.30 | 66.52 | 2.59 |
| End systolic LV mass | 56.46 | 1.57 | 52.34 | 0.75 | 66.08 | 4.28 | 67.84 | 2.70 |

**Supplementary Table S3. Echocardiography data for 12 week male mice with heterozygous PKN2 gene deletion (PKN2Het) and wild-type (WT) mice treated for 7 d with 0.8 mg/kg/d angiotensin 2 or vehicle.** AAT, aortic acceleration time; AET, Aortic expulsion time; VTI, velocity time interval; PAT, pulmonary acceleration time; PET, pulmonary expulsion time; PA, pulmonary artery; LV, left ventricle; ID, internal diameter; AW, anterior wall; PW, posterior wall.

| | **WT: vehicle (n=5)** | | **PKN2Het: vehicle (n=8)** | | **WT: AngII (n=5)** | | **PKN2Het: AngII (n=7)** | |
|---|---|---|---|---|---|---|---|---|
| | Mean | SEM | Mean | SEM | Mean | SEM | Mean | SEM |
| **Aortic flow (Pulsed Wave Doppler)** | | | | | | | | |
| AAT (ms) | 23.50 | 1.19 | 22.76 | 0.86 | 24.42 | 1.23 | 22.53 | 1.44 |
| AET (ms) | 57.71 | 1.57 | 55.21 | 1.39 | 56.29 | 1.65 | 54.49 | 1.23 |
| AAT/AET | 0.41 | 0.03 | 0.41 | 0.01 | 0.43 | 0.02 | 0.41 | 0.02 |
| Aorta VTI (mm) | 61.67 | 5.44 | 58.84 | 4.05 | 58.82 | 3.16 | 61.60 | 5.13 |
| Aorta Mean Velocity (mm/s) | 780.00 | 63.10 | 819.21 | 59.70 | 811.40 | 45.61 | 864.12 | 57.41 |
| Aorta Mean Gradient (mmHg) | 2.53 | 0.41 | 2.80 | 0.37 | 2.68 | 0.30 | 3.08 | 0.42 |
| Aorta Peak Velocity (mm/s | 1583.20 | 128.93 | 1607.53 | 109.14 | 1498.86 | 29.58 | 1689.25 | 81.15 |
| Aorta Peak Grad (mmHg) | 10.32 | 1.68 | 10.71 | 1.32 | 9.01 | 0.35 | 11.60 | 1.08 |
| Aorta Peak Pressure (mmHg) | 10.20 | 1.68 | 10.61 | 1.30 | 8.91 | 0.35 | 11.50 | 1.09 |
| **Pulmonary flow (Pulsed Wave Doppler)** | | | | | | | | |
| PAT (ms) | 26.13 | 1.02 | 25.44 | 1.09 | 24.54 | 0.58 | 22.71 | 1.25 |
| PET (ms) | 62.38 | 1.29 | 63.13 | 1.47 | 58.68 | 2.93 | 56.67 | 1.72 |
| PAT/PET | 0.42 | 0.01 | 0.40 | 0.01 | 0.42 | 0.02 | 0.40 | 0.02 |
| PA VTI | 31.66 | 1.74 | 31.73 | 1.05 | 28.44 | 2.24 | 25.39 | 1.39 |
| PA Mean Velocity (mm/s) | -362.61 | 16.48 | -363.12 | 13.26 | -333.86 | 18.50 | -310.38 | 10.82 |
| PA Mean Gradient (mmHg) | 0.53 | 0.05 | 0.53 | 0.04 | 0.45 | 0.05 | 0.39 | 0.03 |
| PA Peak Velocity (mm/s | -755.55 | 28.94 | -744.09 | 26.15 | -704.73 | 33.94 | -682.22 | 25.60 |
| PA Peak Grad (mmHg) | 2.30 | 0.18 | 2.24 | 0.16 | 2.01 | 0.20 | 1.88 | 0.14 |
| **Aorta diameter (B-Mode)** | | | | | | | | |
| Widest (mm) | 1.51 | 0.02 | 1.51 | 0.01 | 1.63 | 0.06 | 1.69 | 0.08 |
| Narrowest (mm | 1.21 | 0.02 | 1.27 | 0.03 | 1.44 | 0.09 | 1.56 | 0.09 |
| Wide/narrow | 1.24 | 0.02 | 1.19 | 0.03 | 1.14 | 0.04 | 1.09 | 0.01 |
| **Left ventricle dimensions (short axis M-mode)** | | | | | | | | |
| Heart Rate (bpm | 501.28 | 13.71 | 506.12 | 12.71 | 528.57 | 15.67 | 521.25 | 17.10 |
| LVID;s (mm) | 2.96 | 0.11 | 3.01 | 0.07 | 2.59 | 0.08 | 2.68 | 0.11 |
| LVID;d (mm) | 4.17 | 0.10 | 4.15 | 0.05 | 3.86 | 0.13 | 3.80 | 0.14 |
| LVAW;s (mm) | 1.08 | 0.04 | 1.07 | 0.01 | 1.29 | 0.06 | 1.15 | 0.02 |
| LVAW;d (mm) | 0.82 | 0.03 | 0.81 | 0.02 | 1.00 | 0.05 | 0.91 | 0.02 |
| LVPW;s (mm) | 1.03 | 0.06 | 1.02 | 0.03 | 1.31 | 0.07 | 1.18 | 0.02 |
| LVPW;d (mm) | 0.72 | 0.05 | 0.72 | 0.03 | 1.02 | 0.09 | 0.85 | 0.04 |
| **Cardiac function (speckle-tracking strain analysis; long axis B-mode)** | | | | | | | | |
| Heart rate (bpm | 462.84 | 9.65 | 503.48 | 16.63 | 525.26 | 14.72 | 512.86 | 19.44 |
| Stroke volume (µl) | 28.52 | 2.63 | 28.07 | 1.03 | 27.49 | 3.19 | 21.12 | 2.08 |
| Fractional shortening (%) | 24.54 | 1.58 | 27.42 | 1.62 | 33.15 | 2.06 | 25.56 | 1.69 |
| Ejection fraction (%) | 49.85 | 1.59 | 53.86 | 1.99 | 60.29 | 3.22 | 52.70 | 1.28 |
| Cardiac output (ml/min) | 13.21 | 1.31 | 14.22 | 0.90 | 14.41 | 1.70 | 10.68 | 0.90 |
| End diastolic LV mass | 54.15 | 4.23 | 51.75 | 2.14 | 71.11 | 5.80 | 54.44 | 4.21 |
| End systolic LV mass | 57.33 | 5.02 | 56.87 | 2.64 | 76.33 | 5.51 | 57.72 | 4.56 |

**Supplementary Table S4.** RNASeq analysis of effects of angiotensin II (AngII) on mRNA expression in hearts from PKN2Het vs WT littermates: mRNAs significantly upregulated by AngII in PKN2Het or WT hearts.

| Gene Symbol | Ensembl gene id | WT Vehicle | | WT AngII | | PKN2Het Vehicle | | PKN2Het AngII | |
|---|---|---|---|---|---|---|---|---|---|
| | | Mean | SD | Mean | SD | Mean | SD | Mean | SD |
| Sept5 | ENSMUSG00000072214 | 83 | 20 | 175 | 64 | 94 | 15 | 166 | 50 |
| Sept9 | ENSMUSG00000059248 | 865 | 99 | 1148 | 72 | 882 | 89 | 1148 | 182 |
| Sept11 | ENSMUSG00000058013 | 781 | 129 | 1170 | 161 | 705 | 99 | 998 | 196 |
| 1500009L16Rik | ENSMUSG00000087651 | 50 | 8 | 89 | 32 | 43 | 11 | 85 | 10 |
| 1700120C14Rik | ENSMUSG00000100599 | 84 | 4 | 153 | 41 | 76 | 23 | 122 | 23 |
| 2010111I01Rik | ENSMUSG00000021458 | 946 | 73 | 1185 | 106 | 902 | 74 | 1185 | 115 |
| Abcg1 | ENSMUSG00000024030 | 121 | 26 | 192 | 24 | 107 | 25 | 163 | 40 |
| Acan | ENSMUSG00000030607 | 4 | 3 | 66 | 49 | 3 | 1 | 32 | 34 |
| Ace | ENSMUSG00000020681 | 1467 | 152 | 2473 | 345 | 1594 | 190 | 2408 | 305 |
| Acta1 | ENSMUSG00000031972 | 6773 | 1395 | 24772 | 10833 | 8570 | 3863 | 28601 | 14894 |
| Actn1 | ENSMUSG00000015143 | 522 | 79 | 925 | 273 | 524 | 61 | 761 | 132 |
| Actr3 | ENSMUSG00000026341 | 2536 | 120 | 3001 | 264 | 2399 | 43 | 2908 | 199 |
| Adam12 | ENSMUSG00000054555 | 55 | 9 | 167 | 53 | 41 | 13 | 115 | 49 |
| Adam15 | ENSMUSG00000028041 | 1010 | 154 | 1434 | 80 | 997 | 96 | 1364 | 151 |
| Adamts12 | ENSMUSG00000047497 | 91 | 12 | 261 | 100 | 100 | 8 | 204 | 80 |
| Adamts2 | ENSMUSG00000036545 | 499 | 49 | 1369 | 604 | 526 | 39 | 1080 | 305 |
| Adamts8 | ENSMUSG00000031994 | 16 | 4 | 76 | 21 | 15 | 2 | 57 | 28 |
| Adamtsl2 | ENSMUSG00000036040 | 230 | 49 | 554 | 141 | 251 | 43 | 511 | 150 |
| Adcy7 | ENSMUSG00000031659 | 355 | 49 | 714 | 204 | 342 | 76 | 589 | 111 |
| Adgre1 | ENSMUSG00000004730 | 375 | 73 | 763 | 304 | 349 | 35 | 608 | 121 |
| AI506816 | ENSMUSG00000105987 | 463 | 36 | 655 | 41 | 454 | 52 | 676 | 86 |
| Aif1 | ENSMUSG00000024397 | 63 | 12 | 133 | 73 | 51 | 11 | 95 | 13 |
| Aldh1a2 | ENSMUSG00000013584 | 143 | 34 | 307 | 88 | 145 | 27 | 322 | 123 |
| Ankrd1 | ENSMUSG00000024803 | 20150 | 6483 | 57246 | 17002 | 19310 | 6244 | 49450 | 10874 |
| Ankrd23 | ENSMUSG00000067653 | 8223 | 1670 | 13521 | 3447 | 9104 | 579 | 14731 | 4085 |
| Anln | ENSMUSG00000036777 | 56 | 22 | 225 | 55 | 56 | 7 | 174 | 50 |
| Anxa1 | ENSMUSG00000024659 | 596 | 70 | 1111 | 254 | 555 | 84 | 919 | 152 |
| Anxa2 | ENSMUSG00000032231 | 1328 | 74 | 2290 | 204 | 1236 | 149 | 1957 | 280 |
| Anxa5 | ENSMUSG00000027712 | 1991 | 157 | 2628 | 137 | 1995 | 184 | 2495 | 432 |
| Apbb1ip | ENSMUSG00000026786 | 129 | 14 | 240 | 62 | 146 | 18 | 205 | 16 |
| Apod | ENSMUSG00000022548 | 115 | 26 | 229 | 22 | 122 | 16 | 198 | 30 |
| Apoe | ENSMUSG00000002985 | 3788 | 666 | 6913 | 2610 | 4103 | 455 | 5864 | 858 |
| Arhgap1 | ENSMUSG00000027247 | 496 | 38 | 622 | 41 | 520 | 32 | 610 | 36 |
| Arhgap11a | ENSMUSG00000041219 | 76 | 15 | 183 | 32 | 69 | 9 | 158 | 44 |
| Arhgap30 | ENSMUSG00000048865 | 154 | 24 | 265 | 55 | 147 | 24 | 206 | 37 |
| Arhgdib | ENSMUSG00000030220 | 506 | 27 | 664 | 56 | 483 | 58 | 627 | 60 |
| Arhgef40 | ENSMUSG00000004562 | 530 | 50 | 747 | 111 | 504 | 59 | 674 | 68 |
| Arid5a | ENSMUSG00000037447 | 177 | 60 | 315 | 53 | 146 | 28 | 315 | 68 |
| Arl4c | ENSMUSG00000049866 | 257 | 44 | 447 | 45 | 299 | 36 | 401 | 52 |
| Arl6ip1 | ENSMUSG00000030654 | 555 | 29 | 783 | 95 | 531 | 41 | 675 | 58 |
| Arpc1b | ENSMUSG00000029622 | 814 | 81 | 1186 | 179 | 737 | 113 | 1071 | 150 |
| Arpc3 | ENSMUSG00000029465 | 1134 | 39 | 1490 | 168 | 1115 | 115 | 1392 | 88 |
| Arpc5 | ENSMUSG00000008475 | 864 | 45 | 1233 | 156 | 871 | 66 | 1121 | 40 |
| Asap2 | ENSMUSG00000052632 | 656 | 118 | 829 | 131 | 621 | 58 | 818 | 121 |
| Aspm | ENSMUSG00000033952 | 46 | 20 | 147 | 29 | 43 | 10 | 129 | 39 |
| Atad2 | ENSMUSG00000022360 | 124 | 14 | 243 | 23 | 132 | 24 | 212 | 55 |
| Atf4 | ENSMUSG00000042406 | 1171 | 85 | 1374 | 69 | 1169 | 114 | 1319 | 150 |
| Atp10a | ENSMUSG00000025324 | 58 | 4 | 114 | 20 | 63 | 11 | 98 | 18 |
| Atp8a2 | ENSMUSG00000021983 | 247 | 26 | 365 | 72 | 234 | 36 | 355 | 74 |
| Atp8b1 | ENSMUSG00000039529 | 239 | 30 | 380 | 49 | 213 | 28 | 351 | 75 |
| Aurka | ENSMUSG00000027496 | 18 | 5 | 51 | 16 | 19 | 6 | 46 | 15 |
| Aurkb | ENSMUSG00000020897 | 18 | 6 | 47 | 18 | 16 | 5 | 39 | 13 |
| Axl | ENSMUSG00000002602 | 1982 | 499 | 2556 | 293 | 1813 | 317 | 2326 | 298 |
| B3galnt1 | ENSMUSG00000043300 | 50 | 8 | 82 | 21 | 50 | 8 | 80 | 7 |
| Basp1 | ENSMUSG00000045763 | 41 | 7 | 103 | 33 | 42 | 3 | 76 | 15 |

| | | | | | | | | | |
|---|---|---|---|---|---|---|---|---|---|
| BC028528 | ENSMUSG00000038543 | 144 | 14 | 227 | 38 | 130 | 24 | 206 | 24 |
| Bgn | ENSMUSG00000031375 | 5175 | 520 | 12514 | 5386 | 5152 | 464 | 9491 | 2127 |
| Birc5 | ENSMUSG00000017716 | 20 | 10 | 66 | 13 | 18 | 6 | 66 | 24 |
| Bmp1 | ENSMUSG00000022098 | 505 | 77 | 897 | 297 | 496 | 46 | 796 | 219 |
| Bub1 | ENSMUSG00000027379 | 12 | 2 | 48 | 14 | 16 | 9 | 44 | 19 |
| Bub1b | ENSMUSG00000040084 | 41 | 19 | 125 | 41 | 51 | 17 | 103 | 38 |
| C1qa | ENSMUSG00000036887 | 895 | 96 | 1523 | 250 | 883 | 97 | 1362 | 183 |
| C1qb | ENSMUSG00000036905 | 826 | 111 | 1513 | 429 | 782 | 128 | 1330 | 245 |
| C1qc | ENSMUSG00000036896 | 886 | 106 | 1454 | 199 | 845 | 131 | 1290 | 171 |
| C1qtnf6 | ENSMUSG00000022440 | 122 | 21 | 426 | 250 | 120 | 11 | 348 | 107 |
| C4b | ENSMUSG00000073418 | 231 | 28 | 604 | 141 | 295 | 41 | 861 | 703 |
| Cald1 | ENSMUSG00000029761 | 1649 | 323 | 2616 | 191 | 1624 | 256 | 2205 | 422 |
| Cap1 | ENSMUSG00000028656 | 1199 | 155 | 1633 | 55 | 1115 | 167 | 1503 | 127 |
| Capg | ENSMUSG00000056737 | 183 | 29 | 364 | 36 | 182 | 36 | 328 | 70 |
| Capza1 | ENSMUSG00000070372 | 1251 | 78 | 1608 | 138 | 1166 | 148 | 1428 | 124 |
| Carhsp1 | ENSMUSG00000008393 | 592 | 69 | 889 | 70 | 517 | 68 | 796 | 96 |
| Casp4 | ENSMUSG00000033538 | 79 | 11 | 119 | 19 | 74 | 18 | 113 | 27 |
| Casp8 | ENSMUSG00000026029 | 191 | 12 | 303 | 44 | 192 | 23 | 274 | 37 |
| Cavin3 | ENSMUSG00000037060 | 300 | 26 | 506 | 43 | 295 | 23 | 430 | 88 |
| Ccdc80 | ENSMUSG00000022665 | 1816 | 295 | 3479 | 1261 | 1843 | 168 | 2847 | 551 |
| Ccl8 | ENSMUSG00000009185 | 11 | 7 | 75 | 31 | 17 | 9 | 73 | 59 |
| Ccna2 | ENSMUSG00000027715 | 65 | 39 | 245 | 89 | 48 | 9 | 184 | 81 |
| Ccnb1 | ENSMUSG00000041431 | 24 | 11 | 73 | 20 | 14 | 4 | 71 | 30 |
| Ccnb2 | ENSMUSG00000032218 | 27 | 10 | 82 | 39 | 18 | 5 | 72 | 21 |
| Cd109 | ENSMUSG00000046186 | 101 | 22 | 257 | 86 | 92 | 18 | 181 | 52 |
| Cd14 | ENSMUSG00000051439 | 89 | 13 | 165 | 34 | 94 | 6 | 143 | 26 |
| Cd248 | ENSMUSG00000056481 | 285 | 55 | 497 | 49 | 278 | 22 | 431 | 86 |
| Cd300c2 | ENSMUSG00000044811 | 33 | 5 | 109 | 72 | 32 | 8 | 66 | 20 |
| Cd300ld | ENSMUSG00000034641 | 151 | 30 | 245 | 51 | 133 | 23 | 238 | 49 |
| Cd34 | ENSMUSG00000016494 | 3352 | 231 | 5107 | 337 | 3106 | 396 | 4820 | 637 |
| Cd44 | ENSMUSG00000005087 | 292 | 53 | 505 | 44 | 273 | 42 | 409 | 82 |
| Cd48 | ENSMUSG00000015355 | 78 | 10 | 145 | 48 | 67 | 12 | 120 | 19 |
| Cd68 | ENSMUSG00000018774 | 205 | 25 | 321 | 48 | 181 | 27 | 308 | 61 |
| Cd72 | ENSMUSG00000028459 | 30 | 9 | 159 | 128 | 27 | 7 | 92 | 56 |
| Cd84 | ENSMUSG00000038147 | 88 | 19 | 178 | 51 | 80 | 9 | 164 | 32 |
| Cd9 | ENSMUSG00000030342 | 590 | 47 | 756 | 24 | 581 | 71 | 670 | 65 |
| Cd93 | ENSMUSG00000027435 | 3708 | 657 | 5435 | 821 | 3856 | 571 | 5157 | 633 |
| Cdc20 | ENSMUSG00000006398 | 28 | 11 | 91 | 19 | 23 | 7 | 80 | 21 |
| Cdca3 | ENSMUSG00000023505 | 16 | 5 | 56 | 20 | 12 | 5 | 59 | 28 |
| Cdca5 | ENSMUSG00000024791 | 5 | 2 | 21 | 8 | 5 | 3 | 21 | 10 |
| Cdca8 | ENSMUSG00000028873 | 14 | 3 | 49 | 5 | 13 | 6 | 38 | 18 |
| Cdk1 | ENSMUSG00000019942 | 33 | 11 | 194 | 49 | 35 | 17 | 130 | 47 |
| Cdkn1a | ENSMUSG00000023067 | 351 | 153 | 609 | 89 | 367 | 175 | 544 | 152 |
| Cenpe | ENSMUSG00000045328 | 62 | 41 | 172 | 56 | 38 | 21 | 116 | 38 |
| Cep55 | ENSMUSG00000024989 | 17 | 5 | 63 | 15 | 12 | 2 | 58 | 27 |
| Cfb | ENSMUSG00000090231 | 36 | 7 | 140 | 61 | 33 | 15 | 114 | 78 |
| Cfl1 | ENSMUSG00000056201 | 2052 | 87 | 2670 | 304 | 1872 | 236 | 2490 | 204 |
| Ch25h | ENSMUSG00000050370 | 11 | 6 | 32 | 14 | 13 | 5 | 34 | 17 |
| Chd9 | ENSMUSG00000056608 | 1089 | 68 | 1541 | 297 | 1030 | 53 | 1413 | 164 |
| Cilp | ENSMUSG00000042254 | 294 | 76 | 3092 | 3273 | 401 | 134 | 2249 | 1515 |
| Ckap2 | ENSMUSG00000037725 | 33 | 16 | 131 | 46 | 25 | 5 | 95 | 45 |
| Ckap2l | ENSMUSG00000048327 | 30 | 10 | 123 | 39 | 25 | 8 | 110 | 29 |
| Ckap4 | ENSMUSG00000046841 | 756 | 159 | 1094 | 75 | 815 | 103 | 1104 | 165 |
| Cks2 | ENSMUSG00000062248 | 13 | 1 | 50 | 13 | 9 | 2 | 35 | 11 |
| Clec4d | ENSMUSG00000030144 | 9 | 3 | 26 | 14 | 4 | 2 | 18 | 9 |
| Clec4n | ENSMUSG00000023349 | 32 | 6 | 110 | 48 | 34 | 9 | 86 | 29 |
| Clec5a | ENSMUSG00000029915 | 57 | 11 | 99 | 10 | 54 | 23 | 89 | 10 |
| Clic1 | ENSMUSG00000007041 | 641 | 18 | 1053 | 159 | 606 | 66 | 943 | 94 |
| Cmtm3 | ENSMUSG00000031875 | 323 | 45 | 510 | 56 | 301 | 38 | 439 | 77 |
| Cnn3 | ENSMUSG00000053931 | 1009 | 125 | 1438 | 123 | 955 | 121 | 1319 | 119 |
| Cnot6 | ENSMUSG00000020362 | 832 | 68 | 999 | 41 | 747 | 64 | 909 | 122 |
| Col12a1 | ENSMUSG00000032332 | 47 | 12 | 495 | 623 | 42 | 10 | 315 | 288 |
| Col14a1 | ENSMUSG00000022371 | 466 | 75 | 1493 | 1179 | 377 | 75 | 1075 | 556 |
| Col15a1 | ENSMUSG00000028339 | 2833 | 383 | 5668 | 1135 | 3175 | 509 | 5335 | 1392 |

| | | | | | | | | | |
|---|---|---|---|---|---|---|---|---|---|
| Col18a1 | ENSMUSG00000001435 | 248 | 28 | 790 | 244 | 293 | 35 | 541 | 181 |
| Col3a1 | ENSMUSG00000026043 | 5494 | 744 | 25906 | 20347 | 5871 | 824 | 16747 | 9657 |
| Col4a1 | ENSMUSG00000031502 | 12856 | 2211 | 23909 | 1751 | 13123 | 855 | 20837 | 3837 |
| Col4a2 | ENSMUSG00000031503 | 9490 | 1105 | 15270 | 591 | 9814 | 526 | 14603 | 2361 |
| Col4a4 | ENSMUSG00000067158 | 345 | 73 | 497 | 114 | 364 | 54 | 494 | 37 |
| Col5a1 | ENSMUSG00000026837 | 1205 | 173 | 3410 | 1633 | 1238 | 85 | 2682 | 1054 |
| Col5a2 | ENSMUSG00000026042 | 878 | 65 | 4055 | 3237 | 927 | 105 | 2738 | 1563 |
| Col6a1 | ENSMUSG00000001119 | 2071 | 252 | 3943 | 1376 | 2047 | 155 | 3242 | 704 |
| Col6a2 | ENSMUSG00000020241 | 2038 | 291 | 3824 | 981 | 2062 | 139 | 3256 | 743 |
| Col6a3 | ENSMUSG00000048126 | 1170 | 165 | 2685 | 1137 | 1119 | 203 | 1895 | 452 |
| Col8a1 | ENSMUSG00000068196 | 704 | 96 | 3461 | 2170 | 747 | 153 | 2473 | 1081 |
| Cotl1 | ENSMUSG00000031827 | 234 | 40 | 369 | 66 | 208 | 50 | 335 | 22 |
| Creb5 | ENSMUSG00000053007 | 235 | 36 | 347 | 57 | 199 | 28 | 323 | 65 |
| Crip1 | ENSMUSG00000006360 | 686 | 77 | 907 | 72 | 645 | 76 | 846 | 125 |
| Crlf1 | ENSMUSG00000007888 | 11 | 7 | 82 | 61 | 12 | 6 | 50 | 29 |
| Csf1r | ENSMUSG00000024621 | 1029 | 156 | 1681 | 243 | 1120 | 141 | 1491 | 195 |
| Csrp2 | ENSMUSG00000020186 | 180 | 37 | 542 | 354 | 165 | 20 | 315 | 127 |
| Ctgf | ENSMUSG00000019997 | 1416 | 407 | 4434 | 1114 | 1381 | 216 | 3773 | 916 |
| Ctla2a | ENSMUSG00000044258 | 445 | 55 | 611 | 63 | 463 | 136 | 614 | 55 |
| Ctsc | ENSMUSG00000030560 | 1077 | 100 | 1506 | 162 | 1055 | 115 | 1329 | 214 |
| Ctsz | ENSMUSG00000016256 | 454 | 61 | 730 | 152 | 421 | 29 | 666 | 78 |
| Cxcl16 | ENSMUSG00000018920 | 209 | 30 | 426 | 208 | 210 | 55 | 332 | 47 |
| Dab2 | ENSMUSG00000022150 | 1262 | 200 | 1666 | 115 | 1122 | 167 | 1571 | 239 |
| Dbn1 | ENSMUSG00000034675 | 170 | 35 | 385 | 71 | 169 | 25 | 330 | 74 |
| Dchs1 | ENSMUSG00000036862 | 677 | 75 | 906 | 45 | 647 | 56 | 875 | 104 |
| Depdc1a | ENSMUSG00000028175 | 7 | 6 | 46 | 33 | 10 | 5 | 34 | 14 |
| Diaph3 | ENSMUSG00000022021 | 16 | 5 | 59 | 16 | 23 | 10 | 55 | 26 |
| Dio2 | ENSMUSG00000007682 | 59 | 18 | 224 | 44 | 57 | 23 | 165 | 64 |
| Dlgap5 | ENSMUSG00000037544 | 17 | 6 | 48 | 15 | 12 | 6 | 41 | 13 |
| Dpysl3 | ENSMUSG00000024501 | 1022 | 66 | 1824 | 480 | 1008 | 79 | 1535 | 363 |
| Dtl | ENSMUSG00000037474 | 13 | 10 | 42 | 13 | 12 | 3 | 42 | 11 |
| Dynll1 | ENSMUSG00000009013 | 571 | 82 | 870 | 205 | 474 | 104 | 727 | 130 |
| E2f1 | ENSMUSG00000027490 | 21 | 5 | 59 | 13 | 23 | 8 | 56 | 20 |
| E2f7 | ENSMUSG00000020185 | 42 | 13 | 85 | 32 | 34 | 13 | 79 | 29 |
| Ecm1 | ENSMUSG00000028108 | 452 | 86 | 696 | 127 | 464 | 82 | 720 | 136 |
| Ecscr | ENSMUSG00000073599 | 254 | 10 | 366 | 51 | 246 | 20 | 337 | 40 |
| Ect2 | ENSMUSG00000027699 | 21 | 8 | 80 | 27 | 24 | 9 | 75 | 24 |
| Edem1 | ENSMUSG00000030104 | 377 | 69 | 588 | 97 | 278 | 84 | 510 | 147 |
| Eef1a1 | ENSMUSG00000037742 | 19836 | 2225 | 28730 | 7994 | 17884 | 2426 | 23968 | 2075 |
| Efhd2 | ENSMUSG00000040659 | 427 | 47 | 656 | 180 | 426 | 31 | 620 | 78 |
| Elf4 | ENSMUSG00000031103 | 320 | 62 | 496 | 40 | 293 | 54 | 437 | 22 |
| Emilin1 | ENSMUSG00000029163 | 473 | 106 | 835 | 204 | 519 | 28 | 812 | 91 |
| Emp1 | ENSMUSG00000030208 | 1538 | 159 | 4210 | 1587 | 1504 | 221 | 3295 | 791 |
| Emp3 | ENSMUSG00000040212 | 145 | 17 | 216 | 17 | 152 | 17 | 192 | 12 |
| Enah | ENSMUSG00000022995 | 2820 | 352 | 4077 | 860 | 3138 | 213 | 4243 | 467 |
| Endod1 | ENSMUSG00000037419 | 225 | 35 | 387 | 85 | 188 | 20 | 328 | 98 |
| Entpd1 | ENSMUSG00000048120 | 473 | 39 | 717 | 165 | 458 | 60 | 637 | 114 |
| Ereg | ENSMUSG00000029377 | 2 | 1 | 20 | 9 | 2 | 1 | 13 | 5 |
| Esco2 | ENSMUSG00000022034 | 12 | 5 | 43 | 10 | 11 | 6 | 36 | 16 |
| Esyt1 | ENSMUSG00000025366 | 578 | 37 | 738 | 64 | 571 | 38 | 697 | 35 |
| F2r | ENSMUSG00000048376 | 819 | 41 | 1217 | 181 | 861 | 54 | 1112 | 37 |
| F2rl1 | ENSMUSG00000021678 | 27 | 6 | 56 | 22 | 20 | 7 | 44 | 13 |
| Fads1 | ENSMUSG00000010663 | 476 | 36 | 619 | 11 | 473 | 39 | 586 | 42 |
| Fam111a | ENSMUSG00000024691 | 311 | 25 | 543 | 98 | 278 | 57 | 446 | 66 |
| Fam114a1 | ENSMUSG00000029185 | 316 | 44 | 511 | 131 | 316 | 48 | 398 | 46 |
| Fam129b | ENSMUSG00000026796 | 610 | 83 | 785 | 85 | 572 | 21 | 715 | 78 |
| Fam198b | ENSMUSG00000027955 | 1065 | 183 | 1845 | 423 | 1058 | 202 | 1912 | 390 |
| Fbln2 | ENSMUSG00000064080 | 1827 | 201 | 3200 | 493 | 1966 | 214 | 3008 | 422 |
| Fbn1 | ENSMUSG00000027204 | 2671 | 543 | 7690 | 2514 | 3011 | 303 | 6532 | 2277 |
| Fcgr2b | ENSMUSG00000026656 | 275 | 47 | 507 | 137 | 255 | 41 | 437 | 94 |
| Fcgr3 | ENSMUSG00000059498 | 355 | 58 | 659 | 143 | 324 | 56 | 543 | 112 |
| Fcgr4 | ENSMUSG00000059089 | 20 | 9 | 70 | 53 | 17 | 3 | 41 | 16 |
| Fcrls | ENSMUSG00000015852 | 197 | 22 | 463 | 184 | 186 | 29 | 357 | 109 |
| Fignl1 | ENSMUSG00000035455 | 26 | 6 | 57 | 22 | 21 | 4 | 51 | 17 |

| | | | | | | | | | |
|---|---|---|---|---|---|---|---|---|---|
| Filip1l | ENSMUSG00000043336 | 880 | 82 | 1259 | 105 | 842 | 147 | 1084 | 69 |
| Fkbp10 | ENSMUSG00000001555 | 345 | 47 | 461 | 58 | 288 | 35 | 402 | 56 |
| Flna | ENSMUSG00000031328 | 4014 | 455 | 5769 | 673 | 3783 | 774 | 4951 | 309 |
| Fmnl3 | ENSMUSG00000023008 | 757 | 91 | 963 | 63 | 730 | 56 | 931 | 86 |
| Fndc1 | ENSMUSG00000071984 | 473 | 60 | 1081 | 567 | 452 | 50 | 911 | 402 |
| Foxm1 | ENSMUSG00000001517 | 42 | 12 | 122 | 30 | 47 | 13 | 101 | 53 |
| Frzb | ENSMUSG00000027004 | 70 | 18 | 272 | 183 | 82 | 24 | 208 | 55 |
| Fscn1 | ENSMUSG00000029581 | 817 | 47 | 1186 | 92 | 831 | 115 | 1140 | 241 |
| Fstl1 | ENSMUSG00000022816 | 2281 | 246 | 7581 | 4030 | 2354 | 314 | 5589 | 1854 |
| Fstl3 | ENSMUSG00000020325 | 58 | 10 | 114 | 21 | 45 | 10 | 101 | 24 |
| Fuca2 | ENSMUSG00000019810 | 4743 | 786 | 5774 | 464 | 4738 | 302 | 6270 | 543 |
| Fxyd5 | ENSMUSG00000009687 | 283 | 49 | 603 | 81 | 271 | 56 | 473 | 80 |
| Gab2 | ENSMUSG00000004508 | 804 | 49 | 1077 | 134 | 805 | 51 | 1017 | 130 |
| Gas2l3 | ENSMUSG00000074802 | 49 | 17 | 117 | 44 | 58 | 17 | 123 | 41 |
| Gdf6 | ENSMUSG00000051279 | 21 | 6 | 60 | 18 | 20 | 4 | 52 | 18 |
| Glipr2 | ENSMUSG00000028480 | 116 | 26 | 213 | 33 | 120 | 25 | 192 | 33 |
| Gm42417 | ENSMUSG00000109510 | 601 | 184 | 1052 | 250 | 572 | 93 | 1085 | 285 |
| Gm47302 | ENSMUSG00000105211 | 24 | 9 | 110 | 97 | 27 | 2 | 68 | 30 |
| Gm4739 | ENSMUSG00000112808 | 128 | 31 | 209 | 69 | 125 | 23 | 223 | 29 |
| Gng2 | ENSMUSG00000043004 | 106 | 30 | 205 | 31 | 90 | 14 | 161 | 60 |
| Gpr153 | ENSMUSG00000042804 | 289 | 39 | 472 | 87 | 284 | 32 | 390 | 64 |
| Gprc5b | ENSMUSG00000008734 | 417 | 88 | 644 | 60 | 371 | 45 | 622 | 68 |
| Gpx1 | ENSMUSG00000063856 | 1007 | 59 | 1482 | 189 | 1015 | 47 | 1424 | 176 |
| Grb10 | ENSMUSG00000020176 | 1827 | 116 | 2380 | 316 | 1880 | 141 | 2271 | 196 |
| Grn | ENSMUSG00000034708 | 1349 | 120 | 1891 | 216 | 1368 | 123 | 1853 | 161 |
| Gtse1 | ENSMUSG00000022385 | 12 | 4 | 37 | 7 | 11 | 3 | 35 | 14 |
| Gusb | ENSMUSG00000025534 | 484 | 36 | 644 | 46 | 462 | 48 | 631 | 40 |
| Haspin | ENSMUSG00000050107 | 7 | 3 | 26 | 4 | 7 | 2 | 20 | 10 |
| Hcls1 | ENSMUSG00000022831 | 234 | 38 | 352 | 48 | 191 | 20 | 308 | 52 |
| Hectd2os | ENSMUSG00000087579 | 181 | 28 | 259 | 45 | 192 | 32 | 288 | 75 |
| Hells | ENSMUSG00000025001 | 29 | 11 | 78 | 18 | 36 | 7 | 79 | 13 |
| Hhipl1 | ENSMUSG00000021260 | 67 | 22 | 127 | 40 | 63 | 22 | 109 | 33 |
| Hist1h2ap | ENSMUSG00000094777 | 37 | 19 | 118 | 39 | 33 | 8 | 97 | 65 |
| Hmmr | ENSMUSG00000020330 | 21 | 9 | 98 | 40 | 21 | 6 | 82 | 33 |
| Hspa1l | ENSMUSG00000007033 | 134 | 25 | 210 | 54 | 130 | 20 | 216 | 41 |
| Ifi204 | ENSMUSG00000073489 | 206 | 34 | 433 | 95 | 196 | 40 | 334 | 43 |
| Ifi27l2a | ENSMUSG00000079017 | 237 | 35 | 439 | 84 | 218 | 39 | 420 | 49 |
| Ifitm2 | ENSMUSG00000060591 | 875 | 131 | 1176 | 80 | 841 | 117 | 1097 | 135 |
| Ifitm3 | ENSMUSG00000025492 | 1226 | 258 | 1689 | 136 | 1126 | 147 | 1515 | 182 |
| Ifngr1 | ENSMUSG00000020009 | 1109 | 126 | 1400 | 78 | 1101 | 128 | 1324 | 109 |
| Ift122 | ENSMUSG00000030323 | 278 | 32 | 496 | 94 | 309 | 39 | 488 | 78 |
| Igfbp7 | ENSMUSG00000036256 | 2248 | 254 | 4619 | 1238 | 2198 | 173 | 3722 | 608 |
| Igsf6 | ENSMUSG00000035004 | 49 | 7 | 92 | 27 | 46 | 7 | 79 | 10 |
| Il10ra | ENSMUSG00000032089 | 138 | 27 | 237 | 86 | 108 | 21 | 194 | 46 |
| Il1rl2 | ENSMUSG00000070942 | 95 | 8 | 139 | 17 | 75 | 7 | 130 | 22 |
| Il2rg | ENSMUSG00000031304 | 249 | 26 | 354 | 25 | 214 | 37 | 325 | 48 |
| Il4ra | ENSMUSG00000030748 | 359 | 66 | 598 | 83 | 333 | 48 | 463 | 59 |
| Incenp | ENSMUSG00000024660 | 92 | 17 | 144 | 29 | 81 | 14 | 144 | 40 |
| Inhba | ENSMUSG00000041324 | 65 | 15 | 199 | 57 | 61 | 18 | 178 | 55 |
| Iqgap1 | ENSMUSG00000030536 | 1656 | 342 | 2416 | 368 | 1438 | 279 | 2181 | 405 |
| Iqgap3 | ENSMUSG00000028068 | 31 | 20 | 115 | 32 | 27 | 10 | 119 | 46 |
| Irf7 | ENSMUSG00000025498 | 220 | 65 | 413 | 114 | 189 | 26 | 332 | 52 |
| Itga5 | ENSMUSG00000000555 | 986 | 110 | 1409 | 68 | 945 | 123 | 1366 | 142 |
| Itga9 | ENSMUSG00000039115 | 1864 | 139 | 2467 | 423 | 1953 | 131 | 2644 | 386 |
| Itgam | ENSMUSG00000030786 | 190 | 26 | 365 | 31 | 200 | 68 | 269 | 58 |
| Itih5 | ENSMUSG00000025780 | 368 | 63 | 749 | 273 | 377 | 54 | 597 | 122 |
| Itpripl2 | ENSMUSG00000095115 | 1256 | 118 | 1751 | 114 | 1222 | 89 | 1508 | 115 |
| Kcne4 | ENSMUSG00000047330 | 64 | 17 | 119 | 36 | 59 | 9 | 95 | 7 |
| Kctd11 | ENSMUSG00000046731 | 99 | 17 | 174 | 7 | 96 | 16 | 157 | 31 |
| Kctd17 | ENSMUSG00000033287 | 312 | 32 | 468 | 35 | 317 | 29 | 437 | 78 |
| Kif11 | ENSMUSG00000012443 | 54 | 18 | 219 | 84 | 56 | 14 | 169 | 71 |
| Kif15 | ENSMUSG00000036768 | 14 | 6 | 40 | 11 | 16 | 8 | 57 | 20 |
| Kif18b | ENSMUSG00000051378 | 13 | 3 | 50 | 17 | 14 | 3 | 49 | 16 |
| Kif20a | ENSMUSG00000003779 | 45 | 12 | 135 | 43 | 44 | 5 | 120 | 29 |

| | | | | | | | | | |
|---|---|---|---|---|---|---|---|---|---|
| Kif22 | ENSMUSG00000030677 | 26 | 8 | 61 | 17 | 19 | 10 | 51 | 20 |
| Kif23 | ENSMUSG00000032254 | 53 | 21 | 153 | 63 | 39 | 12 | 144 | 59 |
| Kif2c | ENSMUSG00000028678 | 15 | 5 | 39 | 8 | 8 | 2 | 33 | 12 |
| Kif4 | ENSMUSG00000034311 | 35 | 11 | 86 | 21 | 32 | 13 | 86 | 26 |
| Kif5b | ENSMUSG00000006740 | 4524 | 195 | 5272 | 357 | 4610 | 339 | 5543 | 496 |
| Kifc1 | ENSMUSG00000079553 | 21 | 17 | 44 | 7 | 18 | 7 | 48 | 18 |
| Knl1 | ENSMUSG00000027326 | 29 | 18 | 66 | 14 | 21 | 8 | 59 | 16 |
| Knstrn | ENSMUSG00000027331 | 28 | 8 | 90 | 28 | 24 | 3 | 76 | 21 |
| Kntc1 | ENSMUSG00000029414 | 13 | 8 | 61 | 43 | 19 | 11 | 63 | 21 |
| Lama4 | ENSMUSG00000019846 | 2464 | 138 | 3226 | 192 | 2291 | 264 | 3050 | 270 |
| Lamb1 | ENSMUSG00000002900 | 2656 | 308 | 3500 | 191 | 2690 | 265 | 3282 | 467 |
| Lamc1 | ENSMUSG00000026478 | 4969 | 490 | 7157 | 380 | 5155 | 283 | 6688 | 486 |
| Laptm5 | ENSMUSG00000028581 | 500 | 82 | 885 | 205 | 478 | 56 | 708 | 122 |
| Lcp1 | ENSMUSG00000021998 | 803 | 118 | 1421 | 200 | 781 | 112 | 1229 | 174 |
| Lgals3 | ENSMUSG00000050335 | 56 | 18 | 203 | 128 | 57 | 20 | 148 | 73 |
| Lgals3bp | ENSMUSG00000033880 | 711 | 128 | 1195 | 145 | 721 | 83 | 1079 | 113 |
| Lgals9 | ENSMUSG00000001123 | 575 | 96 | 870 | 73 | 533 | 82 | 766 | 58 |
| Lhfpl2 | ENSMUSG00000045312 | 121 | 17 | 351 | 198 | 128 | 21 | 269 | 96 |
| Lilr4b | ENSMUSG00000112023 | 154 | 32 | 309 | 106 | 138 | 41 | 252 | 97 |
| Litaf | ENSMUSG00000022500 | 341 | 97 | 525 | 67 | 315 | 74 | 445 | 72 |
| Lman1l | ENSMUSG00000056271 | 6 | 2 | 21 | 5 | 6 | 5 | 23 | 11 |
| Lockd | ENSMUSG00000098318 | 4 | 4 | 22 | 11 | 5 | 3 | 21 | 12 |
| Loxl1 | ENSMUSG00000032334 | 618 | 97 | 1447 | 604 | 687 | 57 | 1181 | 257 |
| Loxl2 | ENSMUSG00000034205 | 724 | 133 | 1635 | 127 | 683 | 84 | 1403 | 383 |
| Lrp1 | ENSMUSG00000040249 | 4010 | 495 | 5574 | 985 | 4250 | 358 | 5472 | 456 |
| Lrp8 | ENSMUSG00000028613 | 9 | 5 | 40 | 13 | 13 | 7 | 51 | 44 |
| Ly6e | ENSMUSG00000022587 | 2929 | 530 | 3882 | 293 | 2782 | 305 | 3608 | 375 |
| Ly86 | ENSMUSG00000021423 | 93 | 6 | 221 | 112 | 92 | 20 | 181 | 43 |
| Lyz2 | ENSMUSG00000069516 | 4037 | 355 | 6768 | 1318 | 3852 | 675 | 5981 | 848 |
| Mall | ENSMUSG00000027377 | 160 | 25 | 233 | 27 | 157 | 20 | 226 | 33 |
| Map1b | ENSMUSG00000052727 | 582 | 102 | 873 | 51 | 444 | 82 | 777 | 113 |
| Map4k4 | ENSMUSG00000026074 | 3034 | 462 | 3619 | 383 | 2841 | 174 | 3784 | 234 |
| Marcksl1 | ENSMUSG00000047945 | 104 | 18 | 211 | 70 | 107 | 9 | 183 | 4 |
| Masp1 | ENSMUSG00000022887 | 205 | 16 | 307 | 114 | 219 | 36 | 315 | 61 |
| Mcam | ENSMUSG00000032135 | 1064 | 99 | 1678 | 301 | 970 | 129 | 1515 | 234 |
| Medag | ENSMUSG00000029659 | 437 | 58 | 771 | 121 | 486 | 78 | 684 | 41 |
| Melk | ENSMUSG00000035683 | 19 | 16 | 45 | 19 | 13 | 4 | 40 | 14 |
| Meox1 | ENSMUSG00000001493 | 296 | 46 | 905 | 284 | 323 | 49 | 767 | 246 |
| Mest | ENSMUSG00000051855 | 193 | 22 | 598 | 325 | 156 | 26 | 417 | 154 |
| Mfap4 | ENSMUSG00000042436 | 202 | 18 | 1249 | 1153 | 245 | 36 | 712 | 324 |
| Mfap5 | ENSMUSG00000030116 | 411 | 43 | 1634 | 1033 | 404 | 45 | 1132 | 430 |
| Mis18bp1 | ENSMUSG00000047534 | 15 | 3 | 59 | 20 | 16 | 6 | 44 | 14 |
| Mki67 | ENSMUSG00000031004 | 305 | 108 | 1186 | 340 | 284 | 62 | 1009 | 362 |
| Mmp2 | ENSMUSG00000031740 | 991 | 124 | 2606 | 1435 | 1013 | 93 | 2013 | 656 |
| Mmp23 | ENSMUSG00000029061 | 80 | 8 | 179 | 97 | 80 | 19 | 146 | 38 |
| Mpp1 | ENSMUSG00000031402 | 302 | 30 | 434 | 51 | 294 | 38 | 387 | 21 |
| Mrc1 | ENSMUSG00000026712 | 1125 | 59 | 1641 | 55 | 1195 | 105 | 1622 | 319 |
| Ms4a6b | ENSMUSG00000024677 | 163 | 35 | 265 | 79 | 109 | 27 | 237 | 58 |
| Ms4a6c | ENSMUSG00000079419 | 117 | 24 | 273 | 106 | 108 | 30 | 198 | 50 |
| Msn | ENSMUSG00000031207 | 4381 | 327 | 6174 | 266 | 4286 | 400 | 5734 | 498 |
| Msr1 | ENSMUSG00000025044 | 91 | 16 | 165 | 30 | 95 | 12 | 158 | 35 |
| Mxd3 | ENSMUSG00000021485 | 3 | 2 | 21 | 8 | 5 | 2 | 23 | 10 |
| Mxra7 | ENSMUSG00000020814 | 353 | 43 | 549 | 114 | 334 | 33 | 485 | 81 |
| Mybl2 | ENSMUSG00000017861 | 12 | 4 | 32 | 11 | 10 | 3 | 29 | 7 |
| Mybpc2 | ENSMUSG00000038670 | 292 | 36 | 901 | 354 | 321 | 110 | 773 | 309 |
| Myh10 | ENSMUSG00000020900 | 964 | 132 | 1462 | 186 | 988 | 96 | 1366 | 213 |
| Myl1 | ENSMUSG00000061816 | 777 | 84 | 1354 | 639 | 674 | 113 | 1186 | 282 |
| Myl6 | ENSMUSG00000090841 | 3039 | 174 | 4606 | 743 | 2959 | 365 | 4098 | 492 |
| Myo1d | ENSMUSG00000035441 | 367 | 44 | 489 | 47 | 360 | 56 | 474 | 39 |
| Myo1f | ENSMUSG00000024300 | 97 | 11 | 170 | 34 | 88 | 10 | 163 | 43 |
| Myof | ENSMUSG00000048612 | 346 | 89 | 640 | 167 | 312 | 56 | 541 | 78 |
| Nbl1 | ENSMUSG00000041120 | 214 | 26 | 349 | 63 | 215 | 19 | 307 | 61 |
| Ncapg | ENSMUSG00000015880 | 17 | 11 | 88 | 26 | 18 | 10 | 55 | 17 |
| Ncapg2 | ENSMUSG00000042029 | 78 | 9 | 184 | 50 | 101 | 31 | 166 | 54 |

| | | | | | | | | | |
|---|---|---|---|---|---|---|---|---|---|
| Ncaph | ENSMUSG00000034906 | 32 | 7 | 80 | 27 | 35 | 11 | 78 | 26 |
| Ncf1 | ENSMUSG00000015950 | 112 | 33 | 201 | 54 | 124 | 31 | 190 | 42 |
| Nckap1l | ENSMUSG00000022488 | 260 | 42 | 511 | 159 | 240 | 40 | 412 | 73 |
| Ndc80 | ENSMUSG00000024056 | 18 | 8 | 69 | 23 | 19 | 4 | 56 | 15 |
| Necap2 | ENSMUSG00000028923 | 266 | 29 | 348 | 33 | 232 | 24 | 305 | 28 |
| Nek2 | ENSMUSG00000026622 | 16 | 10 | 60 | 22 | 15 | 3 | 46 | 11 |
| Nes | ENSMUSG00000004891 | 1779 | 216 | 2677 | 688 | 1621 | 280 | 2378 | 347 |
| Nid1 | ENSMUSG00000005397 | 3926 | 530 | 6347 | 1387 | 3634 | 332 | 5323 | 725 |
| Nid2 | ENSMUSG00000021806 | 561 | 130 | 914 | 118 | 537 | 62 | 895 | 114 |
| Nkd2 | ENSMUSG00000021567 | 39 | 11 | 118 | 51 | 36 | 6 | 77 | 24 |
| Nlrc3 | ENSMUSG00000049871 | 58 | 17 | 139 | 45 | 66 | 12 | 140 | 43 |
| Nppa | ENSMUSG00000041616 | 2919 | 1471 | 17568 | 21348 | 2998 | 848 | 11493 | 6461 |
| Nppb | ENSMUSG00000029019 | 3896 | 1226 | 10218 | 3100 | 4287 | 995 | 11088 | 3450 |
| Nuf2 | ENSMUSG00000026683 | 21 | 9 | 86 | 40 | 16 | 4 | 77 | 34 |
| Oaf | ENSMUSG00000032014 | 264 | 77 | 399 | 146 | 208 | 41 | 331 | 55 |
| Olfml3 | ENSMUSG00000027848 | 303 | 52 | 492 | 108 | 323 | 21 | 470 | 29 |
| Otulin | ENSMUSG00000046034 | 372 | 28 | 492 | 30 | 374 | 38 | 498 | 73 |
| P2ry6 | ENSMUSG00000048779 | 103 | 15 | 191 | 33 | 95 | 15 | 152 | 28 |
| P3h3 | ENSMUSG00000023191 | 185 | 14 | 319 | 100 | 179 | 28 | 278 | 54 |
| Pabpc1 | ENSMUSG00000022283 | 2750 | 396 | 4424 | 589 | 2940 | 313 | 3983 | 663 |
| Pam | ENSMUSG00000026335 | 13364 | 1232 | 16353 | 2118 | 13089 | 520 | 17039 | 1382 |
| Pamr1 | ENSMUSG00000027188 | 50 | 16 | 271 | 244 | 44 | 23 | 183 | 117 |
| Pbk | ENSMUSG00000022033 | 18 | 15 | 71 | 30 | 16 | 9 | 69 | 28 |
| Pcdhgc3 | ENSMUSG00000102918 | 783 | 97 | 981 | 69 | 752 | 51 | 988 | 95 |
| Pclaf | ENSMUSG00000040204 | 20 | 11 | 70 | 26 | 13 | 4 | 61 | 24 |
| Pcolce | ENSMUSG00000029718 | 778 | 81 | 1236 | 178 | 675 | 91 | 1133 | 173 |
| Pea15a | ENSMUSG00000013698 | 2146 | 275 | 2690 | 102 | 2035 | 242 | 2530 | 274 |
| Pfkp | ENSMUSG00000021196 | 1083 | 101 | 1607 | 246 | 1116 | 108 | 1508 | 216 |
| Pfn1 | ENSMUSG00000018293 | 2349 | 50 | 2712 | 75 | 2362 | 104 | 2642 | 191 |
| Phf11d | ENSMUSG00000068245 | 263 | 46 | 403 | 78 | 242 | 30 | 341 | 41 |
| Phlda3 | ENSMUSG00000041801 | 128 | 18 | 222 | 62 | 128 | 4 | 216 | 43 |
| Pi16 | ENSMUSG00000024011 | 1042 | 154 | 1987 | 337 | 1030 | 148 | 1793 | 499 |
| Picalm | ENSMUSG00000039361 | 3682 | 311 | 4950 | 606 | 3495 | 243 | 4537 | 508 |
| Pimreg | ENSMUSG00000020808 | 18 | 10 | 79 | 24 | 18 | 4 | 67 | 18 |
| Pirb | ENSMUSG00000058818 | 142 | 22 | 283 | 84 | 134 | 30 | 266 | 92 |
| Pla2g4a | ENSMUSG00000056220 | 118 | 46 | 201 | 50 | 118 | 39 | 188 | 22 |
| Pla2g7 | ENSMUSG00000023913 | 171 | 21 | 280 | 45 | 172 | 34 | 252 | 45 |
| Plat | ENSMUSG00000031538 | 475 | 62 | 725 | 65 | 464 | 41 | 699 | 128 |
| Plcg2 | ENSMUSG00000034330 | 248 | 18 | 352 | 61 | 256 | 18 | 349 | 46 |
| Plek | ENSMUSG00000020120 | 194 | 26 | 382 | 104 | 207 | 35 | 336 | 73 |
| Plekhg2 | ENSMUSG00000037552 | 415 | 79 | 602 | 42 | 355 | 76 | 498 | 81 |
| Plekhh2 | ENSMUSG00000040852 | 172 | 20 | 250 | 25 | 168 | 11 | 228 | 26 |
| Plekho1 | ENSMUSG00000015745 | 873 | 48 | 1229 | 109 | 927 | 54 | 1262 | 202 |
| Plekho2 | ENSMUSG00000050721 | 497 | 98 | 707 | 90 | 525 | 90 | 666 | 108 |
| Pls3 | ENSMUSG00000016382 | 1718 | 79 | 2436 | 255 | 1640 | 157 | 2091 | 235 |
| Pmepa1 | ENSMUSG00000038400 | 495 | 93 | 798 | 197 | 520 | 37 | 753 | 189 |
| Pmp22 | ENSMUSG00000018217 | 799 | 98 | 1125 | 49 | 732 | 79 | 1029 | 87 |
| Postn | ENSMUSG00000027750 | 1288 | 354 | 16369 | 19036 | 1325 | 317 | 10262 | 8108 |
| Ppic | ENSMUSG00000024538 | 424 | 29 | 954 | 416 | 413 | 41 | 706 | 107 |
| Ppp1r9b | ENSMUSG00000038976 | 1114 | 96 | 1479 | 122 | 1143 | 53 | 1390 | 134 |
| Praf2 | ENSMUSG00000031149 | 91 | 14 | 155 | 27 | 90 | 17 | 137 | 19 |
| Prc1 | ENSMUSG00000038943 | 59 | 10 | 277 | 77 | 74 | 20 | 211 | 83 |
| Prcp | ENSMUSG00000061119 | 378 | 31 | 583 | 113 | 369 | 43 | 525 | 62 |
| Prelid1 | ENSMUSG00000021486 | 540 | 47 | 753 | 58 | 476 | 49 | 647 | 93 |
| Prnd | ENSMUSG00000027338 | 170 | 30 | 416 | 121 | 167 | 33 | 335 | 59 |
| Prr11 | ENSMUSG00000020493 | 26 | 9 | 79 | 21 | 18 | 6 | 65 | 19 |
| Prrg3 | ENSMUSG00000033361 | 304 | 49 | 476 | 60 | 309 | 45 | 455 | 53 |
| Psat1 | ENSMUSG00000024640 | 65 | 15 | 115 | 35 | 66 | 18 | 116 | 28 |
| Psrc1 | ENSMUSG00000068744 | 10 | 4 | 29 | 15 | 9 | 4 | 29 | 18 |
| Ptgfrn | ENSMUSG00000027864 | 1116 | 158 | 1460 | 136 | 1089 | 77 | 1442 | 156 |
| Ptgis | ENSMUSG00000017969 | 202 | 28 | 364 | 114 | 200 | 27 | 323 | 64 |
| Ptma | ENSMUSG00000026238 | 4087 | 376 | 5328 | 376 | 3885 | 518 | 4897 | 607 |
| Ptprj | ENSMUSG00000025314 | 404 | 60 | 631 | 75 | 427 | 56 | 571 | 83 |
| Pxdn | ENSMUSG00000020674 | 2049 | 186 | 2790 | 214 | 2026 | 252 | 2556 | 286 |

| | | | | | | | | | |
|---|---|---|---|---|---|---|---|---|---|
| Qsox1 | ENSMUSG00000033684 | 551 | 17 | 792 | 82 | 588 | 46 | 765 | 102 |
| Rab13 | ENSMUSG00000027935 | 64 | 9 | 113 | 32 | 60 | 13 | 95 | 9 |
| Rab31 | ENSMUSG00000056515 | 515 | 47 | 841 | 272 | 486 | 52 | 743 | 141 |
| Rab7b | ENSMUSG00000052688 | 114 | 17 | 216 | 47 | 113 | 25 | 187 | 28 |
| Racgap1 | ENSMUSG00000023015 | 51 | 13 | 168 | 39 | 47 | 22 | 147 | 53 |
| Rbl1 | ENSMUSG00000027641 | 92 | 18 | 142 | 31 | 88 | 10 | 137 | 16 |
| Rcan1 | ENSMUSG00000022951 | 2025 | 588 | 3577 | 1067 | 2237 | 187 | 3873 | 1294 |
| Rcc2 | ENSMUSG00000040945 | 364 | 24 | 453 | 17 | 354 | 21 | 462 | 57 |
| Rcn3 | ENSMUSG00000019539 | 304 | 49 | 526 | 177 | 289 | 35 | 431 | 74 |
| Rgs16 | ENSMUSG00000026475 | 42 | 13 | 97 | 25 | 41 | 22 | 76 | 17 |
| Rhoc | ENSMUSG00000002233 | 1016 | 82 | 1471 | 147 | 963 | 60 | 1399 | 180 |
| Ripk1 | ENSMUSG00000021408 | 280 | 44 | 414 | 37 | 283 | 47 | 366 | 18 |
| Rnase4 | ENSMUSG00000021876 | 967 | 55 | 1292 | 169 | 990 | 104 | 1212 | 120 |
| Rnd3 | ENSMUSG00000017144 | 446 | 50 | 573 | 42 | 410 | 17 | 542 | 43 |
| Rnf213 | ENSMUSG00000070327 | 1921 | 265 | 2664 | 212 | 1732 | 232 | 2287 | 172 |
| Rnf4 | ENSMUSG00000029110 | 719 | 66 | 881 | 34 | 682 | 79 | 832 | 58 |
| Rrm2 | ENSMUSG00000020649 | 47 | 3 | 106 | 36 | 46 | 11 | 90 | 24 |
| Rsad2 | ENSMUSG00000020641 | 558 | 59 | 808 | 127 | 569 | 74 | 833 | 251 |
| Rsu1 | ENSMUSG00000026727 | 628 | 30 | 848 | 58 | 638 | 25 | 765 | 30 |
| Runx3 | ENSMUSG00000070691 | 15 | 6 | 40 | 18 | 11 | 6 | 33 | 8 |
| S100a10 | ENSMUSG00000041959 | 648 | 48 | 917 | 65 | 558 | 90 | 873 | 124 |
| S100a11 | ENSMUSG00000027907 | 541 | 79 | 1009 | 138 | 513 | 59 | 798 | 136 |
| S100a4 | ENSMUSG00000001020 | 101 | 23 | 185 | 48 | 95 | 20 | 157 | 48 |
| S100a6 | ENSMUSG00000001025 | 463 | 35 | 675 | 47 | 446 | 74 | 610 | 100 |
| Samd9l | ENSMUSG00000047735 | 766 | 77 | 1115 | 179 | 714 | 95 | 898 | 111 |
| Scml4 | ENSMUSG00000044770 | 68 | 22 | 133 | 24 | 73 | 13 | 155 | 38 |
| Sdc3 | ENSMUSG00000025743 | 1679 | 201 | 2152 | 200 | 1603 | 123 | 1955 | 125 |
| Sec61a1 | ENSMUSG00000030082 | 772 | 59 | 1001 | 99 | 760 | 61 | 922 | 32 |
| Sema3f | ENSMUSG00000034684 | 281 | 62 | 437 | 37 | 248 | 21 | 450 | 63 |
| Serpinb1c | ENSMUSG00000079049 | 2 | 1 | 40 | 44 | 2 | 1 | 21 | 10 |
| Serpinf1 | ENSMUSG00000000753 | 424 | 109 | 1021 | 469 | 419 | 39 | 783 | 209 |
| Sgo1 | ENSMUSG00000023940 | 10 | 3 | 30 | 10 | 11 | 4 | 27 | 6 |
| Sh3bgrl3 | ENSMUSG00000028843 | 283 | 45 | 439 | 75 | 294 | 39 | 393 | 63 |
| Sh3bp2 | ENSMUSG00000054520 | 69 | 26 | 114 | 31 | 61 | 7 | 106 | 12 |
| Sh3pxd2b | ENSMUSG00000040711 | 249 | 93 | 530 | 157 | 247 | 52 | 411 | 85 |
| Siglec1 | ENSMUSG00000027322 | 120 | 21 | 202 | 53 | 128 | 32 | 195 | 28 |
| Slamf9 | ENSMUSG00000026548 | 102 | 15 | 214 | 66 | 105 | 27 | 174 | 36 |
| Slc7a5 | ENSMUSG00000040010 | 92 | 11 | 140 | 11 | 101 | 4 | 149 | 21 |
| Slfn2 | ENSMUSG00000072620 | 233 | 30 | 361 | 80 | 223 | 50 | 310 | 16 |
| Slfn9 | ENSMUSG00000069793 | 117 | 11 | 264 | 62 | 100 | 26 | 248 | 59 |
| Slmap | ENSMUSG00000021870 | 4454 | 592 | 5686 | 709 | 4379 | 533 | 6028 | 274 |
| Smc2 | ENSMUSG00000028312 | 165 | 38 | 318 | 105 | 129 | 27 | 242 | 61 |
| Snca | ENSMUSG00000025889 | 79 | 27 | 187 | 57 | 121 | 41 | 270 | 139 |
| Sntb2 | ENSMUSG00000041308 | 632 | 76 | 913 | 145 | 638 | 68 | 853 | 99 |
| Socs3 | ENSMUSG00000053113 | 83 | 19 | 176 | 67 | 83 | 9 | 171 | 89 |
| Sox9 | ENSMUSG00000000567 | 49 | 4 | 128 | 64 | 53 | 5 | 103 | 17 |
| Spag5 | ENSMUSG00000002055 | 26 | 31 | 54 | 13 | 19 | 10 | 55 | 17 |
| Sparc | ENSMUSG00000018593 | 7512 | 635 | 18831 | 6579 | 7355 | 933 | 14409 | 2942 |
| Spc25 | ENSMUSG00000005233 | 14 | 8 | 61 | 17 | 18 | 8 | 51 | 25 |
| Spdl1 | ENSMUSG00000069910 | 14 | 4 | 33 | 8 | 11 | 3 | 28 | 11 |
| Specc1 | ENSMUSG00000042331 | 92 | 10 | 188 | 35 | 117 | 18 | 192 | 33 |
| Sprr1a | ENSMUSG00000050359 | 2 | 1 | 59 | 85 | 1 | 1 | 31 | 38 |
| Sptlc2 | ENSMUSG00000021036 | 532 | 46 | 684 | 85 | 500 | 56 | 646 | 54 |
| Sri | ENSMUSG00000003161 | 820 | 36 | 947 | 43 | 794 | 58 | 937 | 33 |
| Srpx2 | ENSMUSG00000031253 | 164 | 22 | 353 | 215 | 151 | 23 | 300 | 80 |
| Ssc5d | ENSMUSG00000035279 | 158 | 37 | 405 | 221 | 163 | 32 | 347 | 116 |
| Stab1 | ENSMUSG00000042286 | 1603 | 325 | 2076 | 219 | 1569 | 318 | 2072 | 151 |
| Stc1 | ENSMUSG00000014813 | 72 | 10 | 132 | 17 | 71 | 21 | 124 | 24 |
| Stil | ENSMUSG00000028718 | 13 | 7 | 37 | 11 | 18 | 15 | 36 | 14 |
| Stmn1 | ENSMUSG00000028832 | 255 | 12 | 400 | 110 | 218 | 34 | 386 | 85 |
| Sulf1 | ENSMUSG00000016918 | 588 | 72 | 1266 | 375 | 636 | 106 | 916 | 142 |
| Svep1 | ENSMUSG00000028369 | 454 | 54 | 1081 | 419 | 488 | 75 | 1015 | 246 |
| Syk | ENSMUSG00000021457 | 661 | 54 | 816 | 97 | 624 | 37 | 818 | 114 |
| Synpo2l | ENSMUSG00000039376 | 2429 | 160 | 4589 | 1397 | 2488 | 266 | 4960 | 1191 |

| | | | | | | | | | |
|---|---|---|---|---|---|---|---|---|---|
| Tacc3 | ENSMUSG00000037313 | 55 | 17 | 109 | 27 | 41 | 13 | 101 | 32 |
| Tagln2 | ENSMUSG00000026547 | 1222 | 205 | 1863 | 181 | 1131 | 206 | 1710 | 250 |
| Tax1bp3 | ENSMUSG00000040158 | 636 | 52 | 758 | 26 | 603 | 49 | 731 | 37 |
| Tcf19 | ENSMUSG00000050410 | 39 | 7 | 78 | 17 | 36 | 10 | 69 | 16 |
| Tead2 | ENSMUSG00000030796 | 87 | 19 | 131 | 19 | 76 | 18 | 119 | 12 |
| Tgfb1 | ENSMUSG00000002603 | 550 | 90 | 766 | 73 | 584 | 72 | 754 | 96 |
| Tgfbi | ENSMUSG00000035493 | 530 | 77 | 732 | 148 | 489 | 68 | 651 | 78 |
| Tgif1 | ENSMUSG00000047407 | 80 | 8 | 172 | 68 | 80 | 12 | 125 | 17 |
| Thbs1 | ENSMUSG00000040152 | 409 | 149 | 2336 | 862 | 430 | 59 | 1679 | 388 |
| Thbs3 | ENSMUSG00000028047 | 103 | 23 | 225 | 128 | 92 | 40 | 196 | 95 |
| Thbs4 | ENSMUSG00000021702 | 98 | 24 | 980 | 1112 | 104 | 23 | 729 | 572 |
| Thy1 | ENSMUSG00000032011 | 161 | 30 | 284 | 38 | 162 | 24 | 265 | 26 |
| Timp1 | ENSMUSG00000001131 | 29 | 17 | 298 | 217 | 27 | 8 | 170 | 133 |
| Timp2 | ENSMUSG00000017466 | 2020 | 279 | 2851 | 462 | 2083 | 233 | 2707 | 361 |
| Tk1 | ENSMUSG00000025574 | 27 | 6 | 65 | 23 | 23 | 5 | 56 | 16 |
| Tln1 | ENSMUSG00000028465 | 3433 | 355 | 4129 | 304 | 3528 | 301 | 4361 | 274 |
| Tlr13 | ENSMUSG00000033777 | 67 | 17 | 156 | 73 | 54 | 17 | 129 | 34 |
| Tlr4 | ENSMUSG00000039005 | 491 | 150 | 724 | 86 | 471 | 25 | 747 | 193 |
| Tmem173 | ENSMUSG00000024349 | 169 | 26 | 277 | 47 | 150 | 31 | 216 | 31 |
| Tmem176b | ENSMUSG00000029810 | 447 | 59 | 695 | 169 | 422 | 18 | 612 | 87 |
| Tmem254b | ENSMUSG00000021867 | 94 | 24 | 152 | 22 | 90 | 34 | 151 | 21 |
| Tmsb10 | ENSMUSG00000079523 | 1059 | 131 | 1943 | 416 | 1045 | 141 | 1515 | 201 |
| Tmsb4x | ENSMUSG00000049775 | 6492 | 390 | 9057 | 1900 | 6337 | 792 | 8278 | 983 |
| Tnc | ENSMUSG00000028364 | 43 | 6 | 544 | 443 | 40 | 8 | 407 | 402 |
| Tnfaip6 | ENSMUSG00000053475 | 16 | 7 | 49 | 21 | 15 | 3 | 40 | 15 |
| Tnfaip8l1 | ENSMUSG00000044469 | 59 | 9 | 106 | 14 | 51 | 10 | 88 | 20 |
| Tnfrsf11b | ENSMUSG00000063727 | 4 | 4 | 30 | 9 | 8 | 3 | 25 | 12 |
| Tnfrsf12a | ENSMUSG00000023905 | 569 | 211 | 1115 | 309 | 580 | 83 | 1245 | 254 |
| Tnfrsf1a | ENSMUSG00000030341 | 862 | 76 | 1114 | 56 | 808 | 94 | 1077 | 63 |
| Top2a | ENSMUSG00000020914 | 156 | 64 | 602 | 224 | 141 | 28 | 511 | 197 |
| Tpm2 | ENSMUSG00000028464 | 338 | 60 | 565 | 58 | 358 | 58 | 477 | 79 |
| Tpm3 | ENSMUSG00000027940 | 1515 | 192 | 2133 | 211 | 1390 | 166 | 1854 | 103 |
| Tpm4 | ENSMUSG00000031799 | 3181 | 181 | 4770 | 635 | 3020 | 369 | 4360 | 322 |
| Tpx2 | ENSMUSG00000027469 | 56 | 21 | 186 | 55 | 49 | 21 | 165 | 46 |
| Trem2 | ENSMUSG00000023992 | 35 | 6 | 92 | 48 | 34 | 8 | 74 | 19 |
| Trim47 | ENSMUSG00000020773 | 322 | 56 | 432 | 50 | 294 | 39 | 391 | 53 |
| Trim59 | ENSMUSG00000034317 | 45 | 13 | 121 | 42 | 39 | 10 | 82 | 19 |
| Tspan6 | ENSMUSG00000067377 | 201 | 18 | 351 | 99 | 188 | 36 | 291 | 45 |
| Ttc9 | ENSMUSG00000042734 | 48 | 13 | 109 | 12 | 46 | 8 | 91 | 22 |
| Ttk | ENSMUSG00000038379 | 11 | 6 | 50 | 21 | 9 | 2 | 35 | 14 |
| Tuba1a | ENSMUSG00000072235 | 1839 | 223 | 2349 | 176 | 1762 | 189 | 2229 | 379 |
| Tubb5 | ENSMUSG00000001525 | 1852 | 114 | 2383 | 316 | 1680 | 188 | 2353 | 298 |
| Tyms | ENSMUSG00000025747 | 54 | 5 | 125 | 33 | 37 | 11 | 97 | 35 |
| Tyrobp | ENSMUSG00000030579 | 196 | 25 | 385 | 134 | 191 | 36 | 314 | 57 |
| Ube2c | ENSMUSG00000001403 | 26 | 8 | 84 | 26 | 25 | 10 | 76 | 27 |
| Uck2 | ENSMUSG00000026558 | 821 | 87 | 1664 | 260 | 843 | 95 | 1668 | 393 |
| Ugt1a7c | ENSMUSG00000090124 | 31 | 6 | 81 | 25 | 36 | 7 | 76 | 31 |
| Uhrf1 | ENSMUSG00000001228 | 52 | 9 | 132 | 55 | 40 | 18 | 111 | 47 |
| Ulbp1 | ENSMUSG00000079685 | 96 | 27 | 174 | 24 | 75 | 13 | 122 | 11 |
| Unc93b1 | ENSMUSG00000036908 | 322 | 73 | 550 | 131 | 355 | 31 | 500 | 38 |
| Vat1 | ENSMUSG00000034993 | 610 | 35 | 884 | 140 | 594 | 35 | 798 | 66 |
| Vcan | ENSMUSG00000021614 | 730 | 115 | 1503 | 151 | 665 | 105 | 1354 | 355 |
| Vim | ENSMUSG00000026728 | 3763 | 407 | 7380 | 1381 | 3521 | 520 | 6206 | 983 |
| Xirp2 | ENSMUSG00000027022 | 23026 | 1933 | 46990 | 12463 | 24085 | 3291 | 47862 | 16560 |
| Xylt1 | ENSMUSG00000030657 | 76 | 8 | 167 | 19 | 86 | 14 | 139 | 34 |
| Ywhaz | ENSMUSG00000022285 | 2464 | 152 | 2960 | 156 | 2391 | 141 | 2749 | 56 |
| Zyx | ENSMUSG00000029860 | 896 | 129 | 1214 | 76 | 876 | 143 | 1120 | 93 |

**Supplementary Table S5.** RNASeq analysis of effects of angiotensin II (AngII) on mRNA expression in hearts from PKN2Het vs WT littermates: mRNAs significantly downregulated by AngII in PKN2Het or WT hearts.

| Gene Symbol | Ensembl gene id | WT Vehicle | | WT AngII | | PKN2Het Vehicle | | PKN2Het AngII | |
|---|---|---|---|---|---|---|---|---|---|
| | | Mean | SD | Mean | SD | Mean | SD | Mean | SD |
| March6 | ENSMUSG00000039100 | 4345 | 470 | 3364 | 215 | 4521 | 533 | 3717 | 463 |
| A530016L24Rik | ENSMUSG00000043122 | 486 | 59 | 252 | 64 | 537 | 84 | 302 | 85 |
| Abca12 | ENSMUSG00000050296 | 186 | 25 | 118 | 22 | 190 | 14 | 113 | 26 |
| Abcc9 | ENSMUSG00000030249 | 9613 | 652 | 6487 | 953 | 9739 | 961 | 7503 | 994 |
| Acad11 | ENSMUSG00000090150 | 4332 | 376 | 3012 | 486 | 4068 | 456 | 3113 | 804 |
| Acss1 | ENSMUSG00000027452 | 6051 | 350 | 3795 | 623 | 6116 | 511 | 4694 | 536 |
| Adcy9 | ENSMUSG00000005580 | 523 | 37 | 386 | 56 | 548 | 56 | 435 | 28 |
| Adi1 | ENSMUSG00000020629 | 777 | 43 | 587 | 41 | 770 | 25 | 625 | 82 |
| Adra1a | ENSMUSG00000045875 | 454 | 44 | 309 | 38 | 449 | 28 | 324 | 43 |
| Adrb1 | ENSMUSG00000035283 | 299 | 27 | 186 | 24 | 312 | 44 | 237 | 7 |
| Aes | ENSMUSG00000054452 | 8100 | 309 | 6557 | 669 | 8573 | 728 | 7022 | 440 |
| Ak4 | ENSMUSG00000028527 | 1110 | 101 | 673 | 88 | 1095 | 120 | 841 | 121 |
| Aldh2 | ENSMUSG00000029455 | 3628 | 278 | 2621 | 303 | 3753 | 239 | 3023 | 178 |
| Aldh4a1 | ENSMUSG00000028737 | 2051 | 114 | 1277 | 213 | 2215 | 312 | 1569 | 163 |
| Aldh6a1 | ENSMUSG00000021238 | 4712 | 450 | 3354 | 316 | 4858 | 561 | 3659 | 653 |
| Aldob | ENSMUSG00000028307 | 153 | 33 | 37 | 15 | 130 | 33 | 51 | 29 |
| Angpt1 | ENSMUSG00000022309 | 983 | 254 | 585 | 130 | 825 | 181 | 510 | 175 |
| Ano10 | ENSMUSG00000037949 | 698 | 182 | 344 | 58 | 669 | 131 | 405 | 96 |
| Apbb1 | ENSMUSG00000037032 | 1485 | 207 | 973 | 270 | 1582 | 132 | 1096 | 145 |
| Arel1 | ENSMUSG00000042350 | 1385 | 91 | 997 | 117 | 1373 | 120 | 1084 | 87 |
| Arfgef1 | ENSMUSG00000067851 | 4031 | 401 | 3145 | 313 | 4160 | 428 | 3482 | 356 |
| Asb10 | ENSMUSG00000038204 | 1171 | 63 | 812 | 130 | 1209 | 144 | 936 | 72 |
| Asb14 | ENSMUSG00000021898 | 1693 | 190 | 1270 | 203 | 1859 | 69 | 1299 | 90 |
| Asb15 | ENSMUSG00000029685 | 1620 | 111 | 997 | 132 | 1719 | 167 | 1139 | 111 |
| Atp2a2 | ENSMUSG00000029467 | 276075 | 9196 | 182185 | 30616 | 276406 | 28699 | 212684 | 20219 |
| Bckdha | ENSMUSG00000060376 | 3198 | 271 | 2027 | 281 | 3491 | 476 | 2369 | 149 |
| Bckdhb | ENSMUSG00000032263 | 843 | 53 | 580 | 105 | 820 | 64 | 644 | 127 |
| Blcap | ENSMUSG00000067787 | 674 | 60 | 518 | 51 | 708 | 57 | 566 | 19 |
| Cacna1s | ENSMUSG00000026407 | 289 | 41 | 167 | 50 | 318 | 106 | 207 | 31 |
| Calcoco1 | ENSMUSG00000023055 | 2244 | 90 | 1682 | 135 | 2260 | 208 | 1798 | 84 |
| Camk2a | ENSMUSG00000024617 | 894 | 42 | 584 | 78 | 907 | 103 | 635 | 63 |
| Cbx7 | ENSMUSG00000053411 | 375 | 36 | 283 | 55 | 434 | 66 | 306 | 28 |
| Cdnf | ENSMUSG00000039496 | 693 | 83 | 462 | 69 | 638 | 86 | 470 | 25 |
| Clasp1 | ENSMUSG00000064302 | 10042 | 1348 | 7016 | 1532 | 10721 | 1184 | 8158 | 674 |
| Clasp2 | ENSMUSG00000033392 | 1555 | 68 | 1255 | 49 | 1588 | 122 | 1315 | 122 |
| Clcn1 | ENSMUSG00000029862 | 122 | 31 | 56 | 16 | 126 | 19 | 52 | 17 |
| Clpx | ENSMUSG00000015357 | 2210 | 180 | 1776 | 257 | 2247 | 186 | 1932 | 167 |
| Cmtm8 | ENSMUSG00000041012 | 193 | 16 | 122 | 26 | 198 | 19 | 133 | 20 |
| Cmya5 | ENSMUSG00000047419 | 30807 | 1640 | 21498 | 2766 | 31721 | 3064 | 25206 | 2497 |
| Cngb3 | ENSMUSG00000056494 | 85 | 21 | 34 | 5 | 96 | 10 | 48 | 18 |
| Cnst | ENSMUSG00000038949 | 1417 | 109 | 1052 | 101 | 1496 | 138 | 1149 | 127 |
| Coq8a | ENSMUSG00000026489 | 8867 | 811 | 6241 | 1021 | 9606 | 1491 | 7168 | 332 |
| Cpeb3 | ENSMUSG00000039652 | 1712 | 172 | 1204 | 231 | 1874 | 247 | 1337 | 250 |
| Creg1 | ENSMUSG00000040713 | 2146 | 83 | 1671 | 135 | 2006 | 90 | 1667 | 106 |
| Crip2 | ENSMUSG00000006356 | 12559 | 568 | 9239 | 1165 | 12739 | 1064 | 10202 | 689 |
| D10Jhu81e | ENSMUSG00000053329 | 4564 | 257 | 3218 | 428 | 4591 | 348 | 3636 | 306 |
| Dcaf11 | ENSMUSG00000022214 | 2685 | 174 | 1996 | 224 | 2643 | 170 | 2142 | 182 |
| Dcaf8 | ENSMUSG00000026554 | 3228 | 148 | 2835 | 139 | 3162 | 236 | 2954 | 198 |
| Dcun1d2 | ENSMUSG00000038506 | 1528 | 101 | 1151 | 175 | 1592 | 86 | 1245 | 86 |
| Dglucy | ENSMUSG00000021185 | 1025 | 27 | 630 | 95 | 1021 | 154 | 706 | 146 |
| Dhdh | ENSMUSG00000011382 | 539 | 37 | 412 | 40 | 541 | 58 | 441 | 60 |
| Dsg2 | ENSMUSG00000044393 | 2214 | 147 | 1452 | 293 | 2105 | 275 | 1573 | 277 |
| Ehhadh | ENSMUSG00000022853 | 319 | 54 | 215 | 36 | 367 | 52 | 263 | 31 |
| Entpd5 | ENSMUSG00000021236 | 4587 | 555 | 2853 | 380 | 4318 | 388 | 3001 | 409 |
| Epha4 | ENSMUSG00000026235 | 1430 | 183 | 778 | 139 | 1532 | 195 | 924 | 149 |

| | | | | | | | | | |
|---|---|---|---|---|---|---|---|---|---|
| Esrra | ENSMUSG00000024955 | 1498 | 69 | 1122 | 160 | 1612 | 352 | 1242 | 55 |
| Fam174b | ENSMUSG00000078670 | 5146 | 309 | 3201 | 609 | 5011 | 412 | 3808 | 373 |
| Fblim1 | ENSMUSG00000006219 | 4020 | 400 | 2963 | 478 | 4442 | 528 | 3364 | 287 |
| Fgf1 | ENSMUSG00000036585 | 4273 | 125 | 2800 | 289 | 4103 | 316 | 3148 | 422 |
| Fgf13 | ENSMUSG00000031137 | 912 | 28 | 624 | 84 | 957 | 100 | 687 | 122 |
| Fgf16 | ENSMUSG00000031230 | 350 | 35 | 197 | 36 | 325 | 28 | 212 | 47 |
| Fitm2 | ENSMUSG00000048486 | 4343 | 253 | 2687 | 652 | 4389 | 466 | 3182 | 252 |
| Fktn | ENSMUSG00000028414 | 919 | 77 | 713 | 82 | 966 | 86 | 734 | 38 |
| Fyco1 | ENSMUSG00000025241 | 8256 | 626 | 6068 | 819 | 8149 | 1328 | 6602 | 1438 |
| Gadd45a | ENSMUSG00000036390 | 237 | 30 | 173 | 8 | 280 | 38 | 199 | 29 |
| Gal3st3 | ENSMUSG00000047658 | 235 | 33 | 123 | 31 | 253 | 42 | 156 | 27 |
| Gcat | ENSMUSG00000006378 | 139 | 15 | 88 | 16 | 122 | 23 | 81 | 20 |
| Gcdh | ENSMUSG00000003809 | 1192 | 92 | 826 | 152 | 1218 | 47 | 907 | 64 |
| Ghr | ENSMUSG00000055737 | 3256 | 90 | 2495 | 135 | 3239 | 69 | 2574 | 165 |
| Gid4 | ENSMUSG00000018415 | 1429 | 66 | 1151 | 186 | 1528 | 101 | 1208 | 39 |
| Gm10435 | ENSMUSG00000072902 | 350 | 47 | 240 | 57 | 374 | 40 | 256 | 85 |
| Gm10635 | ENSMUSG00000111765 | 62 | 15 | 26 | 11 | 57 | 10 | 29 | 5 |
| Gm37691 | ENSMUSG00000104348 | 120 | 14 | 62 | 18 | 123 | 24 | 83 | 16 |
| Gpd1l | ENSMUSG00000050627 | 1532 | 131 | 1271 | 72 | 1610 | 184 | 1401 | 84 |
| Gpt2 | ENSMUSG00000031700 | 649 | 55 | 444 | 64 | 689 | 85 | 483 | 73 |
| Gramd1b | ENSMUSG00000040111 | 802 | 117 | 546 | 52 | 929 | 115 | 696 | 158 |
| Grcc10 | ENSMUSG00000072772 | 1142 | 101 | 905 | 96 | 1137 | 51 | 918 | 108 |
| Grm1 | ENSMUSG00000019828 | 796 | 113 | 605 | 58 | 918 | 161 | 761 | 121 |
| Gstm2 | ENSMUSG00000040562 | 1088 | 63 | 815 | 54 | 1016 | 120 | 792 | 38 |
| Hadha | ENSMUSG00000025745 | 29451 | 2086 | 19576 | 3952 | 28687 | 2036 | 22309 | 1220 |
| Hdac11 | ENSMUSG00000034245 | 385 | 30 | 247 | 18 | 404 | 32 | 306 | 52 |
| Hdlbp | ENSMUSG00000034088 | 16562 | 1228 | 12760 | 1097 | 16946 | 1402 | 13742 | 294 |
| Herpud1 | ENSMUSG00000031770 | 2271 | 311 | 1495 | 173 | 2164 | 218 | 1572 | 242 |
| Idh3g | ENSMUSG00000002010 | 6376 | 459 | 4785 | 580 | 6418 | 422 | 4954 | 479 |
| Ift81 | ENSMUSG00000029469 | 1002 | 129 | 666 | 78 | 994 | 49 | 708 | 78 |
| Il15 | ENSMUSG00000031712 | 439 | 60 | 265 | 38 | 422 | 29 | 277 | 62 |
| Inmt | ENSMUSG00000003477 | 120 | 30 | 62 | 9 | 125 | 27 | 60 | 31 |
| Iqsec1 | ENSMUSG00000034312 | 2343 | 237 | 1649 | 239 | 2475 | 375 | 1887 | 147 |
| Isoc1 | ENSMUSG00000024601 | 1090 | 72 | 835 | 76 | 1154 | 85 | 886 | 108 |
| Ivd | ENSMUSG00000027332 | 5467 | 219 | 3645 | 686 | 5738 | 467 | 4074 | 283 |
| Kcnd2 | ENSMUSG00000060882 | 633 | 53 | 397 | 84 | 534 | 96 | 384 | 46 |
| Kcnj11 | ENSMUSG00000096146 | 2367 | 103 | 1635 | 312 | 2506 | 312 | 1913 | 137 |
| Kcnj12 | ENSMUSG00000042529 | 362 | 32 | 215 | 47 | 369 | 42 | 263 | 36 |
| Kcnj3 | ENSMUSG00000026824 | 1558 | 124 | 892 | 227 | 1560 | 194 | 966 | 194 |
| Kcnj5 | ENSMUSG00000032034 | 1526 | 103 | 978 | 122 | 1446 | 143 | 1137 | 106 |
| Kcnv2 | ENSMUSG00000047298 | 217 | 40 | 101 | 27 | 221 | 15 | 94 | 19 |
| Klf15 | ENSMUSG00000030087 | 629 | 39 | 379 | 62 | 616 | 39 | 429 | 68 |
| Klhdc1 | ENSMUSG00000051890 | 679 | 63 | 505 | 66 | 737 | 123 | 482 | 72 |
| Klhdc7a | ENSMUSG00000078234 | 208 | 40 | 115 | 26 | 215 | 51 | 132 | 33 |
| Klhl24 | ENSMUSG00000062901 | 8344 | 685 | 6893 | 406 | 8712 | 1112 | 7374 | 755 |
| Klhl30 | ENSMUSG00000026308 | 803 | 90 | 596 | 82 | 845 | 153 | 665 | 56 |
| Klhl38 | ENSMUSG00000022357 | 594 | 49 | 396 | 86 | 598 | 80 | 398 | 64 |
| Ldhd | ENSMUSG00000031958 | 690 | 79 | 434 | 51 | 734 | 70 | 479 | 49 |
| Lgals4 | ENSMUSG00000053964 | 258 | 38 | 159 | 45 | 261 | 30 | 142 | 30 |
| Lrrc14b | ENSMUSG00000021579 | 1627 | 72 | 1181 | 214 | 1726 | 119 | 1267 | 58 |
| Lrtm1 | ENSMUSG00000045776 | 10990 | 1420 | 7948 | 2173 | 9808 | 949 | 7556 | 656 |
| Macrod1 | ENSMUSG00000036278 | 1985 | 82 | 1475 | 206 | 2115 | 228 | 1644 | 157 |
| Maob | ENSMUSG00000040147 | 1029 | 117 | 683 | 136 | 1003 | 67 | 758 | 145 |
| Mccc2 | ENSMUSG00000021646 | 1044 | 126 | 719 | 104 | 1066 | 137 | 833 | 122 |
| Me3 | ENSMUSG00000030621 | 1244 | 36 | 938 | 86 | 1271 | 81 | 1046 | 52 |
| Mfap3l | ENSMUSG00000031647 | 512 | 38 | 351 | 27 | 511 | 51 | 378 | 15 |
| Mgea5 | ENSMUSG00000025220 | 2744 | 156 | 2258 | 103 | 2825 | 202 | 2463 | 245 |
| Mitf | ENSMUSG00000035158 | 1007 | 108 | 715 | 103 | 983 | 124 | 817 | 111 |
| Mlycd | ENSMUSG00000074064 | 1449 | 99 | 1002 | 186 | 1535 | 222 | 1119 | 122 |
| Mmab | ENSMUSG00000029575 | 681 | 48 | 453 | 44 | 638 | 56 | 508 | 62 |
| Mrgprh | ENSMUSG00000059408 | 120 | 15 | 59 | 19 | 121 | 13 | 67 | 15 |
| mt-Rnr2 | ENSMUSG00000064339 | 356240 | 61989 | 247940 | 69255 | 362883 | 53161 | 276407 | 38517 |
| Mut | ENSMUSG00000023921 | 2916 | 284 | 2169 | 324 | 2826 | 245 | 2373 | 210 |
| Mylk3 | ENSMUSG00000031698 | 13367 | 1180 | 7949 | 948 | 12901 | 1032 | 9303 | 1403 |

| | | | | | | | | | |
|---|---|---|---|---|---|---|---|---|---|
| Nadk2 | ENSMUSG00000022253 | 1247 | 102 | 792 | 168 | 1254 | 222 | 893 | 160 |
| Nceh1 | ENSMUSG00000027698 | 4775 | 531 | 3631 | 517 | 4935 | 459 | 4158 | 196 |
| Nipsnap2 | ENSMUSG00000029432 | 12499 | 468 | 8522 | 1242 | 13021 | 796 | 9974 | 916 |
| Osbp2 | ENSMUSG00000020435 | 538 | 31 | 388 | 52 | 578 | 65 | 451 | 35 |
| Oxr1 | ENSMUSG00000022307 | 1785 | 149 | 1475 | 130 | 1747 | 115 | 1456 | 95 |
| Oxsm | ENSMUSG00000021786 | 592 | 47 | 470 | 26 | 633 | 71 | 520 | 20 |
| Oxsr1 | ENSMUSG00000036737 | 1469 | 64 | 1253 | 72 | 1563 | 110 | 1374 | 101 |
| P2ry1 | ENSMUSG00000027765 | 592 | 33 | 336 | 93 | 614 | 60 | 420 | 44 |
| Pank1 | ENSMUSG00000033610 | 1036 | 54 | 700 | 77 | 1098 | 87 | 753 | 63 |
| Paqr9 | ENSMUSG00000064225 | 1387 | 188 | 959 | 139 | 1468 | 178 | 1110 | 132 |
| Pcca | ENSMUSG00000041650 | 1754 | 109 | 1194 | 210 | 1808 | 136 | 1417 | 61 |
| Pcnt | ENSMUSG00000001151 | 1516 | 71 | 1137 | 205 | 1490 | 140 | 1194 | 68 |
| Pde4a | ENSMUSG00000032177 | 1479 | 69 | 1032 | 200 | 1596 | 227 | 1163 | 137 |
| Pde4d | ENSMUSG00000021699 | 566 | 55 | 373 | 47 | 575 | 76 | 416 | 43 |
| Pdha1 | ENSMUSG00000031299 | 24973 | 2883 | 19882 | 1983 | 24844 | 2183 | 21298 | 1610 |
| Pdp1 | ENSMUSG00000049225 | 1052 | 157 | 827 | 97 | 1038 | 87 | 873 | 99 |
| Pdp2 | ENSMUSG00000048371 | 822 | 182 | 433 | 87 | 839 | 116 | 560 | 111 |
| Pdpr | ENSMUSG00000033624 | 2409 | 176 | 1771 | 254 | 2523 | 330 | 2118 | 147 |
| Pex11a | ENSMUSG00000030545 | 362 | 28 | 251 | 44 | 360 | 53 | 277 | 16 |
| Pfkfb1 | ENSMUSG00000025271 | 181 | 54 | 76 | 15 | 125 | 29 | 76 | 31 |
| Pink1 | ENSMUSG00000028756 | 7908 | 460 | 5095 | 582 | 8315 | 930 | 5887 | 645 |
| Pkia | ENSMUSG00000027499 | 6489 | 587 | 5105 | 667 | 6499 | 450 | 5564 | 198 |
| Pkig | ENSMUSG00000035268 | 1911 | 58 | 1637 | 67 | 1943 | 106 | 1679 | 127 |
| Pkm | ENSMUSG00000032294 | 19295 | 1479 | 15108 | 1318 | 19558 | 1824 | 16780 | 652 |
| Pla2g5 | ENSMUSG00000041193 | 703 | 60 | 323 | 118 | 617 | 100 | 370 | 76 |
| Pln | ENSMUSG00000038583 | 103669 | 9371 | 75161 | 8133 | 106163 | 7666 | 81558 | 8775 |
| Plxnb1 | ENSMUSG00000053646 | 817 | 113 | 486 | 68 | 762 | 119 | 534 | 62 |
| Pm20d2 | ENSMUSG00000054659 | 458 | 21 | 337 | 55 | 500 | 64 | 387 | 32 |
| Pnpla8 | ENSMUSG00000036257 | 3505 | 336 | 2968 | 271 | 3603 | 216 | 3140 | 212 |
| Ppargc1a | ENSMUSG00000029167 | 1975 | 310 | 1550 | 238 | 2310 | 255 | 1647 | 126 |
| Ppfibp2 | ENSMUSG00000036528 | 632 | 102 | 494 | 48 | 689 | 69 | 541 | 29 |
| Ppip5k2 | ENSMUSG00000040648 | 2823 | 288 | 1947 | 183 | 2670 | 258 | 1967 | 582 |
| Ppm1l | ENSMUSG00000027784 | 1625 | 159 | 1031 | 184 | 1684 | 206 | 1260 | 149 |
| Ppp1r14c | ENSMUSG00000040653 | 3104 | 380 | 2506 | 85 | 3332 | 336 | 2807 | 364 |
| Pptc7 | ENSMUSG00000038582 | 3687 | 299 | 2559 | 451 | 3939 | 504 | 3081 | 324 |
| Prkab1 | ENSMUSG00000029513 | 625 | 39 | 486 | 57 | 684 | 60 | 543 | 69 |
| Rap1gap2 | ENSMUSG00000038807 | 2208 | 274 | 1592 | 353 | 2292 | 294 | 1806 | 108 |
| Rbfox1 | ENSMUSG00000008658 | 634 | 47 | 360 | 59 | 657 | 70 | 426 | 78 |
| Reep1 | ENSMUSG00000052852 | 440 | 28 | 345 | 23 | 454 | 53 | 354 | 26 |
| Reep5 | ENSMUSG00000005873 | 6212 | 260 | 4792 | 333 | 6407 | 427 | 5212 | 316 |
| Rgs2 | ENSMUSG00000026360 | 831 | 133 | 501 | 64 | 860 | 81 | 573 | 166 |
| Ric8b | ENSMUSG00000035620 | 1056 | 84 | 764 | 88 | 1036 | 119 | 844 | 32 |
| Rilpl1 | ENSMUSG00000029392 | 2712 | 176 | 2148 | 322 | 2829 | 181 | 2289 | 127 |
| Rmnd5a | ENSMUSG00000002222 | 2488 | 67 | 2108 | 149 | 2641 | 123 | 2117 | 196 |
| Rpl3l | ENSMUSG00000002500 | 3330 | 271 | 1888 | 478 | 3250 | 187 | 2275 | 264 |
| Rtn2 | ENSMUSG00000030401 | 979 | 45 | 649 | 92 | 997 | 112 | 733 | 89 |
| Sdha | ENSMUSG00000021577 | 26565 | 1713 | 17836 | 2991 | 26775 | 2129 | 20539 | 1624 |
| Sec31b | ENSMUSG00000051984 | 190 | 12 | 140 | 15 | 206 | 17 | 142 | 18 |
| Selenbp1 | ENSMUSG00000068874 | 1432 | 58 | 903 | 170 | 1401 | 157 | 1028 | 139 |
| Sgcb | ENSMUSG00000029156 | 2725 | 146 | 2351 | 41 | 2817 | 190 | 2512 | 150 |
| Slc20a2 | ENSMUSG00000037656 | 3299 | 120 | 2345 | 335 | 3156 | 313 | 2538 | 136 |
| Slc22a5 | ENSMUSG00000018900 | 515 | 48 | 392 | 19 | 513 | 45 | 415 | 24 |
| Slc25a34 | ENSMUSG00000040740 | 2458 | 124 | 1716 | 276 | 2459 | 202 | 1859 | 71 |
| Slc25a42 | ENSMUSG00000002346 | 919 | 100 | 557 | 97 | 966 | 156 | 583 | 82 |
| Slc27a1 | ENSMUSG00000031808 | 1220 | 145 | 877 | 126 | 1202 | 129 | 879 | 108 |
| Slc4a3 | ENSMUSG00000006576 | 2856 | 245 | 2018 | 244 | 3026 | 328 | 2319 | 297 |
| Smim20 | ENSMUSG00000061461 | 843 | 73 | 641 | 62 | 853 | 63 | 664 | 59 |
| Stom | ENSMUSG00000026880 | 2813 | 112 | 2237 | 204 | 2994 | 287 | 2470 | 159 |
| Stum | ENSMUSG00000053963 | 160 | 19 | 64 | 17 | 122 | 38 | 54 | 18 |
| Syde2 | ENSMUSG00000036863 | 475 | 95 | 306 | 46 | 437 | 39 | 305 | 68 |
| Synj2 | ENSMUSG00000023805 | 1330 | 103 | 1004 | 60 | 1390 | 140 | 1030 | 43 |
| Taf1a | ENSMUSG00000072258 | 316 | 29 | 238 | 17 | 323 | 11 | 239 | 23 |
| Tbc1d10c | ENSMUSG00000040247 | 79 | 13 | 45 | 9 | 80 | 16 | 46 | 8 |
| Tbc1d16 | ENSMUSG00000039976 | 1680 | 162 | 1163 | 183 | 1773 | 217 | 1329 | 136 |

| | | | | | | | | | |
|---|---|---|---|---|---|---|---|---|---|
| Tbc1d4 | ENSMUSG00000033083 | 2228 | 153 | 1576 | 167 | 2172 | 256 | 1634 | 160 |
| Tbx5 | ENSMUSG00000018263 | 456 | 56 | 299 | 60 | 527 | 87 | 347 | 57 |
| Tcea3 | ENSMUSG00000001604 | 1429 | 35 | 1076 | 111 | 1549 | 116 | 1154 | 106 |
| Tmem150c | ENSMUSG00000050640 | 201 | 25 | 109 | 24 | 249 | 43 | 146 | 26 |
| Tmem182 | ENSMUSG00000079588 | 4532 | 444 | 3422 | 366 | 4747 | 305 | 3856 | 392 |
| Tmem245 | ENSMUSG00000055296 | 3203 | 419 | 2375 | 169 | 3028 | 247 | 2485 | 165 |
| Tmem63b | ENSMUSG00000036026 | 1741 | 102 | 1317 | 164 | 1858 | 169 | 1449 | 124 |
| Tmem65 | ENSMUSG00000062373 | 3732 | 474 | 2882 | 185 | 3814 | 575 | 3099 | 401 |
| Tnfrsf19 | ENSMUSG00000060548 | 97 | 27 | 54 | 15 | 91 | 22 | 56 | 12 |
| Tnni3k | ENSMUSG00000040086 | 2347 | 262 | 1468 | 283 | 2247 | 145 | 1743 | 187 |
| Trap1 | ENSMUSG00000005981 | 2287 | 176 | 1864 | 73 | 2431 | 195 | 1969 | 70 |
| Trim7 | ENSMUSG00000040350 | 466 | 53 | 297 | 42 | 440 | 107 | 331 | 53 |
| Trip10 | ENSMUSG00000019487 | 1361 | 60 | 1155 | 94 | 1369 | 30 | 1142 | 93 |
| Ttll1 | ENSMUSG00000022442 | 921 | 104 | 531 | 110 | 965 | 54 | 680 | 86 |
| Txlnb | ENSMUSG00000039891 | 15534 | 643 | 11828 | 883 | 16172 | 1476 | 12752 | 915 |
| Uckl1os | ENSMUSG00000010492 | 61 | 16 | 21 | 7 | 56 | 24 | 31 | 10 |
| Vldlr | ENSMUSG00000024924 | 9623 | 566 | 7598 | 786 | 10008 | 777 | 8275 | 415 |
| Ybx2 | ENSMUSG00000018554 | 158 | 19 | 86 | 33 | 158 | 21 | 98 | 24 |
| Zfp612 | ENSMUSG00000044676 | 301 | 27 | 215 | 18 | 304 | 9 | 201 | 39 |
| Zyg11b | ENSMUSG00000034636 | 2665 | 223 | 2061 | 190 | 2682 | 291 | 2292 | 252 |

**Supplementary Table S6.** RNASeq analysis of effects of angiotensin II (AngII) on mRNA expression in hearts from PKN2Het vs WT littermates: mRNAs significantly upregulated by AngII in WT hearts.

| **Gene Symbol** | **Ensembl gene id** | **WT Vehicle** | | **WT AngII** | | **PKN2Het Vehicle** | | **PKN2Het AngII** | |
|---|---|---|---|---|---|---|---|---|---|
| | | **Mean** | **SD** | **Mean** | **SD** | **Mean** | **SD** | **Mean** | **SD** |
| March1 | ENSMUSG00000036469 | 56 | 11 | 114 | 66 | 64 | 15 | 90 | 21 |
| 1500011B03Rik | ENSMUSG00000072694 | 30 | 9 | 54 | 13 | 33 | 8 | 48 | 9 |
| 1500015O10Rik | ENSMUSG00000026051 | 3 | 2 | 49 | 80 | 5 | 3 | 18 | 14 |
| 4930503L19Rik | ENSMUSG00000044906 | 133 | 18 | 213 | 40 | 136 | 22 | 156 | 26 |
| 9930111J21Rik2 | ENSMUSG00000069892 | 725 | 103 | 926 | 89 | 721 | 107 | 783 | 91 |
| Abca9 | ENSMUSG00000041797 | 849 | 55 | 1144 | 187 | 818 | 92 | 953 | 67 |
| Abhd2 | ENSMUSG00000039202 | 898 | 103 | 1060 | 61 | 933 | 139 | 1000 | 123 |
| AC105304.1 | ENSMUSG00000117110 | 1 | 2 | 19 | 19 | 2 | 2 | 11 | 8 |
| Acp5 | ENSMUSG00000001348 | 5 | 2 | 22 | 17 | 6 | 4 | 15 | 9 |
| Actb | ENSMUSG00000029580 | 7471 | 1804 | 10178 | 1677 | 7763 | 1384 | 9365 | 1200 |
| Actg1 | ENSMUSG00000062825 | 6824 | 1047 | 8916 | 1194 | 7413 | 1015 | 8405 | 1266 |
| Actg2 | ENSMUSG00000059430 | 11 | 6 | 37 | 14 | 12 | 6 | 20 | 12 |
| Actn4 | ENSMUSG00000054808 | 2213 | 272 | 2742 | 211 | 2012 | 203 | 2417 | 171 |
| Actr2 | ENSMUSG00000020152 | 2099 | 200 | 2493 | 267 | 2101 | 71 | 2352 | 201 |
| Adam10 | ENSMUSG00000054693 | 1295 | 82 | 1540 | 84 | 1289 | 31 | 1416 | 75 |
| Adamts4 | ENSMUSG00000006403 | 26 | 22 | 120 | 29 | 38 | 22 | 83 | 38 |
| Adgra2 | ENSMUSG00000031486 | 286 | 38 | 404 | 66 | 291 | 29 | 400 | 70 |
| Adss | ENSMUSG00000015961 | 296 | 25 | 391 | 63 | 284 | 33 | 321 | 15 |
| Aebp1 | ENSMUSG00000020473 | 297 | 22 | 678 | 352 | 334 | 100 | 516 | 123 |
| Agrn | ENSMUSG00000041936 | 958 | 125 | 1295 | 154 | 947 | 79 | 1136 | 70 |
| Ahnak2 | ENSMUSG00000072812 | 300 | 55 | 611 | 330 | 327 | 47 | 473 | 130 |
| Aida | ENSMUSG00000042901 | 767 | 113 | 1024 | 80 | 877 | 107 | 902 | 132 |
| Akr1b8 | ENSMUSG00000029762 | 67 | 13 | 106 | 31 | 68 | 10 | 93 | 26 |
| Akt3 | ENSMUSG00000019699 | 651 | 50 | 856 | 71 | 640 | 112 | 760 | 47 |
| Aldh1a3 | ENSMUSG00000015134 | 25 | 2 | 51 | 12 | 25 | 7 | 49 | 8 |
| Amot | ENSMUSG00000041688 | 452 | 43 | 612 | 111 | 514 | 116 | 652 | 97 |
| Antxr1 | ENSMUSG00000033420 | 343 | 48 | 747 | 479 | 332 | 36 | 524 | 184 |
| Anxa3 | ENSMUSG00000029484 | 637 | 92 | 881 | 124 | 558 | 119 | 714 | 80 |
| Anxa4 | ENSMUSG00000029994 | 390 | 35 | 552 | 99 | 380 | 26 | 495 | 39 |
| Anxa8 | ENSMUSG00000021950 | 10 | 5 | 45 | 20 | 13 | 5 | 25 | 12 |
| Aoah | ENSMUSG00000021322 | 51 | 20 | 105 | 50 | 48 | 14 | 83 | 23 |
| Ap2b1 | ENSMUSG00000035152 | 1179 | 64 | 1470 | 164 | 1223 | 59 | 1427 | 130 |
| Ap3s1 | ENSMUSG00000024480 | 226 | 22 | 364 | 126 | 251 | 27 | 316 | 37 |
| Apaf1 | ENSMUSG00000019979 | 202 | 42 | 332 | 75 | 213 | 32 | 293 | 28 |
| Apobec1 | ENSMUSG00000040613 | 119 | 28 | 241 | 144 | 120 | 8 | 163 | 31 |
| Apobr | ENSMUSG00000042759 | 32 | 3 | 70 | 17 | 32 | 9 | 51 | 18 |
| Apol11b | ENSMUSG00000091694 | 7 | 5 | 34 | 9 | 17 | 6 | 37 | 25 |
| App | ENSMUSG00000022892 | 3981 | 381 | 4939 | 605 | 3820 | 129 | 4513 | 126 |
| Aqp8 | ENSMUSG00000030762 | 83 | 23 | 156 | 68 | 106 | 26 | 182 | 68 |
| Arf3 | ENSMUSG00000051853 | 700 | 31 | 853 | 47 | 714 | 16 | 777 | 34 |
| Arfip1 | ENSMUSG00000074513 | 425 | 63 | 585 | 108 | 425 | 15 | 463 | 26 |
| Arhgap23 | ENSMUSG00000049807 | 496 | 48 | 632 | 37 | 504 | 37 | 586 | 47 |
| Arhgap45 | ENSMUSG00000035697 | 124 | 18 | 203 | 68 | 131 | 23 | 172 | 35 |
| Arhgdia | ENSMUSG00000025132 | 2274 | 183 | 2741 | 90 | 2286 | 149 | 2537 | 205 |
| Arhgef2 | ENSMUSG00000028059 | 885 | 125 | 1137 | 110 | 951 | 83 | 1051 | 107 |
| Arhgef39 | ENSMUSG00000051517 | 7 | 3 | 25 | 5 | 7 | 4 | 18 | 6 |
| Armcx2 | ENSMUSG00000033436 | 204 | 22 | 300 | 52 | 209 | 17 | 270 | 18 |
| Arpc2 | ENSMUSG00000006304 | 2337 | 43 | 2757 | 185 | 2338 | 47 | 2652 | 121 |
| Arpin | ENSMUSG00000039043 | 239 | 19 | 299 | 19 | 258 | 25 | 280 | 26 |
| Arrb2 | ENSMUSG00000060216 | 182 | 31 | 313 | 100 | 186 | 36 | 253 | 55 |
| Arsb | ENSMUSG00000042082 | 304 | 35 | 427 | 70 | 298 | 13 | 358 | 32 |
| Aspn | ENSMUSG00000021388 | 750 | 159 | 3970 | 5012 | 798 | 171 | 1825 | 1043 |
| Ass1 | ENSMUSG00000076441 | 79 | 24 | 147 | 71 | 80 | 5 | 118 | 24 |
| Atl3 | ENSMUSG00000024759 | 741 | 53 | 1024 | 135 | 748 | 50 | 882 | 82 |
| Atp6v0a4 | ENSMUSG00000038600 | 9 | 7 | 42 | 23 | 11 | 7 | 25 | 11 |

| | | | | | | | | | |
|---|---|---|---|---|---|---|---|---|---|
| Atp6v1h | ENSMUSG00000033793 | 486 | 36 | 596 | 56 | 470 | 28 | 561 | 47 |
| Atp7a | ENSMUSG00000033792 | 198 | 11 | 253 | 13 | 197 | 21 | 209 | 17 |
| Atp9a | ENSMUSG00000027546 | 1025 | 69 | 1233 | 124 | 1046 | 42 | 1120 | 130 |
| AW551984 | ENSMUSG00000038112 | 5 | 2 | 17 | 8 | 7 | 4 | 17 | 10 |
| B2m | ENSMUSG00000060802 | 4421 | 1037 | 6550 | 1100 | 4422 | 785 | 4929 | 246 |
| B3galt2 | ENSMUSG00000033849 | 372 | 49 | 542 | 57 | 383 | 37 | 471 | 157 |
| B4galnt1 | ENSMUSG00000006731 | 31 | 7 | 66 | 39 | 32 | 6 | 48 | 11 |
| B4galt1 | ENSMUSG00000028413 | 1799 | 176 | 2095 | 181 | 1836 | 106 | 2087 | 131 |
| Baalc | ENSMUSG00000022296 | 20 | 5 | 38 | 10 | 18 | 4 | 29 | 7 |
| Bax | ENSMUSG00000003873 | 210 | 31 | 278 | 16 | 218 | 17 | 271 | 24 |
| BC037034 | ENSMUSG00000036948 | 67 | 7 | 102 | 13 | 66 | 9 | 74 | 11 |
| Bcl10 | ENSMUSG00000028191 | 379 | 38 | 473 | 26 | 405 | 34 | 468 | 34 |
| Bcl2 | ENSMUSG00000057329 | 294 | 11 | 378 | 46 | 307 | 30 | 367 | 43 |
| Bcl2a1b | ENSMUSG00000089929 | 18 | 7 | 68 | 61 | 17 | 5 | 44 | 19 |
| Bcl3 | ENSMUSG00000053175 | 60 | 35 | 113 | 24 | 70 | 15 | 98 | 24 |
| Bcl6b | ENSMUSG00000000317 | 820 | 250 | 1194 | 317 | 923 | 223 | 1077 | 280 |
| Bicc1 | ENSMUSG00000014329 | 681 | 54 | 1034 | 279 | 743 | 66 | 924 | 94 |
| Bin1 | ENSMUSG00000024381 | 163 | 26 | 241 | 35 | 165 | 22 | 215 | 15 |
| Bmp2k | ENSMUSG00000034663 | 222 | 15 | 328 | 62 | 234 | 20 | 306 | 14 |
| Bora | ENSMUSG00000022070 | 17 | 4 | 37 | 7 | 18 | 8 | 30 | 4 |
| C1qtnf3 | ENSMUSG00000058914 | 3 | 2 | 341 | 632 | 3 | 2 | 118 | 157 |
| C1qtnf5 | ENSMUSG00000079592 | 53 | 6 | 100 | 38 | 61 | 17 | 87 | 20 |
| C1qtnf7 | ENSMUSG00000061535 | 178 | 26 | 294 | 78 | 174 | 31 | 227 | 32 |
| C3ar1 | ENSMUSG00000040552 | 234 | 31 | 502 | 241 | 242 | 41 | 380 | 97 |
| Cacnb3 | ENSMUSG00000003352 | 53 | 8 | 108 | 38 | 53 | 6 | 75 | 17 |
| Calhm5 | ENSMUSG00000049872 | 49 | 5 | 97 | 18 | 60 | 11 | 89 | 26 |
| Calm2 | ENSMUSG00000036438 | 2322 | 263 | 2921 | 526 | 2281 | 193 | 2735 | 103 |
| Camk1d | ENSMUSG00000039145 | 125 | 28 | 197 | 13 | 146 | 26 | 154 | 32 |
| Camkk1 | ENSMUSG00000020785 | 29 | 5 | 54 | 12 | 30 | 5 | 42 | 7 |
| Capn6 | ENSMUSG00000067276 | 8 | 3 | 42 | 52 | 12 | 8 | 24 | 13 |
| Casp12 | ENSMUSG00000025887 | 208 | 53 | 318 | 96 | 186 | 49 | 260 | 34 |
| Casp3 | ENSMUSG00000031628 | 113 | 29 | 191 | 45 | 108 | 17 | 171 | 41 |
| Cbfb | ENSMUSG00000031885 | 509 | 26 | 664 | 130 | 518 | 37 | 561 | 32 |
| Ccdc88a | ENSMUSG00000032740 | 355 | 54 | 511 | 63 | 310 | 60 | 391 | 78 |
| Ccdc88b | ENSMUSG00000047810 | 26 | 4 | 52 | 17 | 31 | 7 | 43 | 4 |
| Ccl12 | ENSMUSG00000035352 | 28 | 12 | 76 | 30 | 20 | 5 | 61 | 36 |
| Ccr2 | ENSMUSG00000049103 | 85 | 30 | 408 | 389 | 93 | 25 | 233 | 135 |
| Ccr5 | ENSMUSG00000079227 | 186 | 40 | 341 | 137 | 159 | 68 | 244 | 42 |
| Cd180 | ENSMUSG00000021624 | 33 | 12 | 77 | 31 | 53 | 18 | 71 | 11 |
| Cd24a | ENSMUSG00000047139 | 79 | 19 | 145 | 26 | 90 | 14 | 161 | 75 |
| Cd2ap | ENSMUSG00000061665 | 690 | 32 | 831 | 97 | 667 | 57 | 750 | 87 |
| Cd33 | ENSMUSG00000004609 | 150 | 35 | 242 | 42 | 149 | 35 | 189 | 29 |
| Cd52 | ENSMUSG00000000682 | 61 | 22 | 171 | 113 | 54 | 14 | 95 | 23 |
| Cd53 | ENSMUSG00000040747 | 129 | 18 | 286 | 154 | 144 | 34 | 221 | 59 |
| Cd55 | ENSMUSG00000026399 | 345 | 57 | 518 | 88 | 369 | 49 | 468 | 57 |
| Cd63 | ENSMUSG00000025351 | 1383 | 97 | 1809 | 371 | 1430 | 52 | 1693 | 169 |
| Cd80 | ENSMUSG00000075122 | 19 | 7 | 60 | 17 | 24 | 9 | 47 | 22 |
| Cd83 | ENSMUSG00000015396 | 132 | 30 | 202 | 45 | 132 | 31 | 168 | 13 |
| Cd86 | ENSMUSG00000022901 | 75 | 19 | 139 | 14 | 88 | 32 | 109 | 23 |
| Cdc25b | ENSMUSG00000027330 | 92 | 22 | 163 | 19 | 92 | 22 | 150 | 40 |
| Cdc42se1 | ENSMUSG00000046722 | 468 | 71 | 617 | 41 | 442 | 53 | 554 | 68 |
| Cdc6 | ENSMUSG00000017499 | 18 | 6 | 40 | 11 | 16 | 6 | 33 | 8 |
| Cdc7 | ENSMUSG00000029283 | 20 | 3 | 41 | 10 | 29 | 11 | 30 | 8 |
| Cdca2 | ENSMUSG00000048922 | 13 | 3 | 38 | 8 | 27 | 15 | 36 | 10 |
| Cdca4 | ENSMUSG00000047832 | 122 | 18 | 180 | 26 | 120 | 24 | 154 | 24 |
| Cdk14 | ENSMUSG00000028926 | 200 | 28 | 273 | 53 | 192 | 35 | 226 | 37 |
| Cdkn3 | ENSMUSG00000037628 | 13 | 6 | 27 | 8 | 12 | 2 | 21 | 5 |
| Cdr2 | ENSMUSG00000030878 | 234 | 28 | 362 | 52 | 247 | 34 | 334 | 25 |
| Cdr2l | ENSMUSG00000050910 | 97 | 21 | 164 | 22 | 96 | 12 | 136 | 9 |
| Cdt1 | ENSMUSG00000006585 | 24 | 1 | 51 | 16 | 26 | 8 | 47 | 12 |
| Cemip | ENSMUSG00000052353 | 1 | 1 | 29 | 37 | 0 | 0 | 17 | 18 |
| Cenpn | ENSMUSG00000031756 | 14 | 5 | 38 | 13 | 16 | 3 | 27 | 10 |
| Cep192 | ENSMUSG00000024542 | 181 | 30 | 290 | 42 | 205 | 41 | 226 | 32 |
| Cercam | ENSMUSG00000039787 | 51 | 4 | 142 | 102 | 51 | 14 | 90 | 27 |

| | | | | | | | | | |
|---|---|---|---|---|---|---|---|---|---|
| Cggbp1 | ENSMUSG00000054604 | 811 | 96 | 987 | 62 | 796 | 114 | 900 | 95 |
| Chaf1a | ENSMUSG00000002835 | 62 | 28 | 147 | 37 | 69 | 14 | 117 | 37 |
| Chaf1b | ENSMUSG00000022945 | 12 | 2 | 37 | 18 | 18 | 4 | 36 | 9 |
| Chst2 | ENSMUSG00000033350 | 33 | 6 | 59 | 8 | 44 | 9 | 45 | 10 |
| Chsy3 | ENSMUSG00000058152 | 5 | 1 | 14 | 5 | 5 | 2 | 9 | 3 |
| Cip2a | ENSMUSG00000033031 | 56 | 15 | 113 | 30 | 63 | 19 | 81 | 15 |
| Cks1b | ENSMUSG00000028044 | 53 | 13 | 94 | 14 | 55 | 9 | 83 | 13 |
| Clca3a1 | ENSMUSG00000056025 | 11 | 2 | 33 | 21 | 15 | 8 | 17 | 9 |
| Cldn15 | ENSMUSG00000001739 | 49 | 9 | 95 | 9 | 66 | 28 | 67 | 17 |
| Clec11a | ENSMUSG00000004473 | 16 | 2 | 53 | 48 | 14 | 4 | 31 | 14 |
| Clec12a | ENSMUSG00000053063 | 76 | 11 | 211 | 106 | 102 | 28 | 134 | 18 |
| Clec4a1 | ENSMUSG00000049037 | 90 | 14 | 212 | 105 | 96 | 15 | 147 | 36 |
| Clec4a2 | ENSMUSG00000030148 | 46 | 12 | 104 | 63 | 40 | 6 | 79 | 23 |
| Clec4a3 | ENSMUSG00000043832 | 54 | 12 | 126 | 46 | 52 | 9 | 86 | 21 |
| Clspn | ENSMUSG00000042489 | 17 | 7 | 42 | 12 | 17 | 3 | 38 | 23 |
| Cnn1 | ENSMUSG00000001349 | 37 | 23 | 130 | 75 | 42 | 17 | 53 | 24 |
| Cnn2 | ENSMUSG00000004665 | 793 | 93 | 1039 | 52 | 856 | 80 | 1002 | 98 |
| Cnrip1 | ENSMUSG00000044629 | 67 | 5 | 110 | 32 | 77 | 10 | 96 | 9 |
| Cntln | ENSMUSG00000038070 | 189 | 18 | 279 | 45 | 181 | 18 | 235 | 37 |
| Cntrl | ENSMUSG00000057110 | 277 | 35 | 379 | 18 | 284 | 40 | 270 | 21 |
| Col11a1 | ENSMUSG00000027966 | 2 | 3 | 52 | 76 | 2 | 2 | 12 | 11 |
| Col16a1 | ENSMUSG00000040690 | 187 | 39 | 603 | 471 | 192 | 60 | 420 | 186 |
| Col1a1 | ENSMUSG00000001506 | 1941 | 334 | 10243 | 9888 | 2115 | 63 | 5954 | 3420 |
| Col1a2 | ENSMUSG00000029661 | 2657 | 250 | 11812 | 10867 | 2786 | 231 | 7551 | 4300 |
| Col4a5 | ENSMUSG00000031274 | 763 | 91 | 1159 | 304 | 773 | 91 | 971 | 147 |
| Col5a3 | ENSMUSG00000004098 | 576 | 178 | 985 | 63 | 666 | 103 | 847 | 222 |
| Col7a1 | ENSMUSG00000025650 | 4 | 6 | 26 | 21 | 5 | 4 | 11 | 6 |
| Col8a2 | ENSMUSG00000056174 | 9 | 5 | 180 | 298 | 14 | 4 | 86 | 85 |
| Col9a2 | ENSMUSG00000028626 | 6 | 1 | 31 | 38 | 7 | 4 | 21 | 11 |
| Comp | ENSMUSG00000031849 | 39 | 8 | 325 | 426 | 69 | 9 | 179 | 129 |
| Copb1 | ENSMUSG00000030754 | 983 | 86 | 1212 | 193 | 1045 | 57 | 1076 | 62 |
| Coro1a | ENSMUSG00000030707 | 175 | 29 | 270 | 73 | 171 | 24 | 228 | 26 |
| Coro1b | ENSMUSG00000024835 | 688 | 47 | 858 | 53 | 695 | 47 | 802 | 76 |
| Cplx2 | ENSMUSG00000025867 | 149 | 21 | 210 | 31 | 160 | 16 | 218 | 24 |
| Cpne8 | ENSMUSG00000052560 | 161 | 16 | 246 | 70 | 161 | 15 | 217 | 22 |
| Creb3l2 | ENSMUSG00000038648 | 1002 | 88 | 1355 | 234 | 1018 | 47 | 1231 | 121 |
| Crtap | ENSMUSG00000032431 | 384 | 57 | 511 | 72 | 379 | 32 | 471 | 67 |
| Csf2ra | ENSMUSG00000059326 | 56 | 11 | 101 | 28 | 65 | 12 | 82 | 17 |
| Csrp1 | ENSMUSG00000026421 | 868 | 80 | 1281 | 109 | 889 | 121 | 1075 | 145 |
| Cstb | ENSMUSG00000005054 | 171 | 14 | 238 | 43 | 175 | 15 | 218 | 32 |
| Cthrc1 | ENSMUSG00000054196 | 3 | 3 | 221 | 380 | 5 | 2 | 92 | 112 |
| Ctsk | ENSMUSG00000028111 | 95 | 23 | 247 | 204 | 90 | 21 | 156 | 59 |
| Ctss | ENSMUSG00000038642 | 458 | 93 | 1279 | 617 | 459 | 103 | 843 | 274 |
| Cttn | ENSMUSG00000031078 | 793 | 94 | 1056 | 113 | 811 | 56 | 962 | 95 |
| Cttnbp2nl | ENSMUSG00000062127 | 644 | 157 | 873 | 93 | 653 | 151 | 770 | 81 |
| Cx3cr1 | ENSMUSG00000052336 | 153 | 23 | 369 | 148 | 170 | 25 | 253 | 86 |
| Cxcl10 | ENSMUSG00000034855 | 12 | 4 | 57 | 37 | 12 | 8 | 27 | 6 |
| Cybb | ENSMUSG00000015340 | 327 | 42 | 648 | 221 | 360 | 106 | 406 | 69 |
| Cysltr1 | ENSMUSG00000052821 | 54 | 23 | 122 | 68 | 67 | 22 | 119 | 47 |
| Cyth3 | ENSMUSG00000018001 | 939 | 142 | 1235 | 98 | 1006 | 115 | 1159 | 89 |
| Cyth4 | ENSMUSG00000018008 | 207 | 45 | 374 | 116 | 216 | 29 | 315 | 54 |
| D1Ertd622e | ENSMUSG00000044768 | 81 | 14 | 134 | 22 | 88 | 9 | 109 | 16 |
| Dap | ENSMUSG00000039168 | 262 | 38 | 433 | 140 | 293 | 35 | 371 | 55 |
| Dbf4 | ENSMUSG00000002297 | 39 | 11 | 81 | 24 | 39 | 8 | 63 | 4 |
| Dbnl | ENSMUSG00000020476 | 472 | 46 | 575 | 30 | 501 | 31 | 573 | 37 |
| Dck | ENSMUSG00000029366 | 104 | 18 | 188 | 15 | 109 | 4 | 138 | 16 |
| Dhx58 | ENSMUSG00000017830 | 37 | 15 | 76 | 24 | 54 | 15 | 63 | 11 |
| Dkk3 | ENSMUSG00000030772 | 52 | 16 | 247 | 258 | 56 | 20 | 169 | 78 |
| Dnm1 | ENSMUSG00000026825 | 162 | 38 | 244 | 43 | 172 | 24 | 214 | 33 |
| Dock11 | ENSMUSG00000031093 | 198 | 35 | 312 | 74 | 204 | 30 | 252 | 29 |
| Dock7 | ENSMUSG00000028556 | 383 | 56 | 520 | 72 | 415 | 36 | 422 | 61 |
| Dok3 | ENSMUSG00000035711 | 38 | 8 | 70 | 27 | 33 | 8 | 52 | 10 |
| Dpep2 | ENSMUSG00000053687 | 21 | 8 | 44 | 15 | 16 | 3 | 33 | 12 |
| Dpp7 | ENSMUSG00000026958 | 48 | 11 | 85 | 22 | 56 | 8 | 80 | 17 |

| | | | | | | | | | |
|---|---|---|---|---|---|---|---|---|---|
| Dpy19l1 | ENSMUSG00000043067 | 214 | 23 | 340 | 91 | 229 | 13 | 276 | 30 |
| Dram1 | ENSMUSG00000020057 | 109 | 11 | 158 | 29 | 104 | 11 | 139 | 11 |
| Dse | ENSMUSG00000039497 | 182 | 24 | 294 | 74 | 230 | 55 | 247 | 27 |
| Dsel | ENSMUSG00000038702 | 152 | 30 | 302 | 163 | 181 | 17 | 233 | 34 |
| E2f2 | ENSMUSG00000018983 | 15 | 9 | 47 | 2 | 35 | 28 | 45 | 12 |
| E2f3 | ENSMUSG00000016477 | 259 | 22 | 315 | 27 | 280 | 21 | 306 | 28 |
| E2f8 | ENSMUSG00000046179 | 31 | 20 | 49 | 20 | 32 | 12 | 50 | 22 |
| Egr2 | ENSMUSG00000037868 | 32 | 17 | 104 | 54 | 67 | 11 | 86 | 35 |
| Egr3 | ENSMUSG00000033730 | 55 | 46 | 127 | 33 | 126 | 43 | 192 | 110 |
| Eif2ak2 | ENSMUSG00000024079 | 457 | 54 | 612 | 78 | 452 | 48 | 564 | 48 |
| Eif4ebp1 | ENSMUSG00000031490 | 600 | 42 | 768 | 49 | 633 | 27 | 785 | 117 |
| Elf1 | ENSMUSG00000036461 | 427 | 66 | 547 | 56 | 479 | 70 | 499 | 46 |
| Eln | ENSMUSG00000029675 | 512 | 176 | 2249 | 1534 | 591 | 55 | 1327 | 606 |
| Enpp1 | ENSMUSG00000037370 | 187 | 9 | 495 | 317 | 181 | 24 | 350 | 178 |
| Epb41l2 | ENSMUSG00000019978 | 1216 | 183 | 1533 | 194 | 1128 | 76 | 1296 | 65 |
| Epsti1 | ENSMUSG00000022014 | 40 | 10 | 90 | 32 | 45 | 8 | 66 | 11 |
| Ercc6l | ENSMUSG00000051220 | 19 | 8 | 48 | 17 | 17 | 6 | 34 | 13 |
| Esm1 | ENSMUSG00000042379 | 32 | 4 | 64 | 10 | 34 | 9 | 54 | 17 |
| Etv4 | ENSMUSG00000017724 | 8 | 5 | 29 | 5 | 8 | 3 | 26 | 18 |
| Etv6 | ENSMUSG00000030199 | 527 | 36 | 643 | 63 | 544 | 47 | 566 | 37 |
| Evi2a | ENSMUSG00000078771 | 85 | 15 | 172 | 58 | 103 | 15 | 147 | 30 |
| Evi2b | ENSMUSG00000093938 | 37 | 8 | 106 | 37 | 54 | 8 | 77 | 17 |
| Ezh2 | ENSMUSG00000029687 | 114 | 28 | 176 | 40 | 98 | 20 | 147 | 19 |
| Fads2 | ENSMUSG00000024665 | 75 | 13 | 108 | 13 | 83 | 14 | 97 | 12 |
| Fam129a | ENSMUSG00000026483 | 681 | 70 | 925 | 196 | 630 | 59 | 727 | 64 |
| Fam167b | ENSMUSG00000050493 | 7 | 3 | 20 | 8 | 8 | 4 | 17 | 5 |
| Fam171b | ENSMUSG00000048388 | 42 | 7 | 93 | 55 | 48 | 6 | 66 | 20 |
| Fam177a | ENSMUSG00000095595 | 702 | 134 | 843 | 123 | 776 | 69 | 888 | 27 |
| Fam83d | ENSMUSG00000027654 | 6 | 3 | 27 | 7 | 5 | 2 | 17 | 9 |
| Fam91a1 | ENSMUSG00000037119 | 520 | 42 | 707 | 99 | 525 | 28 | 606 | 35 |
| Fancd2 | ENSMUSG00000034023 | 13 | 4 | 33 | 8 | 13 | 6 | 28 | 18 |
| Fap | ENSMUSG00000000392 | 100 | 8 | 185 | 87 | 98 | 34 | 134 | 42 |
| Farp1 | ENSMUSG00000025555 | 294 | 13 | 417 | 92 | 304 | 53 | 390 | 72 |
| Fat1 | ENSMUSG00000070047 | 995 | 230 | 1619 | 361 | 886 | 123 | 1302 | 330 |
| Fbn2 | ENSMUSG00000024598 | 25 | 8 | 112 | 120 | 38 | 10 | 67 | 32 |
| Fcer1g | ENSMUSG00000058715 | 141 | 25 | 259 | 105 | 135 | 20 | 195 | 28 |
| Fcgr1 | ENSMUSG00000015947 | 58 | 17 | 140 | 69 | 68 | 8 | 103 | 30 |
| Fermt3 | ENSMUSG00000024965 | 71 | 21 | 121 | 22 | 77 | 11 | 108 | 24 |
| Fes | ENSMUSG00000053158 | 170 | 34 | 266 | 19 | 189 | 43 | 230 | 55 |
| Fgd3 | ENSMUSG00000037946 | 46 | 14 | 84 | 22 | 51 | 10 | 76 | 20 |
| Fgl2 | ENSMUSG00000039899 | 863 | 65 | 1693 | 575 | 912 | 106 | 1331 | 391 |
| Fgr | ENSMUSG00000028874 | 10 | 5 | 30 | 20 | 9 | 4 | 17 | 11 |
| Fhl1 | ENSMUSG00000023092 | 1773 | 169 | 2590 | 350 | 1838 | 138 | 2464 | 537 |
| Fibin | ENSMUSG00000074971 | 181 | 32 | 511 | 420 | 193 | 39 | 346 | 121 |
| Flt3 | ENSMUSG00000042817 | 2 | 2 | 16 | 13 | 4 | 3 | 8 | 7 |
| Fmod | ENSMUSG00000041559 | 30 | 16 | 384 | 621 | 81 | 65 | 169 | 122 |
| Fmr1 | ENSMUSG00000000838 | 269 | 36 | 431 | 176 | 300 | 35 | 342 | 37 |
| Fn1 | ENSMUSG00000026193 | 1579 | 300 | 7830 | 6886 | 1494 | 199 | 4711 | 3367 |
| Foxs1 | ENSMUSG00000074676 | 34 | 11 | 64 | 13 | 46 | 5 | 62 | 22 |
| Frem1 | ENSMUSG00000059049 | 8 | 10 | 51 | 68 | 8 | 4 | 31 | 24 |
| Fut11 | ENSMUSG00000039357 | 204 | 20 | 286 | 68 | 209 | 19 | 246 | 19 |
| Fxyd6 | ENSMUSG00000066705 | 485 | 36 | 809 | 394 | 515 | 53 | 620 | 96 |
| Fyb | ENSMUSG00000022148 | 104 | 44 | 269 | 144 | 111 | 29 | 216 | 57 |
| Fzd1 | ENSMUSG00000044674 | 164 | 18 | 259 | 65 | 173 | 23 | 223 | 25 |
| G2e3 | ENSMUSG00000035293 | 143 | 24 | 208 | 59 | 161 | 20 | 204 | 41 |
| Gak | ENSMUSG00000062234 | 517 | 39 | 618 | 25 | 529 | 64 | 553 | 30 |
| Garem2 | ENSMUSG00000044576 | 3 | 3 | 13 | 9 | 3 | 2 | 7 | 4 |
| Gas7 | ENSMUSG00000033066 | 461 | 108 | 644 | 104 | 455 | 58 | 590 | 51 |
| Gatm | ENSMUSG00000027199 | 79 | 9 | 129 | 34 | 80 | 14 | 103 | 20 |
| Gcnt1 | ENSMUSG00000038843 | 85 | 7 | 138 | 43 | 89 | 8 | 117 | 12 |
| Gcnt4 | ENSMUSG00000091387 | 3 | 2 | 26 | 31 | 3 | 1 | 14 | 10 |
| Gem | ENSMUSG00000028214 | 71 | 22 | 119 | 13 | 95 | 18 | 126 | 28 |
| Gen1 | ENSMUSG00000051235 | 17 | 8 | 43 | 15 | 30 | 19 | 41 | 16 |
| Gjc1 | ENSMUSG00000034520 | 443 | 31 | 538 | 31 | 438 | 34 | 472 | 43 |

| | | | | | | | | | |
|---|---|---|---|---|---|---|---|---|---|
| Glipr1 | ENSMUSG00000056888 | 19 | 3 | 61 | 50 | 17 | 5 | 34 | 19 |
| Gm15675 | ENSMUSG00000086825 | 23 | 4 | 56 | 23 | 34 | 12 | 43 | 15 |
| Gm1966 | ENSMUSG00000073902 | 34 | 9 | 90 | 39 | 50 | 9 | 72 | 22 |
| Gm2026 | ENSMUSG00000078886 | 29 | 10 | 71 | 41 | 64 | 45 | 50 | 32 |
| Gm20559 | ENSMUSG00000106734 | 158 | 25 | 249 | 51 | 169 | 19 | 173 | 37 |
| Gm30873 | ENSMUSG00000109341 | 6 | 4 | 25 | 10 | 11 | 7 | 20 | 4 |
| Gm36161 | ENSMUSG00000114608 | 18 | 6 | 51 | 47 | 19 | 6 | 34 | 17 |
| Gm3636 | ENSMUSG00000091754 | 16 | 5 | 43 | 19 | 24 | 10 | 23 | 5 |
| Gm39214 | ENSMUSG00000109754 | 252 | 32 | 351 | 45 | 256 | 25 | 363 | 83 |
| Gm42047 | ENSMUSG00000110631 | 86 | 19 | 248 | 70 | 75 | 22 | 169 | 103 |
| Gm45705 | ENSMUSG00000110481 | 13 | 3 | 29 | 7 | 14 | 4 | 21 | 3 |
| Gm47761 | ENSMUSG00000112478 | 9 | 2 | 26 | 8 | 7 | 2 | 17 | 10 |
| Gm49342 | ENSMUSG00000021871 | 79 | 19 | 135 | 20 | 95 | 11 | 98 | 23 |
| Gm5431 | ENSMUSG00000058163 | 36 | 7 | 65 | 15 | 44 | 6 | 50 | 7 |
| Gm6377 | ENSMUSG00000048621 | 4 | 2 | 18 | 12 | 13 | 16 | 12 | 6 |
| Gm8995 | ENSMUSG00000063286 | 770 | 80 | 1033 | 133 | 749 | 121 | 889 | 142 |
| Gmip | ENSMUSG00000036246 | 79 | 23 | 131 | 23 | 79 | 20 | 106 | 20 |
| Gnai2 | ENSMUSG00000032562 | 4185 | 325 | 4943 | 197 | 4287 | 161 | 4768 | 360 |
| Gnai3 | ENSMUSG00000000001 | 721 | 11 | 942 | 133 | 676 | 66 | 788 | 53 |
| Gnao1 | ENSMUSG00000031748 | 446 | 40 | 615 | 131 | 524 | 114 | 584 | 73 |
| Golim4 | ENSMUSG00000034109 | 732 | 55 | 1009 | 144 | 790 | 46 | 896 | 57 |
| Gpc6 | ENSMUSG00000058571 | 240 | 29 | 347 | 74 | 228 | 14 | 308 | 59 |
| Gpnmb | ENSMUSG00000029816 | 23 | 2 | 58 | 33 | 25 | 12 | 38 | 17 |
| Gpr176 | ENSMUSG00000040133 | 6 | 3 | 37 | 26 | 7 | 1 | 23 | 18 |
| Gpr34 | ENSMUSG00000040229 | 63 | 25 | 126 | 68 | 77 | 29 | 101 | 26 |
| Gpr39 | ENSMUSG00000026343 | 4 | 2 | 28 | 20 | 6 | 3 | 21 | 14 |
| Gpr65 | ENSMUSG00000021886 | 44 | 9 | 108 | 76 | 42 | 12 | 70 | 21 |
| Gpr68 | ENSMUSG00000047415 | 3 | 3 | 18 | 4 | 8 | 2 | 10 | 7 |
| Gpx8 | ENSMUSG00000021760 | 491 | 35 | 682 | 134 | 493 | 63 | 644 | 76 |
| Gria3 | ENSMUSG00000001986 | 27 | 6 | 74 | 55 | 40 | 11 | 71 | 37 |
| Gsap | ENSMUSG00000039934 | 66 | 8 | 121 | 24 | 74 | 13 | 95 | 10 |
| Gxylt2 | ENSMUSG00000030074 | 171 | 37 | 519 | 541 | 157 | 23 | 308 | 184 |
| Hacd4 | ENSMUSG00000028497 | 224 | 6 | 334 | 92 | 257 | 33 | 288 | 32 |
| Has2 | ENSMUSG00000022367 | 27 | 6 | 50 | 13 | 28 | 8 | 42 | 6 |
| Haus8 | ENSMUSG00000035439 | 242 | 44 | 381 | 64 | 261 | 50 | 360 | 59 |
| Havcr2 | ENSMUSG00000020399 | 17 | 6 | 42 | 21 | 12 | 4 | 23 | 11 |
| Hbb-bt | ENSMUSG00000073940 | 3565 | 1643 | 6280 | 1861 | 4671 | 1644 | 8009 | 3238 |
| Hck | ENSMUSG00000003283 | 35 | 10 | 88 | 47 | 39 | 12 | 58 | 8 |
| Hdac1 | ENSMUSG00000028800 | 455 | 34 | 557 | 48 | 447 | 37 | 529 | 38 |
| Hexa | ENSMUSG00000025232 | 1121 | 111 | 1417 | 213 | 1139 | 67 | 1382 | 81 |
| Hexb | ENSMUSG00000021665 | 487 | 58 | 816 | 323 | 492 | 50 | 666 | 110 |
| Hjurp | ENSMUSG00000044783 | 577 | 122 | 775 | 43 | 589 | 90 | 674 | 87 |
| Hmgb2 | ENSMUSG00000054717 | 115 | 21 | 211 | 30 | 135 | 37 | 202 | 60 |
| Hmgn3 | ENSMUSG00000066456 | 78 | 20 | 130 | 43 | 71 | 15 | 108 | 14 |
| Hpgd | ENSMUSG00000031613 | 182 | 26 | 300 | 44 | 240 | 31 | 258 | 32 |
| Hpgds | ENSMUSG00000029919 | 67 | 16 | 152 | 74 | 91 | 33 | 92 | 21 |
| Hspg2 | ENSMUSG00000028763 | 12506 | 1017 | 15279 | 1628 | 12610 | 707 | 14990 | 1190 |
| Iffo2 | ENSMUSG00000041025 | 210 | 56 | 296 | 34 | 208 | 41 | 260 | 55 |
| Ifi203 | ENSMUSG00000039997 | 891 | 161 | 1230 | 255 | 880 | 117 | 940 | 84 |
| Ifi209 | ENSMUSG00000043263 | 52 | 15 | 126 | 69 | 50 | 17 | 92 | 45 |
| Ifi211 | ENSMUSG00000026536 | 185 | 36 | 351 | 104 | 178 | 18 | 256 | 21 |
| Ifi30 | ENSMUSG00000031838 | 101 | 27 | 193 | 76 | 87 | 20 | 127 | 31 |
| Ifih1 | ENSMUSG00000026896 | 299 | 32 | 446 | 46 | 329 | 56 | 378 | 53 |
| Ifit1 | ENSMUSG00000034459 | 187 | 43 | 329 | 102 | 161 | 24 | 234 | 21 |
| Ifit2 | ENSMUSG00000045932 | 547 | 92 | 934 | 140 | 539 | 58 | 669 | 52 |
| Ifit3 | ENSMUSG00000074896 | 336 | 93 | 626 | 210 | 311 | 45 | 430 | 29 |
| Ifit3b | ENSMUSG00000062488 | 122 | 30 | 208 | 57 | 123 | 22 | 150 | 9 |
| Igf1 | ENSMUSG00000020053 | 384 | 78 | 1064 | 595 | 422 | 57 | 770 | 126 |
| Igf2bp2 | ENSMUSG00000033581 | 72 | 26 | 122 | 41 | 102 | 14 | 135 | 38 |
| Igfbp2 | ENSMUSG00000039323 | 1 | 1 | 24 | 20 | 0 | 1 | 4 | 4 |
| Igfbp5 | ENSMUSG00000026185 | 2981 | 305 | 5165 | 1633 | 3493 | 830 | 4402 | 921 |
| Ighm | ENSMUSG00000076617 | 206 | 22 | 302 | 49 | 234 | 29 | 292 | 34 |
| Igsf10 | ENSMUSG00000036334 | 145 | 33 | 315 | 123 | 162 | 38 | 247 | 69 |
| Ikzf1 | ENSMUSG00000018654 | 64 | 21 | 119 | 32 | 75 | 10 | 86 | 10 |

| | | | | | | | | | |
|---|---|---|---|---|---|---|---|---|---|
| Il13ra1 | ENSMUSG00000017057 | 1206 | 209 | 1526 | 234 | 1225 | 65 | 1443 | 213 |
| Il18rap | ENSMUSG00000026068 | 5 | 4 | 17 | 7 | 3 | 2 | 6 | 4 |
| Il1b | ENSMUSG00000027398 | 20 | 8 | 71 | 68 | 17 | 7 | 29 | 8 |
| Il21r | ENSMUSG00000030745 | 15 | 3 | 46 | 25 | 17 | 2 | 35 | 6 |
| Ildr2 | ENSMUSG00000040612 | 25 | 9 | 63 | 21 | 39 | 23 | 51 | 20 |
| Irf5 | ENSMUSG00000029771 | 73 | 13 | 148 | 22 | 78 | 12 | 119 | 35 |
| Irf8 | ENSMUSG00000041515 | 118 | 13 | 229 | 76 | 128 | 21 | 193 | 26 |
| Isg15 | ENSMUSG00000035692 | 59 | 12 | 120 | 28 | 67 | 16 | 93 | 12 |
| Isg20 | ENSMUSG00000039236 | 86 | 30 | 130 | 16 | 102 | 27 | 144 | 32 |
| Islr | ENSMUSG00000037206 | 560 | 95 | 865 | 285 | 608 | 50 | 774 | 128 |
| Itga4 | ENSMUSG00000027009 | 88 | 24 | 170 | 60 | 100 | 20 | 120 | 28 |
| Itgav | ENSMUSG00000027087 | 584 | 81 | 920 | 293 | 645 | 110 | 863 | 204 |
| Itgax | ENSMUSG00000030789 | 14 | 13 | 49 | 40 | 11 | 3 | 41 | 38 |
| Itgb1 | ENSMUSG00000025809 | 10648 | 748 | 12926 | 1642 | 10469 | 431 | 12265 | 1179 |
| Itgb2 | ENSMUSG00000000290 | 175 | 40 | 280 | 60 | 156 | 29 | 225 | 46 |
| Itgb3 | ENSMUSG00000020689 | 55 | 19 | 135 | 26 | 74 | 35 | 101 | 24 |
| Itgb8 | ENSMUSG00000025321 | 44 | 17 | 94 | 45 | 58 | 14 | 57 | 7 |
| Itgbl1 | ENSMUSG00000032925 | 238 | 42 | 795 | 674 | 281 | 46 | 513 | 150 |
| Itih2 | ENSMUSG00000037254 | 10 | 2 | 36 | 22 | 10 | 1 | 20 | 5 |
| Itm2a | ENSMUSG00000031239 | 204 | 26 | 483 | 405 | 214 | 32 | 305 | 66 |
| Kcne1 | ENSMUSG00000039639 | 91 | 46 | 147 | 42 | 92 | 14 | 127 | 27 |
| Kctd10 | ENSMUSG00000001098 | 1166 | 95 | 1425 | 67 | 1193 | 154 | 1325 | 119 |
| Kctd12 | ENSMUSG00000098557 | 889 | 110 | 1244 | 276 | 915 | 43 | 1073 | 34 |
| Kctd12b | ENSMUSG00000041633 | 620 | 130 | 959 | 330 | 830 | 137 | 891 | 240 |
| Kctd15 | ENSMUSG00000030499 | 38 | 5 | 65 | 13 | 39 | 8 | 54 | 12 |
| Kdelr3 | ENSMUSG00000010830 | 122 | 17 | 231 | 83 | 128 | 25 | 188 | 48 |
| Kif18a | ENSMUSG00000027115 | 33 | 7 | 55 | 18 | 33 | 3 | 53 | 12 |
| Kif20b | ENSMUSG00000024795 | 44 | 17 | 104 | 56 | 35 | 19 | 80 | 25 |
| Kirrel | ENSMUSG00000041734 | 354 | 32 | 515 | 106 | 351 | 32 | 432 | 45 |
| Klhl29 | ENSMUSG00000020627 | 12 | 4 | 36 | 26 | 16 | 6 | 21 | 5 |
| Krt18 | ENSMUSG00000023043 | 11 | 4 | 41 | 28 | 17 | 8 | 20 | 7 |
| Lacc1 | ENSMUSG00000044350 | 110 | 10 | 170 | 40 | 111 | 18 | 129 | 16 |
| Lair1 | ENSMUSG00000055541 | 93 | 25 | 193 | 59 | 88 | 18 | 152 | 37 |
| Lbp | ENSMUSG00000016024 | 114 | 14 | 211 | 51 | 141 | 27 | 174 | 37 |
| Ldlrad4 | ENSMUSG00000024544 | 207 | 23 | 293 | 57 | 198 | 24 | 230 | 26 |
| Lhfp | ENSMUSG00000048332 | 883 | 63 | 1348 | 372 | 958 | 40 | 1113 | 122 |
| Lilrb4a | ENSMUSG00000112148 | 184 | 42 | 406 | 123 | 195 | 44 | 266 | 91 |
| Lix1l | ENSMUSG00000049288 | 862 | 45 | 1020 | 63 | 903 | 22 | 948 | 26 |
| Lmna | ENSMUSG00000028063 | 1287 | 73 | 1552 | 83 | 1330 | 102 | 1508 | 161 |
| Lmnb1 | ENSMUSG00000024590 | 122 | 21 | 210 | 37 | 129 | 15 | 185 | 47 |
| Lox | ENSMUSG00000024529 | 89 | 9 | 1050 | 1321 | 94 | 21 | 476 | 436 |
| Loxl3 | ENSMUSG00000000693 | 104 | 36 | 311 | 233 | 87 | 12 | 196 | 70 |
| Lpcat2 | ENSMUSG00000033192 | 36 | 10 | 94 | 53 | 31 | 6 | 64 | 11 |
| Lpp | ENSMUSG00000033306 | 1398 | 218 | 1794 | 95 | 1378 | 115 | 1689 | 226 |
| Lpxn | ENSMUSG00000024696 | 18 | 5 | 55 | 29 | 22 | 7 | 34 | 9 |
| Lrmp | ENSMUSG00000030263 | 36 | 7 | 68 | 14 | 45 | 10 | 53 | 5 |
| Lrrc32 | ENSMUSG00000090958 | 328 | 91 | 455 | 36 | 361 | 64 | 432 | 52 |
| Lrrc59 | ENSMUSG00000020869 | 766 | 27 | 883 | 28 | 772 | 44 | 888 | 71 |
| Lsp1 | ENSMUSG00000018819 | 614 | 136 | 984 | 350 | 600 | 112 | 808 | 176 |
| Lst1 | ENSMUSG00000073412 | 16 | 5 | 43 | 19 | 20 | 5 | 34 | 7 |
| Ltbp2 | ENSMUSG00000002020 | 94 | 45 | 964 | 1120 | 126 | 62 | 578 | 514 |
| Ltbp3 | ENSMUSG00000024940 | 530 | 28 | 924 | 412 | 618 | 50 | 772 | 112 |
| Lum | ENSMUSG00000036446 | 1447 | 238 | 3824 | 2686 | 1553 | 192 | 2795 | 564 |
| Lxn | ENSMUSG00000047557 | 65 | 9 | 139 | 25 | 70 | 19 | 104 | 26 |
| Lyl1 | ENSMUSG00000034041 | 75 | 11 | 111 | 12 | 85 | 3 | 91 | 4 |
| Maf | ENSMUSG00000055435 | 389 | 56 | 552 | 118 | 424 | 20 | 460 | 80 |
| Maff | ENSMUSG00000042622 | 133 | 18 | 220 | 64 | 128 | 36 | 187 | 43 |
| Malt1 | ENSMUSG00000032688 | 196 | 29 | 270 | 27 | 193 | 17 | 212 | 27 |
| Mapre1 | ENSMUSG00000027479 | 1639 | 151 | 1876 | 92 | 1651 | 91 | 1859 | 64 |
| Marcks | ENSMUSG00000069662 | 819 | 115 | 1485 | 784 | 999 | 154 | 1070 | 121 |
| Mastl | ENSMUSG00000026779 | 21 | 19 | 58 | 17 | 22 | 7 | 58 | 28 |
| Matn2 | ENSMUSG00000022324 | 417 | 57 | 667 | 192 | 429 | 35 | 559 | 39 |
| Mcm5 | ENSMUSG00000005410 | 84 | 12 | 162 | 45 | 83 | 19 | 169 | 35 |
| Mcm6 | ENSMUSG00000026355 | 230 | 32 | 397 | 77 | 216 | 18 | 363 | 70 |

| | | | | | | | | | |
|---|---|---|---|---|---|---|---|---|---|
| Mcub | ENSMUSG00000027994 | 41 | 14 | 77 | 15 | 37 | 4 | 50 | 10 |
| Mdk | ENSMUSG00000027239 | 26 | 4 | 95 | 130 | 24 | 6 | 43 | 34 |
| Meg3 | ENSMUSG00000021268 | 123 | 46 | 231 | 107 | 121 | 17 | 140 | 38 |
| Megf10 | ENSMUSG00000024593 | 19 | 6 | 49 | 31 | 30 | 3 | 40 | 15 |
| Mex3c | ENSMUSG00000037253 | 486 | 50 | 607 | 52 | 503 | 15 | 557 | 37 |
| Mfap2 | ENSMUSG00000060572 | 20 | 2 | 69 | 61 | 23 | 4 | 41 | 14 |
| Mfap3 | ENSMUSG00000020522 | 454 | 25 | 541 | 33 | 431 | 20 | 503 | 22 |
| Mgam | ENSMUSG00000068587 | 9 | 4 | 32 | 24 | 15 | 4 | 30 | 21 |
| Mgat2 | ENSMUSG00000043998 | 423 | 27 | 547 | 64 | 443 | 34 | 503 | 31 |
| Mgp | ENSMUSG00000030218 | 1671 | 257 | 3299 | 1832 | 1627 | 149 | 2424 | 267 |
| Milr1 | ENSMUSG00000040528 | 16 | 6 | 43 | 18 | 16 | 4 | 35 | 10 |
| Mis18a | ENSMUSG00000022978 | 46 | 8 | 79 | 17 | 61 | 14 | 68 | 8 |
| Mkrn1 | ENSMUSG00000029922 | 598 | 102 | 779 | 95 | 681 | 119 | 829 | 128 |
| Mmp14 | ENSMUSG00000000957 | 291 | 39 | 862 | 471 | 332 | 50 | 582 | 198 |
| Mmp16 | ENSMUSG00000028226 | 14 | 6 | 34 | 13 | 18 | 6 | 22 | 9 |
| Mns1 | ENSMUSG00000032221 | 9 | 2 | 49 | 8 | 14 | 7 | 28 | 16 |
| Mob1a | ENSMUSG00000043131 | 1089 | 121 | 1399 | 155 | 1053 | 73 | 1248 | 99 |
| Mpeg1 | ENSMUSG00000046805 | 387 | 93 | 1323 | 966 | 395 | 77 | 809 | 361 |
| Mpzl1 | ENSMUSG00000026566 | 252 | 38 | 353 | 54 | 258 | 43 | 287 | 41 |
| Mrc2 | ENSMUSG00000020695 | 462 | 41 | 913 | 394 | 473 | 31 | 716 | 159 |
| Ms4a14 | ENSMUSG00000099398 | 18 | 2 | 74 | 52 | 16 | 6 | 34 | 17 |
| Ms4a4b | ENSMUSG00000056290 | 4 | 2 | 18 | 15 | 8 | 4 | 9 | 4 |
| Ms4a4c | ENSMUSG00000024675 | 8 | 4 | 38 | 33 | 10 | 3 | 16 | 8 |
| Ms4a6d | ENSMUSG00000024679 | 61 | 21 | 136 | 54 | 54 | 15 | 109 | 45 |
| Ms4a7 | ENSMUSG00000024672 | 65 | 11 | 296 | 303 | 51 | 5 | 166 | 95 |
| Mtfr2 | ENSMUSG00000019992 | 4 | 2 | 15 | 5 | 3 | 1 | 10 | 2 |
| Mthfd2 | ENSMUSG00000005667 | 41 | 11 | 81 | 14 | 49 | 16 | 81 | 18 |
| Mtmr11 | ENSMUSG00000045934 | 69 | 26 | 114 | 28 | 68 | 30 | 107 | 32 |
| Mtpn | ENSMUSG00000029840 | 1161 | 76 | 1505 | 277 | 1222 | 89 | 1380 | 115 |
| Mx1 | ENSMUSG00000000386 | 20 | 10 | 55 | 32 | 17 | 6 | 22 | 11 |
| Mxra8 | ENSMUSG00000029070 | 431 | 61 | 782 | 329 | 511 | 145 | 714 | 119 |
| Myc | ENSMUSG00000022346 | 48 | 8 | 98 | 19 | 58 | 7 | 95 | 39 |
| Myef2 | ENSMUSG00000027201 | 185 | 24 | 262 | 27 | 174 | 6 | 207 | 19 |
| Myh7 | ENSMUSG00000053093 | 1672 | 510 | 20844 | 27743 | 2490 | 1006 | 11438 | 8811 |
| Myh9 | ENSMUSG00000022443 | 4467 | 917 | 6229 | 813 | 4465 | 679 | 5733 | 609 |
| Myl9 | ENSMUSG00000067818 | 507 | 113 | 744 | 105 | 504 | 152 | 536 | 41 |
| Myo1e | ENSMUSG00000032220 | 382 | 52 | 541 | 82 | 377 | 40 | 487 | 59 |
| Myo1g | ENSMUSG00000020437 | 50 | 9 | 89 | 21 | 55 | 12 | 90 | 33 |
| Myo5a | ENSMUSG00000034593 | 359 | 55 | 615 | 165 | 371 | 102 | 518 | 152 |
| Naalad2 | ENSMUSG00000043943 | 352 | 43 | 575 | 192 | 346 | 54 | 447 | 64 |
| Nab2 | ENSMUSG00000025402 | 178 | 29 | 277 | 77 | 177 | 31 | 248 | 30 |
| Nav1 | ENSMUSG00000009418 | 1498 | 290 | 1893 | 171 | 1457 | 90 | 1737 | 179 |
| Ncapd2 | ENSMUSG00000038252 | 298 | 45 | 437 | 112 | 363 | 44 | 394 | 55 |
| Nedd9 | ENSMUSG00000021365 | 728 | 336 | 1059 | 86 | 794 | 203 | 936 | 202 |
| Neil3 | ENSMUSG00000039396 | 10 | 3 | 35 | 11 | 13 | 11 | 23 | 8 |
| Neurl3 | ENSMUSG00000047180 | 159 | 33 | 303 | 74 | 176 | 25 | 246 | 35 |
| Nfam1 | ENSMUSG00000058099 | 110 | 22 | 202 | 30 | 118 | 21 | 160 | 20 |
| Nfkbie | ENSMUSG00000023947 | 37 | 6 | 81 | 15 | 38 | 5 | 56 | 8 |
| Nfkbiz | ENSMUSG00000035356 | 92 | 25 | 164 | 59 | 103 | 9 | 151 | 32 |
| Nhsl2 | ENSMUSG00000079481 | 276 | 31 | 399 | 49 | 272 | 39 | 339 | 24 |
| Nkd1 | ENSMUSG00000031661 | 22 | 8 | 42 | 10 | 25 | 6 | 30 | 7 |
| Nlgn2 | ENSMUSG00000051790 | 167 | 30 | 260 | 63 | 167 | 33 | 228 | 39 |
| Nmrk2 | ENSMUSG00000004939 | 73 | 9 | 158 | 71 | 68 | 15 | 132 | 76 |
| Nmt2 | ENSMUSG00000026643 | 286 | 28 | 378 | 59 | 277 | 25 | 339 | 48 |
| Nnmt | ENSMUSG00000032271 | 42 | 5 | 88 | 24 | 47 | 10 | 60 | 8 |
| Nox4 | ENSMUSG00000030562 | 31 | 5 | 162 | 173 | 29 | 9 | 64 | 30 |
| Npdc1 | ENSMUSG00000015094 | 238 | 45 | 344 | 63 | 227 | 27 | 280 | 28 |
| Npl | ENSMUSG00000042684 | 26 | 8 | 47 | 8 | 28 | 8 | 51 | 12 |
| Npnt | ENSMUSG00000040998 | 22 | 7 | 57 | 25 | 31 | 14 | 28 | 9 |
| Nptxr | ENSMUSG00000022421 | 69 | 12 | 105 | 10 | 64 | 14 | 94 | 14 |
| Nrep | ENSMUSG00000042834 | 535 | 52 | 997 | 503 | 559 | 60 | 711 | 91 |
| Nrros | ENSMUSG00000052384 | 157 | 25 | 231 | 32 | 162 | 27 | 220 | 28 |
| Nts | ENSMUSG00000019890 | 21 | 8 | 42 | 6 | 25 | 4 | 33 | 13 |
| Nucb2 | ENSMUSG00000030659 | 137 | 19 | 239 | 55 | 144 | 20 | 174 | 24 |

| | | | | | | | | | |
|---|---|---|---|---|---|---|---|---|---|
| Nupr1 | ENSMUSG00000030717 | 60 | 10 | 196 | 120 | 74 | 8 | 130 | 33 |
| Nusap1 | ENSMUSG00000027306 | 48 | 17 | 132 | 40 | 45 | 9 | 116 | 36 |
| Nxpe4 | ENSMUSG00000044229 | 315 | 33 | 430 | 45 | 320 | 35 | 379 | 48 |
| Nxpe5 | ENSMUSG00000047592 | 11 | 6 | 35 | 8 | 9 | 4 | 28 | 15 |
| Oas1a | ENSMUSG00000052776 | 82 | 18 | 149 | 24 | 88 | 14 | 121 | 8 |
| Oas3 | ENSMUSG00000032661 | 6 | 1 | 34 | 18 | 9 | 3 | 18 | 9 |
| Oasl1 | ENSMUSG00000041827 | 32 | 9 | 56 | 13 | 38 | 7 | 48 | 5 |
| Oasl2 | ENSMUSG00000029561 | 269 | 62 | 498 | 109 | 332 | 69 | 402 | 43 |
| Olfr558 | ENSMUSG00000070423 | 167 | 26 | 236 | 25 | 154 | 13 | 204 | 33 |
| Olfr56 | ENSMUSG00000040328 | 6 | 3 | 19 | 7 | 8 | 2 | 10 | 5 |
| Omd | ENSMUSG00000048368 | 16 | 4 | 58 | 62 | 32 | 17 | 29 | 10 |
| Otulinl | ENSMUSG00000056069 | 53 | 9 | 133 | 67 | 61 | 15 | 119 | 66 |
| P2rx7 | ENSMUSG00000029468 | 140 | 30 | 223 | 46 | 141 | 24 | 188 | 49 |
| P3h1 | ENSMUSG00000028641 | 217 | 25 | 313 | 43 | 242 | 18 | 308 | 55 |
| P4ha3 | ENSMUSG00000051048 | 2 | 2 | 23 | 32 | 1 | 1 | 11 | 10 |
| Pak1 | ENSMUSG00000030774 | 34 | 10 | 89 | 41 | 49 | 20 | 57 | 18 |
| Panx1 | ENSMUSG00000031934 | 35 | 11 | 98 | 49 | 42 | 21 | 76 | 30 |
| Parp9 | ENSMUSG00000022906 | 467 | 40 | 651 | 71 | 484 | 24 | 548 | 44 |
| Parpbp | ENSMUSG00000035365 | 8 | 4 | 24 | 9 | 7 | 2 | 18 | 11 |
| Pcdh17 | ENSMUSG00000035566 | 299 | 64 | 458 | 76 | 316 | 37 | 426 | 89 |
| Pcsk5 | ENSMUSG00000024713 | 67 | 10 | 129 | 24 | 84 | 4 | 119 | 16 |
| Pdgfrl | ENSMUSG00000031595 | 48 | 13 | 182 | 194 | 44 | 4 | 108 | 62 |
| Pdk3 | ENSMUSG00000035232 | 44 | 14 | 83 | 17 | 53 | 21 | 69 | 7 |
| Pdlim2 | ENSMUSG00000022090 | 106 | 9 | 175 | 59 | 121 | 17 | 153 | 29 |
| Pdlim3 | ENSMUSG00000031636 | 187 | 22 | 301 | 87 | 190 | 20 | 249 | 48 |
| Pdpn | ENSMUSG00000028583 | 91 | 16 | 188 | 37 | 105 | 17 | 159 | 30 |
| Pgm2 | ENSMUSG00000029171 | 116 | 24 | 174 | 30 | 127 | 15 | 159 | 19 |
| Phf11b | ENSMUSG00000091649 | 41 | 10 | 91 | 36 | 39 | 9 | 65 | 19 |
| Phldb2 | ENSMUSG00000033149 | 633 | 66 | 824 | 141 | 626 | 28 | 795 | 54 |
| Pi15 | ENSMUSG00000067780 | 74 | 15 | 180 | 121 | 89 | 17 | 105 | 28 |
| Piezo1 | ENSMUSG00000014444 | 764 | 311 | 1009 | 184 | 762 | 82 | 868 | 67 |
| Piezo2 | ENSMUSG00000041482 | 24 | 11 | 103 | 100 | 27 | 10 | 60 | 26 |
| Pif1 | ENSMUSG00000041064 | 7 | 7 | 20 | 9 | 7 | 4 | 13 | 5 |
| Pik3ap1 | ENSMUSG00000025017 | 55 | 12 | 132 | 58 | 64 | 10 | 102 | 14 |
| Pik3c2a | ENSMUSG00000030660 | 644 | 90 | 821 | 49 | 571 | 41 | 614 | 49 |
| Pik3cd | ENSMUSG00000039936 | 104 | 25 | 184 | 49 | 123 | 16 | 148 | 46 |
| Pik3cg | ENSMUSG00000020573 | 96 | 11 | 139 | 24 | 110 | 14 | 125 | 19 |
| Pik3r5 | ENSMUSG00000020901 | 24 | 7 | 70 | 31 | 28 | 9 | 52 | 21 |
| Pkd2 | ENSMUSG00000034462 | 934 | 68 | 1341 | 308 | 928 | 43 | 1049 | 85 |
| Pkhd1l1 | ENSMUSG00000038725 | 113 | 45 | 211 | 84 | 195 | 90 | 172 | 61 |
| Pkn3 | ENSMUSG00000026785 | 143 | 28 | 220 | 23 | 138 | 15 | 195 | 21 |
| Plac8 | ENSMUSG00000029322 | 10 | 2 | 54 | 49 | 9 | 5 | 21 | 10 |
| Plau | ENSMUSG00000021822 | 120 | 25 | 172 | 15 | 122 | 29 | 160 | 33 |
| Plaur | ENSMUSG00000046223 | 36 | 7 | 71 | 21 | 37 | 23 | 64 | 15 |
| Pld4 | ENSMUSG00000052160 | 158 | 21 | 294 | 102 | 174 | 44 | 228 | 58 |
| Plekha4 | ENSMUSG00000040428 | 34 | 16 | 87 | 40 | 51 | 13 | 73 | 38 |
| Plk1 | ENSMUSG00000030867 | 19 | 10 | 48 | 16 | 17 | 7 | 40 | 14 |
| Plk4 | ENSMUSG00000025758 | 70 | 9 | 110 | 26 | 68 | 16 | 112 | 44 |
| Plod3 | ENSMUSG00000004846 | 475 | 118 | 658 | 87 | 472 | 51 | 605 | 62 |
| Plp2 | ENSMUSG00000031146 | 570 | 31 | 793 | 128 | 592 | 14 | 724 | 133 |
| Plpp1 | ENSMUSG00000021759 | 1000 | 50 | 1160 | 56 | 1013 | 21 | 1087 | 43 |
| Plxdc2 | ENSMUSG00000026748 | 599 | 71 | 840 | 221 | 565 | 51 | 696 | 96 |
| Pou2f2 | ENSMUSG00000008496 | 37 | 5 | 109 | 53 | 61 | 19 | 105 | 35 |
| Ppfia1 | ENSMUSG00000037519 | 773 | 148 | 999 | 166 | 867 | 163 | 931 | 191 |
| Ppfibp1 | ENSMUSG00000016487 | 1707 | 135 | 2004 | 276 | 1903 | 142 | 1868 | 195 |
| Ppib | ENSMUSG00000032383 | 1085 | 51 | 1440 | 242 | 1046 | 78 | 1281 | 107 |
| Ppp1r15b | ENSMUSG00000046062 | 750 | 74 | 889 | 76 | 775 | 49 | 880 | 44 |
| Ppp1r18 | ENSMUSG00000034595 | 415 | 70 | 651 | 61 | 435 | 37 | 557 | 57 |
| Pqlc3 | ENSMUSG00000045679 | 95 | 9 | 171 | 87 | 101 | 15 | 126 | 18 |
| Prelp | ENSMUSG00000041577 | 1273 | 232 | 1575 | 178 | 1325 | 81 | 1511 | 142 |
| Prex1 | ENSMUSG00000039621 | 261 | 20 | 427 | 60 | 278 | 27 | 341 | 38 |
| Prim1 | ENSMUSG00000025395 | 42 | 9 | 75 | 26 | 45 | 8 | 65 | 14 |
| Prkcd | ENSMUSG00000021948 | 246 | 36 | 398 | 115 | 245 | 27 | 325 | 36 |
| Prr5l | ENSMUSG00000032841 | 24 | 3 | 49 | 21 | 25 | 8 | 44 | 16 |

| | | | | | | | | | |
|---|---|---|---|---|---|---|---|---|---|
| Prrt4 | ENSMUSG00000079654 | 41 | 13 | 75 | 15 | 44 | 3 | 68 | 12 |
| Prss23 | ENSMUSG00000039405 | 314 | 52 | 488 | 126 | 314 | 35 | 361 | 78 |
| Pstpip1 | ENSMUSG00000032322 | 21 | 5 | 54 | 31 | 21 | 3 | 37 | 15 |
| Ptafr | ENSMUSG00000056529 | 107 | 31 | 167 | 16 | 115 | 20 | 154 | 29 |
| Ptbp1 | ENSMUSG00000006498 | 741 | 116 | 1045 | 159 | 740 | 51 | 886 | 127 |
| Pthlh | ENSMUSG00000048776 | 10 | 3 | 24 | 10 | 10 | 4 | 20 | 5 |
| Ptk2b | ENSMUSG00000059456 | 161 | 11 | 248 | 52 | 179 | 24 | 246 | 35 |
| Ptk7 | ENSMUSG00000023972 | 64 | 11 | 121 | 51 | 79 | 10 | 101 | 16 |
| Ptms | ENSMUSG00000030122 | 1096 | 180 | 1556 | 255 | 1175 | 69 | 1354 | 123 |
| Ptn | ENSMUSG00000029838 | 35 | 10 | 527 | 950 | 47 | 13 | 109 | 85 |
| Ptpn1 | ENSMUSG00000027540 | 383 | 52 | 516 | 71 | 406 | 46 | 484 | 90 |
| Ptpn12 | ENSMUSG00000028771 | 720 | 48 | 925 | 82 | 719 | 64 | 779 | 35 |
| Ptpn18 | ENSMUSG00000026126 | 35 | 12 | 64 | 9 | 54 | 15 | 62 | 12 |
| Ptpn6 | ENSMUSG00000004266 | 114 | 27 | 202 | 70 | 107 | 24 | 164 | 25 |
| Ptprc | ENSMUSG00000026395 | 246 | 49 | 533 | 285 | 234 | 45 | 382 | 129 |
| Ptpre | ENSMUSG00000041836 | 163 | 23 | 260 | 20 | 183 | 21 | 214 | 24 |
| Ptprf | ENSMUSG00000033295 | 111 | 21 | 196 | 66 | 107 | 19 | 177 | 20 |
| Rab23 | ENSMUSG00000004768 | 193 | 52 | 284 | 32 | 226 | 32 | 252 | 58 |
| Rab32 | ENSMUSG00000019832 | 40 | 12 | 85 | 42 | 46 | 3 | 70 | 12 |
| Rab3il1 | ENSMUSG00000024663 | 211 | 18 | 296 | 51 | 216 | 26 | 283 | 32 |
| Rab5c | ENSMUSG00000019173 | 838 | 81 | 1059 | 21 | 918 | 29 | 1032 | 128 |
| Rab8b | ENSMUSG00000036943 | 406 | 32 | 589 | 93 | 439 | 43 | 531 | 48 |
| Rad50 | ENSMUSG00000020380 | 310 | 33 | 381 | 31 | 339 | 21 | 385 | 26 |
| Rai14 | ENSMUSG00000022246 | 173 | 18 | 300 | 114 | 173 | 15 | 241 | 57 |
| Rap1b | ENSMUSG00000052681 | 1590 | 124 | 2099 | 385 | 1642 | 155 | 1892 | 156 |
| Rasa4 | ENSMUSG00000004952 | 143 | 26 | 261 | 113 | 138 | 14 | 162 | 33 |
| Rbbp8 | ENSMUSG00000041238 | 177 | 8 | 233 | 32 | 161 | 22 | 196 | 17 |
| Rbm3 | ENSMUSG00000031167 | 1065 | 243 | 1408 | 328 | 1097 | 137 | 1294 | 160 |
| Rbp1 | ENSMUSG00000046402 | 203 | 16 | 395 | 267 | 185 | 21 | 311 | 56 |
| Reps2 | ENSMUSG00000040855 | 67 | 11 | 117 | 35 | 105 | 41 | 101 | 23 |
| Rflnb | ENSMUSG00000020846 | 1333 | 290 | 1883 | 230 | 1446 | 172 | 1704 | 352 |
| Rfx7 | ENSMUSG00000037674 | 332 | 31 | 421 | 33 | 374 | 29 | 376 | 17 |
| Rgs10 | ENSMUSG00000030844 | 114 | 9 | 181 | 62 | 108 | 10 | 146 | 24 |
| Rhoa | ENSMUSG00000007815 | 4789 | 93 | 5216 | 118 | 4862 | 194 | 5112 | 92 |
| Rhod | ENSMUSG00000041845 | 59 | 10 | 92 | 15 | 68 | 16 | 66 | 10 |
| Rhoj | ENSMUSG00000046768 | 564 | 95 | 704 | 75 | 573 | 74 | 656 | 48 |
| Rhou | ENSMUSG00000039960 | 110 | 28 | 173 | 59 | 135 | 9 | 173 | 41 |
| Rnase6 | ENSMUSG00000021880 | 9 | 5 | 29 | 15 | 13 | 7 | 14 | 2 |
| Robo1 | ENSMUSG00000022883 | 56 | 11 | 110 | 44 | 68 | 14 | 97 | 13 |
| Rpl10-ps3 | ENSMUSG00000058443 | 1037 | 413 | 1554 | 348 | 1563 | 299 | 1415 | 424 |
| Rpl3 | ENSMUSG00000060036 | 3377 | 296 | 4471 | 607 | 3263 | 335 | 4320 | 344 |
| Rps6ka1 | ENSMUSG00000003644 | 155 | 10 | 219 | 28 | 140 | 11 | 177 | 26 |
| Rtn4 | ENSMUSG00000020458 | 1597 | 115 | 2738 | 963 | 1691 | 68 | 2291 | 384 |
| Rtp4 | ENSMUSG00000033355 | 158 | 33 | 260 | 39 | 177 | 33 | 212 | 32 |
| Runx1 | ENSMUSG00000022952 | 63 | 29 | 176 | 93 | 54 | 10 | 115 | 46 |
| S1pr2 | ENSMUSG00000043895 | 122 | 33 | 225 | 28 | 161 | 13 | 210 | 36 |
| Samd14 | ENSMUSG00000047181 | 37 | 7 | 72 | 30 | 41 | 5 | 56 | 15 |
| Samsn1 | ENSMUSG00000022876 | 14 | 3 | 34 | 12 | 17 | 4 | 26 | 8 |
| Sat1 | ENSMUSG00000025283 | 655 | 35 | 1037 | 357 | 699 | 101 | 854 | 124 |
| Sbno2 | ENSMUSG00000035673 | 285 | 44 | 452 | 89 | 290 | 31 | 409 | 72 |
| Scara3 | ENSMUSG00000034463 | 56 | 13 | 114 | 45 | 58 | 2 | 87 | 26 |
| Scd2 | ENSMUSG00000025203 | 637 | 102 | 838 | 65 | 763 | 157 | 803 | 46 |
| Scn1b | ENSMUSG00000019194 | 235 | 13 | 365 | 140 | 268 | 19 | 330 | 60 |
| Scpep1 | ENSMUSG00000000278 | 287 | 48 | 480 | 174 | 308 | 44 | 404 | 65 |
| Scrn1 | ENSMUSG00000019124 | 212 | 35 | 302 | 33 | 202 | 45 | 226 | 61 |
| Scube3 | ENSMUSG00000038677 | 5 | 4 | 20 | 12 | 8 | 3 | 16 | 11 |
| Sdc1 | ENSMUSG00000020592 | 116 | 15 | 220 | 98 | 124 | 18 | 164 | 39 |
| Sdcbp | ENSMUSG00000028249 | 2167 | 214 | 2679 | 438 | 2118 | 92 | 2530 | 189 |
| Sdk1 | ENSMUSG00000039683 | 24 | 3 | 57 | 13 | 40 | 21 | 57 | 22 |
| Sec16b | ENSMUSG00000026589 | 57 | 7 | 120 | 43 | 63 | 10 | 73 | 13 |
| Sele | ENSMUSG00000026582 | 35 | 8 | 70 | 23 | 43 | 4 | 63 | 17 |
| Selplg | ENSMUSG00000048163 | 63 | 15 | 127 | 38 | 67 | 10 | 99 | 16 |
| Sema6d | ENSMUSG00000027200 | 1026 | 136 | 1377 | 210 | 1090 | 137 | 1238 | 97 |
| Serp1 | ENSMUSG00000027808 | 586 | 74 | 881 | 294 | 591 | 79 | 738 | 92 |

| | | | | | | | | |
|---|---|---|---|---|---|---|---|---|
| Serpina3g | ENSMUSG00000041481 | 9 | 5 | 29 | 16 | 12 | 5 | 21 | 10 |
| Serpina3i | ENSMUSG00000079014 | 3 | 2 | 17 | 15 | 2 | 2 | 10 | 8 |
| Serpina3n | ENSMUSG00000021091 | 176 | 100 | 698 | 430 | 134 | 41 | 495 | 374 |
| Serpinb1a | ENSMUSG00000044734 | 77 | 13 | 216 | 200 | 93 | 25 | 137 | 59 |
| Serpine1 | ENSMUSG00000037411 | 794 | 250 | 1688 | 848 | 832 | 284 | 1309 | 495 |
| Serpine2 | ENSMUSG00000026249 | 478 | 38 | 814 | 242 | 497 | 57 | 623 | 60 |
| Serping1 | ENSMUSG00000023224 | 1822 | 273 | 2473 | 457 | 1827 | 186 | 2279 | 296 |
| Sertad4 | ENSMUSG00000016262 | 105 | 17 | 221 | 144 | 112 | 18 | 169 | 32 |
| Sfrp1 | ENSMUSG00000031548 | 345 | 30 | 981 | 861 | 365 | 40 | 683 | 270 |
| Sfrp2 | ENSMUSG00000027996 | 43 | 4 | 313 | 445 | 47 | 8 | 154 | 132 |
| Sfxn3 | ENSMUSG00000025212 | 216 | 19 | 310 | 29 | 224 | 5 | 272 | 31 |
| Sgce | ENSMUSG00000004631 | 294 | 28 | 387 | 53 | 287 | 38 | 342 | 18 |
| Sh3bgrl | ENSMUSG00000031246 | 918 | 89 | 1283 | 246 | 887 | 122 | 1015 | 78 |
| Shc2 | ENSMUSG00000020312 | 50 | 9 | 88 | 30 | 52 | 6 | 71 | 5 |
| Shcbp1 | ENSMUSG00000022322 | 19 | 11 | 66 | 49 | 19 | 6 | 42 | 16 |
| Shisa4 | ENSMUSG00000041889 | 45 | 8 | 85 | 28 | 49 | 10 | 83 | 18 |
| Shisa5 | ENSMUSG00000025647 | 705 | 100 | 1028 | 56 | 749 | 29 | 862 | 118 |
| Shtn1 | ENSMUSG00000041362 | 111 | 11 | 174 | 21 | 133 | 10 | 158 | 33 |
| Siglece | ENSMUSG00000030474 | 31 | 11 | 61 | 22 | 29 | 7 | 49 | 8 |
| Sirpa | ENSMUSG00000037902 | 816 | 78 | 1138 | 211 | 857 | 29 | 986 | 159 |
| Ska1 | ENSMUSG00000036223 | 3 | 3 | 16 | 11 | 2 | 2 | 11 | 8 |
| Ska3 | ENSMUSG00000021965 | 7 | 5 | 20 | 4 | 7 | 1 | 19 | 6 |
| Skil | ENSMUSG00000027660 | 581 | 92 | 881 | 203 | 556 | 78 | 727 | 130 |
| Skp2 | ENSMUSG00000054115 | 26 | 2 | 53 | 9 | 32 | 9 | 43 | 12 |
| Sla | ENSMUSG00000022372 | 69 | 14 | 123 | 46 | 60 | 7 | 74 | 15 |
| Slamf7 | ENSMUSG00000038179 | 16 | 7 | 59 | 44 | 20 | 9 | 31 | 22 |
| Slbp | ENSMUSG00000004642 | 270 | 27 | 364 | 58 | 264 | 33 | 311 | 22 |
| Slc15a3 | ENSMUSG00000024737 | 89 | 19 | 150 | 23 | 90 | 18 | 130 | 28 |
| Slc20a1 | ENSMUSG00000027397 | 373 | 83 | 570 | 141 | 383 | 75 | 430 | 73 |
| Slc25a24 | ENSMUSG00000040322 | 184 | 16 | 278 | 63 | 197 | 22 | 246 | 35 |
| Slc25a45 | ENSMUSG00000024818 | 85 | 11 | 134 | 30 | 87 | 26 | 111 | 13 |
| Slc39a6 | ENSMUSG00000024270 | 135 | 18 | 213 | 64 | 147 | 13 | 187 | 32 |
| Slc7a8 | ENSMUSG00000022180 | 85 | 8 | 120 | 20 | 95 | 6 | 114 | 11 |
| Slc9a1 | ENSMUSG00000028854 | 394 | 28 | 509 | 56 | 454 | 36 | 435 | 54 |
| Slc9a9 | ENSMUSG00000031129 | 122 | 18 | 169 | 14 | 122 | 7 | 161 | 18 |
| Slco2a1 | ENSMUSG00000032548 | 79 | 12 | 158 | 76 | 109 | 32 | 151 | 52 |
| Slfn1 | ENSMUSG00000078763 | 6 | 3 | 30 | 22 | 7 | 4 | 14 | 5 |
| Slfn4 | ENSMUSG00000000204 | 4 | 3 | 37 | 45 | 3 | 3 | 7 | 6 |
| Slfn8 | ENSMUSG00000035208 | 92 | 15 | 162 | 48 | 81 | 9 | 133 | 41 |
| Smc4 | ENSMUSG00000034349 | 507 | 95 | 871 | 221 | 482 | 64 | 678 | 208 |
| Smg1 | ENSMUSG00000030655 | 1722 | 260 | 2176 | 77 | 1762 | 176 | 1909 | 93 |
| Snhg18 | ENSMUSG00000096956 | 105 | 9 | 186 | 61 | 111 | 26 | 157 | 23 |
| Snx18 | ENSMUSG00000042364 | 553 | 47 | 696 | 92 | 562 | 61 | 610 | 52 |
| Snx20 | ENSMUSG00000031662 | 19 | 11 | 47 | 15 | 23 | 2 | 34 | 9 |
| Snx5 | ENSMUSG00000027423 | 1439 | 125 | 1707 | 156 | 1541 | 86 | 1646 | 122 |
| Soat1 | ENSMUSG00000026600 | 139 | 29 | 253 | 90 | 127 | 33 | 216 | 79 |
| Sp3 | ENSMUSG00000027109 | 1028 | 55 | 1219 | 141 | 1088 | 81 | 1139 | 66 |
| Spi1 | ENSMUSG00000002111 | 75 | 11 | 144 | 49 | 95 | 22 | 131 | 8 |
| Spidr | ENSMUSG00000041974 | 67 | 16 | 111 | 18 | 68 | 12 | 82 | 12 |
| Spin4 | ENSMUSG00000071722 | 30 | 7 | 51 | 5 | 34 | 7 | 38 | 7 |
| Spn | ENSMUSG00000051457 | 31 | 12 | 75 | 26 | 34 | 10 | 54 | 21 |
| Spp1 | ENSMUSG00000029304 | 7 | 3 | 219 | 250 | 5 | 2 | 107 | 189 |
| Spred1 | ENSMUSG00000027351 | 771 | 71 | 991 | 172 | 810 | 79 | 960 | 76 |
| Sprr2a2 | ENSMUSG00000068893 | 5 | 4 | 45 | 45 | 7 | 3 | 19 | 6 |
| Spty2d1 | ENSMUSG00000049516 | 303 | 36 | 432 | 55 | 340 | 21 | 357 | 39 |
| Sqle | ENSMUSG00000022351 | 22 | 13 | 44 | 9 | 32 | 12 | 32 | 12 |
| Srgap1 | ENSMUSG00000020121 | 122 | 17 | 179 | 21 | 119 | 24 | 155 | 30 |
| Ssh1 | ENSMUSG00000042121 | 389 | 56 | 482 | 27 | 429 | 54 | 403 | 41 |
| St14 | ENSMUSG00000031995 | 4 | 1 | 21 | 10 | 6 | 4 | 18 | 19 |
| Star | ENSMUSG00000031574 | 18 | 11 | 140 | 152 | 32 | 27 | 93 | 89 |
| Stk26 | ENSMUSG00000031112 | 14 | 4 | 39 | 26 | 33 | 20 | 26 | 15 |
| Stk32c | ENSMUSG00000015981 | 20 | 5 | 41 | 5 | 22 | 5 | 29 | 4 |
| Stt3a | ENSMUSG00000032116 | 850 | 44 | 1211 | 182 | 933 | 75 | 948 | 77 |
| Stxbp2 | ENSMUSG00000004626 | 39 | 12 | 87 | 17 | 33 | 5 | 50 | 10 |

| | | | | | | | | | |
|---|---|---|---|---|---|---|---|---|---|
| Sulf2 | ENSMUSG00000006800 | 1067 | 166 | 1425 | 253 | 1164 | 107 | 1386 | 119 |
| Susd2 | ENSMUSG00000006342 | 39 | 13 | 86 | 24 | 45 | 10 | 49 | 10 |
| Susd5 | ENSMUSG00000086596 | 20 | 9 | 62 | 31 | 26 | 9 | 31 | 13 |
| Synpo | ENSMUSG00000043079 | 2231 | 119 | 2747 | 352 | 2295 | 170 | 2766 | 301 |
| Syt12 | ENSMUSG00000049303 | 54 | 21 | 128 | 28 | 50 | 24 | 89 | 45 |
| Tagln | ENSMUSG00000032085 | 483 | 160 | 931 | 216 | 458 | 122 | 542 | 128 |
| Taok3 | ENSMUSG00000061288 | 238 | 28 | 320 | 48 | 253 | 36 | 307 | 26 |
| Tbpl1 | ENSMUSG00000071359 | 277 | 18 | 347 | 9 | 269 | 16 | 294 | 34 |
| Tbx15 | ENSMUSG00000027868 | 14 | 5 | 36 | 21 | 18 | 11 | 36 | 10 |
| Tbxas1 | ENSMUSG00000029925 | 62 | 14 | 104 | 27 | 75 | 23 | 89 | 12 |
| Tcaf1 | ENSMUSG00000036667 | 298 | 59 | 423 | 88 | 292 | 16 | 368 | 39 |
| Tceal9 | ENSMUSG00000042712 | 276 | 42 | 454 | 184 | 290 | 28 | 356 | 21 |
| Tcirg1 | ENSMUSG00000001750 | 143 | 21 | 252 | 50 | 181 | 41 | 211 | 53 |
| Tent5a | ENSMUSG00000032265 | 352 | 77 | 521 | 110 | 348 | 40 | 469 | 106 |
| Tent5c | ENSMUSG00000044468 | 180 | 62 | 415 | 151 | 316 | 121 | 530 | 280 |
| Tep1 | ENSMUSG00000006281 | 281 | 10 | 425 | 52 | 275 | 19 | 343 | 44 |
| Tgfb1i1 | ENSMUSG00000030782 | 331 | 73 | 487 | 33 | 326 | 41 | 417 | 38 |
| Tgfb3 | ENSMUSG00000021253 | 358 | 51 | 819 | 586 | 360 | 30 | 577 | 163 |
| Tgfbr1 | ENSMUSG00000007613 | 473 | 61 | 783 | 295 | 536 | 62 | 608 | 144 |
| Tgfbr2 | ENSMUSG00000032440 | 1323 | 221 | 1904 | 229 | 1360 | 84 | 1637 | 48 |
| Tgif2 | ENSMUSG00000062175 | 15 | 4 | 33 | 9 | 22 | 11 | 27 | 10 |
| Themis2 | ENSMUSG00000037731 | 94 | 14 | 167 | 25 | 103 | 12 | 120 | 24 |
| Thoc1 | ENSMUSG00000024287 | 393 | 27 | 507 | 81 | 425 | 40 | 457 | 36 |
| Tifab | ENSMUSG00000049625 | 51 | 11 | 93 | 26 | 49 | 6 | 72 | 19 |
| Tlr1 | ENSMUSG00000044827 | 25 | 12 | 79 | 59 | 23 | 4 | 49 | 22 |
| Tlr2 | ENSMUSG00000027995 | 103 | 10 | 196 | 69 | 107 | 15 | 152 | 23 |
| Tlr6 | ENSMUSG00000051498 | 15 | 4 | 36 | 10 | 18 | 9 | 18 | 10 |
| Tlr7 | ENSMUSG00000044583 | 100 | 10 | 218 | 51 | 119 | 44 | 182 | 57 |
| Tm6sf1 | ENSMUSG00000038623 | 189 | 36 | 283 | 38 | 193 | 35 | 228 | 22 |
| Tmem119 | ENSMUSG00000054675 | 65 | 21 | 180 | 126 | 65 | 6 | 118 | 38 |
| Tmem132a | ENSMUSG00000024736 | 173 | 25 | 238 | 29 | 190 | 15 | 225 | 33 |
| Tmem165 | ENSMUSG00000029234 | 367 | 42 | 504 | 119 | 398 | 39 | 467 | 29 |
| Tmem184c | ENSMUSG00000031617 | 277 | 31 | 380 | 62 | 313 | 36 | 352 | 23 |
| Tmem198b | ENSMUSG00000047090 | 101 | 18 | 156 | 32 | 99 | 12 | 129 | 16 |
| Tmem273 | ENSMUSG00000041707 | 12 | 5 | 33 | 15 | 17 | 6 | 24 | 6 |
| Tmem45a | ENSMUSG00000022754 | 60 | 25 | 158 | 66 | 68 | 11 | 112 | 16 |
| Tmem51 | ENSMUSG00000040616 | 54 | 4 | 82 | 7 | 68 | 8 | 69 | 12 |
| Tnfrsf1b | ENSMUSG00000028599 | 240 | 51 | 378 | 73 | 263 | 41 | 340 | 52 |
| Tnnt3 | ENSMUSG00000061723 | 6 | 2 | 29 | 39 | 5 | 3 | 17 | 17 |
| Tns3 | ENSMUSG00000020422 | 616 | 54 | 885 | 194 | 633 | 54 | 721 | 71 |
| Tor3a | ENSMUSG00000060519 | 274 | 45 | 395 | 49 | 295 | 22 | 340 | 25 |
| Tram2 | ENSMUSG00000041779 | 231 | 36 | 333 | 42 | 225 | 59 | 272 | 55 |
| Trf | ENSMUSG00000032554 | 860 | 217 | 1129 | 103 | 862 | 83 | 1027 | 78 |
| Trim30a | ENSMUSG00000030921 | 325 | 27 | 481 | 93 | 321 | 43 | 427 | 50 |
| Trip13 | ENSMUSG00000021569 | 9 | 5 | 30 | 12 | 10 | 2 | 26 | 8 |
| Trp53i11 | ENSMUSG00000068735 | 646 | 173 | 926 | 105 | 688 | 75 | 760 | 211 |
| Tshz3 | ENSMUSG00000021217 | 83 | 14 | 129 | 7 | 86 | 15 | 111 | 16 |
| Tsku | ENSMUSG00000049580 | 67 | 12 | 104 | 26 | 53 | 7 | 91 | 27 |
| Tspan18 | ENSMUSG00000027217 | 290 | 49 | 403 | 41 | 317 | 28 | 366 | 89 |
| Tspo | ENSMUSG00000041736 | 297 | 21 | 484 | 201 | 308 | 28 | 397 | 63 |
| Ttyh3 | ENSMUSG00000036565 | 273 | 38 | 397 | 63 | 320 | 15 | 336 | 32 |
| Tubb2b | ENSMUSG00000045136 | 49 | 15 | 127 | 28 | 62 | 34 | 105 | 43 |
| Txndc5 | ENSMUSG00000038991 | 889 | 51 | 1133 | 196 | 889 | 39 | 1068 | 85 |
| Ube2l6 | ENSMUSG00000027078 | 317 | 72 | 521 | 98 | 342 | 63 | 524 | 145 |
| Uchl1 | ENSMUSG00000029223 | 115 | 13 | 232 | 135 | 124 | 16 | 194 | 68 |
| Ugdh | ENSMUSG00000029201 | 332 | 33 | 438 | 36 | 370 | 92 | 419 | 88 |
| Upp1 | ENSMUSG00000020407 | 78 | 13 | 127 | 37 | 87 | 15 | 115 | 29 |
| Usp18 | ENSMUSG00000030107 | 82 | 19 | 133 | 27 | 74 | 13 | 106 | 20 |
| Usp6nl | ENSMUSG00000039046 | 217 | 31 | 297 | 27 | 260 | 46 | 269 | 17 |
| Vash1 | ENSMUSG00000021256 | 340 | 36 | 487 | 83 | 322 | 15 | 460 | 115 |
| Vasn | ENSMUSG00000039646 | 131 | 17 | 195 | 17 | 135 | 19 | 160 | 13 |
| Vasp | ENSMUSG00000030403 | 602 | 62 | 764 | 80 | 624 | 92 | 708 | 80 |
| Vgll3 | ENSMUSG00000091243 | 127 | 49 | 249 | 84 | 167 | 31 | 215 | 72 |
| Vsir | ENSMUSG00000020101 | 544 | 130 | 797 | 85 | 643 | 20 | 694 | 88 |

| | | | | | | | | | |
|---|---|---|---|---|---|---|---|---|---|
| Was | ENSMUSG00000031165 | 34 | 9 | 71 | 17 | 40 | 6 | 48 | 12 |
| Wipf1 | ENSMUSG00000075284 | 572 | 54 | 783 | 47 | 562 | 70 | 635 | 43 |
| Wisp1 | ENSMUSG00000005124 | 27 | 8 | 174 | 241 | 30 | 12 | 82 | 66 |
| Wnt9b | ENSMUSG00000018486 | 9 | 5 | 34 | 16 | 8 | 4 | 21 | 10 |
| Wsb1 | ENSMUSG00000017677 | 413 | 65 | 588 | 117 | 430 | 59 | 499 | 86 |
| Wwtr1 | ENSMUSG00000027803 | 1788 | 70 | 2201 | 189 | 1899 | 86 | 2118 | 63 |
| Xaf1 | ENSMUSG00000040483 | 194 | 44 | 280 | 32 | 185 | 32 | 215 | 10 |
| Xcr1 | ENSMUSG00000060509 | 4 | 4 | 16 | 13 | 3 | 2 | 5 | 4 |
| Yes1 | ENSMUSG00000014932 | 493 | 48 | 631 | 79 | 517 | 38 | 549 | 53 |
| Yipf5 | ENSMUSG00000024487 | 306 | 19 | 418 | 92 | 321 | 20 | 373 | 55 |
| Ywhaq | ENSMUSG00000076432 | 1324 | 72 | 1556 | 97 | 1256 | 62 | 1410 | 136 |
| Zbp1 | ENSMUSG00000027514 | 37 | 19 | 92 | 33 | 37 | 4 | 63 | 13 |
| Zdhhc20 | ENSMUSG00000021969 | 297 | 35 | 512 | 168 | 404 | 125 | 438 | 107 |
| Zfas1 | ENSMUSG00000074578 | 115 | 9 | 200 | 96 | 116 | 17 | 147 | 30 |
| Zfp185 | ENSMUSG00000031351 | 15 | 3 | 69 | 100 | 28 | 22 | 34 | 12 |
| Zfp36l2 | ENSMUSG00000045817 | 834 | 75 | 1082 | 70 | 960 | 66 | 1042 | 116 |
| Zfp385b | ENSMUSG00000027016 | 105 | 21 | 153 | 38 | 106 | 21 | 148 | 29 |
| Zfp948 | ENSMUSG00000067931 | 153 | 16 | 226 | 55 | 139 | 21 | 197 | 20 |
| Zmat3 | ENSMUSG00000027663 | 267 | 23 | 355 | 37 | 287 | 4 | 337 | 50 |

**Supplementary Table S7.** RNASeq analysis of effects of angiotensin II (AngII) on mRNA expression in hearts from PKN2Het vs WT littermates: mRNAs significantly downregulated by AngII in WT hearts.

| Gene Symbol | Ensembl gene id | WT Vehicle | | WT AngII | | PKN2Het Vehicle | | PKN2Het AngII | |
|---|---|---|---|---|---|---|---|---|---|
| | | Mean | SD | Mean | SD | Mean | SD | Mean | SD |
| 0610040J01Rik | ENSMUSG00000060512 | 86 | 12 | 50 | 19 | 78 | 10 | 62 | 9 |
| 1600014C10Rik | ENSMUSG00000054676 | 741 | 15 | 595 | 54 | 669 | 41 | 652 | 77 |
| 1700123M08Rik | ENSMUSG00000085614 | 65 | 5 | 40 | 9 | 59 | 6 | 47 | 7 |
| 2010001K21Rik | ENSMUSG00000051606 | 55 | 21 | 18 | 7 | 54 | 24 | 32 | 15 |
| 2310015K22Rik | ENSMUSG00000101257 | 17 | 5 | 5 | 2 | 14 | 3 | 6 | 4 |
| 2700097O09Rik | ENSMUSG00000062198 | 195 | 23 | 141 | 15 | 172 | 25 | 159 | 20 |
| 2900097C17Rik | ENSMUSG00000102869 | 6799 | 384 | 5461 | 338 | 6885 | 478 | 6050 | 659 |
| 5830417I10Rik | ENSMUSG00000078684 | 1015 | 96 | 743 | 87 | 964 | 152 | 776 | 61 |
| A330023F24Rik | ENSMUSG00000096929 | 201 | 38 | 119 | 33 | 138 | 27 | 115 | 44 |
| Abcb10 | ENSMUSG00000031974 | 999 | 42 | 743 | 124 | 1019 | 76 | 871 | 93 |
| Abcb8 | ENSMUSG00000028973 | 1023 | 86 | 729 | 145 | 1062 | 134 | 871 | 63 |
| Abcd3 | ENSMUSG00000028127 | 2749 | 274 | 2096 | 283 | 2698 | 193 | 2334 | 334 |
| Abhd10 | ENSMUSG00000033157 | 265 | 34 | 203 | 14 | 258 | 14 | 237 | 14 |
| AC131339.2 | ENSMUSG00000116656 | 8254 | 1201 | 5715 | 1421 | 7792 | 541 | 6468 | 972 |
| Acaa2 | ENSMUSG00000036880 | 13252 | 1380 | 8259 | 2391 | 12324 | 1059 | 9258 | 1215 |
| Acacb | ENSMUSG00000042010 | 7443 | 571 | 5153 | 1077 | 7777 | 973 | 6190 | 276 |
| Acad12 | ENSMUSG00000042647 | 1869 | 179 | 1262 | 208 | 1828 | 141 | 1493 | 156 |
| Acad8 | ENSMUSG00000031969 | 768 | 60 | 549 | 63 | 690 | 83 | 672 | 41 |
| Acadm | ENSMUSG00000062908 | 23340 | 1838 | 16479 | 3425 | 22611 | 922 | 17864 | 1871 |
| Acads | ENSMUSG00000029545 | 2580 | 67 | 1841 | 326 | 2630 | 310 | 2033 | 223 |
| Acadsb | ENSMUSG00000030861 | 2976 | 193 | 2400 | 192 | 2851 | 242 | 2623 | 299 |
| Acadvl | ENSMUSG00000018574 | 16703 | 1281 | 11802 | 3040 | 16035 | 839 | 13194 | 640 |
| Acat1 | ENSMUSG00000032047 | 11595 | 1306 | 8601 | 2045 | 11394 | 1310 | 9863 | 770 |
| Aco2 | ENSMUSG00000022477 | 41047 | 2512 | 30674 | 4778 | 41075 | 2487 | 35785 | 2269 |
| Acot2 | ENSMUSG00000021226 | 1314 | 144 | 912 | 240 | 1349 | 225 | 975 | 53 |
| Acot7 | ENSMUSG00000028937 | 819 | 45 | 640 | 57 | 804 | 63 | 691 | 52 |
| Acox1 | ENSMUSG00000020777 | 5964 | 375 | 4807 | 434 | 5983 | 629 | 5289 | 513 |
| Acp6 | ENSMUSG00000028093 | 375 | 19 | 285 | 28 | 360 | 30 | 302 | 31 |
| Acsl1 | ENSMUSG00000018796 | 21401 | 1598 | 15516 | 3710 | 21505 | 1628 | 18159 | 1993 |
| Acsl6 | ENSMUSG00000020333 | 322 | 47 | 199 | 38 | 318 | 50 | 264 | 61 |
| Acsm5 | ENSMUSG00000030972 | 83 | 17 | 31 | 11 | 61 | 9 | 36 | 8 |
| Acy3 | ENSMUSG00000024866 | 300 | 15 | 192 | 35 | 310 | 25 | 238 | 34 |
| Adamts7 | ENSMUSG00000032363 | 320 | 21 | 222 | 35 | 323 | 29 | 246 | 59 |
| Adck1 | ENSMUSG00000021044 | 486 | 19 | 381 | 49 | 484 | 24 | 447 | 35 |
| Adcy1 | ENSMUSG00000020431 | 128 | 33 | 66 | 23 | 142 | 11 | 108 | 33 |
| Adhfe1 | ENSMUSG00000025911 | 1463 | 160 | 926 | 241 | 1480 | 91 | 1152 | 46 |
| Adipor2 | ENSMUSG00000030168 | 2471 | 47 | 2151 | 151 | 2493 | 164 | 2345 | 116 |
| Adra1b | ENSMUSG00000050541 | 489 | 73 | 272 | 64 | 473 | 98 | 320 | 63 |
| Afg1l | ENSMUSG00000038302 | 1113 | 135 | 844 | 176 | 1027 | 95 | 958 | 51 |
| Afg3l1 | ENSMUSG00000031967 | 1302 | 67 | 991 | 162 | 1222 | 123 | 1128 | 81 |
| Afg3l2 | ENSMUSG00000024527 | 3400 | 269 | 2686 | 386 | 3367 | 188 | 3160 | 98 |
| Agl | ENSMUSG00000033400 | 6620 | 574 | 5262 | 429 | 6615 | 587 | 5784 | 471 |
| Agtpbp1 | ENSMUSG00000021557 | 3401 | 239 | 2676 | 267 | 3243 | 152 | 2747 | 370 |
| Agtr1a | ENSMUSG00000049115 | 735 | 120 | 525 | 56 | 733 | 94 | 602 | 75 |
| AI464131 | ENSMUSG00000046312 | 274 | 30 | 170 | 37 | 300 | 45 | 212 | 57 |
| Ak1 | ENSMUSG00000026817 | 8786 | 736 | 6676 | 1141 | 8306 | 245 | 7478 | 570 |
| Ak3 | ENSMUSG00000024782 | 2865 | 206 | 2428 | 194 | 2737 | 50 | 2646 | 143 |
| Akap1 | ENSMUSG00000018428 | 3302 | 350 | 2545 | 516 | 3554 | 412 | 3012 | 130 |
| Akr1b3 | ENSMUSG00000001642 | 4099 | 333 | 3107 | 569 | 4024 | 229 | 3471 | 308 |
| Akr1e1 | ENSMUSG00000045410 | 600 | 74 | 471 | 42 | 501 | 55 | 559 | 71 |
| Akr7a5 | ENSMUSG00000028743 | 466 | 29 | 337 | 42 | 459 | 49 | 383 | 48 |
| Akt2 | ENSMUSG00000004056 | 3610 | 431 | 2539 | 315 | 3460 | 299 | 2896 | 149 |
| Aktip | ENSMUSG00000031667 | 1230 | 73 | 958 | 87 | 1211 | 51 | 1043 | 104 |
| Aldh5a1 | ENSMUSG00000035936 | 1096 | 126 | 715 | 113 | 1087 | 105 | 893 | 121 |
| Aldoa | ENSMUSG00000030695 | 35858 | 1289 | 30804 | 2263 | 36705 | 2352 | 34386 | 2290 |

| | | | | | | | | | |
|---|---|---|---|---|---|---|---|---|---|
| Alkbh5 | ENSMUSG00000042650 | 2176 | 139 | 1835 | 156 | 2250 | 270 | 1962 | 35 |
| Alkbh7 | ENSMUSG00000002661 | 389 | 36 | 284 | 50 | 398 | 51 | 341 | 39 |
| Amd1 | ENSMUSG00000075232 | 2921 | 695 | 1957 | 248 | 2352 | 376 | 2046 | 448 |
| Anapc13 | ENSMUSG00000035048 | 991 | 52 | 835 | 73 | 936 | 44 | 898 | 50 |
| Ank | ENSMUSG00000022265 | 4065 | 84 | 3017 | 442 | 4145 | 294 | 3446 | 202 |
| Ank2 | ENSMUSG00000032826 | 3236 | 377 | 2426 | 262 | 3245 | 406 | 2934 | 430 |
| Anks1 | ENSMUSG00000024219 | 1418 | 123 | 1011 | 201 | 1347 | 177 | 1118 | 141 |
| Anxa11 | ENSMUSG00000021866 | 1492 | 129 | 1141 | 185 | 1489 | 130 | 1259 | 107 |
| Apba3 | ENSMUSG00000004931 | 676 | 58 | 507 | 64 | 671 | 72 | 610 | 77 |
| Apobec2 | ENSMUSG00000040694 | 3232 | 195 | 2680 | 271 | 3255 | 84 | 2930 | 192 |
| Arfgap2 | ENSMUSG00000027255 | 1072 | 50 | 921 | 18 | 1095 | 72 | 979 | 38 |
| Arhgap26 | ENSMUSG00000036452 | 1156 | 166 | 889 | 88 | 1003 | 90 | 905 | 186 |
| Arhgef17 | ENSMUSG00000032875 | 1622 | 105 | 1309 | 145 | 1648 | 171 | 1396 | 65 |
| Arhgef19 | ENSMUSG00000028919 | 631 | 130 | 395 | 97 | 630 | 134 | 502 | 132 |
| Arl2 | ENSMUSG00000024944 | 429 | 20 | 345 | 25 | 432 | 32 | 371 | 18 |
| Armc2 | ENSMUSG00000071324 | 597 | 48 | 445 | 104 | 615 | 73 | 517 | 100 |
| Art3 | ENSMUSG00000034842 | 4846 | 519 | 3712 | 598 | 4603 | 497 | 4111 | 546 |
| Art5 | ENSMUSG00000070424 | 186 | 40 | 126 | 25 | 137 | 17 | 109 | 25 |
| As3mt | ENSMUSG00000003559 | 1028 | 40 | 757 | 113 | 983 | 51 | 838 | 64 |
| Asb18 | ENSMUSG00000067081 | 499 | 48 | 355 | 63 | 568 | 74 | 485 | 101 |
| Asb8 | ENSMUSG00000048175 | 1440 | 34 | 1106 | 130 | 1405 | 108 | 1243 | 39 |
| Atp5a1 | ENSMUSG00000025428 | 73890 | 5007 | 52519 | 8597 | 73310 | 4969 | 61155 | 3605 |
| Atp5b | ENSMUSG00000025393 | 81983 | 4027 | 60525 | 7907 | 80175 | 3926 | 68956 | 5579 |
| Atp5d | ENSMUSG00000003072 | 5810 | 313 | 4470 | 769 | 5730 | 539 | 5074 | 535 |
| Atp5e | ENSMUSG00000016252 | 4682 | 460 | 3624 | 408 | 4707 | 229 | 4130 | 508 |
| Atp5g3 | ENSMUSG00000018770 | 18004 | 1331 | 13448 | 2291 | 18271 | 996 | 15327 | 1271 |
| Atp5o | ENSMUSG00000022956 | 14204 | 841 | 11103 | 1882 | 13615 | 536 | 12332 | 665 |
| Atrip | ENSMUSG00000025646 | 240 | 48 | 178 | 13 | 214 | 7 | 169 | 22 |
| Auh | ENSMUSG00000021460 | 1320 | 84 | 937 | 128 | 1302 | 50 | 1114 | 103 |
| B4gat1 | ENSMUSG00000047379 | 686 | 31 | 550 | 51 | 714 | 60 | 593 | 58 |
| Banf2os | ENSMUSG00000086384 | 37 | 10 | 18 | 9 | 34 | 8 | 23 | 6 |
| Bap1 | ENSMUSG00000021901 | 906 | 50 | 726 | 88 | 873 | 45 | 830 | 87 |
| BB218582 | ENSMUSG00000085218 | 104 | 20 | 66 | 19 | 93 | 12 | 74 | 18 |
| BC025920 | ENSMUSG00000074862 | 63 | 12 | 38 | 10 | 58 | 6 | 40 | 6 |
| Bcas2 | ENSMUSG00000005687 | 603 | 47 | 499 | 33 | 607 | 16 | 526 | 34 |
| Bcat2 | ENSMUSG00000030826 | 1208 | 96 | 917 | 126 | 1160 | 104 | 1012 | 77 |
| Bcl2l13 | ENSMUSG00000009112 | 1949 | 142 | 1520 | 239 | 2093 | 293 | 1695 | 182 |
| Bsg | ENSMUSG00000023175 | 18883 | 1349 | 15208 | 1807 | 18987 | 778 | 17057 | 1251 |
| Btbd2 | ENSMUSG00000003344 | 522 | 41 | 395 | 30 | 524 | 83 | 414 | 33 |
| Bzw2 | ENSMUSG00000020547 | 2728 | 225 | 2062 | 260 | 2680 | 103 | 2352 | 189 |
| C030006K11Rik | ENSMUSG00000116138 | 646 | 57 | 430 | 81 | 638 | 91 | 512 | 103 |
| C530005A16Rik | ENSMUSG00000085408 | 50 | 10 | 25 | 6 | 47 | 2 | 34 | 5 |
| Cacfd1 | ENSMUSG00000015488 | 733 | 54 | 563 | 73 | 706 | 29 | 560 | 60 |
| Cacnb2 | ENSMUSG00000057914 | 1546 | 266 | 1014 | 149 | 1369 | 192 | 1040 | 154 |
| Cacng6 | ENSMUSG00000078815 | 34 | 9 | 13 | 6 | 28 | 11 | 13 | 6 |
| Cadm4 | ENSMUSG00000054793 | 295 | 26 | 180 | 39 | 288 | 32 | 216 | 22 |
| Calr3 | ENSMUSG00000019732 | 237 | 30 | 185 | 7 | 218 | 18 | 193 | 10 |
| Cars2 | ENSMUSG00000056228 | 540 | 40 | 408 | 58 | 563 | 48 | 458 | 53 |
| Ccdc85c | ENSMUSG00000084883 | 758 | 52 | 536 | 118 | 780 | 117 | 613 | 80 |
| Ccl11 | ENSMUSG00000020676 | 47 | 16 | 18 | 6 | 28 | 8 | 22 | 12 |
| Cd59a | ENSMUSG00000032679 | 1574 | 127 | 1303 | 120 | 1547 | 105 | 1280 | 69 |
| Cd99l2 | ENSMUSG00000035776 | 1839 | 81 | 1579 | 95 | 1863 | 109 | 1697 | 59 |
| Cdh2 | ENSMUSG00000024304 | 7013 | 593 | 5762 | 276 | 7154 | 806 | 6155 | 511 |
| Cdip1 | ENSMUSG00000004071 | 1222 | 78 | 994 | 157 | 1254 | 99 | 1121 | 69 |
| Cdkl5 | ENSMUSG00000031292 | 275 | 46 | 190 | 15 | 246 | 19 | 225 | 37 |
| Cdkn1c | ENSMUSG00000037664 | 381 | 66 | 236 | 40 | 329 | 26 | 256 | 83 |
| Cds2 | ENSMUSG00000058793 | 4266 | 154 | 3435 | 371 | 4238 | 261 | 3575 | 214 |
| Cep128 | ENSMUSG00000061533 | 288 | 24 | 199 | 59 | 257 | 34 | 220 | 53 |
| Cep63 | ENSMUSG00000032534 | 767 | 47 | 629 | 48 | 738 | 59 | 684 | 37 |
| Ces1d | ENSMUSG00000056973 | 2184 | 133 | 1205 | 405 | 1975 | 180 | 1372 | 236 |
| Chchd10 | ENSMUSG00000049422 | 5597 | 303 | 4264 | 732 | 5732 | 395 | 4864 | 565 |
| Chchd2 | ENSMUSG00000070493 | 5866 | 649 | 4709 | 645 | 5829 | 370 | 5166 | 564 |
| Chrm2 | ENSMUSG00000045613 | 2913 | 381 | 2078 | 129 | 2822 | 350 | 2290 | 539 |
| Chrna2 | ENSMUSG00000022041 | 51 | 20 | 23 | 5 | 42 | 10 | 23 | 6 |

| | | | | | | | | | |
|---|---|---|---|---|---|---|---|---|---|
| Cirbp | ENSMUSG00000045193 | 506 | 100 | 381 | 38 | 528 | 31 | 425 | 70 |
| Cisd1 | ENSMUSG00000037710 | 2524 | 194 | 2038 | 264 | 2499 | 82 | 2259 | 149 |
| Clcn3 | ENSMUSG00000004319 | 1219 | 147 | 945 | 108 | 1163 | 87 | 993 | 110 |
| Clpp | ENSMUSG00000002660 | 608 | 48 | 457 | 72 | 598 | 7 | 499 | 95 |
| Clstn1 | ENSMUSG00000039953 | 1769 | 192 | 1482 | 75 | 1642 | 58 | 1584 | 93 |
| Cluh | ENSMUSG00000020741 | 5745 | 385 | 4121 | 818 | 6093 | 799 | 5032 | 263 |
| Cmbl | ENSMUSG00000022235 | 621 | 59 | 470 | 81 | 594 | 54 | 495 | 52 |
| Cobll1 | ENSMUSG00000034903 | 1935 | 295 | 1391 | 206 | 1921 | 162 | 1717 | 245 |
| Cog7 | ENSMUSG00000034951 | 461 | 24 | 365 | 35 | 450 | 38 | 403 | 33 |
| Colq | ENSMUSG00000057606 | 238 | 48 | 142 | 30 | 274 | 25 | 195 | 33 |
| Coq10a | ENSMUSG00000039914 | 3593 | 171 | 2600 | 450 | 3601 | 150 | 3083 | 271 |
| Coq2 | ENSMUSG00000029319 | 1358 | 71 | 1066 | 109 | 1353 | 65 | 1154 | 62 |
| Coq7 | ENSMUSG00000030652 | 1122 | 100 | 786 | 155 | 1106 | 99 | 884 | 88 |
| Coq9 | ENSMUSG00000031782 | 6190 | 445 | 4627 | 906 | 6154 | 410 | 5544 | 503 |
| Corin | ENSMUSG00000005220 | 3746 | 79 | 2720 | 217 | 3571 | 374 | 2889 | 594 |
| Cox4i1 | ENSMUSG00000031818 | 22284 | 812 | 17754 | 1611 | 22070 | 1141 | 19696 | 1055 |
| Cox5a | ENSMUSG00000000088 | 11595 | 608 | 8951 | 1587 | 11127 | 414 | 10060 | 740 |
| Cox5b | ENSMUSG00000061518 | 10424 | 672 | 8385 | 1343 | 10221 | 301 | 9039 | 577 |
| Cox7a1 | ENSMUSG00000074218 | 8954 | 697 | 6119 | 1540 | 8610 | 451 | 7040 | 827 |
| Cox8b | ENSMUSG00000025488 | 6504 | 621 | 4961 | 1023 | 6394 | 477 | 5556 | 903 |
| Cpt2 | ENSMUSG00000028607 | 3334 | 182 | 2311 | 520 | 3272 | 298 | 2688 | 241 |
| Crat | ENSMUSG00000026853 | 8544 | 365 | 5905 | 1336 | 8462 | 672 | 6705 | 567 |
| Cs | ENSMUSG00000005683 | 24899 | 2214 | 17482 | 3393 | 24790 | 1691 | 20831 | 1885 |
| Ctnna1 | ENSMUSG00000037815 | 10146 | 341 | 7973 | 731 | 9700 | 499 | 8679 | 771 |
| Ctsf | ENSMUSG00000083282 | 632 | 56 | 508 | 41 | 584 | 26 | 503 | 28 |
| Cul4a | ENSMUSG00000031446 | 2213 | 84 | 1944 | 75 | 2181 | 132 | 2162 | 67 |
| Cuta | ENSMUSG00000024194 | 357 | 31 | 286 | 29 | 351 | 28 | 325 | 26 |
| Cux2 | ENSMUSG00000042589 | 305 | 54 | 194 | 43 | 272 | 63 | 232 | 37 |
| Cxadr | ENSMUSG00000022865 | 914 | 145 | 688 | 47 | 929 | 123 | 924 | 91 |
| Cyb5d2 | ENSMUSG00000057778 | 378 | 41 | 287 | 32 | 380 | 40 | 322 | 22 |
| Cyfip2 | ENSMUSG00000020340 | 4903 | 247 | 3825 | 494 | 5110 | 622 | 4425 | 316 |
| Cyhr1 | ENSMUSG00000053929 | 1686 | 63 | 1426 | 122 | 1706 | 105 | 1462 | 118 |
| Cyp1a1 | ENSMUSG00000032315 | 14 | 10 | 3 | 1 | 10 | 6 | 3 | 1 |
| Cyth1 | ENSMUSG00000017132 | 1083 | 120 | 785 | 146 | 1045 | 114 | 943 | 140 |
| D17H6S53E | ENSMUSG00000043311 | 205 | 17 | 148 | 28 | 206 | 24 | 200 | 19 |
| D2hgdh | ENSMUSG00000073609 | 648 | 30 | 491 | 28 | 640 | 53 | 550 | 62 |
| D5Ertd579e | ENSMUSG00000029190 | 3086 | 284 | 2426 | 238 | 2994 | 183 | 2459 | 146 |
| Dap3 | ENSMUSG00000068921 | 1533 | 99 | 1172 | 166 | 1441 | 33 | 1291 | 61 |
| Ddt | ENSMUSG00000001666 | 477 | 14 | 371 | 59 | 468 | 31 | 410 | 47 |
| Decr1 | ENSMUSG00000028223 | 8479 | 863 | 5924 | 1566 | 7835 | 344 | 6512 | 520 |
| Dele1 | ENSMUSG00000024442 | 2437 | 95 | 1683 | 363 | 2478 | 270 | 2004 | 115 |
| Dgat2 | ENSMUSG00000030747 | 4953 | 461 | 3276 | 860 | 4835 | 519 | 3984 | 545 |
| Dhodh | ENSMUSG00000031730 | 247 | 18 | 159 | 24 | 232 | 20 | 214 | 23 |
| Dhrs11 | ENSMUSG00000034449 | 676 | 56 | 455 | 62 | 690 | 48 | 551 | 73 |
| Dhrs4 | ENSMUSG00000022210 | 1099 | 57 | 894 | 97 | 1070 | 35 | 930 | 39 |
| Diablo | ENSMUSG00000029433 | 1178 | 83 | 980 | 91 | 1146 | 35 | 1105 | 48 |
| Dip2c | ENSMUSG00000048264 | 1657 | 88 | 1255 | 180 | 1673 | 186 | 1363 | 98 |
| Dirc2 | ENSMUSG00000022848 | 917 | 63 | 720 | 74 | 897 | 73 | 778 | 36 |
| Dis3l | ENSMUSG00000032396 | 666 | 19 | 538 | 31 | 657 | 38 | 594 | 19 |
| Dlst | ENSMUSG00000004789 | 11654 | 750 | 8617 | 1895 | 11270 | 555 | 10153 | 788 |
| Dmpk | ENSMUSG00000030409 | 5934 | 219 | 4888 | 565 | 5867 | 505 | 5423 | 498 |
| Dnaaf3 | ENSMUSG00000055809 | 112 | 12 | 66 | 8 | 98 | 18 | 67 | 9 |
| Dnajb2 | ENSMUSG00000026203 | 1197 | 57 | 1005 | 70 | 1211 | 30 | 1066 | 80 |
| Dnajb9 | ENSMUSG00000014905 | 945 | 82 | 775 | 43 | 973 | 79 | 828 | 109 |
| Dnajc28 | ENSMUSG00000039763 | 1292 | 140 | 949 | 125 | 1268 | 112 | 1025 | 90 |
| Doc2g | ENSMUSG00000024871 | 2878 | 347 | 2069 | 419 | 3127 | 217 | 2451 | 320 |
| Drosha | ENSMUSG00000022191 | 1389 | 86 | 1151 | 83 | 1380 | 106 | 1191 | 65 |
| Dsc2 | ENSMUSG00000024331 | 1098 | 132 | 758 | 132 | 1038 | 80 | 880 | 140 |
| Dusp18 | ENSMUSG00000047205 | 1193 | 123 | 695 | 186 | 1165 | 123 | 828 | 151 |
| Dusp23 | ENSMUSG00000026544 | 177 | 25 | 131 | 14 | 183 | 13 | 153 | 7 |
| Dym | ENSMUSG00000035765 | 1390 | 74 | 1135 | 99 | 1419 | 90 | 1253 | 76 |
| Dynll2 | ENSMUSG00000020483 | 8450 | 781 | 6147 | 789 | 8457 | 316 | 7374 | 749 |
| E2f6 | ENSMUSG00000057469 | 1947 | 96 | 1553 | 144 | 1933 | 82 | 1731 | 69 |
| Ech1 | ENSMUSG00000053898 | 18987 | 2037 | 11742 | 3859 | 18009 | 1245 | 12749 | 1792 |

| | | | | | | | | | |
|---|---|---|---|---|---|---|---|---|---|
| Echdc3 | ENSMUSG00000039063 | 427 | 19 | 300 | 49 | 428 | 30 | 350 | 42 |
| Echs1 | ENSMUSG00000025465 | 4828 | 268 | 3779 | 465 | 4896 | 284 | 4167 | 357 |
| Eci1 | ENSMUSG00000024132 | 3819 | 262 | 2675 | 691 | 3634 | 65 | 3027 | 306 |
| Ecpas | ENSMUSG00000050812 | 5520 | 139 | 4646 | 405 | 5619 | 641 | 4989 | 290 |
| Ecsit | ENSMUSG00000066839 | 1352 | 92 | 1050 | 152 | 1386 | 47 | 1218 | 79 |
| Eef1a2 | ENSMUSG00000016349 | 19816 | 1103 | 16353 | 1766 | 20590 | 904 | 18385 | 1605 |
| Eef1d | ENSMUSG00000055762 | 2212 | 97 | 1867 | 139 | 2077 | 103 | 2044 | 101 |
| Efcab2 | ENSMUSG00000026495 | 2803 | 355 | 2211 | 241 | 2798 | 209 | 2383 | 245 |
| Efnb3 | ENSMUSG00000003934 | 1231 | 119 | 643 | 238 | 1266 | 236 | 811 | 246 |
| Egflam | ENSMUSG00000042961 | 374 | 53 | 240 | 63 | 279 | 55 | 262 | 40 |
| Egln1 | ENSMUSG00000031987 | 9248 | 229 | 6578 | 1044 | 9122 | 656 | 7541 | 559 |
| Eid2b | ENSMUSG00000070705 | 191 | 16 | 132 | 14 | 196 | 26 | 154 | 37 |
| Eml2 | ENSMUSG00000040811 | 379 | 40 | 285 | 35 | 408 | 44 | 337 | 30 |
| Endog | ENSMUSG00000015337 | 455 | 64 | 312 | 72 | 475 | 50 | 367 | 54 |
| Eno3 | ENSMUSG00000060600 | 20868 | 2272 | 14909 | 3162 | 20388 | 959 | 17554 | 1469 |
| Enpp5 | ENSMUSG00000023960 | 482 | 40 | 367 | 31 | 452 | 24 | 401 | 59 |
| Entpd4b | ENSMUSG00000022066 | 1245 | 62 | 1009 | 104 | 1249 | 57 | 1115 | 37 |
| Ephx2 | ENSMUSG00000022040 | 5045 | 182 | 3683 | 568 | 4643 | 192 | 4079 | 372 |
| Epm2a | ENSMUSG00000055493 | 816 | 72 | 609 | 39 | 825 | 73 | 714 | 58 |
| Erc1 | ENSMUSG00000030172 | 1343 | 106 | 1011 | 106 | 1205 | 149 | 1073 | 45 |
| Esrrb | ENSMUSG00000021255 | 570 | 46 | 424 | 77 | 607 | 56 | 500 | 36 |
| Esrrg | ENSMUSG00000026610 | 1115 | 169 | 852 | 111 | 1188 | 192 | 986 | 138 |
| Etfa | ENSMUSG00000032314 | 11167 | 800 | 7804 | 1484 | 10775 | 533 | 8929 | 812 |
| Etfb | ENSMUSG00000004610 | 8827 | 431 | 6069 | 1637 | 8450 | 419 | 6927 | 725 |
| Etfdh | ENSMUSG00000027809 | 13588 | 811 | 9757 | 2314 | 12812 | 772 | 10703 | 771 |
| Extl1 | ENSMUSG00000028838 | 505 | 28 | 354 | 91 | 508 | 68 | 427 | 38 |
| Fam131a | ENSMUSG00000050821 | 452 | 30 | 325 | 65 | 445 | 59 | 370 | 46 |
| Fam20b | ENSMUSG00000033557 | 2462 | 33 | 2046 | 249 | 2441 | 169 | 2285 | 137 |
| Fam210a | ENSMUSG00000038121 | 5630 | 318 | 3910 | 672 | 5594 | 632 | 4548 | 152 |
| Farp2 | ENSMUSG00000034066 | 407 | 27 | 323 | 41 | 400 | 28 | 340 | 23 |
| Fastkd2 | ENSMUSG00000025962 | 921 | 102 | 730 | 94 | 816 | 39 | 814 | 51 |
| Fbp2 | ENSMUSG00000021456 | 422 | 72 | 243 | 83 | 363 | 23 | 250 | 9 |
| Fbxo21 | ENSMUSG00000032898 | 802 | 70 | 590 | 81 | 829 | 129 | 684 | 55 |
| Fbxo31 | ENSMUSG00000052934 | 1386 | 125 | 967 | 187 | 1389 | 148 | 1079 | 71 |
| Fbxo32 | ENSMUSG00000022358 | 4438 | 248 | 3382 | 496 | 4609 | 317 | 3689 | 462 |
| Fdft1 | ENSMUSG00000021273 | 812 | 58 | 545 | 97 | 811 | 49 | 633 | 82 |
| Fem1a | ENSMUSG00000043683 | 4314 | 280 | 3250 | 616 | 4473 | 492 | 3867 | 370 |
| Fh1 | ENSMUSG00000026526 | 7500 | 576 | 5659 | 1050 | 7018 | 180 | 6388 | 586 |
| Fhod3 | ENSMUSG00000034295 | 6968 | 897 | 4787 | 812 | 6949 | 852 | 5943 | 560 |
| Fign | ENSMUSG00000075324 | 264 | 50 | 152 | 48 | 282 | 77 | 222 | 62 |
| Fkbp4 | ENSMUSG00000030357 | 7033 | 609 | 4753 | 1205 | 6950 | 342 | 5630 | 654 |
| Flad1 | ENSMUSG00000042642 | 928 | 42 | 703 | 107 | 840 | 64 | 785 | 91 |
| Fmc1 | ENSMUSG00000019689 | 622 | 55 | 458 | 87 | 580 | 55 | 484 | 71 |
| Fn3k | ENSMUSG00000025175 | 262 | 26 | 173 | 19 | 252 | 25 | 208 | 27 |
| Fndc5 | ENSMUSG00000001334 | 7755 | 469 | 5429 | 1330 | 7876 | 453 | 6619 | 559 |
| Foxo3 | ENSMUSG00000048756 | 1906 | 138 | 1489 | 235 | 1807 | 100 | 1507 | 232 |
| Foxo4 | ENSMUSG00000042903 | 1026 | 58 | 818 | 81 | 1020 | 106 | 883 | 76 |
| Foxo6os | ENSMUSG00000084929 | 123 | 20 | 70 | 22 | 91 | 13 | 97 | 30 |
| Fsd2 | ENSMUSG00000038663 | 7461 | 469 | 5809 | 1017 | 7438 | 469 | 6588 | 359 |
| Fth1 | ENSMUSG00000024661 | 24620 | 1681 | 21525 | 749 | 25292 | 1474 | 22765 | 1350 |
| Fuz | ENSMUSG00000011658 | 154 | 17 | 113 | 4 | 143 | 29 | 114 | 25 |
| Fxr2 | ENSMUSG00000018765 | 2087 | 79 | 1668 | 103 | 2112 | 177 | 1949 | 154 |
| Gart | ENSMUSG00000022962 | 791 | 63 | 621 | 72 | 729 | 62 | 724 | 31 |
| Gcsh | ENSMUSG00000034424 | 1021 | 36 | 874 | 74 | 1000 | 34 | 954 | 42 |
| Gfm1 | ENSMUSG00000027774 | 4053 | 240 | 3035 | 442 | 3973 | 430 | 3372 | 259 |
| Gfm2 | ENSMUSG00000021666 | 1266 | 144 | 1001 | 118 | 1229 | 127 | 1128 | 100 |
| Gfra1 | ENSMUSG00000025089 | 261 | 28 | 179 | 47 | 276 | 25 | 195 | 40 |
| Ghitm | ENSMUSG00000041028 | 10888 | 1003 | 9249 | 687 | 10712 | 845 | 9689 | 126 |
| Glo1 | ENSMUSG00000024026 | 1952 | 124 | 1549 | 162 | 1830 | 76 | 1663 | 101 |
| Gm10644 | ENSMUSG00000074219 | 30 | 3 | 13 | 5 | 21 | 6 | 18 | 7 |
| Gm20619 | ENSMUSG00000093482 | 85 | 17 | 53 | 4 | 91 | 12 | 73 | 11 |
| Gm29170 | ENSMUSG00000100455 | 84 | 16 | 48 | 12 | 73 | 11 | 60 | 14 |
| Gm33543 | ENSMUSG00000110353 | 38 | 12 | 18 | 6 | 30 | 9 | 24 | 10 |
| Gm36827 | ENSMUSG00000112327 | 289 | 42 | 195 | 41 | 280 | 51 | 188 | 31 |

| | | | | | | | | | |
|---|---|---|---|---|---|---|---|---|---|
| Gm37829 | ENSMUSG00000104453 | 1373 | 82 | 1009 | 161 | 1372 | 115 | 1141 | 137 |
| Gm40604 | ENSMUSG00000112800 | 20 | 10 | 8 | 4 | 11 | 3 | 8 | 3 |
| Gm43672 | ENSMUSG00000106019 | 412 | 67 | 247 | 76 | 363 | 75 | 322 | 88 |
| Gm45012 | ENSMUSG00000109052 | 77 | 14 | 44 | 18 | 83 | 8 | 50 | 6 |
| Gm47547 | ENSMUSG00000114196 | 4410 | 804 | 3053 | 766 | 4397 | 589 | 3371 | 749 |
| Gm49083 | ENSMUSG00000115354 | 278 | 31 | 195 | 62 | 260 | 7 | 202 | 31 |
| Gm49130 | ENSMUSG00000115234 | 52 | 12 | 23 | 9 | 54 | 13 | 29 | 9 |
| Gm49477 | ENSMUSG00000116066 | 359 | 37 | 242 | 50 | 366 | 16 | 279 | 49 |
| Gm826 | ENSMUSG00000074623 | 78 | 4 | 53 | 10 | 73 | 9 | 60 | 9 |
| Gna12 | ENSMUSG00000000149 | 3048 | 223 | 2381 | 429 | 3083 | 284 | 2577 | 277 |
| Gnpat | ENSMUSG00000031985 | 4943 | 161 | 3883 | 434 | 4842 | 240 | 4332 | 164 |
| Got1 | ENSMUSG00000025190 | 12937 | 538 | 8755 | 1664 | 12760 | 639 | 10615 | 1253 |
| Got2 | ENSMUSG00000031672 | 15403 | 947 | 11510 | 1822 | 15607 | 1320 | 13207 | 1045 |
| Gpd2 | ENSMUSG00000026827 | 401 | 24 | 334 | 32 | 392 | 48 | 398 | 25 |
| Gpn1 | ENSMUSG00000064037 | 443 | 41 | 349 | 55 | 348 | 21 | 379 | 15 |
| Gpr155 | ENSMUSG00000041762 | 432 | 52 | 326 | 29 | 393 | 31 | 316 | 35 |
| Gpr22 | ENSMUSG00000044067 | 1368 | 315 | 717 | 244 | 1339 | 221 | 933 | 230 |
| Gpr27 | ENSMUSG00000072875 | 177 | 17 | 110 | 8 | 189 | 36 | 153 | 20 |
| Gpt | ENSMUSG00000022546 | 376 | 38 | 239 | 62 | 389 | 47 | 298 | 58 |
| Grb14 | ENSMUSG00000026888 | 3020 | 238 | 2239 | 419 | 3074 | 220 | 2499 | 294 |
| Gsta4 | ENSMUSG00000032348 | 1102 | 100 | 835 | 91 | 1063 | 51 | 979 | 106 |
| Gstk1 | ENSMUSG00000029864 | 1045 | 99 | 635 | 177 | 1034 | 85 | 737 | 172 |
| Gstm1 | ENSMUSG00000058135 | 2900 | 228 | 2007 | 410 | 2765 | 233 | 2247 | 280 |
| Gstm7 | ENSMUSG00000004035 | 451 | 36 | 268 | 66 | 397 | 64 | 313 | 83 |
| Gstp1 | ENSMUSG00000060803 | 1530 | 60 | 1249 | 118 | 1564 | 76 | 1343 | 113 |
| Gstt1 | ENSMUSG00000001663 | 186 | 22 | 128 | 20 | 182 | 23 | 129 | 13 |
| Gypc | ENSMUSG00000090523 | 589 | 35 | 453 | 36 | 560 | 48 | 489 | 47 |
| Gys1 | ENSMUSG00000003865 | 2308 | 136 | 1765 | 214 | 2462 | 310 | 2076 | 150 |
| Gzmm | ENSMUSG00000054206 | 67 | 4 | 40 | 12 | 60 | 12 | 51 | 9 |
| H2-Ke6 | ENSMUSG00000073422 | 624 | 79 | 486 | 54 | 621 | 40 | 499 | 38 |
| H2afv | ENSMUSG00000041126 | 582 | 47 | 479 | 48 | 560 | 29 | 511 | 53 |
| Hadh | ENSMUSG00000027984 | 9800 | 579 | 6798 | 1554 | 9433 | 502 | 7758 | 928 |
| Hadhb | ENSMUSG00000059447 | 37787 | 3939 | 27391 | 5861 | 37795 | 3637 | 30089 | 2124 |
| Hbs1l | ENSMUSG00000019977 | 1596 | 64 | 1363 | 99 | 1582 | 55 | 1431 | 99 |
| Hcn4 | ENSMUSG00000032338 | 215 | 29 | 145 | 22 | 226 | 22 | 167 | 29 |
| Hdhd2 | ENSMUSG00000025421 | 925 | 39 | 718 | 100 | 905 | 51 | 782 | 86 |
| Heatr5b | ENSMUSG00000039414 | 1026 | 107 | 806 | 74 | 965 | 134 | 837 | 66 |
| Helt | ENSMUSG00000047171 | 27 | 9 | 8 | 2 | 21 | 5 | 16 | 10 |
| Herc3 | ENSMUSG00000029804 | 1105 | 188 | 784 | 144 | 1110 | 191 | 995 | 112 |
| Hibadh | ENSMUSG00000029776 | 4598 | 354 | 3506 | 554 | 4704 | 186 | 4030 | 384 |
| Hikeshi | ENSMUSG00000062797 | 431 | 18 | 351 | 14 | 437 | 49 | 395 | 29 |
| Hk2 | ENSMUSG00000000628 | 7061 | 664 | 5062 | 953 | 7161 | 711 | 6174 | 483 |
| Hlf | ENSMUSG00000003949 | 702 | 180 | 377 | 124 | 587 | 150 | 396 | 181 |
| Hmgcs2 | ENSMUSG00000027875 | 274 | 33 | 184 | 30 | 225 | 17 | 152 | 50 |
| Hnmt | ENSMUSG00000026986 | 222 | 26 | 145 | 18 | 233 | 31 | 181 | 38 |
| Hopx | ENSMUSG00000059325 | 3360 | 233 | 2164 | 627 | 3438 | 319 | 2591 | 663 |
| Hrc | ENSMUSG00000038239 | 19722 | 920 | 12511 | 2864 | 19104 | 720 | 15198 | 1393 |
| Hsd17b10 | ENSMUSG00000025260 | 1979 | 163 | 1487 | 242 | 1984 | 101 | 1624 | 232 |
| Hsdl2 | ENSMUSG00000028383 | 5524 | 757 | 4049 | 946 | 5605 | 677 | 4510 | 583 |
| Hspa5 | ENSMUSG00000026864 | 12991 | 1134 | 9421 | 1954 | 11342 | 633 | 10018 | 1044 |
| Hspa9 | ENSMUSG00000024359 | 16027 | 1269 | 12152 | 2141 | 15522 | 684 | 14164 | 919 |
| Hspd1 | ENSMUSG00000025980 | 8960 | 680 | 6784 | 1395 | 8344 | 426 | 7562 | 640 |
| Htra1 | ENSMUSG00000006205 | 1381 | 97 | 1023 | 118 | 1387 | 133 | 1174 | 135 |
| Iars2 | ENSMUSG00000026618 | 2409 | 99 | 1873 | 136 | 2398 | 100 | 2171 | 120 |
| Idh2 | ENSMUSG00000030541 | 25832 | 1566 | 18548 | 3665 | 26906 | 2049 | 21457 | 2197 |
| Idh3a | ENSMUSG00000032279 | 10211 | 1028 | 7707 | 867 | 10029 | 629 | 8753 | 799 |
| Idh3b | ENSMUSG00000027406 | 11455 | 1144 | 8621 | 1973 | 11227 | 459 | 9312 | 738 |
| Ids | ENSMUSG00000035847 | 844 | 113 | 686 | 22 | 705 | 81 | 678 | 78 |
| Idua | ENSMUSG00000033540 | 381 | 15 | 288 | 34 | 343 | 21 | 312 | 19 |
| Il10rb | ENSMUSG00000022969 | 2740 | 90 | 2327 | 186 | 2766 | 76 | 2592 | 101 |
| Immt | ENSMUSG00000052337 | 11137 | 723 | 8672 | 1433 | 10924 | 555 | 9873 | 547 |
| Imp3 | ENSMUSG00000032288 | 386 | 37 | 293 | 37 | 353 | 24 | 333 | 39 |
| Insyn1 | ENSMUSG00000066607 | 332 | 30 | 244 | 41 | 353 | 22 | 296 | 17 |
| Isca1 | ENSMUSG00000044792 | 5523 | 385 | 4556 | 545 | 5614 | 212 | 5191 | 315 |

| | | | | | | | | | |
|---|---|---|---|---|---|---|---|---|---|
| Iscu | ENSMUSG00000025825 | 1538 | 101 | 1296 | 84 | 1533 | 74 | 1368 | 132 |
| Isoc2a | ENSMUSG00000086784 | 619 | 51 | 458 | 90 | 619 | 73 | 511 | 50 |
| Kars | ENSMUSG00000031948 | 2130 | 77 | 1818 | 133 | 2087 | 64 | 1922 | 99 |
| Kcnb1 | ENSMUSG00000050556 | 1529 | 86 | 1149 | 198 | 1362 | 171 | 1226 | 75 |
| Kcnd3 | ENSMUSG00000040896 | 419 | 52 | 304 | 54 | 437 | 49 | 338 | 38 |
| Kcng2 | ENSMUSG00000059852 | 1131 | 114 | 738 | 153 | 1222 | 266 | 957 | 140 |
| Kcnip2 | ENSMUSG00000025221 | 2873 | 266 | 2038 | 321 | 2700 | 318 | 2226 | 310 |
| Kcnk3 | ENSMUSG00000049265 | 4069 | 294 | 2872 | 660 | 4507 | 692 | 3530 | 463 |
| Kctd9 | ENSMUSG00000034327 | 1736 | 135 | 1293 | 231 | 1770 | 104 | 1588 | 97 |
| Khdrbs3 | ENSMUSG00000022332 | 876 | 67 | 638 | 92 | 898 | 73 | 738 | 119 |
| Kif16b | ENSMUSG00000038844 | 1579 | 142 | 1284 | 141 | 1624 | 143 | 1458 | 52 |
| Kif1c | ENSMUSG00000020821 | 13040 | 401 | 10864 | 1290 | 13509 | 1540 | 11810 | 481 |
| Kif21a | ENSMUSG00000022629 | 911 | 67 | 703 | 90 | 899 | 40 | 823 | 111 |
| Klf9 | ENSMUSG00000033863 | 2073 | 327 | 1580 | 218 | 1929 | 244 | 1672 | 332 |
| Klhl21 | ENSMUSG00000073700 | 1596 | 96 | 1232 | 219 | 1629 | 106 | 1373 | 96 |
| Klhl33 | ENSMUSG00000090799 | 489 | 94 | 262 | 86 | 446 | 37 | 313 | 87 |
| Kpna6 | ENSMUSG00000003731 | 2025 | 137 | 1642 | 186 | 1978 | 88 | 1813 | 64 |
| Ky | ENSMUSG00000035606 | 445 | 33 | 251 | 85 | 458 | 49 | 367 | 57 |
| Kyat1 | ENSMUSG00000039648 | 379 | 27 | 294 | 42 | 380 | 16 | 314 | 20 |
| L2hgdh | ENSMUSG00000020988 | 1075 | 56 | 746 | 101 | 1069 | 51 | 876 | 104 |
| Lclat1 | ENSMUSG00000054469 | 3111 | 302 | 2399 | 376 | 3091 | 319 | 2588 | 234 |
| Ldhb | ENSMUSG00000030246 | 22959 | 1639 | 15955 | 3136 | 22470 | 933 | 18647 | 1684 |
| Letm1 | ENSMUSG00000005299 | 2120 | 191 | 1513 | 340 | 2063 | 206 | 1803 | 214 |
| Lias | ENSMUSG00000029199 | 931 | 93 | 751 | 90 | 903 | 34 | 834 | 53 |
| Limch1 | ENSMUSG00000037736 | 5052 | 422 | 3923 | 326 | 5049 | 205 | 4581 | 477 |
| Lingo3 | ENSMUSG00000051067 | 229 | 41 | 120 | 47 | 252 | 73 | 159 | 48 |
| Lmbrd1 | ENSMUSG00000073725 | 1143 | 53 | 960 | 76 | 1105 | 87 | 1035 | 39 |
| Lmo7 | ENSMUSG00000033060 | 8272 | 652 | 6224 | 773 | 7881 | 612 | 7367 | 993 |
| Lonp1 | ENSMUSG00000041168 | 2317 | 80 | 2025 | 163 | 2376 | 150 | 2252 | 87 |
| Lonrf2 | ENSMUSG00000048814 | 202 | 35 | 126 | 24 | 204 | 39 | 153 | 25 |
| Lpin1 | ENSMUSG00000020593 | 4444 | 676 | 2968 | 594 | 4769 | 532 | 3724 | 386 |
| Lpl | ENSMUSG00000015568 | 90429 | 5374 | 71420 | 8174 | 93258 | 8317 | 80525 | 6094 |
| Lrpprc | ENSMUSG00000024120 | 4237 | 451 | 2986 | 400 | 4181 | 326 | 3537 | 377 |
| Lrrc3b | ENSMUSG00000045201 | 506 | 44 | 333 | 72 | 477 | 50 | 367 | 53 |
| Lsm14b | ENSMUSG00000039108 | 972 | 35 | 742 | 85 | 914 | 50 | 896 | 76 |
| Lynx1 | ENSMUSG00000022594 | 5026 | 322 | 3722 | 716 | 5034 | 369 | 4308 | 264 |
| Maf1 | ENSMUSG00000022553 | 1093 | 59 | 891 | 57 | 1121 | 107 | 1002 | 47 |
| Magi3 | ENSMUSG00000052539 | 1616 | 181 | 1196 | 121 | 1556 | 176 | 1327 | 166 |
| Magt1 | ENSMUSG00000031232 | 1232 | 137 | 942 | 126 | 1216 | 90 | 994 | 120 |
| Malsu1 | ENSMUSG00000029815 | 281 | 28 | 208 | 15 | 260 | 23 | 236 | 30 |
| Map10 | ENSMUSG00000050930 | 149 | 21 | 98 | 15 | 151 | 17 | 112 | 10 |
| Map1lc3a | ENSMUSG00000027602 | 4603 | 222 | 3864 | 305 | 4713 | 378 | 4049 | 447 |
| Mapk8ip3 | ENSMUSG00000024163 | 1609 | 148 | 1286 | 127 | 1571 | 128 | 1366 | 102 |
| Mars2 | ENSMUSG00000046994 | 370 | 47 | 284 | 34 | 364 | 34 | 304 | 25 |
| Mccc1 | ENSMUSG00000027709 | 2112 | 94 | 1589 | 261 | 2026 | 95 | 1662 | 118 |
| Mdga1 | ENSMUSG00000043557 | 283 | 40 | 180 | 25 | 255 | 46 | 211 | 43 |
| Mdh2 | ENSMUSG00000019179 | 16876 | 703 | 12970 | 2129 | 16369 | 1034 | 14499 | 734 |
| Me1 | ENSMUSG00000032418 | 2846 | 130 | 2349 | 226 | 2697 | 138 | 2438 | 237 |
| Med12l | ENSMUSG00000056476 | 211 | 24 | 139 | 35 | 199 | 27 | 142 | 38 |
| Med9 | ENSMUSG00000061650 | 383 | 8 | 302 | 30 | 383 | 23 | 347 | 23 |
| Mettl7a1 | ENSMUSG00000054619 | 740 | 30 | 561 | 48 | 684 | 59 | 597 | 63 |
| Mfn1 | ENSMUSG00000027668 | 8217 | 628 | 6382 | 1129 | 8242 | 170 | 7059 | 753 |
| Mfn2 | ENSMUSG00000029020 | 15414 | 955 | 11184 | 2042 | 15810 | 1644 | 13241 | 705 |
| Mgme1 | ENSMUSG00000027424 | 350 | 28 | 255 | 46 | 337 | 12 | 267 | 30 |
| Mgrn1 | ENSMUSG00000022517 | 3397 | 191 | 2489 | 420 | 3480 | 322 | 2787 | 167 |
| Mhrt | ENSMUSG00000097652 | 413 | 27 | 294 | 60 | 414 | 38 | 328 | 38 |
| Miga1 | ENSMUSG00000054942 | 412 | 61 | 323 | 8 | 409 | 40 | 383 | 35 |
| Mipep | ENSMUSG00000021993 | 1364 | 63 | 983 | 167 | 1394 | 126 | 1103 | 71 |
| Mllt6 | ENSMUSG00000038437 | 1839 | 74 | 1515 | 116 | 1801 | 105 | 1533 | 96 |
| Mlxip | ENSMUSG00000038342 | 1090 | 98 | 876 | 108 | 1082 | 85 | 963 | 38 |
| Mmaa | ENSMUSG00000037022 | 603 | 23 | 460 | 70 | 572 | 47 | 517 | 37 |
| Mmadhc | ENSMUSG00000026766 | 1760 | 184 | 1400 | 199 | 1684 | 61 | 1526 | 84 |
| Mov10l1 | ENSMUSG00000015365 | 912 | 70 | 518 | 82 | 908 | 73 | 735 | 108 |
| Mpi | ENSMUSG00000032306 | 1513 | 105 | 1202 | 137 | 1517 | 24 | 1387 | 104 |

| | | | | | | | | | |
|---|---|---|---|---|---|---|---|---|---|
| Mpped2 | ENSMUSG00000016386 | 351 | 88 | 229 | 30 | 340 | 24 | 247 | 36 |
| Mpst | ENSMUSG00000071711 | 347 | 22 | 269 | 21 | 344 | 35 | 280 | 23 |
| Mpv17 | ENSMUSG00000107283 | 1098 | 32 | 873 | 90 | 1118 | 80 | 934 | 104 |
| Mrpl14 | ENSMUSG00000023939 | 682 | 54 | 516 | 77 | 739 | 35 | 586 | 87 |
| Mrpl16 | ENSMUSG00000024683 | 961 | 36 | 784 | 75 | 930 | 52 | 848 | 97 |
| Mrpl28 | ENSMUSG00000024181 | 1210 | 39 | 882 | 132 | 1171 | 81 | 1020 | 121 |
| Mrpl37 | ENSMUSG00000028622 | 1184 | 43 | 904 | 135 | 1161 | 34 | 1046 | 74 |
| Mrpl38 | ENSMUSG00000020775 | 704 | 31 | 534 | 72 | 674 | 41 | 611 | 58 |
| Mrpl39 | ENSMUSG00000022889 | 1365 | 91 | 1053 | 180 | 1335 | 40 | 1087 | 56 |
| Mrpl4 | ENSMUSG00000003299 | 1028 | 38 | 792 | 138 | 1029 | 93 | 915 | 137 |
| Mrpl45 | ENSMUSG00000018882 | 1187 | 97 | 837 | 153 | 1100 | 58 | 1043 | 54 |
| Mrps26 | ENSMUSG00000037740 | 355 | 39 | 258 | 24 | 351 | 18 | 326 | 26 |
| Mrps35 | ENSMUSG00000040112 | 1312 | 76 | 991 | 219 | 1236 | 59 | 1132 | 63 |
| Mrps6 | ENSMUSG00000039680 | 361 | 20 | 285 | 16 | 354 | 19 | 300 | 4 |
| Msrb2 | ENSMUSG00000023094 | 1067 | 66 | 775 | 115 | 1023 | 25 | 865 | 63 |
| Mtfp1 | ENSMUSG00000004748 | 1636 | 82 | 1103 | 322 | 1626 | 114 | 1320 | 90 |
| Mtfr1l | ENSMUSG00000046671 | 2563 | 65 | 2169 | 189 | 2622 | 122 | 2390 | 105 |
| Mtg2 | ENSMUSG00000039069 | 366 | 31 | 275 | 26 | 387 | 9 | 312 | 16 |
| Mtmr4 | ENSMUSG00000018401 | 533 | 95 | 415 | 52 | 508 | 47 | 429 | 38 |
| Mto1 | ENSMUSG00000032342 | 481 | 27 | 368 | 46 | 450 | 40 | 405 | 41 |
| Mtr | ENSMUSG00000021311 | 3225 | 471 | 2012 | 528 | 2736 | 435 | 2132 | 259 |
| Mtus2 | ENSMUSG00000029651 | 2962 | 248 | 2267 | 102 | 3029 | 325 | 2381 | 425 |
| Myadml2 | ENSMUSG00000025141 | 267 | 6 | 178 | 37 | 256 | 34 | 227 | 25 |
| Mybbp1a | ENSMUSG00000040463 | 1302 | 174 | 1041 | 48 | 1189 | 167 | 1099 | 79 |
| Myh14 | ENSMUSG00000030739 | 1825 | 190 | 1436 | 214 | 1992 | 381 | 1613 | 123 |
| Myh6 | ENSMUSG00000040752 | 416670 | 31586 | 295124 | 67491 | 432469 | 60382 | 365274 | 31724 |
| Mylip | ENSMUSG00000038175 | 408 | 59 | 322 | 55 | 361 | 21 | 336 | 42 |
| Myzap | ENSMUSG00000041361 | 8078 | 741 | 6069 | 777 | 8163 | 545 | 7067 | 721 |
| Nampt | ENSMUSG00000020572 | 4880 | 529 | 3965 | 206 | 4631 | 388 | 3944 | 359 |
| Nbas | ENSMUSG00000020576 | 829 | 71 | 596 | 58 | 720 | 104 | 703 | 64 |
| Ndrg2 | ENSMUSG00000004558 | 19416 | 536 | 16383 | 1310 | 19419 | 1235 | 17364 | 725 |
| Ndufa10 | ENSMUSG00000026260 | 9566 | 706 | 7436 | 1132 | 9920 | 331 | 8450 | 886 |
| Ndufa8 | ENSMUSG00000026895 | 4878 | 294 | 3771 | 635 | 4757 | 148 | 4279 | 331 |
| Ndufa9 | ENSMUSG00000000399 | 9480 | 996 | 7084 | 1337 | 9287 | 369 | 7867 | 703 |
| Ndufs1 | ENSMUSG00000025968 | 15975 | 1815 | 11734 | 2163 | 15650 | 1579 | 13236 | 1027 |
| Ndufs2 | ENSMUSG00000013593 | 16031 | 729 | 11634 | 1773 | 15858 | 991 | 13476 | 1336 |
| Ndufs3 | ENSMUSG00000005510 | 6047 | 349 | 4620 | 872 | 5917 | 121 | 5197 | 447 |
| Ndufs7 | ENSMUSG00000020153 | 3525 | 187 | 2485 | 513 | 3595 | 317 | 2923 | 370 |
| Ndufv1 | ENSMUSG00000037916 | 9773 | 762 | 6974 | 1216 | 9669 | 615 | 8148 | 626 |
| Nectin2 | ENSMUSG00000062300 | 413 | 36 | 310 | 34 | 417 | 41 | 335 | 50 |
| Nek9 | ENSMUSG00000034290 | 4497 | 135 | 3774 | 292 | 4482 | 346 | 4215 | 107 |
| Nfe2l1 | ENSMUSG00000038615 | 10454 | 393 | 8704 | 788 | 10418 | 916 | 9618 | 241 |
| Nfs1 | ENSMUSG00000027618 | 1552 | 137 | 1235 | 133 | 1562 | 117 | 1353 | 125 |
| Ngrn | ENSMUSG00000047084 | 316 | 43 | 243 | 18 | 297 | 10 | 268 | 29 |
| Nmnat3 | ENSMUSG00000032456 | 318 | 32 | 227 | 37 | 294 | 26 | 237 | 22 |
| Nomo1 | ENSMUSG00000030835 | 2843 | 188 | 2392 | 196 | 2879 | 272 | 2584 | 198 |
| Npepps | ENSMUSG00000001441 | 2325 | 152 | 1944 | 99 | 2270 | 128 | 2110 | 152 |
| Nprl2 | ENSMUSG00000010057 | 233 | 23 | 177 | 25 | 218 | 16 | 188 | 17 |
| Nqo2 | ENSMUSG00000046949 | 946 | 76 | 768 | 84 | 944 | 37 | 822 | 27 |
| Nr1d1 | ENSMUSG00000020889 | 2205 | 482 | 1594 | 392 | 2278 | 416 | 1686 | 114 |
| Nr3c2 | ENSMUSG00000031618 | 554 | 97 | 352 | 33 | 533 | 70 | 406 | 87 |
| Nsmce1 | ENSMUSG00000030750 | 572 | 26 | 425 | 55 | 506 | 32 | 480 | 49 |
| Nsun4 | ENSMUSG00000028706 | 668 | 48 | 514 | 69 | 656 | 35 | 592 | 35 |
| Nt5c1a | ENSMUSG00000054958 | 221 | 21 | 132 | 21 | 214 | 28 | 180 | 22 |
| Nt5dc3 | ENSMUSG00000054027 | 1591 | 190 | 1169 | 296 | 1701 | 183 | 1426 | 165 |
| Ntn1 | ENSMUSG00000020902 | 1895 | 146 | 1435 | 153 | 2064 | 197 | 1793 | 251 |
| Nudc | ENSMUSG00000028851 | 1219 | 28 | 997 | 119 | 1222 | 68 | 1116 | 61 |
| Nudt3 | ENSMUSG00000024213 | 2420 | 197 | 1955 | 179 | 2428 | 117 | 2144 | 103 |
| Nudt6 | ENSMUSG00000050174 | 240 | 31 | 183 | 19 | 270 | 13 | 214 | 24 |
| Oat | ENSMUSG00000030934 | 4023 | 214 | 3428 | 322 | 3893 | 236 | 3570 | 97 |
| Ogdh | ENSMUSG00000020456 | 39640 | 2492 | 29734 | 4328 | 41635 | 5116 | 35518 | 902 |
| Oma1 | ENSMUSG00000035069 | 520 | 27 | 382 | 67 | 517 | 40 | 433 | 46 |
| Opa1 | ENSMUSG00000038084 | 5644 | 514 | 4210 | 706 | 5735 | 550 | 5001 | 562 |
| Oplah | ENSMUSG00000022562 | 803 | 117 | 550 | 66 | 755 | 147 | 645 | 45 |

| | | | | | | | | | |
|---|---|---|---|---|---|---|---|---|---|
| Optn | ENSMUSG00000026672 | 2167 | 261 | 1708 | 294 | 2308 | 86 | 2033 | 94 |
| Osbp | ENSMUSG00000024687 | 2871 | 112 | 2475 | 227 | 3049 | 216 | 2682 | 119 |
| Osbpl2 | ENSMUSG00000039050 | 924 | 46 | 750 | 55 | 906 | 59 | 792 | 22 |
| Osbpl6 | ENSMUSG00000042359 | 560 | 64 | 401 | 56 | 579 | 105 | 481 | 24 |
| Osgep | ENSMUSG00000006289 | 680 | 53 | 499 | 68 | 589 | 47 | 573 | 41 |
| Otud4 | ENSMUSG00000036990 | 1969 | 261 | 1551 | 87 | 1916 | 144 | 1630 | 131 |
| Oxa1l | ENSMUSG00000000959 | 1341 | 57 | 1036 | 150 | 1356 | 195 | 1142 | 95 |
| Oxct1 | ENSMUSG00000022186 | 26538 | 1570 | 20521 | 2412 | 25456 | 2358 | 23051 | 1272 |
| Oxnad1 | ENSMUSG00000021906 | 1147 | 104 | 870 | 148 | 1159 | 96 | 1055 | 90 |
| Pacsin2 | ENSMUSG00000016664 | 4608 | 491 | 3453 | 568 | 4770 | 416 | 3975 | 269 |
| Pccb | ENSMUSG00000032527 | 1749 | 37 | 1306 | 178 | 1783 | 212 | 1458 | 93 |
| Pcp4l1 | ENSMUSG00000038370 | 3448 | 91 | 2643 | 293 | 3460 | 188 | 3005 | 313 |
| Pde2a | ENSMUSG00000110195 | 1585 | 126 | 1282 | 143 | 1439 | 108 | 1257 | 211 |
| Pdf | ENSMUSG00000078931 | 565 | 42 | 418 | 72 | 575 | 29 | 492 | 19 |
| Pdk2 | ENSMUSG00000038967 | 7045 | 426 | 4922 | 991 | 7351 | 661 | 5815 | 537 |
| Pdss2 | ENSMUSG00000038240 | 428 | 19 | 288 | 48 | 415 | 44 | 365 | 47 |
| Pdzd2 | ENSMUSG00000022197 | 1958 | 240 | 1387 | 312 | 1714 | 300 | 1447 | 149 |
| Peg13 | ENSMUSG00000106847 | 1266 | 135 | 996 | 129 | 1259 | 67 | 1062 | 102 |
| Perm1 | ENSMUSG00000078486 | 6061 | 537 | 4239 | 948 | 6388 | 499 | 5164 | 309 |
| Pex10 | ENSMUSG00000029047 | 216 | 17 | 157 | 21 | 236 | 16 | 189 | 19 |
| Pex6 | ENSMUSG00000002763 | 536 | 39 | 383 | 52 | 543 | 100 | 441 | 44 |
| Pfkl | ENSMUSG00000020277 | 1893 | 156 | 1558 | 139 | 1871 | 165 | 1710 | 80 |
| Pfkm | ENSMUSG00000033065 | 18225 | 1341 | 12907 | 1890 | 18352 | 1337 | 14993 | 396 |
| Pgm5 | ENSMUSG00000041731 | 2496 | 117 | 1972 | 181 | 2577 | 106 | 2310 | 218 |
| Phf20 | ENSMUSG00000038116 | 645 | 141 | 463 | 64 | 547 | 79 | 484 | 111 |
| Phkg1 | ENSMUSG00000025537 | 414 | 78 | 217 | 95 | 351 | 52 | 195 | 50 |
| Phpt1 | ENSMUSG00000036504 | 592 | 39 | 477 | 58 | 591 | 19 | 515 | 20 |
| Phyh | ENSMUSG00000026664 | 9819 | 550 | 7600 | 1012 | 9908 | 282 | 8331 | 547 |
| Pigg | ENSMUSG00000029263 | 357 | 15 | 277 | 13 | 360 | 32 | 317 | 44 |
| Pigv | ENSMUSG00000043257 | 190 | 34 | 113 | 16 | 185 | 35 | 150 | 33 |
| Pik3r4 | ENSMUSG00000032571 | 510 | 63 | 421 | 32 | 453 | 52 | 451 | 38 |
| Pim3 | ENSMUSG00000035828 | 1043 | 153 | 638 | 156 | 989 | 141 | 713 | 162 |
| Pip4k2c | ENSMUSG00000025417 | 839 | 38 | 718 | 27 | 859 | 49 | 781 | 35 |
| Pitpnc1 | ENSMUSG00000040430 | 2019 | 131 | 1451 | 306 | 1945 | 148 | 1677 | 225 |
| Pitrm1 | ENSMUSG00000021193 | 1183 | 91 | 917 | 87 | 1076 | 97 | 982 | 47 |
| Plin4 | ENSMUSG00000002831 | 3263 | 103 | 2596 | 321 | 3280 | 102 | 2808 | 366 |
| Plpbp | ENSMUSG00000031485 | 1297 | 158 | 1038 | 125 | 1269 | 103 | 1218 | 88 |
| Pmpca | ENSMUSG00000026926 | 2809 | 172 | 2292 | 228 | 2800 | 157 | 2538 | 100 |
| Pnpla2 | ENSMUSG00000025509 | 4617 | 330 | 3361 | 654 | 4556 | 476 | 3773 | 317 |
| Poldip2 | ENSMUSG00000001100 | 2037 | 140 | 1588 | 200 | 2103 | 131 | 1870 | 118 |
| Polr2b | ENSMUSG00000029250 | 1491 | 97 | 1240 | 88 | 1445 | 75 | 1325 | 88 |
| Polr2e | ENSMUSG00000004667 | 833 | 69 | 697 | 28 | 814 | 44 | 761 | 49 |
| Polr2m | ENSMUSG00000032199 | 4202 | 139 | 3527 | 289 | 4270 | 294 | 3840 | 106 |
| Ppara | ENSMUSG00000022383 | 730 | 69 | 463 | 119 | 639 | 77 | 525 | 69 |
| Ppm1k | ENSMUSG00000037826 | 2243 | 337 | 1519 | 318 | 2006 | 174 | 1607 | 419 |
| Ppp1r12b | ENSMUSG00000073557 | 7349 | 246 | 5778 | 1053 | 8073 | 1048 | 6703 | 244 |
| Ppp1r26 | ENSMUSG00000035829 | 123 | 33 | 63 | 14 | 99 | 6 | 69 | 7 |
| Ppp1r3d | ENSMUSG00000049999 | 323 | 31 | 243 | 23 | 344 | 39 | 274 | 29 |
| Ppp5c | ENSMUSG00000003099 | 1134 | 58 | 913 | 91 | 1135 | 59 | 1032 | 79 |
| Ppt2 | ENSMUSG00000015474 | 570 | 20 | 441 | 69 | 579 | 54 | 505 | 43 |
| Prdx6 | ENSMUSG00000026701 | 3412 | 145 | 2618 | 272 | 3250 | 102 | 2855 | 190 |
| Prkaca | ENSMUSG00000005469 | 4064 | 153 | 3118 | 405 | 4060 | 252 | 3598 | 192 |
| Prkag1 | ENSMUSG00000067713 | 1480 | 97 | 1136 | 121 | 1449 | 53 | 1228 | 110 |
| Prkce | ENSMUSG00000045038 | 1038 | 123 | 798 | 114 | 1083 | 139 | 932 | 92 |
| Prpf19 | ENSMUSG00000024735 | 3570 | 220 | 2865 | 389 | 3749 | 422 | 3240 | 103 |
| Prpf8 | ENSMUSG00000020850 | 3805 | 235 | 3158 | 312 | 3806 | 301 | 3465 | 198 |
| Psap | ENSMUSG00000004207 | 20301 | 713 | 16734 | 1671 | 20510 | 1354 | 18075 | 1122 |
| Psmd2 | ENSMUSG00000006998 | 4504 | 254 | 3688 | 415 | 4294 | 293 | 4129 | 211 |
| Ptcd3 | ENSMUSG00000063884 | 3497 | 444 | 2527 | 522 | 3299 | 255 | 2951 | 253 |
| Ptov1 | ENSMUSG00000038502 | 994 | 34 | 794 | 32 | 947 | 73 | 875 | 61 |
| Ptpn3 | ENSMUSG00000038764 | 1516 | 118 | 1158 | 191 | 1433 | 94 | 1204 | 64 |
| Pttg1 | ENSMUSG00000020415 | 1218 | 104 | 917 | 152 | 1220 | 46 | 1000 | 82 |
| Pxmp2 | ENSMUSG00000029499 | 803 | 44 | 554 | 114 | 790 | 47 | 628 | 75 |
| Pygm | ENSMUSG00000032648 | 19355 | 1268 | 13869 | 2475 | 19815 | 1520 | 16110 | 975 |

| | | | | | | | | | |
|---|---|---|---|---|---|---|---|---|---|
| Qdpr | ENSMUSG00000015806 | 905 | 53 | 716 | 59 | 901 | 38 | 833 | 92 |
| Qsox2 | ENSMUSG00000036327 | 354 | 29 | 255 | 33 | 340 | 26 | 293 | 34 |
| R3hdm4 | ENSMUSG00000035781 | 843 | 29 | 689 | 82 | 840 | 52 | 717 | 45 |
| Rab12 | ENSMUSG00000023460 | 2932 | 232 | 2405 | 100 | 2976 | 158 | 2577 | 278 |
| Rab28 | ENSMUSG00000029128 | 1501 | 127 | 1261 | 120 | 1546 | 89 | 1421 | 64 |
| Rab3a | ENSMUSG00000031840 | 556 | 31 | 385 | 62 | 576 | 29 | 459 | 63 |
| Rai2 | ENSMUSG00000043518 | 607 | 58 | 481 | 89 | 647 | 60 | 562 | 52 |
| Ralgapa2 | ENSMUSG00000037110 | 2617 | 245 | 1978 | 383 | 2581 | 274 | 2195 | 200 |
| Ralgapb | ENSMUSG00000027652 | 1478 | 136 | 1208 | 120 | 1466 | 153 | 1287 | 73 |
| Rapsn | ENSMUSG00000002104 | 164 | 14 | 112 | 16 | 166 | 20 | 118 | 21 |
| Rasd2 | ENSMUSG00000034472 | 85 | 16 | 52 | 8 | 87 | 5 | 67 | 16 |
| Rbbp5 | ENSMUSG00000026439 | 503 | 77 | 394 | 43 | 462 | 27 | 433 | 19 |
| Rbfa | ENSMUSG00000024570 | 537 | 27 | 421 | 57 | 552 | 37 | 476 | 23 |
| Rbm20 | ENSMUSG00000043639 | 4345 | 353 | 3333 | 538 | 4405 | 474 | 3671 | 314 |
| Rbm24 | ENSMUSG00000038132 | 2769 | 152 | 2054 | 213 | 2701 | 211 | 2285 | 323 |
| Rbpms | ENSMUSG00000031586 | 2091 | 178 | 1725 | 171 | 2143 | 145 | 1933 | 119 |
| Rdm1 | ENSMUSG00000010362 | 355 | 44 | 260 | 27 | 395 | 55 | 308 | 50 |
| Retnla | ENSMUSG00000061100 | 77 | 26 | 28 | 12 | 70 | 10 | 47 | 28 |
| Retsat | ENSMUSG00000056666 | 937 | 102 | 713 | 90 | 893 | 134 | 765 | 34 |
| Rgs7 | ENSMUSG00000026527 | 36 | 12 | 15 | 6 | 19 | 3 | 25 | 6 |
| Rhobtb2 | ENSMUSG00000022075 | 496 | 77 | 385 | 66 | 531 | 42 | 452 | 21 |
| Rhot2 | ENSMUSG00000025733 | 3078 | 130 | 2175 | 480 | 3032 | 157 | 2480 | 193 |
| Rmnd1 | ENSMUSG00000019763 | 878 | 53 | 704 | 63 | 926 | 55 | 800 | 97 |
| Rnf114 | ENSMUSG00000006418 | 1224 | 48 | 1047 | 46 | 1232 | 102 | 1117 | 38 |
| Rpa1 | ENSMUSG00000000751 | 1034 | 51 | 817 | 60 | 1036 | 89 | 910 | 49 |
| Rpap1 | ENSMUSG00000034032 | 295 | 38 | 221 | 36 | 319 | 44 | 271 | 27 |
| Rpusd4 | ENSMUSG00000032044 | 283 | 14 | 221 | 31 | 291 | 18 | 238 | 21 |
| Rragd | ENSMUSG00000028278 | 3170 | 310 | 2413 | 232 | 3220 | 162 | 2839 | 371 |
| Rrn3 | ENSMUSG00000022682 | 1172 | 63 | 954 | 93 | 1149 | 64 | 1040 | 78 |
| Rtn4ip1 | ENSMUSG00000019864 | 1524 | 104 | 1118 | 239 | 1538 | 112 | 1315 | 140 |
| Rufy1 | ENSMUSG00000020375 | 709 | 60 | 558 | 43 | 649 | 12 | 610 | 48 |
| Rxrg | ENSMUSG00000015843 | 1040 | 79 | 760 | 162 | 1102 | 94 | 896 | 96 |
| Samm50 | ENSMUSG00000022437 | 4316 | 204 | 3494 | 463 | 4136 | 107 | 3926 | 74 |
| Sbk1 | ENSMUSG00000042978 | 1288 | 124 | 1004 | 171 | 1329 | 89 | 1218 | 118 |
| Scgb1c1 | ENSMUSG00000038801 | 156 | 23 | 101 | 19 | 163 | 23 | 131 | 27 |
| Scn4a | ENSMUSG00000001027 | 691 | 157 | 407 | 125 | 675 | 132 | 480 | 75 |
| Scn4b | ENSMUSG00000046480 | 1055 | 187 | 607 | 202 | 976 | 267 | 669 | 172 |
| Scrn3 | ENSMUSG00000008226 | 772 | 46 | 634 | 36 | 772 | 89 | 686 | 45 |
| Sdhb | ENSMUSG00000009863 | 12188 | 749 | 9065 | 1722 | 11819 | 503 | 9978 | 990 |
| Sdhc | ENSMUSG00000058076 | 7127 | 66 | 5010 | 729 | 7101 | 434 | 5730 | 395 |
| Sdr39u1 | ENSMUSG00000022223 | 1168 | 90 | 858 | 177 | 1171 | 36 | 977 | 118 |
| Sel1l | ENSMUSG00000020964 | 1913 | 94 | 1550 | 157 | 1805 | 94 | 1646 | 186 |
| Sesn1 | ENSMUSG00000038332 | 2443 | 189 | 2090 | 127 | 2405 | 179 | 2161 | 117 |
| Sipa1l2 | ENSMUSG00000001995 | 1430 | 165 | 1122 | 105 | 1540 | 153 | 1327 | 56 |
| Sirt5 | ENSMUSG00000054021 | 574 | 24 | 464 | 39 | 573 | 30 | 507 | 40 |
| Slc16a7 | ENSMUSG00000020102 | 227 | 56 | 144 | 27 | 216 | 67 | 189 | 39 |
| Slc25a11 | ENSMUSG00000014606 | 8077 | 561 | 5928 | 1242 | 7992 | 341 | 6986 | 662 |
| Slc25a12 | ENSMUSG00000027010 | 4184 | 351 | 2893 | 395 | 4194 | 379 | 3611 | 151 |
| Slc25a13 | ENSMUSG00000015112 | 2989 | 324 | 2199 | 145 | 3029 | 276 | 2574 | 268 |
| Slc25a20 | ENSMUSG00000032602 | 3086 | 349 | 2262 | 477 | 3063 | 140 | 2521 | 210 |
| Slc25a3 | ENSMUSG00000061904 | 25015 | 1211 | 18063 | 2479 | 25124 | 1400 | 20806 | 1485 |
| Slc25a33 | ENSMUSG00000028982 | 271 | 36 | 196 | 32 | 254 | 25 | 214 | 18 |
| Slc26a6 | ENSMUSG00000023259 | 159 | 5 | 105 | 24 | 133 | 7 | 122 | 15 |
| Slc2a4 | ENSMUSG00000018566 | 6195 | 156 | 4116 | 759 | 6192 | 600 | 4937 | 362 |
| Slc36a2 | ENSMUSG00000020264 | 315 | 40 | 215 | 36 | 309 | 9 | 239 | 36 |
| Slc38a3 | ENSMUSG00000010064 | 939 | 97 | 638 | 211 | 1037 | 121 | 746 | 126 |
| Slc41a1 | ENSMUSG00000013275 | 1492 | 118 | 1131 | 142 | 1374 | 124 | 1187 | 145 |
| Slc4a4 | ENSMUSG00000060961 | 964 | 160 | 696 | 87 | 861 | 146 | 774 | 132 |
| Slc7a1 | ENSMUSG00000041313 | 1469 | 249 | 990 | 213 | 1397 | 281 | 1151 | 217 |
| Slc9a8 | ENSMUSG00000039463 | 533 | 54 | 422 | 27 | 481 | 53 | 424 | 37 |
| Slco3a1 | ENSMUSG00000025790 | 1897 | 213 | 1577 | 239 | 1938 | 213 | 1809 | 168 |
| Smco1 | ENSMUSG00000046345 | 280 | 33 | 173 | 42 | 270 | 15 | 222 | 35 |
| Smg5 | ENSMUSG00000001415 | 1134 | 42 | 904 | 93 | 1081 | 79 | 1020 | 136 |
| Smim11 | ENSMUSG00000051989 | 669 | 30 | 545 | 65 | 628 | 41 | 590 | 28 |

| | | | | | | | | | |
|---|---|---|---|---|---|---|---|---|---|
| Snai3 | ENSMUSG00000006587 | 49 | 18 | 24 | 5 | 45 | 14 | 35 | 8 |
| Snrpn | ENSMUSG00000102252 | 939 | 55 | 661 | 136 | 989 | 67 | 760 | 142 |
| Sod1 | ENSMUSG00000022982 | 3578 | 226 | 3002 | 251 | 3367 | 134 | 3215 | 151 |
| Sod2 | ENSMUSG00000006818 | 12266 | 831 | 8665 | 1885 | 11982 | 506 | 9979 | 664 |
| Sord | ENSMUSG00000027227 | 4541 | 245 | 3023 | 680 | 4444 | 324 | 3516 | 336 |
| Spr | ENSMUSG00000033735 | 665 | 53 | 505 | 56 | 683 | 71 | 580 | 44 |
| Spsb1 | ENSMUSG00000039911 | 519 | 41 | 383 | 46 | 499 | 68 | 431 | 61 |
| Sptb | ENSMUSG00000021061 | 3814 | 253 | 2892 | 391 | 4015 | 448 | 3358 | 226 |
| St3gal3 | ENSMUSG00000028538 | 1047 | 50 | 840 | 121 | 1048 | 65 | 941 | 63 |
| St6galnac6 | ENSMUSG00000026811 | 1129 | 64 | 895 | 132 | 1127 | 72 | 981 | 112 |
| Stard10 | ENSMUSG00000030688 | 330 | 75 | 189 | 43 | 297 | 12 | 242 | 66 |
| Stard7 | ENSMUSG00000027367 | 3728 | 299 | 2766 | 387 | 3568 | 81 | 3253 | 316 |
| Stk11 | ENSMUSG00000003068 | 1448 | 37 | 1205 | 94 | 1447 | 59 | 1293 | 70 |
| Ston2 | ENSMUSG00000020961 | 279 | 82 | 178 | 41 | 317 | 59 | 217 | 35 |
| Stub1 | ENSMUSG00000039615 | 1096 | 59 | 905 | 66 | 1104 | 71 | 1014 | 92 |
| Suclg1 | ENSMUSG00000052738 | 6364 | 514 | 4599 | 1021 | 6012 | 279 | 5327 | 363 |
| Suclg2 | ENSMUSG00000061838 | 4105 | 321 | 2961 | 428 | 4037 | 119 | 3294 | 228 |
| Suox | ENSMUSG00000049858 | 473 | 53 | 354 | 36 | 448 | 32 | 405 | 21 |
| Susd6 | ENSMUSG00000021133 | 2324 | 150 | 1943 | 183 | 2364 | 108 | 2034 | 109 |
| Svip | ENSMUSG00000074093 | 948 | 93 | 733 | 96 | 873 | 98 | 769 | 67 |
| Swsap1 | ENSMUSG00000051238 | 126 | 10 | 90 | 15 | 118 | 11 | 105 | 11 |
| Syt7 | ENSMUSG00000024743 | 873 | 100 | 595 | 126 | 831 | 116 | 700 | 57 |
| Tango2 | ENSMUSG00000013539 | 2372 | 272 | 1812 | 121 | 2322 | 147 | 1927 | 149 |
| Tango6 | ENSMUSG00000041949 | 231 | 19 | 170 | 17 | 230 | 31 | 201 | 14 |
| Tars2 | ENSMUSG00000028107 | 667 | 59 | 489 | 66 | 606 | 23 | 549 | 66 |
| Tarsl2 | ENSMUSG00000030515 | 1088 | 105 | 839 | 102 | 1083 | 60 | 931 | 19 |
| Taz | ENSMUSG00000009995 | 683 | 45 | 574 | 36 | 641 | 57 | 573 | 45 |
| Tbrg4 | ENSMUSG00000000384 | 909 | 70 | 673 | 136 | 897 | 113 | 763 | 112 |
| Tcaim | ENSMUSG00000046603 | 1091 | 140 | 752 | 164 | 1078 | 129 | 865 | 134 |
| Tcap | ENSMUSG00000007877 | 15023 | 5676 | 9524 | 3481 | 11808 | 3513 | 10365 | 2562 |
| Tcp11l2 | ENSMUSG00000020034 | 2121 | 130 | 1478 | 128 | 2131 | 148 | 1765 | 222 |
| Tecrl | ENSMUSG00000049537 | 1305 | 197 | 1030 | 90 | 1191 | 128 | 1094 | 181 |
| Tent4b | ENSMUSG00000036779 | 610 | 55 | 482 | 40 | 562 | 52 | 508 | 45 |
| Tesc | ENSMUSG00000029359 | 1119 | 84 | 854 | 76 | 1160 | 95 | 937 | 103 |
| Thrb | ENSMUSG00000021779 | 495 | 72 | 351 | 62 | 440 | 76 | 311 | 43 |
| Tmbim6 | ENSMUSG00000023010 | 3804 | 173 | 3339 | 158 | 3817 | 189 | 3501 | 211 |
| Tmc7 | ENSMUSG00000042246 | 203 | 26 | 121 | 17 | 221 | 36 | 169 | 29 |
| Tmem135 | ENSMUSG00000039428 | 983 | 73 | 725 | 115 | 985 | 79 | 815 | 63 |
| Tmem143 | ENSMUSG00000002781 | 1504 | 67 | 919 | 258 | 1446 | 221 | 1083 | 117 |
| Tmem177 | ENSMUSG00000036975 | 283 | 12 | 189 | 34 | 271 | 12 | 216 | 21 |
| Tmem250-ps | ENSMUSG00000087679 | 1255 | 42 | 1017 | 136 | 1234 | 98 | 1140 | 53 |
| Tmem38a | ENSMUSG00000031791 | 7478 | 344 | 5801 | 950 | 7515 | 482 | 6828 | 346 |
| Tmem50b | ENSMUSG00000022964 | 927 | 63 | 728 | 73 | 909 | 32 | 769 | 105 |
| Tmem70 | ENSMUSG00000025940 | 1686 | 231 | 1310 | 188 | 1682 | 176 | 1404 | 135 |
| Tmem82 | ENSMUSG00000043085 | 271 | 51 | 168 | 43 | 257 | 20 | 187 | 17 |
| Tmem94 | ENSMUSG00000020747 | 1735 | 172 | 1255 | 258 | 1762 | 215 | 1313 | 178 |
| Tmod1 | ENSMUSG00000028328 | 6866 | 511 | 5456 | 542 | 6671 | 424 | 6180 | 413 |
| Tmod4 | ENSMUSG00000005628 | 378 | 37 | 254 | 60 | 361 | 43 | 317 | 59 |
| Tnfaip8 | ENSMUSG00000062210 | 1646 | 269 | 1190 | 267 | 1502 | 102 | 1310 | 232 |
| Tnip3 | ENSMUSG00000044162 | 52 | 11 | 27 | 13 | 47 | 4 | 42 | 4 |
| Tnni3 | ENSMUSG00000035458 | 63108 | 4479 | 42722 | 10485 | 61514 | 1321 | 48873 | 5226 |
| Tnnt2 | ENSMUSG00000026414 | 110161 | 8926 | 87055 | 9068 | 105408 | 3834 | 92241 | 8335 |
| Tom1l2 | ENSMUSG00000000538 | 2948 | 122 | 2426 | 287 | 2928 | 258 | 2694 | 186 |
| Trabd2b | ENSMUSG00000070867 | 3038 | 207 | 2320 | 372 | 3187 | 181 | 2778 | 256 |
| Tsc22d1 | ENSMUSG00000022010 | 4439 | 536 | 3676 | 350 | 4405 | 178 | 4019 | 379 |
| Tspan3 | ENSMUSG00000032324 | 2417 | 116 | 2074 | 132 | 2505 | 165 | 2302 | 52 |
| Tspyl4 | ENSMUSG00000039485 | 376 | 62 | 276 | 18 | 382 | 85 | 298 | 73 |
| Ttc19 | ENSMUSG00000042298 | 902 | 120 | 684 | 71 | 896 | 69 | 748 | 78 |
| Tufm | ENSMUSG00000073838 | 3132 | 215 | 2311 | 479 | 3147 | 191 | 2714 | 265 |
| Twnk | ENSMUSG00000025209 | 604 | 21 | 452 | 67 | 607 | 59 | 505 | 23 |
| Txnrd2 | ENSMUSG00000075704 | 527 | 35 | 362 | 47 | 464 | 30 | 431 | 49 |
| Ubac2 | ENSMUSG00000041765 | 482 | 28 | 351 | 27 | 440 | 56 | 380 | 38 |
| Ubl7 | ENSMUSG00000055720 | 695 | 84 | 565 | 67 | 663 | 44 | 641 | 37 |
| Ubr2 | ENSMUSG00000023977 | 2578 | 83 | 1942 | 289 | 2546 | 196 | 2215 | 196 |

| | | | | | | | | | |
|---|---|---|---|---|---|---|---|---|---|
| Ucp3 | ENSMUSG00000032942 | 1264 | 355 | 659 | 227 | 1281 | 331 | 698 | 271 |
| Unc45b | ENSMUSG00000018845 | 3432 | 81 | 2694 | 372 | 3138 | 382 | 2858 | 156 |
| Uqcc1 | ENSMUSG00000005882 | 3454 | 270 | 2476 | 553 | 3363 | 150 | 2909 | 228 |
| Uqcrc1 | ENSMUSG00000025651 | 15130 | 710 | 10190 | 2080 | 15168 | 907 | 12233 | 1293 |
| Uqcrfs1 | ENSMUSG00000038462 | 13197 | 914 | 10056 | 1838 | 12772 | 692 | 11183 | 890 |
| Urod | ENSMUSG00000028684 | 800 | 46 | 616 | 62 | 756 | 47 | 678 | 29 |
| Usf2 | ENSMUSG00000058239 | 1261 | 122 | 1046 | 81 | 1203 | 37 | 1081 | 60 |
| Vcp | ENSMUSG00000028452 | 9696 | 493 | 8255 | 496 | 9439 | 494 | 9118 | 332 |
| Vdac1 | ENSMUSG00000020402 | 17517 | 801 | 13326 | 1578 | 17010 | 633 | 14751 | 342 |
| Vdac3 | ENSMUSG00000008892 | 6247 | 446 | 4930 | 675 | 5939 | 119 | 5490 | 330 |
| Vegfb | ENSMUSG00000024962 | 2831 | 191 | 2209 | 301 | 2977 | 265 | 2434 | 179 |
| Vps26a | ENSMUSG00000020078 | 1810 | 210 | 1524 | 43 | 1767 | 104 | 1632 | 155 |
| Vwa8 | ENSMUSG00000058997 | 6184 | 159 | 4148 | 762 | 5901 | 635 | 4704 | 407 |
| Wbp2 | ENSMUSG00000034341 | 1379 | 72 | 1162 | 115 | 1438 | 101 | 1294 | 92 |
| Wdtc1 | ENSMUSG00000037622 | 1386 | 86 | 1042 | 217 | 1413 | 154 | 1245 | 104 |
| Wfs1 | ENSMUSG00000039474 | 1806 | 82 | 1416 | 252 | 1797 | 191 | 1634 | 135 |
| Whrn | ENSMUSG00000039137 | 481 | 110 | 307 | 100 | 542 | 79 | 382 | 97 |
| Wnk2 | ENSMUSG00000037989 | 1441 | 166 | 821 | 252 | 1510 | 204 | 1013 | 246 |
| Ybx1 | ENSMUSG00000028639 | 11917 | 513 | 10296 | 764 | 11978 | 639 | 10588 | 490 |
| Ywhae | ENSMUSG00000020849 | 5973 | 224 | 5142 | 422 | 5882 | 295 | 5570 | 239 |
| Zadh2 | ENSMUSG00000049090 | 1687 | 130 | 1330 | 146 | 1727 | 133 | 1433 | 161 |
| Zfp113 | ENSMUSG00000037007 | 207 | 26 | 137 | 20 | 206 | 31 | 160 | 32 |
| Zfp536 | ENSMUSG00000043456 | 134 | 18 | 70 | 15 | 120 | 18 | 92 | 13 |
| Zfp629 | ENSMUSG00000045639 | 654 | 44 | 510 | 45 | 651 | 90 | 565 | 56 |
| Zfyve21 | ENSMUSG00000021286 | 406 | 15 | 295 | 34 | 394 | 31 | 309 | 42 |
| Znrf1 | ENSMUSG00000033545 | 850 | 52 | 714 | 48 | 788 | 71 | 705 | 40 |

**Supplementary Table S8.** RNASeq analysis of effects of angiotensin II (AngII) on mRNA expression in hearts from PKN2Het vs WT littermates: mRNAs significantly upregulated by AngII in PKN2Het hearts.

| Gene Symbol | Ensembl gene id | WT Vehicle | | WT AngII | | PKN2Het Vehicle | | PKN2Het AngII | |
|---|---|---|---|---|---|---|---|---|---|
| | | Mean | SD | Mean | SD | Mean | SD | Mean | SD |
| 1500004A13Rik | ENSMUSG00000098912 | 28 | 5 | 38 | 8 | 20 | 7 | 48 | 20 |
| 4931406P16Rik | ENSMUSG00000066571 | 1213 | 197 | 1375 | 110 | 1210 | 171 | 1441 | 195 |
| 6430584L05Rik | ENSMUSG00000108228 | 2 | 1 | 7 | 2 | 3 | 1 | 69 | 99 |
| AC165271.1 | ENSMUSG00000116641 | 7 | 2 | 16 | 5 | 5 | 2 | 19 | 12 |
| Adgrd1 | ENSMUSG00000044017 | 191 | 51 | 269 | 56 | 169 | 19 | 249 | 49 |
| Akap2 | ENSMUSG00000038729 | 3768 | 234 | 4279 | 435 | 3521 | 379 | 4095 | 258 |
| Aldh18a1 | ENSMUSG00000025007 | 242 | 27 | 280 | 13 | 215 | 33 | 299 | 35 |
| Ano6 | ENSMUSG00000064210 | 778 | 83 | 929 | 100 | 694 | 56 | 876 | 34 |
| Anp32b | ENSMUSG00000028333 | 1235 | 54 | 1423 | 104 | 1216 | 59 | 1418 | 76 |
| Arhgef12 | ENSMUSG00000059495 | 6349 | 600 | 6661 | 1008 | 6125 | 289 | 7218 | 254 |
| Arrb1 | ENSMUSG00000018909 | 877 | 101 | 1086 | 152 | 816 | 90 | 1086 | 112 |
| Atp8b2 | ENSMUSG00000060671 | 531 | 139 | 704 | 103 | 413 | 28 | 673 | 34 |
| Atxn1l | ENSMUSG00000069895 | 724 | 111 | 785 | 64 | 708 | 86 | 852 | 141 |
| B3gnt3 | ENSMUSG00000031803 | 234 | 39 | 304 | 15 | 213 | 12 | 289 | 30 |
| C5ar1 | ENSMUSG00000049130 | 107 | 17 | 173 | 40 | 78 | 18 | 156 | 56 |
| Capn2 | ENSMUSG00000026509 | 2021 | 126 | 2295 | 152 | 1873 | 94 | 2199 | 142 |
| Ccl12 | ENSMUSG00000035352 | 28 | 12 | 76 | 30 | 20 | 5 | 61 | 36 |
| Ccl2 | ENSMUSG00000035385 | 56 | 15 | 106 | 46 | 42 | 6 | 111 | 80 |
| Cd302 | ENSMUSG00000060703 | 206 | 18 | 297 | 71 | 176 | 17 | 266 | 37 |
| Cdc42ep4 | ENSMUSG00000041598 | 293 | 84 | 306 | 18 | 242 | 49 | 312 | 39 |
| Cdc42se1 | ENSMUSG00000046722 | 468 | 71 | 617 | 41 | 442 | 53 | 554 | 68 |
| Cebpa | ENSMUSG00000034957 | 116 | 16 | 166 | 28 | 115 | 23 | 164 | 26 |
| Cenpi | ENSMUSG00000031262 | 14 | 6 | 28 | 12 | 6 | 3 | 28 | 9 |
| Ces2e | ENSMUSG00000031886 | 82 | 18 | 89 | 16 | 65 | 19 | 108 | 34 |
| Clic5 | ENSMUSG00000023959 | 7647 | 453 | 8817 | 809 | 7809 | 595 | 9603 | 708 |
| Cmklr1 | ENSMUSG00000042190 | 507 | 144 | 618 | 103 | 422 | 100 | 579 | 89 |
| Col4a3 | ENSMUSG00000079465 | 203 | 49 | 299 | 93 | 199 | 27 | 307 | 35 |
| Cyb561 | ENSMUSG00000019590 | 237 | 62 | 314 | 54 | 200 | 32 | 299 | 78 |
| Cyb5r1 | ENSMUSG00000026456 | 363 | 22 | 410 | 23 | 351 | 26 | 444 | 45 |
| Dbndd2 | ENSMUSG00000017734 | 265 | 25 | 310 | 15 | 218 | 53 | 285 | 36 |
| Ddah2 | ENSMUSG00000007039 | 240 | 14 | 299 | 36 | 214 | 30 | 292 | 28 |
| Ddr2 | ENSMUSG00000026674 | 1034 | 140 | 1163 | 154 | 869 | 156 | 1130 | 83 |
| Dpysl2 | ENSMUSG00000022048 | 377 | 57 | 499 | 48 | 329 | 67 | 433 | 23 |
| Dsn1 | ENSMUSG00000027635 | 32 | 10 | 50 | 10 | 25 | 6 | 47 | 8 |
| Eif4a1 | ENSMUSG00000059796 | 3380 | 300 | 3664 | 494 | 3084 | 168 | 3826 | 261 |
| Elmo1 | ENSMUSG00000041112 | 303 | 38 | 395 | 22 | 278 | 56 | 438 | 50 |
| Elovl1 | ENSMUSG00000006390 | 286 | 23 | 361 | 51 | 246 | 37 | 316 | 43 |
| Enc1 | ENSMUSG00000041773 | 293 | 76 | 382 | 15 | 273 | 69 | 390 | 41 |
| Erf | ENSMUSG00000040857 | 309 | 56 | 363 | 54 | 285 | 58 | 372 | 48 |
| Fabp5 | ENSMUSG00000027533 | 788 | 95 | 996 | 167 | 674 | 83 | 989 | 37 |
| Fam102b | ENSMUSG00000040339 | 516 | 79 | 657 | 85 | 447 | 67 | 598 | 79 |
| Fgf6 | ENSMUSG00000000183 | 19 | 11 | 37 | 18 | 16 | 6 | 40 | 17 |
| Fkbp5 | ENSMUSG00000024222 | 356 | 106 | 469 | 65 | 289 | 55 | 412 | 74 |
| Gba | ENSMUSG00000028048 | 219 | 25 | 275 | 46 | 194 | 36 | 270 | 21 |
| Gm10275 | ENSMUSG00000069682 | 2119 | 213 | 2280 | 303 | 1958 | 234 | 2331 | 159 |
| Gm14005 | ENSMUSG00000074813 | 25 | 6 | 32 | 5 | 23 | 3 | 43 | 5 |
| Gm15542 | ENSMUSG00000083396 | 41 | 20 | 74 | 30 | 32 | 14 | 92 | 41 |
| Gm21188 | ENSMUSG00000095609 | 24 | 11 | 52 | 38 | 12 | 6 | 39 | 17 |
| Gm8430 | ENSMUSG00000055093 | 747 | 208 | 891 | 177 | 614 | 182 | 1041 | 66 |
| Gngt2 | ENSMUSG00000038811 | 179 | 19 | 226 | 24 | 170 | 41 | 224 | 28 |
| Hbegf | ENSMUSG00000024486 | 285 | 26 | 375 | 79 | 293 | 20 | 445 | 97 |
| Hmox1 | ENSMUSG00000005413 | 189 | 79 | 278 | 49 | 151 | 19 | 270 | 45 |
| Hnrnpf | ENSMUSG00000042079 | 2929 | 181 | 3120 | 203 | 2667 | 95 | 3177 | 258 |
| Hnrnpk | ENSMUSG00000021546 | 4424 | 263 | 4608 | 377 | 3979 | 132 | 4710 | 317 |
| Hsp90aa1 | ENSMUSG00000021270 | 4809 | 792 | 5821 | 774 | 4154 | 437 | 5635 | 799 |

| | | | | | | | | | |
|---|---|---|---|---|---|---|---|---|---|
| Id3 | ENSMUSG00000007872 | 547 | 96 | 629 | 46 | 503 | 70 | 613 | 81 |
| Ier5 | ENSMUSG00000056708 | 536 | 123 | 783 | 105 | 482 | 61 | 803 | 298 |
| Ilk | ENSMUSG00000030890 | 1549 | 39 | 1714 | 90 | 1475 | 117 | 1685 | 57 |
| Itpr2 | ENSMUSG00000030287 | 666 | 151 | 771 | 158 | 613 | 94 | 776 | 101 |
| Jpt1 | ENSMUSG00000020737 | 841 | 76 | 987 | 49 | 790 | 37 | 1002 | 107 |
| Jpt2 | ENSMUSG00000024165 | 169 | 19 | 197 | 35 | 141 | 33 | 226 | 44 |
| Lama2 | ENSMUSG00000019899 | 3507 | 352 | 3968 | 325 | 3362 | 180 | 4015 | 379 |
| Layn | ENSMUSG00000060594 | 73 | 12 | 116 | 39 | 72 | 19 | 117 | 25 |
| Lima1 | ENSMUSG00000023022 | 869 | 137 | 1044 | 86 | 750 | 124 | 1003 | 145 |
| Ly6a | ENSMUSG00000075602 | 1892 | 294 | 2356 | 309 | 1658 | 287 | 2262 | 203 |
| Mad2l1 | ENSMUSG00000029910 | 111 | 20 | 150 | 33 | 90 | 23 | 154 | 23 |
| Map6 | ENSMUSG00000055407 | 108 | 31 | 142 | 36 | 87 | 9 | 129 | 34 |
| Mcm10 | ENSMUSG00000026669 | 14 | 6 | 30 | 10 | 8 | 7 | 26 | 18 |
| Mcm3 | ENSMUSG00000041859 | 156 | 21 | 214 | 41 | 148 | 27 | 240 | 53 |
| Mcm5 | ENSMUSG00000005410 | 84 | 12 | 162 | 45 | 83 | 19 | 169 | 35 |
| Mcm6 | ENSMUSG00000026355 | 230 | 32 | 397 | 77 | 216 | 18 | 363 | 70 |
| Mgat5b | ENSMUSG00000043857 | 8 | 5 | 23 | 15 | 9 | 5 | 31 | 24 |
| Mmrn2 | ENSMUSG00000041445 | 1117 | 202 | 1171 | 66 | 966 | 115 | 1226 | 136 |
| Mob3a | ENSMUSG00000003348 | 134 | 17 | 176 | 20 | 115 | 24 | 176 | 43 |
| Mybl1 | ENSMUSG00000025912 | 20 | 8 | 40 | 11 | 22 | 4 | 48 | 10 |
| Myo1c | ENSMUSG00000017774 | 2939 | 140 | 3445 | 280 | 2807 | 136 | 3414 | 203 |
| Ncf2 | ENSMUSG00000026480 | 117 | 30 | 183 | 21 | 107 | 29 | 174 | 52 |
| Nrp2 | ENSMUSG00000025969 | 1954 | 236 | 2425 | 238 | 1836 | 224 | 2427 | 166 |
| Nuak1 | ENSMUSG00000020032 | 841 | 90 | 1047 | 199 | 819 | 26 | 1165 | 278 |
| Nudt18 | ENSMUSG00000045211 | 271 | 21 | 330 | 55 | 259 | 36 | 362 | 63 |
| Numb | ENSMUSG00000021224 | 620 | 116 | 600 | 28 | 522 | 54 | 673 | 44 |
| Nusap1 | ENSMUSG00000027306 | 48 | 17 | 132 | 40 | 45 | 9 | 116 | 36 |
| Ophn1 | ENSMUSG00000031214 | 212 | 70 | 246 | 57 | 195 | 73 | 284 | 69 |
| Pald1 | ENSMUSG00000020092 | 474 | 36 | 537 | 61 | 417 | 42 | 544 | 37 |
| Papln | ENSMUSG00000021223 | 369 | 42 | 478 | 84 | 367 | 37 | 494 | 57 |
| Parva | ENSMUSG00000030770 | 1225 | 61 | 1409 | 154 | 1113 | 88 | 1320 | 96 |
| Pcolce2 | ENSMUSG00000015354 | 412 | 82 | 494 | 81 | 352 | 55 | 503 | 72 |
| Pdlim1 | ENSMUSG00000055044 | 1237 | 36 | 1524 | 182 | 1078 | 86 | 1607 | 176 |
| Pecam1 | ENSMUSG00000020717 | 4184 | 537 | 5065 | 334 | 3964 | 407 | 4742 | 324 |
| Pid1 | ENSMUSG00000045658 | 275 | 53 | 338 | 14 | 244 | 37 | 314 | 29 |
| Pilra | ENSMUSG00000046245 | 31 | 5 | 52 | 15 | 24 | 9 | 52 | 20 |
| Pole2 | ENSMUSG00000020974 | 16 | 7 | 31 | 8 | 12 | 5 | 33 | 6 |
| Prg4 | ENSMUSG00000006014 | 323 | 89 | 484 | 130 | 321 | 33 | 596 | 281 |
| Ptx3 | ENSMUSG00000027832 | 9 | 5 | 35 | 23 | 8 | 2 | 42 | 47 |
| Rab15 | ENSMUSG00000021062 | 19 | 12 | 25 | 4 | 11 | 5 | 30 | 8 |
| Rad54l | ENSMUSG00000028702 | 11 | 9 | 22 | 6 | 6 | 2 | 28 | 11 |
| Ralb | ENSMUSG00000004451 | 600 | 34 | 674 | 33 | 559 | 34 | 691 | 29 |
| Raph1 | ENSMUSG00000026014 | 2057 | 244 | 2567 | 254 | 1871 | 214 | 2337 | 313 |
| Rasa3 | ENSMUSG00000031453 | 585 | 66 | 758 | 73 | 582 | 31 | 773 | 108 |
| Rhog | ENSMUSG00000073982 | 297 | 22 | 355 | 35 | 256 | 29 | 324 | 32 |
| Ripk3 | ENSMUSG00000022221 | 56 | 11 | 90 | 28 | 34 | 8 | 84 | 24 |
| Rpl3 | ENSMUSG00000060036 | 3377 | 296 | 4471 | 607 | 3263 | 335 | 4320 | 344 |
| Rpl39 | ENSMUSG00000079641 | 2170 | 187 | 2489 | 536 | 1860 | 274 | 2471 | 431 |
| Rps27l | ENSMUSG00000036781 | 985 | 107 | 1026 | 159 | 889 | 103 | 1140 | 141 |
| Rras | ENSMUSG00000038387 | 425 | 64 | 533 | 56 | 410 | 35 | 523 | 72 |
| Sema7a | ENSMUSG00000038264 | 731 | 163 | 865 | 32 | 659 | 93 | 882 | 92 |
| Sh3gl1 | ENSMUSG00000003200 | 234 | 18 | 296 | 44 | 217 | 29 | 272 | 16 |
| Slc39a1 | ENSMUSG00000052310 | 1324 | 29 | 1471 | 82 | 1256 | 48 | 1467 | 64 |
| Slc3a2 | ENSMUSG00000010095 | 496 | 50 | 522 | 83 | 462 | 51 | 597 | 75 |
| Sorbs3 | ENSMUSG00000022091 | 627 | 75 | 731 | 50 | 570 | 54 | 681 | 36 |
| Spsb4 | ENSMUSG00000046997 | 117 | 24 | 148 | 47 | 116 | 20 | 180 | 48 |
| Srpx | ENSMUSG00000090084 | 56 | 2 | 104 | 41 | 56 | 6 | 108 | 42 |
| Thsd1 | ENSMUSG00000031480 | 201 | 35 | 239 | 30 | 176 | 24 | 246 | 27 |
| Tinagl1 | ENSMUSG00000028776 | 767 | 115 | 841 | 137 | 702 | 142 | 896 | 129 |
| Troap | ENSMUSG00000032783 | 9 | 3 | 20 | 8 | 5 | 3 | 20 | 13 |
| Tspan9 | ENSMUSG00000030352 | 1563 | 131 | 1947 | 401 | 1621 | 98 | 2180 | 218 |
| Tubb2a | ENSMUSG00000058672 | 572 | 55 | 703 | 94 | 505 | 25 | 707 | 148 |
| Tubb6 | ENSMUSG00000001473 | 337 | 29 | 445 | 93 | 318 | 39 | 459 | 50 |
| Vsig4 | ENSMUSG00000044206 | 33 | 11 | 57 | 21 | 28 | 10 | 77 | 29 |

| | | | | | | | | | |
|---|---|---|---|---|---|---|---|---|---|
| Wdr1 | ENSMUSG00000005103 | 2521 | 253 | 2721 | 220 | 2501 | 131 | 3026 | 227 |
| Wfdc17 | ENSMUSG00000069792 | 126 | 27 | 205 | 64 | 117 | 24 | 230 | 60 |
| Ywhab | ENSMUSG00000018326 | 2036 | 61 | 2287 | 69 | 1970 | 113 | 2268 | 129 |
| Ywhah | ENSMUSG00000018965 | 1422 | 86 | 1708 | 173 | 1342 | 151 | 1658 | 195 |

**Supplementary Table S9.** RNASeq analysis of effects of angiotensin II (AngII) on mRNA expression in hearts from PKN2Het vs WT littermates: mRNAs significantly downregulated by AngII in PKN2Het hearts.

| Gene Symbol | Ensembl gene id | WT Vehicle | | WT AngII | | PKN2Het Vehicle | | PKN2Het AngII | |
|---|---|---|---|---|---|---|---|---|---|
| | | Mean | SD | Mean | SD | Mean | SD | Mean | SD |
| Abcb7 | ENSMUSG00000031333 | 1213 | 155 | 1016 | 47 | 1337 | 171 | 1067 | 172 |
| AC161607.1 | ENSMUSG00000116903 | 73 | 23 | 44 | 12 | 85 | 11 | 52 | 12 |
| Acaca | ENSMUSG00000020532 | 559 | 140 | 447 | 82 | 557 | 121 | 494 | 41 |
| Antxr2 | ENSMUSG00000029338 | 3138 | 199 | 2527 | 322 | 3402 | 377 | 2661 | 386 |
| Arl8b | ENSMUSG00000030105 | 1872 | 115 | 1674 | 157 | 2035 | 143 | 1684 | 79 |
| Art4 | ENSMUSG00000030217 | 412 | 21 | 326 | 68 | 438 | 37 | 298 | 89 |
| Atl2 | ENSMUSG00000059811 | 1046 | 62 | 921 | 131 | 1052 | 45 | 882 | 97 |
| Atxn2 | ENSMUSG00000042605 | 1186 | 207 | 1021 | 195 | 1395 | 65 | 1084 | 156 |
| Bicra | ENSMUSG00000070808 | 259 | 26 | 236 | 41 | 304 | 46 | 218 | 25 |
| Carnmt1 | ENSMUSG00000024726 | 1249 | 241 | 1074 | 205 | 1403 | 244 | 1098 | 142 |
| Carns1 | ENSMUSG00000075289 | 373 | 32 | 286 | 27 | 383 | 53 | 289 | 49 |
| Cited2 | ENSMUSG00000039910 | 544 | 99 | 394 | 73 | 713 | 201 | 447 | 51 |
| Cog5 | ENSMUSG00000035933 | 1358 | 209 | 1191 | 133 | 1404 | 218 | 1168 | 103 |
| Dcaf12l1 | ENSMUSG00000045284 | 80 | 13 | 49 | 12 | 91 | 12 | 54 | 16 |
| Dgke | ENSMUSG00000000276 | 588 | 81 | 456 | 38 | 628 | 87 | 483 | 103 |
| Dhtkd1 | ENSMUSG00000025815 | 67 | 13 | 46 | 8 | 98 | 24 | 53 | 13 |
| Eml5 | ENSMUSG00000051166 | 129 | 41 | 106 | 30 | 156 | 42 | 80 | 35 |
| Fam84a | ENSMUSG00000020607 | 17 | 9 | 9 | 4 | 31 | 21 | 9 | 5 |
| Fam84b | ENSMUSG00000072568 | 367 | 45 | 332 | 38 | 416 | 37 | 306 | 21 |
| Fbxo3 | ENSMUSG00000027180 | 1943 | 149 | 1714 | 52 | 1982 | 135 | 1735 | 155 |
| Fgd4 | ENSMUSG00000022788 | 952 | 82 | 788 | 50 | 1006 | 124 | 808 | 116 |
| Fnip1 | ENSMUSG00000035992 | 1689 | 280 | 1464 | 146 | 1814 | 202 | 1491 | 143 |
| Gab1 | ENSMUSG00000031714 | 1294 | 55 | 1103 | 118 | 1362 | 98 | 1139 | 128 |
| Gm47283 | ENSMUSG00000096768 | 121 | 68 | 133 | 40 | 164 | 81 | 81 | 49 |
| Hbp1 | ENSMUSG00000002996 | 1726 | 184 | 1572 | 147 | 1798 | 134 | 1552 | 102 |
| Hccs | ENSMUSG00000031352 | 997 | 84 | 858 | 98 | 1041 | 148 | 890 | 60 |
| Hspa12a | ENSMUSG00000025092 | 841 | 91 | 689 | 87 | 938 | 90 | 730 | 91 |
| Jmy | ENSMUSG00000021690 | 1390 | 158 | 1200 | 90 | 1500 | 137 | 1234 | 114 |
| Kcnh2 | ENSMUSG00000038319 | 847 | 124 | 668 | 110 | 912 | 148 | 635 | 80 |
| Kcnj2 | ENSMUSG00000041695 | 1344 | 337 | 915 | 199 | 1453 | 205 | 945 | 234 |
| Laptm4b | ENSMUSG00000022257 | 953 | 46 | 801 | 57 | 990 | 41 | 803 | 90 |
| Lysmd4 | ENSMUSG00000043831 | 351 | 36 | 275 | 32 | 399 | 40 | 293 | 44 |
| Mef2a | ENSMUSG00000030557 | 4209 | 341 | 3602 | 223 | 4401 | 428 | 3617 | 183 |
| Mef2d | ENSMUSG00000001419 | 2886 | 195 | 2459 | 352 | 3285 | 449 | 2564 | 203 |
| Mif4gd | ENSMUSG00000020743 | 399 | 41 | 345 | 38 | 434 | 39 | 328 | 28 |
| mt-Nd2 | ENSMUSG00000064345 | 563587 | 132806 | 437281 | 99302 | 600917 | 101879 | 466929 | 126476 |
| mt-Nd5 | ENSMUSG00000064367 | 810634 | 162252 | 639879 | 106231 | 854483 | 106054 | 682450 | 132819 |
| Myo5c | ENSMUSG00000033590 | 102 | 18 | 90 | 31 | 150 | 35 | 66 | 21 |
| Nnt | ENSMUSG00000025453 | 5147 | 1691 | 3697 | 588 | 6332 | 1498 | 4488 | 661 |
| Osbpl1a | ENSMUSG00000044252 | 1430 | 127 | 1177 | 138 | 1477 | 68 | 1236 | 73 |
| Pkd2l2 | ENSMUSG00000014503 | 119 | 20 | 90 | 8 | 123 | 12 | 80 | 8 |
| Plag1 | ENSMUSG00000003282 | 111 | 11 | 86 | 16 | 110 | 13 | 71 | 14 |
| Plin5 | ENSMUSG00000011305 | 1427 | 167 | 1055 | 258 | 1589 | 193 | 1116 | 132 |
| Plxnb3 | ENSMUSG00000031385 | 47 | 14 | 32 | 7 | 53 | 9 | 26 | 4 |
| Pnrc1 | ENSMUSG00000040128 | 1394 | 186 | 1223 | 251 | 1670 | 151 | 1176 | 94 |
| Prox1 | ENSMUSG00000010175 | 2013 | 472 | 1483 | 298 | 2390 | 394 | 1685 | 308 |
| Ptpru | ENSMUSG00000028909 | 76 | 23 | 56 | 21 | 65 | 20 | 34 | 14 |
| Pygo1 | ENSMUSG00000034910 | 734 | 66 | 583 | 66 | 821 | 99 | 642 | 73 |
| Retreg1 | ENSMUSG00000022270 | 4914 | 645 | 4621 | 579 | 5580 | 346 | 4633 | 468 |
| Rorc | ENSMUSG00000028150 | 820 | 185 | 636 | 99 | 951 | 111 | 730 | 106 |
| Rsbn1l | ENSMUSG00000039968 | 486 | 32 | 495 | 68 | 547 | 61 | 451 | 62 |
| Sall4 | ENSMUSG00000027547 | 38 | 19 | 25 | 17 | 54 | 31 | 20 | 2 |
| Sc5d | ENSMUSG00000032018 | 453 | 51 | 403 | 59 | 480 | 34 | 405 | 25 |
| Slc25a22 | ENSMUSG00000019082 | 545 | 113 | 458 | 107 | 575 | 57 | 418 | 70 |
| Slc25a46 | ENSMUSG00000024259 | 1632 | 281 | 1428 | 165 | 1805 | 227 | 1528 | 184 |

| | | | | | | | | | |
|---|---|---|---|---|---|---|---|---|---|
| Slc40a1 | ENSMUSG00000025993 | 540 | 85 | 413 | 68 | 525 | 93 | 369 | 33 |
| Slc5a6 | ENSMUSG00000006641 | 210 | 13 | 153 | 42 | 237 | 30 | 148 | 21 |
| Slc9a2 | ENSMUSG00000026062 | 118 | 24 | 98 | 34 | 103 | 34 | 64 | 9 |
| Slf1 | ENSMUSG00000021597 | 700 | 128 | 591 | 68 | 734 | 132 | 561 | 98 |
| Smarcd1 | ENSMUSG00000023018 | 561 | 16 | 444 | 69 | 610 | 82 | 467 | 65 |
| Sobp | ENSMUSG00000038248 | 528 | 23 | 434 | 59 | 594 | 52 | 453 | 47 |
| Sorcs2 | ENSMUSG00000029093 | 323 | 58 | 246 | 26 | 327 | 75 | 216 | 35 |
| Sp4 | ENSMUSG00000025323 | 262 | 26 | 223 | 14 | 286 | 22 | 209 | 16 |
| Stk39 | ENSMUSG00000027030 | 2038 | 169 | 1731 | 250 | 2027 | 268 | 1564 | 136 |
| Tmem161a | ENSMUSG00000002342 | 695 | 40 | 602 | 58 | 708 | 61 | 579 | 56 |
| Tmem170b | ENSMUSG00000087370 | 544 | 52 | 444 | 44 | 603 | 78 | 462 | 26 |
| Tmem238 | ENSMUSG00000030431 | 17 | 16 | 16 | 5 | 31 | 9 | 10 | 4 |
| Tob1 | ENSMUSG00000037573 | 692 | 101 | 631 | 35 | 830 | 74 | 668 | 42 |
| Trmt2b | ENSMUSG00000067369 | 1295 | 120 | 1099 | 134 | 1300 | 53 | 1109 | 91 |
| Ttc30a1 | ENSMUSG00000075271 | 133 | 38 | 110 | 17 | 165 | 28 | 117 | 20 |
| Ttc30b | ENSMUSG00000075273 | 431 | 88 | 337 | 29 | 476 | 71 | 363 | 72 |
| Txnip | ENSMUSG00000038393 | 13919 | 3376 | 11722 | 2478 | 18708 | 3486 | 12475 | 1810 |
| Ubc | ENSMUSG00000008348 | 13447 | 4055 | 10872 | 1508 | 16818 | 2949 | 12228 | 2263 |
| Ube2b | ENSMUSG00000020390 | 4614 | 513 | 4036 | 428 | 4668 | 348 | 4142 | 342 |
| Ulk1 | ENSMUSG00000029512 | 1687 | 77 | 1424 | 156 | 1856 | 253 | 1452 | 92 |
| Vps13a | ENSMUSG00000046230 | 1184 | 126 | 1087 | 104 | 1354 | 142 | 1101 | 159 |
| Ythdf3 | ENSMUSG00000047213 | 1491 | 106 | 1342 | 61 | 1508 | 114 | 1305 | 117 |
| Zbtb18 | ENSMUSG00000063659 | 834 | 109 | 672 | 89 | 963 | 89 | 675 | 105 |
| Zfp292 | ENSMUSG00000039967 | 872 | 86 | 749 | 89 | 898 | 77 | 715 | 84 |

**Supplementary Table S10.** RNASeq data: gene clusters.

| Gene Symbol | Ensembl gene id | WT Vehicle | | WT AngII | | PKN2Het Vehicle | | PKN2Het AngII | |
|---|---|---|---|---|---|---|---|---|---|
| | | **Mean** | **SD** | **Mean** | **SD** | **Mean** | **SD** | **Mean** | **SD** |
| **Complement** | | | | | | | | | |
| C1qa | ENSMUSG00000036887 | 895 | 96 | 1523 | 250 | 883 | 97 | 1362 | 183 |
| C1qb | ENSMUSG00000036905 | 826 | 111 | 1513 | 429 | 782 | 128 | 1330 | 245 |
| C1qc | ENSMUSG00000036896 | 886 | 106 | 1454 | 199 | 845 | 131 | 1290 | 171 |
| C1qtnf3 | ENSMUSG00000058914 | 3 | 2 | 341 | 632 | 3 | 2 | 118 | 157 |
| C1qtnf5 | ENSMUSG00000079592 | 53 | 6 | 100 | 38 | 61 | 17 | 87 | 20 |
| C1qtnf6 | ENSMUSG00000022440 | 122 | 21 | 426 | 250 | 120 | 11 | 348 | 107 |
| C1qtnf7 | ENSMUSG00000061535 | 178 | 26 | 294 | 78 | 174 | 31 | 227 | 32 |
| C3ar1 | ENSMUSG00000040552 | 234 | 31 | 502 | 241 | 242 | 41 | 380 | 97 |
| C4b | ENSMUSG00000073418 | 231 | 28 | 604 | 141 | 295 | 41 | 861 | 703 |
| C5ar1 | ENSMUSG00000049130 | 107 | 17 | 173 | 40 | 78 | 18 | 156 | 56 |
| Cfb | ENSMUSG00000090231 | 36 | 7 | 140 | 61 | 33 | 15 | 114 | 78 |
| **Extracellular matrix** | | | | | | | | | |
| Acan | ENSMUSG00000030607 | 4 | 3 | 66 | 49 | 3 | 1 | 32 | 34 |
| Aspn | ENSMUSG00000021388 | 750 | 159 | 3970 | 5012 | 798 | 171 | 1825 | 1043 |
| Bgn | ENSMUSG00000031375 | 5175 | 520 | 12514 | 5386 | 5152 | 464 | 9491 | 2127 |
| Ccdc80 | ENSMUSG00000022665 | 1816 | 295 | 3479 | 1261 | 1843 | 168 | 2847 | 551 |
| Col11a1 | ENSMUSG00000027966 | 2 | 3 | 52 | 76 | 2 | 2 | 12 | 11 |
| Col12a1 | ENSMUSG00000032332 | 47 | 12 | 495 | 623 | 42 | 10 | 315 | 288 |
| Col14a1 | ENSMUSG00000022371 | 466 | 75 | 1493 | 1179 | 377 | 75 | 1075 | 556 |
| Col15a1 | ENSMUSG00000028339 | 2833 | 383 | 5668 | 1135 | 3175 | 509 | 5335 | 1392 |
| Col16a1 | ENSMUSG00000040690 | 187 | 39 | 603 | 471 | 192 | 60 | 420 | 186 |
| Col18a1 | ENSMUSG00000001435 | 248 | 28 | 790 | 244 | 293 | 35 | 541 | 181 |
| Col1a1 | ENSMUSG00000001506 | 1941 | 334 | 10243 | 9888 | 2115 | 63 | 5954 | 3420 |
| Col1a2 | ENSMUSG00000029661 | 2657 | 250 | 11812 | 10867 | 2786 | 231 | 7551 | 4300 |
| Col3a1 | ENSMUSG00000026043 | 5494 | 744 | 25906 | 20347 | 5871 | 824 | 16747 | 9657 |
| Col4a1 | ENSMUSG00000031502 | 12856 | 2211 | 23909 | 1751 | 13123 | 855 | 20837 | 3837 |
| Col4a2 | ENSMUSG00000031503 | 9490 | 1105 | 15270 | 591 | 9814 | 526 | 14603 | 2361 |
| Col4a3 | ENSMUSG00000079465 | 203 | 49 | 299 | 93 | 199 | 27 | 307 | 35 |
| Col4a4 | ENSMUSG00000067158 | 345 | 73 | 497 | 114 | 364 | 54 | 494 | 37 |
| Col4a5 | ENSMUSG00000031274 | 763 | 91 | 1159 | 304 | 773 | 91 | 971 | 147 |
| Col5a1 | ENSMUSG00000026837 | 1205 | 173 | 3410 | 1633 | 1238 | 85 | 2682 | 1054 |
| Col5a2 | ENSMUSG00000026042 | 878 | 65 | 4055 | 3237 | 927 | 105 | 2738 | 1563 |
| Col5a3 | ENSMUSG00000004098 | 576 | 178 | 985 | 63 | 666 | 103 | 847 | 222 |
| Col6a1 | ENSMUSG00000001119 | 2071 | 252 | 3943 | 1376 | 2047 | 155 | 3242 | 704 |
| Col6a2 | ENSMUSG00000020241 | 2038 | 291 | 3824 | 981 | 2062 | 139 | 3256 | 743 |
| Col6a3 | ENSMUSG00000048126 | 1170 | 165 | 2685 | 1137 | 1119 | 203 | 1895 | 452 |
| Col7a1 | ENSMUSG00000025650 | 4 | 6 | 26 | 21 | 5 | 4 | 11 | 6 |
| Col8a1 | ENSMUSG00000068196 | 704 | 96 | 3461 | 2170 | 747 | 153 | 2473 | 1081 |
| Col8a2 | ENSMUSG00000056174 | 9 | 5 | 180 | 298 | 14 | 4 | 86 | 85 |
| Col9a2 | ENSMUSG00000028626 | 6 | 1 | 31 | 38 | 7 | 4 | 21 | 11 |
| Comp | ENSMUSG00000031849 | 39 | 8 | 325 | 426 | 69 | 9 | 179 | 129 |
| Crtap | ENSMUSG00000032431 | 384 | 57 | 511 | 72 | 379 | 32 | 471 | 67 |
| Cthrc1 | ENSMUSG00000054196 | 3 | 3 | 221 | 380 | 5 | 2 | 92 | 112 |
| Ecm1 | ENSMUSG00000028108 | 452 | 86 | 696 | 127 | 464 | 82 | 720 | 136 |
| Egflam | ENSMUSG00000042961 | 374 | 53 | 240 | 63 | 279 | 55 | 262 | 40 |
| Eln | ENSMUSG00000029675 | 512 | 176 | 2249 | 1534 | 591 | 55 | 1327 | 606 |
| Fbln2 | ENSMUSG00000064080 | 1827 | 201 | 3200 | 493 | 1966 | 214 | 3008 | 422 |
| Fbn1 | ENSMUSG00000027204 | 2671 | 543 | 7690 | 2514 | 3011 | 303 | 6532 | 2277 |
| Fbn2 | ENSMUSG00000024598 | 25 | 8 | 112 | 120 | 38 | 10 | 67 | 32 |
| Fgl2 | ENSMUSG00000039899 | 863 | 65 | 1693 | 575 | 912 | 106 | 1331 | 391 |
| Flna | ENSMUSG00000031328 | 4014 | 455 | 5769 | 673 | 3783 | 774 | 4951 | 309 |
| Fn1 | ENSMUSG00000026193 | 1579 | 300 | 7830 | 6886 | 1494 | 199 | 4711 | 3367 |
| Frem1 | ENSMUSG00000059049 | 8 | 10 | 51 | 68 | 8 | 4 | 31 | 24 |
| Has2 | ENSMUSG00000022367 | 27 | 6 | 50 | 13 | 28 | 8 | 42 | 6 |
| Hspg2 | ENSMUSG00000028763 | 12506 | 1017 | 15279 | 1628 | 12610 | 707 | 14990 | 1190 |
| Lama2 | ENSMUSG00000019899 | 3507 | 352 | 3968 | 325 | 3362 | 180 | 4015 | 379 |

| | | | | | | | | | |
|---|---|---|---|---|---|---|---|---|---|
| Lama4 | ENSMUSG00000019846 | 2464 | 138 | 3226 | 192 | 2291 | 264 | 3050 | 270 |
| Lamb1 | ENSMUSG00000002900 | 2656 | 308 | 3500 | 191 | 2690 | 265 | 3282 | 467 |
| Lamc1 | ENSMUSG00000026478 | 4969 | 490 | 7157 | 380 | 5155 | 283 | 6688 | 486 |
| Lox | ENSMUSG00000024529 | 89 | 9 | 1050 | 1321 | 94 | 21 | 476 | 436 |
| Loxl1 | ENSMUSG00000032334 | 618 | 97 | 1447 | 604 | 687 | 57 | 1181 | 257 |
| Loxl2 | ENSMUSG00000034205 | 724 | 133 | 1635 | 127 | 683 | 84 | 1403 | 383 |
| Loxl3 | ENSMUSG00000000693 | 104 | 36 | 311 | 233 | 87 | 12 | 196 | 70 |
| Lum | ENSMUSG00000036446 | 1447 | 238 | 3824 | 2686 | 1553 | 192 | 2795 | 564 |
| Matn2 | ENSMUSG00000022324 | 417 | 57 | 667 | 192 | 429 | 35 | 559 | 39 |
| Mfap2 | ENSMUSG00000060572 | 20 | 2 | 69 | 61 | 23 | 4 | 41 | 14 |
| Mfap3 | ENSMUSG00000020522 | 454 | 25 | 541 | 33 | 431 | 20 | 503 | 22 |
| Mfap3l | ENSMUSG00000031647 | 512 | 38 | 351 | 27 | 511 | 51 | 378 | 15 |
| Mfap4 | ENSMUSG00000042436 | 202 | 18 | 1249 | 1153 | 245 | 36 | 712 | 324 |
| Mfap5 | ENSMUSG00000030116 | 411 | 43 | 1634 | 1033 | 404 | 45 | 1132 | 430 |
| Mgp | ENSMUSG00000030218 | 1671 | 257 | 3299 | 1832 | 1627 | 149 | 2424 | 267 |
| Mxra7 | ENSMUSG00000020814 | 353 | 43 | 549 | 114 | 334 | 33 | 485 | 81 |
| Mxra8 | ENSMUSG00000029070 | 431 | 61 | 782 | 329 | 511 | 145 | 714 | 119 |
| Nid1 | ENSMUSG00000005397 | 3926 | 530 | 6347 | 1387 | 3634 | 332 | 5323 | 725 |
| Nid2 | ENSMUSG00000021806 | 561 | 130 | 914 | 118 | 537 | 62 | 895 | 114 |
| Ntn1 | ENSMUSG00000020902 | 1895 | 146 | 1435 | 153 | 2064 | 197 | 1793 | 251 |
| P3h1 | ENSMUSG00000028641 | 217 | 25 | 313 | 43 | 242 | 18 | 308 | 55 |
| P3h3 | ENSMUSG00000023191 | 185 | 14 | 319 | 100 | 179 | 28 | 278 | 54 |
| P4ha3 | ENSMUSG00000051048 | 2 | 2 | 23 | 32 | 1 | 1 | 11 | 10 |
| Pcolce | ENSMUSG00000029718 | 778 | 81 | 1236 | 178 | 675 | 91 | 1133 | 173 |
| Pcolce2 | ENSMUSG00000015354 | 412 | 82 | 494 | 81 | 352 | 55 | 503 | 72 |
| Plod3 | ENSMUSG00000004846 | 475 | 118 | 658 | 87 | 472 | 51 | 605 | 62 |
| Postn | ENSMUSG00000027750 | 1288 | 354 | 16369 | 19036 | 1325 | 317 | 10262 | 8108 |
| Prg4 | ENSMUSG00000006014 | 323 | 89 | 484 | 130 | 321 | 33 | 596 | 281 |
| Sparc | ENSMUSG00000018593 | 7512 | 635 | 18831 | 6579 | 7355 | 933 | 14409 | 2942 |
| Vcan | ENSMUSG00000021614 | 730 | 115 | 1503 | 151 | 665 | 105 | 1354 | 355 |
| **Interferon signalling** | | | | | | | | | |
| Ifi203 | ENSMUSG00000039997 | 891 | 161 | 1230 | 255 | 880 | 117 | 940 | 84 |
| Ifi204 | ENSMUSG00000073489 | 206 | 34 | 433 | 95 | 196 | 40 | 334 | 43 |
| Ifi209 | ENSMUSG00000043263 | 52 | 15 | 126 | 69 | 50 | 17 | 92 | 45 |
| Ifi211 | ENSMUSG00000026536 | 185 | 36 | 351 | 104 | 178 | 18 | 256 | 21 |
| Ifi27l2a | ENSMUSG00000079017 | 237 | 35 | 439 | 84 | 218 | 39 | 420 | 49 |
| Ifi30 | ENSMUSG00000031838 | 101 | 27 | 193 | 76 | 87 | 20 | 127 | 31 |
| Ifih1 | ENSMUSG00000026896 | 299 | 32 | 446 | 46 | 329 | 56 | 378 | 53 |
| Ifit1 | ENSMUSG00000034459 | 187 | 43 | 329 | 102 | 161 | 24 | 234 | 21 |
| Ifit2 | ENSMUSG00000045932 | 547 | 92 | 934 | 140 | 539 | 58 | 669 | 52 |
| Ifit3 | ENSMUSG00000074896 | 336 | 93 | 626 | 210 | 311 | 45 | 430 | 29 |
| Ifit3b | ENSMUSG00000062488 | 122 | 30 | 208 | 57 | 123 | 22 | 150 | 9 |
| Ifitm2 | ENSMUSG00000060591 | 875 | 131 | 1176 | 80 | 841 | 117 | 1097 | 135 |
| Ifitm3 | ENSMUSG00000025492 | 1226 | 258 | 1689 | 136 | 1126 | 147 | 1515 | 182 |
| Ifngr1 | ENSMUSG00000020009 | 1109 | 126 | 1400 | 78 | 1101 | 128 | 1324 | 109 |
| Irf5 | ENSMUSG00000029771 | 73 | 13 | 148 | 22 | 78 | 12 | 119 | 35 |
| Irf7 | ENSMUSG00000025498 | 220 | 65 | 413 | 114 | 189 | 26 | 332 | 52 |
| Irf8 | ENSMUSG00000041515 | 118 | 13 | 229 | 76 | 128 | 21 | 193 | 26 |
| Isg20 | ENSMUSG00000039236 | 86 | 30 | 130 | 16 | 102 | 27 | 144 | 32 |
| **Mitochondria** | | | | | | | | | |
| Acaa2 | ENSMUSG00000036880 | 13252 | 1380 | 8259 | 2391 | 12324 | 1059 | 9258 | 1215 |
| Aco2 | ENSMUSG00000022477 | 41047 | 2512 | 30674 | 4778 | 41075 | 2487 | 35785 | 2269 |
| Afg1l | ENSMUSG00000038302 | 1113 | 135 | 844 | 176 | 1027 | 95 | 958 | 51 |
| Aldh2 | ENSMUSG00000029455 | 3628 | 278 | 2621 | 303 | 3753 | 239 | 3023 | 178 |
| Atp5a1 | ENSMUSG00000025428 | 73890 | 5007 | 52519 | 8597 | 73310 | 4969 | 61155 | 3605 |
| Atp5b | ENSMUSG00000025393 | 81983 | 4027 | 60525 | 7907 | 80175 | 3926 | 68956 | 5579 |
| Atp5d | ENSMUSG00000003072 | 5810 | 313 | 4470 | 769 | 5730 | 539 | 5074 | 535 |
| Atp5e | ENSMUSG00000016252 | 4682 | 460 | 3624 | 408 | 4707 | 229 | 4130 | 508 |
| Atp5g3 | ENSMUSG00000018770 | 18004 | 1331 | 13448 | 2291 | 18271 | 996 | 15327 | 1271 |
| Atp5o | ENSMUSG00000022956 | 14204 | 841 | 11103 | 1882 | 13615 | 536 | 12332 | 665 |
| Bcat2 | ENSMUSG00000030826 | 1208 | 96 | 917 | 126 | 1160 | 104 | 1012 | 77 |
| Cars2 | ENSMUSG00000056228 | 540 | 40 | 408 | 58 | 563 | 48 | 458 | 53 |

| | | | | | | | | | |
|---|---|---|---|---|---|---|---|---|---|
| Clpp | ENSMUSG00000002660 | 608 | 48 | 457 | 72 | 598 | 7 | 499 | 95 |
| Clpx | ENSMUSG00000015357 | 2210 | 180 | 1776 | 257 | 2247 | 186 | 1932 | 167 |
| Cluh | ENSMUSG00000020741 | 5745 | 385 | 4121 | 818 | 6093 | 799 | 5032 | 263 |
| Coq10a | ENSMUSG00000039914 | 3593 | 171 | 2600 | 450 | 3601 | 150 | 3083 | 271 |
| Coq2 | ENSMUSG00000029319 | 1358 | 71 | 1066 | 109 | 1353 | 65 | 1154 | 62 |
| Coq7 | ENSMUSG00000030652 | 1122 | 100 | 786 | 155 | 1106 | 99 | 884 | 88 |
| Coq8a | ENSMUSG00000026489 | 8867 | 811 | 6241 | 1021 | 9606 | 1491 | 7168 | 332 |
| Coq9 | ENSMUSG00000031782 | 6190 | 445 | 4627 | 906 | 6154 | 410 | 5544 | 503 |
| Cox4i1 | ENSMUSG00000031818 | 22284 | 812 | 17754 | 1611 | 22070 | 1141 | 19696 | 1055 |
| Cox5a | ENSMUSG00000000088 | 11595 | 608 | 8951 | 1587 | 11127 | 414 | 10060 | 740 |
| Cox5b | ENSMUSG00000061518 | 10424 | 672 | 8385 | 1343 | 10221 | 301 | 9039 | 577 |
| Cox7a1 | ENSMUSG00000074218 | 8954 | 697 | 6119 | 1540 | 8610 | 451 | 7040 | 827 |
| Cox8b | ENSMUSG00000025488 | 6504 | 621 | 4961 | 1023 | 6394 | 477 | 5556 | 903 |
| Cs | ENSMUSG00000005683 | 24899 | 2214 | 17482 | 3393 | 24790 | 1691 | 20831 | 1885 |
| Decr1 | ENSMUSG00000028223 | 8479 | 863 | 5924 | 1566 | 7835 | 344 | 6512 | 520 |
| Diablo | ENSMUSG00000029433 | 1178 | 83 | 980 | 91 | 1146 | 35 | 1105 | 48 |
| Dlst | ENSMUSG00000004789 | 11654 | 750 | 8617 | 1895 | 11270 | 555 | 10153 | 788 |
| Echs1 | ENSMUSG00000025465 | 4828 | 268 | 3779 | 465 | 4896 | 284 | 4167 | 357 |
| Etfa | ENSMUSG00000032314 | 11167 | 800 | 7804 | 1484 | 10775 | 533 | 8929 | 812 |
| Etfb | ENSMUSG00000004610 | 8827 | 431 | 6069 | 1637 | 8450 | 419 | 6927 | 725 |
| Etfdh | ENSMUSG00000027809 | 13588 | 811 | 9757 | 2314 | 12812 | 772 | 10703 | 771 |
| Fh1 | ENSMUSG00000026526 | 7500 | 576 | 5659 | 1050 | 7018 | 180 | 6388 | 586 |
| Fmc1 | ENSMUSG00000019689 | 622 | 55 | 458 | 87 | 580 | 55 | 484 | 71 |
| Gfm1 | ENSMUSG00000027774 | 4053 | 240 | 3035 | 442 | 3973 | 430 | 3372 | 259 |
| Gfm2 | ENSMUSG00000021666 | 1266 | 144 | 1001 | 118 | 1229 | 127 | 1128 | 100 |
| Got2 | ENSMUSG00000031672 | 15403 | 947 | 11510 | 1822 | 15607 | 1320 | 13207 | 1045 |
| Gpd2 | ENSMUSG00000026827 | 401 | 24 | 334 | 32 | 392 | 48 | 398 | 25 |
| Hccs | ENSMUSG00000031352 | 997 | 84 | 858 | 98 | 1041 | 148 | 890 | 60 |
| Iars2 | ENSMUSG00000026618 | 2409 | 99 | 1873 | 136 | 2398 | 100 | 2171 | 120 |
| Idh2 | ENSMUSG00000030541 | 25832 | 1566 | 18548 | 3665 | 26906 | 2049 | 21457 | 2197 |
| Idh3a | ENSMUSG00000032279 | 10211 | 1028 | 7707 | 867 | 10029 | 629 | 8753 | 799 |
| Idh3b | ENSMUSG00000027406 | 11455 | 1144 | 8621 | 1973 | 11227 | 459 | 9312 | 738 |
| Idh3g | ENSMUSG00000002010 | 6376 | 459 | 4785 | 580 | 6418 | 422 | 4954 | 479 |
| Immt | ENSMUSG00000052337 | 11137 | 723 | 8672 | 1433 | 10924 | 555 | 9873 | 547 |
| Lonp1 | ENSMUSG00000041168 | 2317 | 80 | 2025 | 163 | 2376 | 150 | 2252 | 87 |
| Malsu1 | ENSMUSG00000029815 | 281 | 28 | 208 | 15 | 260 | 23 | 236 | 30 |
| Mars2 | ENSMUSG00000046994 | 370 | 47 | 284 | 34 | 364 | 34 | 304 | 25 |
| Mcub | ENSMUSG00000027994 | 41 | 14 | 77 | 15 | 37 | 4 | 50 | 10 |
| Mdh2 | ENSMUSG00000019179 | 16876 | 703 | 12970 | 2129 | 16369 | 1034 | 14499 | 734 |
| Me3 | ENSMUSG00000030621 | 1244 | 36 | 938 | 86 | 1271 | 81 | 1046 | 52 |
| Mfn1 | ENSMUSG00000027668 | 8217 | 628 | 6382 | 1129 | 8242 | 170 | 7059 | 753 |
| Mfn2 | ENSMUSG00000029020 | 15414 | 955 | 11184 | 2042 | 15810 | 1644 | 13241 | 705 |
| Mgme1 | ENSMUSG00000027424 | 350 | 28 | 255 | 46 | 337 | 12 | 267 | 30 |
| Miga1 | ENSMUSG00000054942 | 412 | 61 | 323 | 8 | 409 | 40 | 383 | 35 |
| Mipep | ENSMUSG00000021993 | 1364 | 63 | 983 | 167 | 1394 | 126 | 1103 | 71 |
| Mpv17 | ENSMUSG00000107283 | 1098 | 32 | 873 | 90 | 1118 | 80 | 934 | 104 |
| Mrpl14 | ENSMUSG00000023939 | 682 | 54 | 516 | 77 | 739 | 35 | 586 | 87 |
| Mrpl16 | ENSMUSG00000024683 | 961 | 36 | 784 | 75 | 930 | 52 | 848 | 97 |
| Mrpl28 | ENSMUSG00000024181 | 1210 | 39 | 882 | 132 | 1171 | 81 | 1020 | 121 |
| Mrpl37 | ENSMUSG00000028622 | 1184 | 43 | 904 | 135 | 1161 | 34 | 1046 | 74 |
| Mrpl38 | ENSMUSG00000020775 | 704 | 31 | 534 | 72 | 674 | 41 | 611 | 58 |
| Mrpl39 | ENSMUSG00000022889 | 1365 | 91 | 1053 | 180 | 1335 | 40 | 1087 | 56 |
| Mrpl4 | ENSMUSG00000003299 | 1028 | 38 | 792 | 138 | 1029 | 93 | 915 | 137 |
| Mrpl45 | ENSMUSG00000018882 | 1187 | 97 | 837 | 153 | 1100 | 58 | 1043 | 54 |
| Mrps26 | ENSMUSG00000037740 | 355 | 39 | 258 | 24 | 351 | 18 | 326 | 26 |
| Mrps35 | ENSMUSG00000040112 | 1312 | 76 | 991 | 219 | 1236 | 59 | 1132 | 63 |
| Mrps6 | ENSMUSG00000039680 | 361 | 20 | 285 | 16 | 354 | 19 | 300 | 4 |
| mt-Nd2 | ENSMUSG00000064345 | 563587 | 132806 | 437281 | 99302 | 600917 | 101879 | 466929 | 126476 |
| mt-Nd5 | ENSMUSG00000064367 | 810634 | 162252 | 639879 | 106231 | 854483 | 106054 | 682450 | 132819 |
| mt-Rnr2 | ENSMUSG00000064339 | 356240 | 61989 | 247940 | 69255 | 362883 | 53161 | 276407 | 38517 |
| Mtfp1 | ENSMUSG00000004748 | 1636 | 82 | 1103 | 322 | 1626 | 114 | 1320 | 90 |
| Mtfr1l | ENSMUSG00000046671 | 2563 | 65 | 2169 | 189 | 2622 | 122 | 2390 | 105 |
| Mtg2 | ENSMUSG00000039069 | 366 | 31 | 275 | 26 | 387 | 9 | 312 | 16 |
| Mto1 | ENSMUSG00000032342 | 481 | 27 | 368 | 46 | 450 | 40 | 405 | 41 |

| | | | | | | | | | |
|---|---|---|---|---|---|---|---|---|---|
| Nadk2 | ENSMUSG00000022253 | 1247 | 102 | 792 | 168 | 1254 | 222 | 893 | 160 |
| Ndufa10 | ENSMUSG00000026260 | 9566 | 706 | 7436 | 1132 | 9920 | 331 | 8450 | 886 |
| Ndufa8 | ENSMUSG00000026895 | 4878 | 294 | 3771 | 635 | 4757 | 148 | 4279 | 331 |
| Ndufa9 | ENSMUSG00000000399 | 9480 | 996 | 7084 | 1337 | 9287 | 369 | 7867 | 703 |
| Ndufs1 | ENSMUSG00000025968 | 15975 | 1815 | 11734 | 2163 | 15650 | 1579 | 13236 | 1027 |
| Ndufs2 | ENSMUSG00000013593 | 16031 | 729 | 11634 | 1773 | 15858 | 991 | 13476 | 1336 |
| Ndufs3 | ENSMUSG00000005510 | 6047 | 349 | 4620 | 872 | 5917 | 121 | 5197 | 447 |
| Ndufs7 | ENSMUSG00000020153 | 3525 | 187 | 2485 | 513 | 3595 | 317 | 2923 | 370 |
| Ndufv1 | ENSMUSG00000037916 | 9773 | 762 | 6974 | 1216 | 9669 | 615 | 8148 | 626 |
| Ogdh | ENSMUSG00000020456 | 39640 | 2492 | 29734 | 4328 | 41635 | 5116 | 35518 | 902 |
| Opa1 | ENSMUSG00000038084 | 5644 | 514 | 4210 | 706 | 5735 | 550 | 5001 | 562 |
| Oxa1l | ENSMUSG00000000959 | 1341 | 57 | 1036 | 150 | 1356 | 195 | 1142 | 95 |
| Oxct1 | ENSMUSG00000022186 | 26538 | 1570 | 20521 | 2412 | 25456 | 2358 | 23051 | 1272 |
| Oxsm | ENSMUSG00000021786 | 592 | 47 | 470 | 26 | 633 | 71 | 520 | 20 |
| Pdf | ENSMUSG00000078931 | 565 | 42 | 418 | 72 | 575 | 29 | 492 | 19 |
| Pdha1 | ENSMUSG00000031299 | 24973 | 2883 | 19882 | 1983 | 24844 | 2183 | 21298 | 1610 |
| Pdk2 | ENSMUSG00000038967 | 7045 | 426 | 4922 | 991 | 7351 | 661 | 5815 | 537 |
| Pdk3 | ENSMUSG00000035232 | 44 | 14 | 83 | 17 | 53 | 21 | 69 | 7 |
| Pdp1 | ENSMUSG00000049225 | 1052 | 157 | 827 | 97 | 1038 | 87 | 873 | 99 |
| Pdp2 | ENSMUSG00000048371 | 822 | 182 | 433 | 87 | 839 | 116 | 560 | 111 |
| Pdpr | ENSMUSG00000033624 | 2409 | 176 | 1771 | 254 | 2523 | 330 | 2118 | 147 |
| Pmpca | ENSMUSG00000026926 | 2809 | 172 | 2292 | 228 | 2800 | 157 | 2538 | 100 |
| Sdha | ENSMUSG00000021577 | 26565 | 1713 | 17836 | 2991 | 26775 | 2129 | 20539 | 1624 |
| Sdhb | ENSMUSG00000009863 | 12188 | 749 | 9065 | 1722 | 11819 | 503 | 9978 | 990 |
| Sdhc | ENSMUSG00000058076 | 7127 | 66 | 5010 | 729 | 7101 | 434 | 5730 | 395 |
| Sfxn3 | ENSMUSG00000025212 | 216 | 19 | 310 | 29 | 224 | 5 | 272 | 31 |
| Slc25a11 | ENSMUSG00000014606 | 8077 | 561 | 5928 | 1242 | 7992 | 341 | 6986 | 662 |
| Slc25a12 | ENSMUSG00000027010 | 4184 | 351 | 2893 | 395 | 4194 | 379 | 3611 | 151 |
| Slc25a13 | ENSMUSG00000015112 | 2989 | 324 | 2199 | 145 | 3029 | 276 | 2574 | 268 |
| Slc25a20 | ENSMUSG00000032602 | 3086 | 349 | 2262 | 477 | 3063 | 140 | 2521 | 210 |
| Slc25a22 | ENSMUSG00000019082 | 545 | 113 | 458 | 107 | 575 | 57 | 418 | 70 |
| Slc25a24 | ENSMUSG00000040322 | 184 | 16 | 278 | 63 | 197 | 22 | 246 | 35 |
| Slc25a3 | ENSMUSG00000061904 | 25015 | 1211 | 18063 | 2479 | 25124 | 1400 | 20806 | 1485 |
| Sod2 | ENSMUSG00000006818 | 12266 | 831 | 8665 | 1885 | 11982 | 506 | 9979 | 664 |
| Suclg1 | ENSMUSG00000052738 | 6364 | 514 | 4599 | 1021 | 6012 | 279 | 5327 | 363 |
| Suclg2 | ENSMUSG00000061838 | 4105 | 321 | 2961 | 428 | 4037 | 119 | 3294 | 228 |
| Tars2 | ENSMUSG00000028107 | 667 | 59 | 489 | 66 | 606 | 23 | 549 | 66 |
| Tcaim | ENSMUSG00000046603 | 1091 | 140 | 752 | 164 | 1078 | 129 | 865 | 134 |
| Tufm | ENSMUSG00000073838 | 3132 | 215 | 2311 | 479 | 3147 | 191 | 2714 | 265 |
| Twnk | ENSMUSG00000025209 | 604 | 21 | 452 | 67 | 607 | 59 | 505 | 23 |
| Ucp3 | ENSMUSG00000032942 | 1264 | 355 | 659 | 227 | 1281 | 331 | 698 | 271 |
| Uqcc1 | ENSMUSG00000005882 | 3454 | 270 | 2476 | 553 | 3363 | 150 | 2909 | 228 |
| Uqcrc1 | ENSMUSG00000025651 | 15130 | 710 | 10190 | 2080 | 15168 | 907 | 12233 | 1293 |
| Uqcrfs1 | ENSMUSG00000038462 | 13197 | 914 | 10056 | 1838 | 12772 | 692 | 11183 | 890 |
| Vdac1 | ENSMUSG00000020402 | 17517 | 801 | 13326 | 1578 | 17010 | 633 | 14751 | 342 |
| Vdac3 | ENSMUSG00000008892 | 6247 | 446 | 4930 | 675 | 5939 | 119 | 5490 | 330 |
| **Structural** | | | | | | | | | |
| Acta1 | ENSMUSG00000031972 | 6773 | 1395 | 24772 | 10833 | 8570 | 3863 | 28601 | 14894 |
| Actb | ENSMUSG00000029580 | 7471 | 1804 | 10178 | 1677 | 7763 | 1384 | 9365 | 1200 |
| Actg1 | ENSMUSG00000062825 | 6824 | 1047 | 8916 | 1194 | 7413 | 1015 | 8405 | 1266 |
| Actg2 | ENSMUSG00000059430 | 11 | 6 | 37 | 14 | 12 | 6 | 20 | 12 |
| Actn1 | ENSMUSG00000015143 | 522 | 79 | 925 | 273 | 524 | 61 | 761 | 132 |
| Actn4 | ENSMUSG00000054808 | 2213 | 272 | 2742 | 211 | 2012 | 203 | 2417 | 171 |
| Actr2 | ENSMUSG00000020152 | 2099 | 200 | 2493 | 267 | 2101 | 71 | 2352 | 201 |
| Actr3 | ENSMUSG00000026341 | 2536 | 120 | 3001 | 264 | 2399 | 43 | 2908 | 199 |
| Agrn | ENSMUSG00000041936 | 958 | 125 | 1295 | 154 | 947 | 79 | 1136 | 70 |
| Aif1 | ENSMUSG00000024397 | 63 | 12 | 133 | 73 | 51 | 11 | 95 | 13 |
| Ank2 | ENSMUSG00000032826 | 3236 | 377 | 2426 | 262 | 3245 | 406 | 2934 | 430 |
| Ankrd1 | ENSMUSG00000024803 | 20150 | 6483 | 57246 | 17002 | 19310 | 6244 | 49450 | 10874 |
| Ankrd23 | ENSMUSG00000067653 | 8223 | 1670 | 13521 | 3447 | 9104 | 579 | 14731 | 4085 |
| Anks1 | ENSMUSG00000024219 | 1418 | 123 | 1011 | 201 | 1347 | 177 | 1118 | 141 |
| Anln | ENSMUSG00000036777 | 56 | 22 | 225 | 55 | 56 | 7 | 174 | 50 |
| Arpc1b | ENSMUSG00000029622 | 814 | 81 | 1186 | 179 | 737 | 113 | 1071 | 150 |

| | | | | | | | | | |
|---|---|---|---|---|---|---|---|---|---|
| Arpc2 | ENSMUSG00000006304 | 2337 | 43 | 2757 | 185 | 2338 | 47 | 2652 | 121 |
| Arpc3 | ENSMUSG00000029465 | 1134 | 39 | 1490 | 168 | 1115 | 115 | 1392 | 88 |
| Arpc5 | ENSMUSG00000008475 | 864 | 45 | 1233 | 156 | 871 | 66 | 1121 | 40 |
| Arpin | ENSMUSG00000039043 | 239 | 19 | 299 | 19 | 258 | 25 | 280 | 26 |
| Cald1 | ENSMUSG00000029761 | 1649 | 323 | 2616 | 191 | 1624 | 256 | 2205 | 422 |
| Capg | ENSMUSG00000056737 | 183 | 29 | 364 | 36 | 182 | 36 | 328 | 70 |
| Capza1 | ENSMUSG00000070372 | 1251 | 78 | 1608 | 138 | 1166 | 148 | 1428 | 124 |
| Cfl1 | ENSMUSG00000056201 | 2052 | 87 | 2670 | 304 | 1872 | 236 | 2490 | 204 |
| Cilp | ENSMUSG00000042254 | 294 | 76 | 3092 | 3273 | 401 | 134 | 2249 | 1515 |
| Ckap2 | ENSMUSG00000037725 | 33 | 16 | 131 | 46 | 25 | 5 | 95 | 45 |
| Ckap2l | ENSMUSG00000048327 | 30 | 10 | 123 | 39 | 25 | 8 | 110 | 29 |
| Ckap4 | ENSMUSG00000046841 | 756 | 159 | 1094 | 75 | 815 | 103 | 1104 | 165 |
| Cnn1 | ENSMUSG00000001349 | 37 | 23 | 130 | 75 | 42 | 17 | 53 | 24 |
| Cnn2 | ENSMUSG00000004665 | 793 | 93 | 1039 | 52 | 856 | 80 | 1002 | 98 |
| Cnn3 | ENSMUSG00000053931 | 1009 | 125 | 1438 | 123 | 955 | 121 | 1319 | 119 |
| Coro1a | ENSMUSG00000030707 | 175 | 29 | 270 | 73 | 171 | 24 | 228 | 26 |
| Coro1b | ENSMUSG00000024835 | 688 | 47 | 858 | 53 | 695 | 47 | 802 | 76 |
| Cotl1 | ENSMUSG00000031827 | 234 | 40 | 369 | 66 | 208 | 50 | 335 | 22 |
| Csrp1 | ENSMUSG00000026421 | 868 | 80 | 1281 | 109 | 889 | 121 | 1075 | 145 |
| Csrp2 | ENSMUSG00000020186 | 180 | 37 | 542 | 354 | 165 | 20 | 315 | 127 |
| Cttn | ENSMUSG00000031078 | 793 | 94 | 1056 | 113 | 811 | 56 | 962 | 95 |
| Dbn1 | ENSMUSG00000034675 | 170 | 35 | 385 | 71 | 169 | 25 | 330 | 74 |
| Dbnl | ENSMUSG00000020476 | 472 | 46 | 575 | 30 | 501 | 31 | 573 | 37 |
| Emilin1 | ENSMUSG00000029163 | 473 | 106 | 835 | 204 | 519 | 28 | 812 | 91 |
| Eml2 | ENSMUSG00000040811 | 379 | 40 | 285 | 35 | 408 | 44 | 337 | 30 |
| Eml5 | ENSMUSG00000051166 | 129 | 41 | 106 | 30 | 156 | 42 | 80 | 35 |
| Enah | ENSMUSG00000022995 | 2820 | 352 | 4077 | 860 | 3138 | 213 | 4243 | 467 |
| Fscn1 | ENSMUSG00000029581 | 817 | 47 | 1186 | 92 | 831 | 115 | 1140 | 241 |
| Ift122 | ENSMUSG00000030323 | 278 | 32 | 496 | 94 | 309 | 39 | 488 | 78 |
| Ift81 | ENSMUSG00000029469 | 1002 | 129 | 666 | 78 | 994 | 49 | 708 | 78 |
| Jpt1 | ENSMUSG00000020737 | 841 | 76 | 987 | 49 | 790 | 37 | 1002 | 107 |
| Jpt2 | ENSMUSG00000024165 | 169 | 19 | 197 | 35 | 141 | 33 | 226 | 44 |
| Map10 | ENSMUSG00000050930 | 149 | 21 | 98 | 15 | 151 | 17 | 112 | 10 |
| Map1b | ENSMUSG00000052727 | 582 | 102 | 873 | 51 | 444 | 82 | 777 | 113 |
| Map1lc3a | ENSMUSG00000027602 | 4603 | 222 | 3864 | 305 | 4713 | 378 | 4049 | 447 |
| Map6 | ENSMUSG00000055407 | 108 | 31 | 142 | 36 | 87 | 9 | 129 | 34 |
| Mapre1 | ENSMUSG00000027479 | 1639 | 151 | 1876 | 92 | 1651 | 91 | 1859 | 64 |
| Msn | ENSMUSG00000031207 | 4381 | 327 | 6174 | 266 | 4286 | 400 | 5734 | 498 |
| Mybpc2 | ENSMUSG00000038670 | 292 | 36 | 901 | 354 | 321 | 110 | 773 | 309 |
| Myh10 | ENSMUSG00000020900 | 964 | 132 | 1462 | 186 | 988 | 96 | 1366 | 213 |
| Myh14 | ENSMUSG00000030739 | 1825 | 190 | 1436 | 214 | 1992 | 381 | 1613 | 123 |
| Myh6 | ENSMUSG00000040752 | 416670 | 31586 | 295124 | 67491 | 432469 | 60382 | 365274 | 31724 |
| Myh7 | ENSMUSG00000053093 | 1672 | 510 | 20844 | 27743 | 2490 | 1006 | 11438 | 8811 |
| Myh9 | ENSMUSG00000022443 | 4467 | 917 | 6229 | 813 | 4465 | 679 | 5733 | 609 |
| Myl1 | ENSMUSG00000061816 | 777 | 84 | 1354 | 639 | 674 | 113 | 1186 | 282 |
| Myl6 | ENSMUSG00000090841 | 3039 | 174 | 4606 | 743 | 2959 | 365 | 4098 | 492 |
| Myl9 | ENSMUSG00000067818 | 507 | 113 | 744 | 105 | 504 | 152 | 536 | 41 |
| Mylip | ENSMUSG00000038175 | 408 | 59 | 322 | 55 | 361 | 21 | 336 | 42 |
| Myo1c | ENSMUSG00000017774 | 2939 | 140 | 3445 | 280 | 2807 | 136 | 3414 | 203 |
| Myo1d | ENSMUSG00000035441 | 367 | 44 | 489 | 47 | 360 | 56 | 474 | 39 |
| Myo1e | ENSMUSG00000032220 | 382 | 52 | 541 | 82 | 377 | 40 | 487 | 59 |
| Myo1f | ENSMUSG00000024300 | 97 | 11 | 170 | 34 | 88 | 10 | 163 | 43 |
| Myo1g | ENSMUSG00000020437 | 50 | 9 | 89 | 21 | 55 | 12 | 90 | 33 |
| Myo5a | ENSMUSG00000034593 | 359 | 55 | 615 | 165 | 371 | 102 | 518 | 152 |
| Myo5c | ENSMUSG00000033590 | 102 | 18 | 90 | 31 | 150 | 35 | 66 | 21 |
| Myof | ENSMUSG00000048612 | 346 | 89 | 640 | 167 | 312 | 56 | 541 | 78 |
| Nes | ENSMUSG00000004891 | 1779 | 216 | 2677 | 688 | 1621 | 280 | 2378 | 347 |
| Pfn1 | ENSMUSG00000018293 | 2349 | 50 | 2712 | 75 | 2362 | 104 | 2642 | 191 |
| Sgcb | ENSMUSG00000029156 | 2725 | 146 | 2351 | 41 | 2817 | 190 | 2512 | 150 |
| Sgce | ENSMUSG00000004631 | 294 | 28 | 387 | 53 | 287 | 38 | 342 | 18 |
| Smarcd1 | ENSMUSG00000023018 | 561 | 16 | 444 | 69 | 610 | 82 | 467 | 65 |
| Sntb2 | ENSMUSG00000041308 | 632 | 76 | 913 | 145 | 638 | 68 | 853 | 99 |
| Sptb | ENSMUSG00000021061 | 3814 | 253 | 2892 | 391 | 4015 | 448 | 3358 | 226 |
| Tagln | ENSMUSG00000032085 | 483 | 160 | 931 | 216 | 458 | 122 | 542 | 128 |

| | | | | | | | | | |
|---|---|---|---|---|---|---|---|---|---|
| Tagln2 | ENSMUSG00000026547 | 1222 | 205 | 1863 | 181 | 1131 | 206 | 1710 | 250 |
| Tcap | ENSMUSG00000007877 | 15023 | 5676 | 9524 | 3481 | 11808 | 3513 | 10365 | 2562 |
| Tln1 | ENSMUSG00000028465 | 3433 | 355 | 4129 | 304 | 3528 | 301 | 4361 | 274 |
| Tmod1 | ENSMUSG00000028328 | 6866 | 511 | 5456 | 542 | 6671 | 424 | 6180 | 413 |
| Tmod4 | ENSMUSG00000005628 | 378 | 37 | 254 | 60 | 361 | 43 | 317 | 59 |
| Tnni3 | ENSMUSG00000035458 | 63108 | 4479 | 42722 | 10485 | 61514 | 1321 | 48873 | 5226 |
| Tnnt2 | ENSMUSG00000026414 | 110161 | 8926 | 87055 | 9068 | 105408 | 3834 | 92241 | 8335 |
| Tnnt3 | ENSMUSG00000061723 | 6 | 2 | 29 | 39 | 5 | 3 | 17 | 17 |
| Tpm2 | ENSMUSG00000028464 | 338 | 60 | 565 | 58 | 358 | 58 | 477 | 79 |
| Tpm3 | ENSMUSG00000027940 | 1515 | 192 | 2133 | 211 | 1390 | 166 | 1854 | 103 |
| Tpm4 | ENSMUSG00000031799 | 3181 | 181 | 4770 | 635 | 3020 | 369 | 4360 | 322 |
| Ttll1 | ENSMUSG00000022442 | 921 | 104 | 531 | 110 | 965 | 54 | 680 | 86 |
| Tuba1a | ENSMUSG00000072235 | 1839 | 223 | 2349 | 176 | 1762 | 189 | 2229 | 379 |
| Tubb2a | ENSMUSG00000058672 | 572 | 55 | 703 | 94 | 505 | 25 | 707 | 148 |
| Tubb2b | ENSMUSG00000045136 | 49 | 15 | 127 | 28 | 62 | 34 | 105 | 43 |
| Tubb5 | ENSMUSG00000001525 | 1852 | 114 | 2383 | 316 | 1680 | 188 | 2353 | 298 |
| Tubb6 | ENSMUSG00000001473 | 337 | 29 | 445 | 93 | 318 | 39 | 459 | 50 |
| Vim | ENSMUSG00000026728 | 3763 | 407 | 7380 | 1381 | 3521 | 520 | 6206 | 983 |
| Was | ENSMUSG00000031165 | 34 | 9 | 71 | 17 | 40 | 6 | 48 | 12 |
| Whrn | ENSMUSG00000039137 | 481 | 110 | 307 | 100 | 542 | 79 | 382 | 97 |
| Wipf1 | ENSMUSG00000075284 | 572 | 54 | 783 | 47 | 562 | 70 | 635 | 43 |
| Wisp1 | ENSMUSG00000005124 | 27 | 8 | 174 | 241 | 30 | 12 | 82 | 66 |
| Xirp2 | ENSMUSG00000027022 | 23026 | 1933 | 46990 | 12463 | 24085 | 3291 | 47862 | 16560 |
| Zyx | ENSMUSG00000029860 | 896 | 129 | 1214 | 76 | 876 | 143 | 1120 | 93 |

**Supplementary Table S11. Body weights of adult mice.** P values are for body weight after treatment with angiotensin II (AngII) for 7 d relative to baseline body weight (2-way ANOVA with Holm-Sidak's post-test) or for PKN2Het mice relative to wild-type (WT) mice at 42 weeks (t test). There were no significant differences between WT and PKN2Het mice at 12 weeks.

| **12 weeks** | **Baseline body weight (g)** | | | **Body weight after 7 d AngII (g)** | | | **P value** |
|---|---|---|---|---|---|---|---|
| | **Mean** | **SEM** | **n** | **Mean** | **SEM** | **n** | |
| WT/Vehicle | 28.18 | 0.75 | 5 | 28.24 | 0.72 | 5 | P=0.071 |
| PKN2Het/Vehicle | 28.25 | 0.69 | 8 | 27.80 | 0.80 | 7 | **P<0.001** |
| WT/AngII | 28.96 | 0.72 | 5 | 28.74 | 0.56 | 5 | P=0.6235 |
| PKN2/AngII | 29.42 | 0.76 | 8 | 28.47 | 0.72 | 7 | P=0.1847 |
| **42 weeks** | **WT** | | | **PKN2Het** | | | |
| | Mean | SEM | n | Mean | SEM | n | |
| | 35.10 | 1.40 | 8 | 37.24 | 0.67 | 11 | p=0.151 |

**Supplementary Table S12. qPCR primers.**

| **Gene** | **Sense Primer (5'→3')** | **Antisense Primer (5'→3')** |
|---|---|---|
| Gapdh | TCACCACCATGGAGAAGGC | GCTAAGCAGTTGGTGGTGCA |
| Myh7 | CATGCCAACCGTATGGCTG | GTTCCACGATGGCGATGTTC |
| Nppa | GATGGATTTCAAGAACCTGCTAGA | CTTCCTCAGTCTGCTCACTCA |
| Nppb | TCCAGCAGAGACCTCAAAATTC | CAGTGCGTTACAGCCCAAA |
| Tagln | GACTGCACTTCTCGGCTCAT | CCGAAGCTACTCTCCTTCCA |

**Supplementary References.**

1. Quetier I, et al. Knockout of the PKN Family of Rho Effector Kinases Reveals a Non-redundant Role for PKN2 in Developmental Mesoderm Expansion. *Cell Rep*. 2016;14:440-448. (Main Reference 12)
2 . Breckenridge R, Kotecha S, Towers N, Bennett M and Mohun T. Pan-myocardial expression of Cre recombinase throughout mouse development. *Genesis*. 2007;45:135-44. (Main Reference 22)
3. Stuckey DJ, Carr CA, Tyler DJ, Aasum E and Clarke K. Novel MRI method to detect altered left ventricular ejection and filling patterns in rodent models of disease. *Magn Reson Med*. 2008;60:582-7.
4. Geyer SH, Maurer-Gesek B, Reissig LF and Weninger WJ. High-resolution Episcopic Microscopy (HREM) - Simple and Robust Protocols for Processing and Visualizing Organic Materials. *J Vis Exp*. 2017.
5. Mohun TJ and Weninger WJ. Embedding embryos for high-resolution episcopic microscopy (HREM). *Cold Spring Harb Protoc*. 2012;2012:678-80.
6. Weninger WJ, et al. Visualising the Cardiovascular System of Embryos of Biomedical Model Organisms with High Resolution Episcopic Microscopy (HREM). *J Cardiovasc Dev Dis*. 2018;5.
7. Meijles DN, et al. Redox regulation of cardiac ASK1 (Apoptosis Signal-Regulating Kinase 1) controls p38-MAPK (mitogen-activated protein kinase) and orchestrates cardiac remodeling to hypertension. *Hypertension*. 2020;76:1208-1218. (Main Reference 23)
8. Ewels PA, Peltzer A, Fillinger S, Patel H, Alneberg J, Wilm A, Garcia MU, Di Tommaso P and Nahnsen S. The nf-core framework for community-curated bioinformatics pipelines. *Nature Biotechnology*. 2020;38:276-278.
9. Dobin A, Davis CA, Schlesinger F, Drenkow J, Zaleski C, Jha S, Batut P, Chaisson M and Gingeras TR. STAR: ultrafast universal RNA-seq aligner. *Bioinformatics*. 2013;29:15-21.
10. Li B and Dewey CN. RSEM: accurate transcript quantification from RNA-Seq data with or without a reference genome. *BMC Bioinformatics*. 2011;12:323.
11. Soneson C, Love MI and Robinson MD. Differential analyses for RNA-seq: transcript-level estimates improve gene-level inferences. *F1000Res*. 2015;4:1521.
12. Love MI, Huber W and Anders S. Moderated estimation of fold change and dispersion for RNA-seq data with DESeq2. *Genome Biology*. 2014;15:550.
13. Marshall AK, Barrett OPT, Cullingford TE, Shanmugasundram A, Sugden PH and Clerk A. ERK1/2 signaling dominates over RhoA signaling in regulating early changes in RNA expression induced by endothelin-1 in neonatal rat cardiomyocytes. *PLoSOne*. 2010;5:e10027.
14. Heinig M, et al. Natural genetic variation of the cardiac transcriptome in non-diseased donors and patients with dilated cardiomyopathy. *Genome Biology*. 2017;18:170. (Main Reference 21)

**Legends for video files.**

**Video 1.** HREM reconstruction of the heart from a day 14.5 *XMLC2Cre*$^{+/-}$ *Pkn2*$^{fl/fl}$ embryo. Video progresses to show long-axis sections through the heart and features include a large ventricular-septal defect with over-riding aorta, surface nodules which are ventricular diverticula, thin-walled ventricles and an overall abnormally squat shape.

**Video 2.** CT reconstruction of fixed heart and lungs from a 16-week *SM22αCre*$^{+/-}$ *Pkn2*$^{fl/fl}$ male mouse, with segmentation of the tissues: heart (pink, initially opaque, becoming partially transparent from 5 seconds), lungs (blue, partially transparent) and high-intensity signal from calcified plaques under the aortic and mitral valves (white solid).
